# Recording context and data splitting can inflate acoustic re-identification accuracy across species and studies

**DOI:** 10.64898/2026.10.08.757537

**Authors:** Graham Wallington, Jackie Lighten

## Abstract

1. Re-identifying individual animals (re-ID) from their calls with machine learning (ML) could extend individual-based monitoring to species and areas hard to survey by capture and marking, and published studies often report re-ID accuracy above 90%. Two methodological practices can inflate these values: (1) attributing calls to individuals by territory, nest, tag or station, tying each animal to its own recording conditions, and (2) randomly splitting individual clips into training and test sets, allowing recording context to be shared between them. Each effect has so far been documented only in single-species studies.
2. We implement four diagnostics of how methodological choices affect re-ID accuracy: (1) splitting clips by recording session; (2) background tests using ambient recordings without the animal; (3) within-recording permutation and paired-clip tests where animals share recordings; and (4) open-set tests with thresholds set on other individuals. We applied them to 13 public datasets from nine species (eight birds, one bat), using BirdNET, 19 other neural networks trained on animal or general sound and 16 trained on human speech, each with one classifier.
3. Under session splits, BirdNET re-ID accuracy ranged from 0.16 to 0.97. Within one year, ambient sound alone carried substantial identity information for territorial chiffchaffs (BirdNET balanced accuracy 0.71, against 0.79 for song); background identification exceeded chance for 35 of 38 embeddings and controls. Count-matched random clip splits raised mean accuracy by 0.20 to 0.51 in 18 comparisons across three species. Where several animals shared recordings, only one of three datasets showed clearly that re-ID rested on the animal rather than the recording. With unfamiliar animals present and thresholds calibrated to target 10% stranger acceptance, across 36 neural networks and two scoring methods on wild session-split datasets, the highest median share of known birds’ calls accepted and correctly named was 46%, using nearest-clip scoring with four enrolled birds; test-stranger acceptance could exceed the target. No neural network beat BirdNET on every dataset, and network rankings depended on layer choice.
4. The diagnostics need no new models and apply wherever a dataset holds what they require: session labels, background recordings, shared recordings or enough individuals. Re-ID accuracy reported without them can substantially overstate what acoustic re-ID achieves, and should not be used to plan population monitoring until they are applied.

## 1 | INTRODUCTION

Recognising the same individual on repeated occasions, or re-identification (re-ID), underpins estimates of abundance, survival and recruitment, as well as studies of dispersal between subpopulations, measures of territory tenure, mate choice, reproductive success, and the monitoring of rare, threatened or invasive species. Re-ID usually relies on capture and marking, genetic sampling or photographs of natural markings. These methods are labour intensive, can disturb animals, and limit how many individuals, and how large an area a study can cover.

Passive acoustic monitoring offers an alternative. Autonomous recorders run for weeks or months without an observer, and if individuals can be recognised reliably from their calls, the same recordings could estimate how many individuals use a site, whether they return and how long they survive. This is especially valuable for species that are difficult to catch or see, and for monitoring over larger areas and longer periods than mark-recapture allows. Calls differ consistently between individuals in many birds and mammals (Osiecka et al. 2024), in several species these differences persist across breeding seasons (Klenova et al. 2012; Takagi 2020; Calcari et al. 2021), and individual ovenbirds have been recognised from their songs over four years of passive recordings (Lapp et al. 2025).

Identifying species from recordings with machine learning (ML) is now routine (e.g., Kahl et al. 2021; Stowell 2022), but re-ID is harder. Differences between individuals are much finer than differences between species, and individual calls are not fixed: they change as animals mature and, in species that learn their calls, are shaped by the calls around them, so that they differ between populations and change as populations turn over (Podos & Warren 2007; Prat et al. 2017b; Vernes & Wilkinson 2020; Merino Recalde et al. 2025). Few recordings have been labelled with the identity of the calling animal, and in the wild a working system must cope with animals it has never encountered (Stowell 2022). Most current studies use a neural network pretrained on large collections of sound to convert each clip into an embedding, a list of numbers that summarises its acoustic properties, and then train a simple classifier to assign embeddings to known individuals (Ghani et al. 2023). In a closed-set test every test clip comes from an individual the classifier has learned; in an open-set test some come from strangers, animals the classifier has never learned, which it must recognise as unknown rather than wrongly name as one of the known individuals. Only the open-set test matches field monitoring, where many of the animals recorded will never have been heard before.

Reported re-ID accuracy is often high: 0.94 to 0.99 in five bird species (Huang et al. 2025a), 0.939 to 0.983 in seven bird species analysed separately (Gallego et al. 2026), and 0.84 in captive greater leaf-nosed bats when each echolocation call is classified on its own (Li et al. 2026). In a review of acoustic individual identification across all animal taxa, models had been trained on a mean of 20.3 individuals, and only 6.7% of studies had tested the open-set case, with strangers present (Knight et al. 2024).

The methods commonly used to test acoustic re-ID can, however, confound the results and artificially inflate estimated accuracy, in two ways. First, a classifier learns whatever acoustic differences best separate the labelled groups, whether or not they come from the animal. Every recording carries, besides the animal’s voice, the signature of where and how it was made: background sound, the acoustics of vegetation and terrain, the distance from animal to microphone and the recorder itself. The method used to establish which animal is calling usually fixes these conditions too: a territorial songbird is recorded on its territory, a nesting seabird at its nest, a tagged whale by its own tag. Identity and recording conditions then become the same variable (Figure 1a), and a classifier can score highly by recognising sites rather than individuals. Stowell et al. (2019) showed that ambient recordings containing no bird could identify the birds known to use each site, and Tolkova et al. (2026) found BirdNET embeddings identifying North Atlantic right whales at 0.84 from their calls and at 0.79 from tag noise containing no whale.

**Figure 1.**
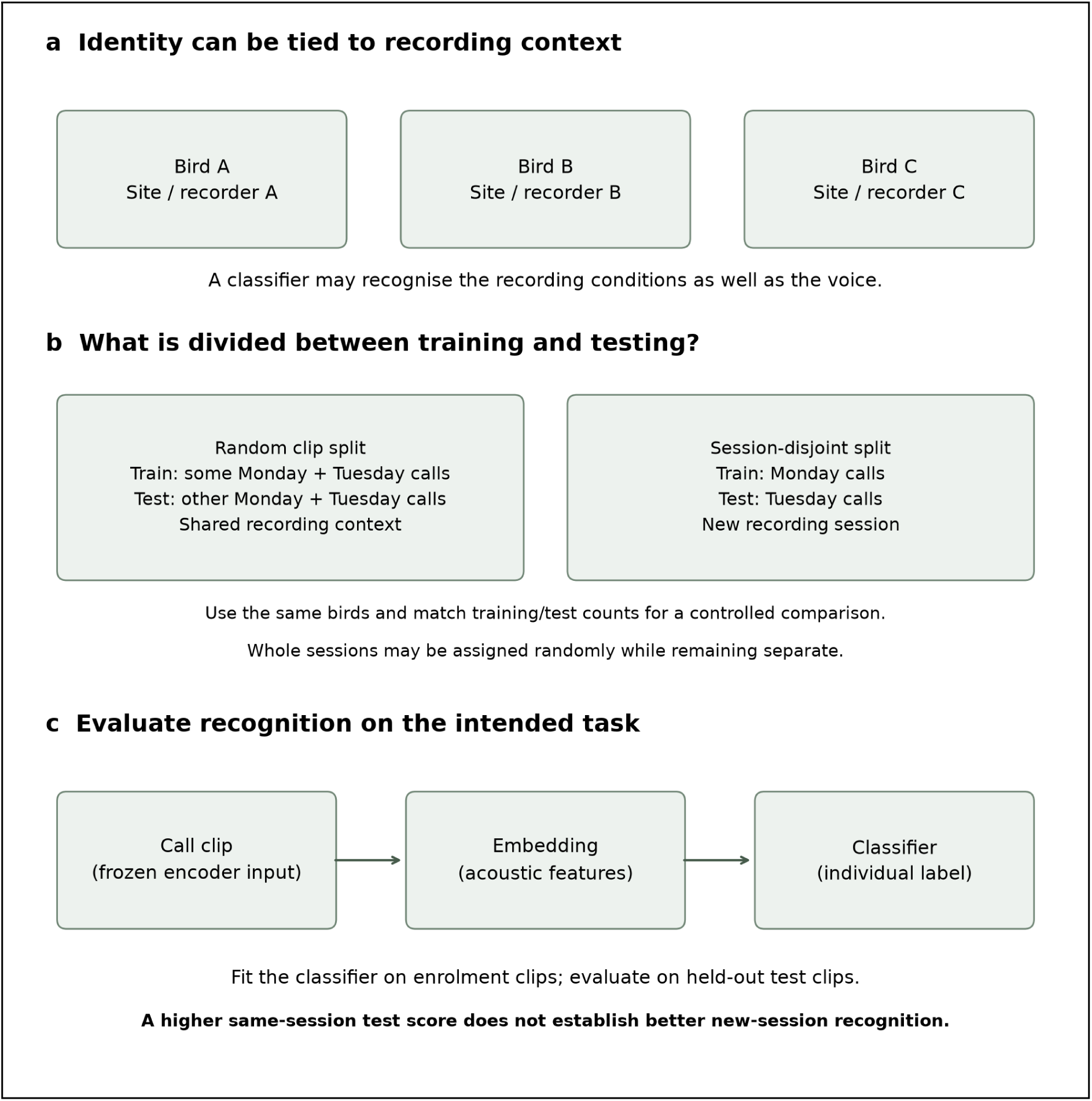
Recording context, split units and the re-ID test. (a) When each animal is associated with its own recording site or equipment, identity and recording conditions can covary. (b) Random allocation of clips can put calls from the same sessions in enrolment and test sets; keeping whole sessions apart tests transfer to a new session. Whole sessions can themselves be assigned randomly without mixing their clips. Monday and Tuesday are illustrative, not the actual timing of every dataset. (c) A frozen encoder converts a call to an embedding; a classifier fitted on enrolment clips assigns a held-out clip to an individual. Match the evaluation to the intended biological use.

Second, most published values come from random splits of clips or calls into training (here, enrolment) and test sets. The unit being split matters: random allocation of individual clips can put calls from the same recording session in both sets, whereas random allocation of whole sessions can keep them separate. For monitoring a known animal in a new recording session, enrolment and test sessions must be separate. For example, training on some of Monday’s calls and testing on other calls from Monday measures recognition within a familiar recording context; training on Monday and testing on Tuesday measures transfer to another session. A higher score on the first task is not evidence of improved performance on the second. Clips from one session can share background, equipment, position and closely related calls, so treating them as independent can overstate both accuracy for new sessions and its precision. Session separation alone does not remove site or equipment cues that persist between sessions. Gallego et al. (2026) note that their call-level splits do not keep sessions apart. Chiffchaff re-ID fell from above 0.85 to about 0.60 in area under the curve between within-year and across-year tests (Stowell et al. 2019), and a random clip split of bonobo calls gave 12.5 percentage points more than a split keeping related calls apart (Arnaud et al. 2023). Neither comparison used the pretrained embeddings now common in acoustic re-ID.

Here we test whether these single-species observations hold across species and studies. We present four diagnostics for evaluating acoustic re-ID, each targeting one route by which accuracy can be inflated or misread (Box 1), and apply them to 13 public datasets from nine species across seven studies. We ask: (1) how much a random split of individual clips inflates re-ID accuracy when embeddings and classifier are held constant; (2) whether the recording site alone can identify individuals; (3) where animals share recordings, whether re-ID survives tests that hold the recording fixed; and (4) how re-ID performs when unfamiliar animals are present. We also ask whether any of 35 other neural networks, trained on animal, general or human sound, is a better starting point than BirdNET, and how the answer depends on which layer of each network is read. We find that both problems are present in the datasets examined and can substantially inflate re-ID accuracy in birds, whichever neural network is used.

### Box 1. Four diagnostics for evaluating acoustic re-ID accuracy

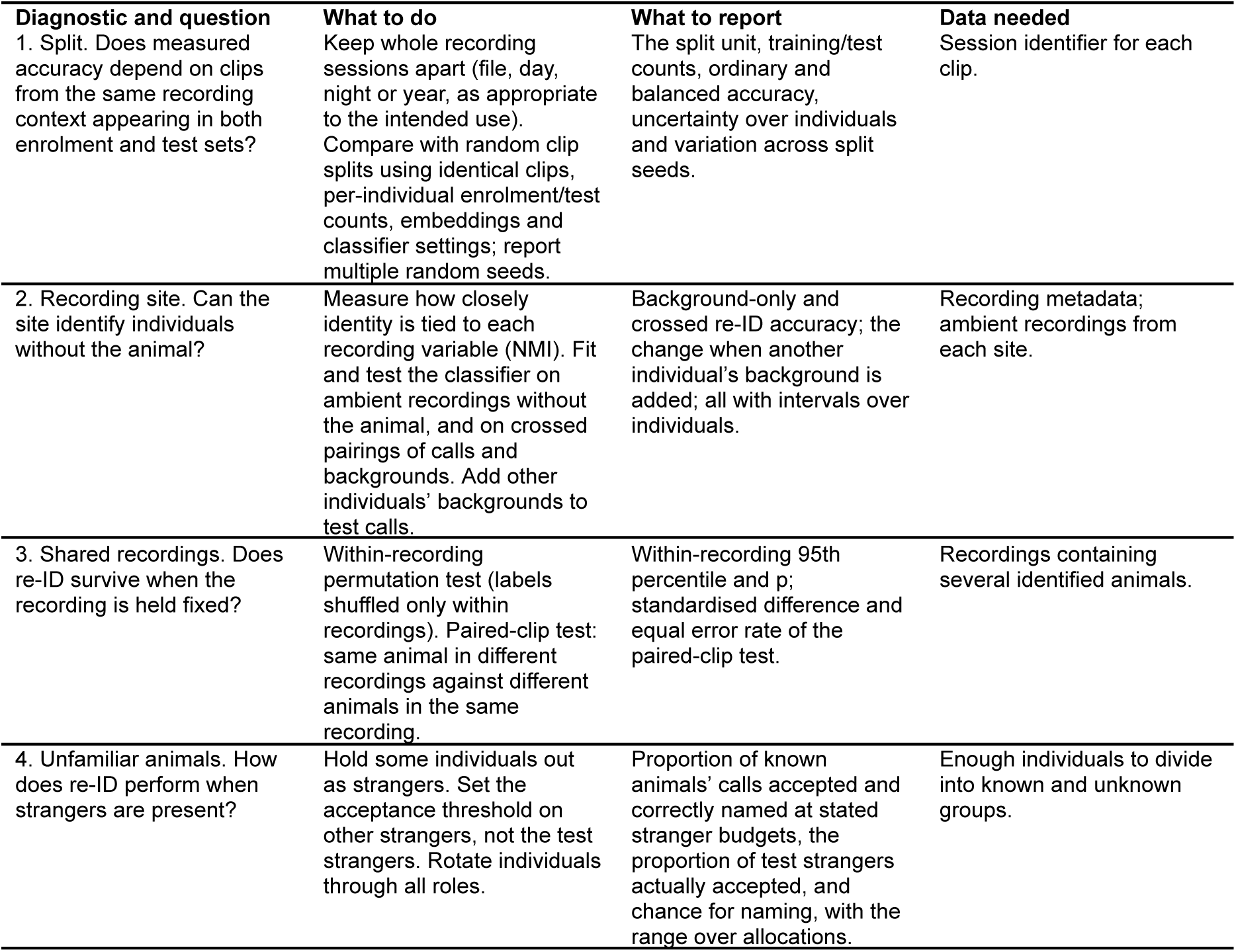

In all cases: report re-ID accuracy with 95% intervals over individuals (not clips), together with uniform chance, the majority-class rate and the 95th percentile of a label-permutation test. A value that exceeds the permutation test but not the majority-class rate has not shown that the classifier beats always naming the commonest individual.

## 2 | MATERIALS AND METHODS

### 2.1 | Overview

Every analysis followed the test in Figure 1c. Clips of known individuals were divided into enrolment clips, used to train the classifier, and test clips, used to measure re-ID accuracy, the proportion of test clips assigned to the correct individual. Each clip was converted to an embedding by a pretrained neural network used without retraining, and one fixed classifier was used for every closed-set test; the open-set test scored clips by similarity to each individual’s mean embedding (Section 2.8). A dataset is recordings of known individuals together with a fixed rule for dividing them into enrolment and test clips; the same animals divided within and across years give two datasets.

### 2.2 | Datasets

We used 13 publicly available datasets from nine species (Table 1), in which the identity of the animal producing each call had been established independently of any ML model, by an observer, the recording design or a sensor. They represent four recording designs that differ in how closely identity is tied to recording conditions (Figure 1a).

**Table 1.** The 13 datasets used in this study.

| Dataset | Species | Source | n | Enrolment / test clips | Recording design | Identity attributed by | Split by | Background recordings | NMI, identity and recording |
| --- | --- | --- | --- | --- | --- | --- | --- | --- | --- |
| Zebra finch, group of four | *Taeniopygia guttata* | Rüttimann et al. (2024) | 4 | 199 / 218 | shared microphone, captive | body-worn transmitter | recording file | no | 0.473 (session) |
| Zebra finch, group of eight | *T. guttata* | Rüttimann et al. (2024) | 8 | 260 / 229 | shared microphone, captive | body-worn transmitter | recording file | no | 0.407 (session) |
| Rook | *Corvus frugilegus* | Martin et al. (2022a, b) | 11 | 990 / 4,417 | shared microphones, captive aviary | leg rings, observer | year | no | 0.549 (recording) |
| Zebra finch, one bird per recording | *T. guttata* | Elie & Theunissen (2016, 2018) | 9 | 90 / 90 | one bird per recording, captive | recordist | recording | no | 0.827 (day) |
| Chiffchaff, within year | *Phylloscopus collybita* | Stowell et al. (2018, 2019) | 13 | 5,107 / 1,131 | own territory, wild | recordist, territory | recording day | yes | not recorded |
| Chiffchaff, across year | *P. collybita* | Stowell et al. (2018, 2019) | 10 | 324 / 200 | own territory, wild | recordist, territory | year | yes | not recorded |
| Little owl, across year | *Athene noctua* | Stowell et al. (2018, 2019) | 16 | 545 / 407 | own territory, wild | recordist, territory | year | yes | not recorded |
| Tree pipit, within year | *Anthus trivialis* | Stowell et al. (2018, 2019) | 10 | 409 / 303 | own territory, wild | recordist, territory | session within year | yes | not recorded |
| Tree pipit, across year | *A. trivialis* | Stowell et al. (2018, 2019) | 10 | 409 / 313 | own territory, wild | recordist, territory | year | yes | not recorded |
| Great tit | *Parus major* | Merino Recalde et al. (2024) | 16 | 160 / 160 | nest box, wild | ringed bird at nest box | year and nest box | no | 1.000 (recording) |
| Little penguin | *Eudyptula minor* | Huang et al. (2025a, b) | 15 (13 in test) | 2,429 / 792 | nest, wild | nest | night | no | 0.493 (day) |
| Red-tailed black cockatoo | *Calyptorhynchus banksii* | Huang et al. (2025a, b) | 16 | 1,428 / 357 | nest, wild | nest | random over clips | no | not recorded |
| Egyptian fruit bat | *Rousettus aegyptiacus* | Prat et al. (2017a) | 6 | 6,252 / 928 | one bat per recording, captive | synchronised video | year, day as session | no | 0.998 (recording); 0.136 (day) |

**Notes:** n, number of individuals enrolled; where fewer had test clips, that number follows in brackets. Recording design: whether each animal was recorded at its own site or several animals shared recordings. Identity attributed by: how the original authors established which animal produced each call; in most designs this also fixes where each animal was recorded. Split by: the recording session that separates enrolment from test clips. Background recordings: whether ambient recordings without the animal are available. NMI: mutual information between identity and the recording variable named, over all clips, divided by the smaller of the two entropies (0, unrelated; 1, either variable determines the other, as when every recording holds one animal); not recorded, the dataset does not identify recordings. The rook row is the dataset with 11 birds; the three chiffchaff and tree pipit rows and the little owl row are separate datasets from one study.

#### Own territory, wild (five datasets)

Stowell et al. (2018, 2019) recorded chiffchaffs, tree pipits and little owls, each on its own territory, divided into sessions within and across years. Beside every call they recorded ambient sound from the same place with no target bird calling (hereafter a background recording).

#### Nest, wild (three datasets)

The great tit dataset (Merino Recalde et al. 2024) comprises 16 ringed birds, each recorded at nest boxes in two of three years. In the little penguin and red-tailed black cockatoo datasets (Huang et al. 2025a, b), calls were attributed by nest; the penguin dataset is divided by night, and the cockatoo dataset offers only a random division of clips, so it serves as a reference for a random split. To test the great tit design on more birds, we built a second dataset from the same study with 50 birds and 1,000 songs, enrolled in one year and tested in the next (Appendix S1). In both great tit datasets a song is attributed to a bird through the nest attempt, not through an observation of the singer. The 50-bird dataset is reported separately and does not enter comparisons across datasets.

#### One animal per recording, captive (two datasets)

The zebra finch dataset (Elie & Theunissen 2016, 2018) comprises nine birds recorded one at a time. The Egyptian fruit bat dataset (Prat et al. 2017a) comprises six bats identified from synchronised video; recordings from 2012 and 2013 were used for enrolment and from 2014 for testing. No neural network used here analyses sound above 24 kHz (Table S12 gives each network’s limit), which excludes the upper part of these bats’ calls.

#### Shared microphone, captive (three datasets)

Several animals were recorded together, so the site and equipment were the same for all. In the BirdPark dataset (Rüttimann et al. 2024), group-housed zebra finches were recorded by one wall microphone and each call was attributed by a transmitter the bird carried; we used a group of four and a group of eight juveniles, divided by recording file. The rook dataset (Martin et al. 2022a, b) comprises ringed birds in one outdoor aviary recorded on 23 days across two years; we used one microphone channel, 2020 recordings for enrolment and 2021 recordings for testing, the 11 birds with at least 20 calls in each year, and at most 90 enrolment clips per bird (Appendix S1).

In every dataset except the cockatoo, enrolment and test clips came from different recording sessions: a recording file, day, night or year depending on the dataset (Table 1). Datasets screened and not used are listed in Table S20.

### 2.3 | Embeddings and classifier

Clips were used as supplied, cut by the original authors around individual calls or songs. We compared 38 embeddings and controls.

#### BirdNET

BirdNET v2.4 (Kahl et al. 2021), a neural network trained to identify bird species, was the reference against which every other neural network was compared. For each 3-s segment of a clip we took the output of the layer before the species predictions, an embedding of 1,024 numbers, and averaged the segments (Appendix S1). Such embeddings are widely used as general-purpose descriptions of animal sound (Ghani et al. 2023).

#### Other neural networks trained on animal or general sound

To ask whether any is a better starting point than BirdNET, we added 19, covering the main published designs (Table S12), including Perch 2.0 (van Merriënboer et al. 2025) and two AVEX networks (Miron et al. 2026). Each clip was brought to the network’s own sample rate, cut into windows of the network’s own input length and passed through the network’s published preprocessing, and the window embeddings were averaged. For networks built from a stack of repeated processing blocks (transformers), the output of every block was also a candidate embedding. BirdNET and these networks were given the sound at its recorded level, except that BirdPark clips were multiplied by one common gain per endpoint (0.95 divided by its peak), preserving relative levels within that endpoint but not absolute levels. Nine of them take sound sampled at 16 kHz and so analyse it only up to 8 kHz; unlike the speech networks, they were not also given slowed playback (Table S12).

#### Speech neural networks

Recognising a person from their voice is the human counterpart of acoustic re-ID, so we included 16 neural networks pretrained on human speech (Table S12). Three are speaker-recognition networks, trained to tell people apart regardless of what they say. Twelve were trained on unlabelled speech, two of which (data2vec Base, 100 h and 960 h) were then fine-tuned to transcribe speech, and XEUS (Chen et al. 2024) on unlabelled speech in more than 4,000 languages; networks of this kind distinguish individual marmosets without retraining (Sarkar & Magimai-Doss 2023), and the same design has been retrained on animal sound (Hagiwara 2023). They analyse sound only up to 8 kHz, so each clip was also played at one half and one third of its speed, bringing sound up to 24 kHz into range. Each clip was scaled to a peak of one, so these networks could not use a clip’s absolute level. Every layer, at each playback speed, was a candidate embedding. A speaker-verification checkpoint of WavLM Base+ gave values identical to WavLM Base+ and is not counted (Appendix S1).

#### Choosing among candidate embeddings

Choosing the layer with the test clips would inflate accuracy, so the layer and playback speed of every network were chosen on the enrolment calls alone, as the candidate with the greatest separation between individuals relative to variation within them (the Fisher ratio). When each animal’s enrolment clips come from its own recordings, the most separating layer may be the one that best separates recordings.

Where the data allowed it, we therefore also chose each layer by a second rule: each individual’s enrolment sessions were dealt into five groups, the classifier was fitted on four and tested on the fifth, and the candidate with the highest accuracy on sessions it had not been fitted on was chosen (Appendix S1). This rule needs at least two enrolment sessions per individual, which six of the 13 datasets and the 50-bird great tit dataset have. After review, we also read every network at its own final output: the output embedding of networks trained on animal or general sound, and the last layer at the recording’s own speed for speech networks. This third rule was specified after the results were seen and is exploratory.

#### Controls

A three-number description of clip loudness and the clip’s duration were passed through the same classifier in place of an embedding. If either identifies individuals, re-ID could rest on trivial differences such as distance to the microphone or how clips were cut.

#### Classifier

We used a deliberately simple classifier with no tuned settings, following standard practice for evaluating pretrained embeddings (Ghani et al. 2023): kernel ridge regression, which learns from the enrolment embeddings a weighted combination of embedding values that predicts each individual, with a fixed penalty against fitting noise. Two classifiers that need no fitting checked that results did not depend on this choice: class mean (the individual whose enrolment clips a test clip resembles most on average) and nearest clip (the individual that produced the most similar enrolment clip).

### 2.4 | Measuring re-ID accuracy

Clips from one animal, and especially from one recording, are not independent, so headline 95% intervals were obtained by resampling individuals with replacement 10,000 times. Initial model-sweep diagnostics used 2,000 draws and 999 label permutations; the final paired network comparisons used 10,000 shared draws, and the headline and background-test nulls used 9,999 permutations. Where recordings were identified, a three-stage interval also resampled recordings within individuals and clips within recordings. All analyses used a fixed random seed so that every value can be reproduced, and Holm’s correction was applied to the network-comparison and background-test families identified below (Holm 1979). The permutation p values in Table 2 are unadjusted.

**Table 2.** Closed-set re-ID accuracy with BirdNET and the kernel ridge classifier.

| Dataset | n | Split by | Re-ID accuracy (95% interval) | Three-stage interval | Majority class | Permutation 95th percentile | p | Within-recording permutation 95th percentile (p) |
| --- | --- | --- | --- | --- | --- | --- | --- | --- |
| Zebra finch, group of four | 4 | recording file | 0.807 (0.669 to 0.963) | 0.613 to 0.963 | 0.445 | 0.491 | 0.0001 | 0.587 (0.0001) |
| Zebra finch, group of eight | 8 | recording file | 0.498 (0.176 to 0.725) | 0.124 to 0.754 | 0.358 | 0.293 | 0.0002 | 0.362 (0.0002) |
| Rook, across year | 11 | year | 0.375 (0.260 to 0.510) | 0.250 to 0.529 | 0.241 | 0.127 | 0.0001 | 0.224 (0.0001) |
| Zebra finch, one bird per recording | 9 | recording | 0.967 (0.933 to 1.000) | 0.906 to 1.000 | 0.111 | 0.244 | 0.0001 | 0.856 (0.0002) |
| Chiffchaff, within year | 13 | recording day | 0.765 (0.681 to 0.846) | n/a | 0.393 | 0.154 | 0.0001 | n/a |
| Chiffchaff, across year | 10 | year | 0.160 (0.035 to 0.333) | n/a | 0.160 | 0.190 | 0.1327 | n/a |
| Little owl, across year | 16 | year | 0.501 (0.371 to 0.634) | n/a | 0.084 | 0.123 | 0.0001 | n/a |
| Tree pipit, within year | 10 | session within year | 0.376 (0.193 to 0.566) | n/a | 0.125 | 0.162 | 0.0001 | n/a |
| Tree pipit, across year | 10 | year | 0.214 (0.116 to 0.334) | n/a | 0.160 | 0.157 | 0.0022 | n/a |
| Great tit, across year | 16 | year and nest box | 0.444 (0.319 to 0.575) | 0.297 to 0.594 | 0.062 | 0.106 | 0.0001 | n/a |
| Little penguin, across night | 15 | night | 0.693 (0.604 to 0.772) | 0.594 to 0.782 | 0.136 | 0.115 | 0.0001 | n/a |
| Red-tailed black cockatoo, random split | 16 | random over clips | 0.961 (0.929 to 0.980) | n/a | 0.216 | 0.182 | 0.0001 | n/a |
| Egyptian fruit bat, across year | 6 | year, day as session | 0.381 (0.189 to 0.569) | 0.188 to 0.565 | 0.297 | 0.239 | 0.0001 | n/a |

*Notes:* The 95% interval resamples individuals 10,000 times; the three-stage interval also resamples recordings within individuals and clips within recordings, where the dataset records them. Majority class: accuracy of always naming the individual with the most test clips. Permutation: 95th percentile and p value from 9,999 shuffles of the enrolment labels (0.0001 is the smallest attainable p). Within-recording permutation: labels shuffled only among clips of the same recording (for the zebra finch recorded singly, the same day). n/a: the dataset records no recordings, or too few hold two individuals for the test to change the labels (fruit bat: 16 of 6,252 enrolment clips could be shuffled). The rook enrolment was capped at 90 clips per bird.

Each re-ID accuracy was compared with three reference levels, each representing a classifier with no knowledge of the animals’ voices: uniform chance (one divided by the number of individuals); the majority-class rate (always naming the individual with most test clips, which matters because individuals rarely call equally often); and a label-permutation test, in which enrolment labels were shuffled 9,999 times for the headline analyses and the classifier refitted each time, giving a 95th percentile and p value. Passing the permutation test shows that embeddings predict the label, not whether they do so through the animal or the recording; the diagnostics below address that question. Beecher’s information statistic (Beecher 1989), the measure of identity information used in studies of vocal individuality, is reported for every embedding in Table S18.

### 2.5 | Diagnostic 1: how the data are split

To compare random clip and session-disjoint splits, we held the clip pool, each individual’s enrolment and test counts, embeddings, preprocessing and classifier settings constant. Using the published BirdNET and Perch embeddings of Huang et al. (2025b), we retained only the recordists’ train/test clips for chiffchaff, little owl and tree pipit. Enrolment/test counts were respectively 5,107/1,131, 545/407 and 409/303. Each individual’s clips were shuffled using ten predetermined seeds (101–110), preserving its counts on both sides. The three classifiers were refitted under each split; no test data chose a classifier or parameter. Table 3 reports the mean accuracy across seeds and a paired interval from 10,000 shared draws of individuals, conditional on these ten splits; seed ranges and balanced accuracies are reported in Appendix S2. This follow-up comparison was specified after review. The original published-fold results remain in Appendix S2: their little-owl fold used 761 enrolment clips, rather than 545, and their tree-pipit fold included additional clips. They therefore changed training size or clip composition as well as session overlap. Descriptive nearest-enrolment similarities were also computed, but do not establish near-duplication or its causal contribution. The rook comparison retains the original count-matched random division within each bird against enrolment in 2020 and testing in 2021. The cockatoo endpoint in Table 2 follows metadata.csv fold labels; these differ from the published embedding-file labels and are not interchangeable.

**Table 3.** Effect of the data split on re-ID accuracy, with embeddings and classifier unchanged.

| Species | Embeddings | Classifier | Random clip split (mean) | Session split | Difference (95% interval) |
| --- | --- | --- | --- | --- | --- |
| Little owl | BirdNET | kernel ridge | 0.963 | 0.477 | 0.486 (0.359 to 0.606) |
| Little owl | BirdNET | class mean | 0.897 | 0.430 | 0.467 (0.343 to 0.582) |
| Little owl | BirdNET | nearest clip | 0.973 | 0.577 | 0.396 (0.249 to 0.546) |
| Little owl | Perch | kernel ridge | 0.964 | 0.631 | 0.333 (0.201 to 0.479) |
| Little owl | Perch | class mean | 0.879 | 0.543 | 0.336 (0.203 to 0.466) |
| Little owl | Perch | nearest clip | 0.963 | 0.582 | 0.381 (0.229 to 0.537) |
| Chiffchaff | BirdNET | kernel ridge | 0.969 | 0.734 | 0.235 (0.147 to 0.315) |
| Chiffchaff | BirdNET | class mean | 0.810 | 0.502 | 0.308 (0.219 to 0.425) |
| Chiffchaff | BirdNET | nearest clip | 0.941 | 0.595 | 0.346 (0.253 to 0.434) |
| Chiffchaff | Perch | kernel ridge | 0.947 | 0.751 | 0.196 (0.120 to 0.270) |
| Chiffchaff | Perch | class mean | 0.622 | 0.404 | 0.218 (0.128 to 0.276) |
| Chiffchaff | Perch | nearest clip | 0.818 | 0.486 | 0.332 (0.225 to 0.389) |
| Tree pipit | BirdNET | kernel ridge | 0.927 | 0.419 | 0.508 (0.328 to 0.688) |
| Tree pipit | BirdNET | class mean | 0.782 | 0.363 | 0.418 (0.291 to 0.548) |
| Tree pipit | BirdNET | nearest clip | 0.865 | 0.406 | 0.459 (0.325 to 0.592) |
| Tree pipit | Perch | kernel ridge | 0.910 | 0.452 | 0.458 (0.275 to 0.635) |
| Tree pipit | Perch | class mean | 0.717 | 0.277 | 0.440 (0.339 to 0.543) |
| Tree pipit | Perch | nearest clip | 0.771 | 0.356 | 0.415 (0.263 to 0.569) |
| Rook, shared aviary | BirdNET (this study) | kernel ridge | 0.547 | 0.422 | 0.126 (0.065 to 0.172) |

Notes: For chiffchaff, little owl and tree pipit, random-clip accuracy is the mean over seeds 101–110, using exactly the recordists' clip pool and preserving each individual's enrolment/test counts. Differences are mean random-clip minus session-disjoint accuracy. Intervals use 10,000 paired draws of individuals and condition on the ten fixed splits; Appendix S2 reports seed ranges, balanced accuracies and the original published-fold comparisons. The rook retains its original single count-matched split, without the Table 2 enrolment cap. None of these differences isolates a particular acoustic mechanism.

### 2.6 | Diagnostic 2: can the recording site identify individuals?

We first measured how closely identity was tied to each recording variable stored with the clips (recording file, day, channel, site) using normalised mutual information (NMI), the mutual information divided by the smaller of the two entropies. NMI is 0 when the variable says nothing about which animal is calling and 1 when either variable determines the other, as when each recording contains one animal but also when each animal belongs to one year; counts of levels holding several individuals, and of individuals on several levels, show which (Table S1). For both great tit datasets, whose splits moved birds between years, we checked whether any nest box or recording contributed to both a bird’s enrolment and test clips, and measured the distance between them.

The five Stowell et al. datasets include background recordings, so the site could be tested directly, for all 38 embeddings and controls. The classifier was fitted on calls or backgrounds and tested on calls or backgrounds, giving four pairings. Backgrounds tested on backgrounds asks whether the site alone identifies the bird; the two crossed pairings (fitted on calls and tested on backgrounds, as in Stowell et al. 2019, and the reverse) ask whether the classifier uses the same features for call and site. For the little owl, an exploratory equivalence test, with a margin set after the background estimate was seen, asked whether background-only accuracy lay close to chance (Appendix S1).

The four pairings show whether the site carries identity; an added-background test shows whether the classifier uses it on real calls. With the classifier fitted on the original enrolment clips, each test clip was mixed with a background from its own individual or from another (the donor, the next individual in sorted order), with the background 10 dB louder than the clip, at equal level or 10 dB quieter. Both backgrounds came from the test side of the split, so in the across-year datasets from the test year. If the classifier relies on the site, another individual’s background should pull answers towards that individual. We report the paired difference in balanced accuracy (the mean of per-individual accuracies) between the two mixtures. How often BirdNET’s errors named the donor, compared with how often its errors on the same clips without any added background named that individual, was an exploratory analysis specified after results were seen. The same sorted-order donor mapping was used throughout. Intervals resample recipient individuals conditional on that mapping; they do not measure sensitivity to the choice of donor. Rotating donors would be a separate robustness experiment and was not performed.

A failed background test does not show that no individual signal exists. To measure what the test can detect, we planted calls from ten within-year chiffchaffs into the across-year chiffchaff backgrounds, one bird per territory, at 10 dB above, equal to and 10 dB below the background, and scored them as a normal dataset. The planted calls were recorded on the birds’ own territories, so each carries its original site as well as the bird, and the test measures detection of the two together.

#### Remedies after recording

We tested three ways of reducing the site’s contribution once recordings exist, each scored through the four pairings (Appendix S1). Background augmentation (Stowell et al. 2019) mixes every enrolment call with a background from each other individual, so that site no longer separates individuals; it was applied to the within-year chiffchaff, the little owl and the within-year tree pipit. Spectral subtraction removes from every clip the average spectrum of its individual’s background recordings; because it uses each test clip’s true identity, it asks only whether the site can be removed when every test clip’s site is known, and a rise in re-ID accuracy cannot be credited to the method. Adaptive score normalisation, used in speaker recognition, rescales each score against the scores the same clip gives to a set of background recordings. We also tested four session-compensation methods from human speaker recognition on the fruit bat, measuring how well the recording day could still be predicted.

#### Right whale

To compare with Tolkova et al. (2026), the four pairings were also run on their right whale calls and tag noise, in which each whale’s clips come from one tag recording (Appendix S1); these data are not among the 13 datasets.

### 2.7 | Diagnostic 3: tests that hold the recording fixed

Where recordings contain several individuals, the recording itself can be held fixed; we counted such recordings in each dataset. In a within-recording permutation test, labels were shuffled only among clips of the same recording, so each recording still predicts its own set of labels; a classifier that recognises recordings passes the ordinary permutation test but fails this one. It was applied, for every embedding, to the rook and both BirdPark groups, with recording files as the unit, and to the zebra finch recorded one bird at a time, with recording days as the unit. In the fruit bat, recordings almost never held two bats, and the little penguin’s test clips record only a date, across nests, so neither test is reported for them. In a paired-clip test (Appendix S1), we compared the similarity of animal pairs (two clips of one individual from different recordings) with recording pairs (clips of two individuals from the same recording). An embedding that captures the animal rates animal pairs as more similar. We report the standardised difference in mean cosine similarity (divided by the square root of the mean of the two population variances) and the equal error rate for telling the two kinds of pair apart from a single pair (0.5, no separation; 0, perfect separation). The paired-clip equal error rates were recomputed from the original cached embeddings with corrected tie handling; all reported values in Tables S3 and S4 were unchanged (Appendix S1.8). These are the only tests of recording context available for shared-microphone datasets, which have no background recordings.

### 2.8 | Diagnostic 4: unfamiliar animals present (open-set test)

On the great tit, the 50-bird great tit, the little owl, the cockatoo and the rook datasets, individuals were divided into four roles, known (enrolled) and unknown (strangers), each for calibration and for testing, over 16 allocations in which every individual held every role (Appendix S1). Each role held four individuals (two or three in the rook, 12 or 13 in the 50-bird great tit). Each known individual’s profile was the mean of its enrolment embeddings, and each test call was scored by its highest cosine similarity to any profile. A call was accepted if its score exceeded a threshold and was then assigned to the best-matching individual. For stranger budgets of 1 to 50% (the maximum proportion of strangers the system may accept), thresholds were set on the calibration strangers and applied to the test animals, as in practice (calibrated threshold), and we report the proportion of test strangers they actually accepted. The best threshold among those accepting at most the budgeted proportion of test strangers (test-derived threshold) cannot be set in practice; it is an upper limit only when the calibrated threshold accepts no more strangers. We report how often the correct known individual was named among the four known individuals of an allocation (chance 0.25), separation (the probability that a known individual’s call outscores a stranger’s; 0.5, none), the equal error rate, the proportion of known individuals’ calls accepted and correctly named, and, following the AnimalCLEF re-ID benchmark (Adam et al. 2025), balanced accuracy on known individuals and on strangers and their geometric mean, as medians and ranges over allocations. The test is reported for all 36 neural networks. Duration and loudness controls are excluded from this cosine-based analysis because unit normalisation removes their magnitude; their kernel-ridge results are retained. For the great tit, we also scored year and nest-box position alone, without audio, and BirdNET on clips with every note of the song removed.

In a follow-up specified after review, we also used each call’s nearest enrolment clip as its score and assigned that clip’s identity, recalibrating thresholds separately for this scorer while retaining all roles and frozen embeddings. The original mean-profile results are retained in Table 6 and Table S15; Figure 6 and Appendix S2 compare both scorers. The 50-bird great-tit calibration sensitivity used fixed nested subsets of 4, 8 and 12 calibration strangers with galleries and test roles unchanged. Neither scorer nor calibration size was chosen on test performance.

### 2.9 | Comparing neural networks

On the 13 datasets, each of the 37 other embeddings and controls was compared with BirdNET over the same 10,000 bootstrap draws of individuals, with Holm’s correction across the 37 comparisons on each dataset, separately for each layer rule. The two-sided bootstrap p calculation is given in the Table S13 note. The held-out rule was summarised only on the six datasets where it can choose among layers; on the others it chooses what the Fisher rule chooses. Consistency of ranking across datasets was tested with Friedman’s test, Kendall’s W and Nemenyi’s critical difference (Demšar 2006).

### 2.10 | Reproducibility

Every measurement was run from version-controlled code on one computer, with each run’s code version, clip lists, splits, random seed and computing environment recorded in its result file, which is archived under a checksum of its contents. Statistics added after review were computed from the same result files on a second computer, after the previous version’s tables and figures had been reproduced there exactly. The 16 speech network checkpoints of an earlier run were run again, and every reported value was identical between the two runs (Appendix S1, Table S19). Before the original tables were produced, an evidence gate checked the presence, provenance and role-specific contracts of the 224 result files used by the paper. It checked resampling counts for most roles, but not the corrected-sweep, speech, selection-refresh and speech-replay roles; all figures and tables were then produced from those files by scripts that record the source of every value. Analyses were specified in the project’s internal repository before being run; those specified after results were seen are labelled exploratory.

## 3 | RESULTS

### 3.1 | Re-ID accuracy under session splits

Under session splits, re-ID accuracy with BirdNET ranged from 0.160 for the chiffchaff across a year (95% interval 0.035 to 0.333) to 0.967 for the zebra finch recorded one bird at a time (0.933 to 1.000; Table 2, Figure 2). All datasets except the chiffchaff across a year exceeded the 95th percentile of their permutation test (chiffchaff: 0.160 against 0.190, p = 0.1327). Fruit bats were re-identified at 0.381 (0.189 to 0.569) against a majority-class rate of 0.297, and the cockatoo, divided at random, at 0.961 (0.929 to 0.980). In seven of 13 datasets the permutation 95th percentile lay below the majority-class rate, so passing the permutation test alone did not show that the classifier beat always naming the commonest individual. In the 50-bird great tit dataset, BirdNET re-identified birds at 0.270 (0.202 to 0.342) against uniform chance of 0.020 (p = 0.0001; Table S3).

**Figure 2.**
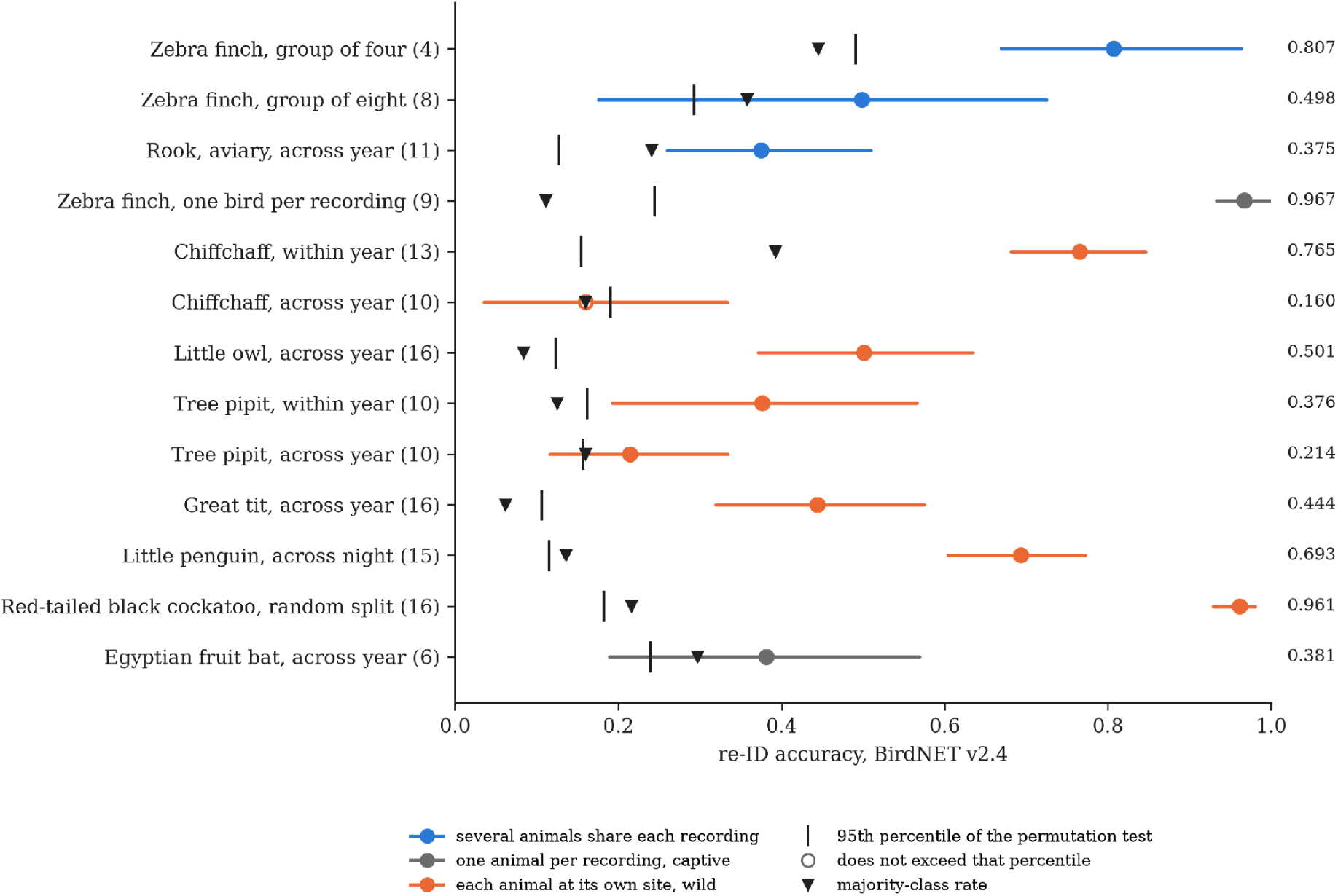
Closed-set re-ID accuracy with BirdNET on 13 datasets; the number of individuals is in parentheses. Points: re-ID accuracy; bars: 95% interval over individuals; vertical ticks: 95th percentile of the label-permutation test; triangles: majority-class rate. Open circles: values that do not exceed that percentile. Colour gives the recording design.

### 3.2 | Diagnostic 1: random clip splits raised measured accuracy

With clip pools and per-individual enrolment/test counts matched, random clip splits raised mean measured accuracy in all 18 comparisons across three species, two neural networks and three classifiers, by 0.196 to 0.508. Every paired individual-bootstrap interval excluded zero (Table 3, Figure 3). For little owl with BirdNET and kernel ridge, session-disjoint accuracy was 0.477 and the mean random-clip accuracy was 0.963 (seed range 0.953 to 0.973), a difference of 0.486 (0.359 to 0.606). The largest mean difference, 0.508, was for tree pipit with BirdNET and kernel ridge, using the same 712 clips and the same 409/303 enrolment/test counts in both arms. These are changes in measured accuracy under different testing conditions, not demonstrated improvements in recognition in new sessions. Median similarity to the closest enrolment clip from the same individual was higher in all six species–embedding combinations under every random split, consistent with greater within-block acoustic similarity but not a direct test of near-duplication. In the rook, the original count-matched random division raised accuracy from 0.422 to 0.547 (difference 0.126, 0.065 to 0.172); it also mixed the two recording years.

**Figure 3.**
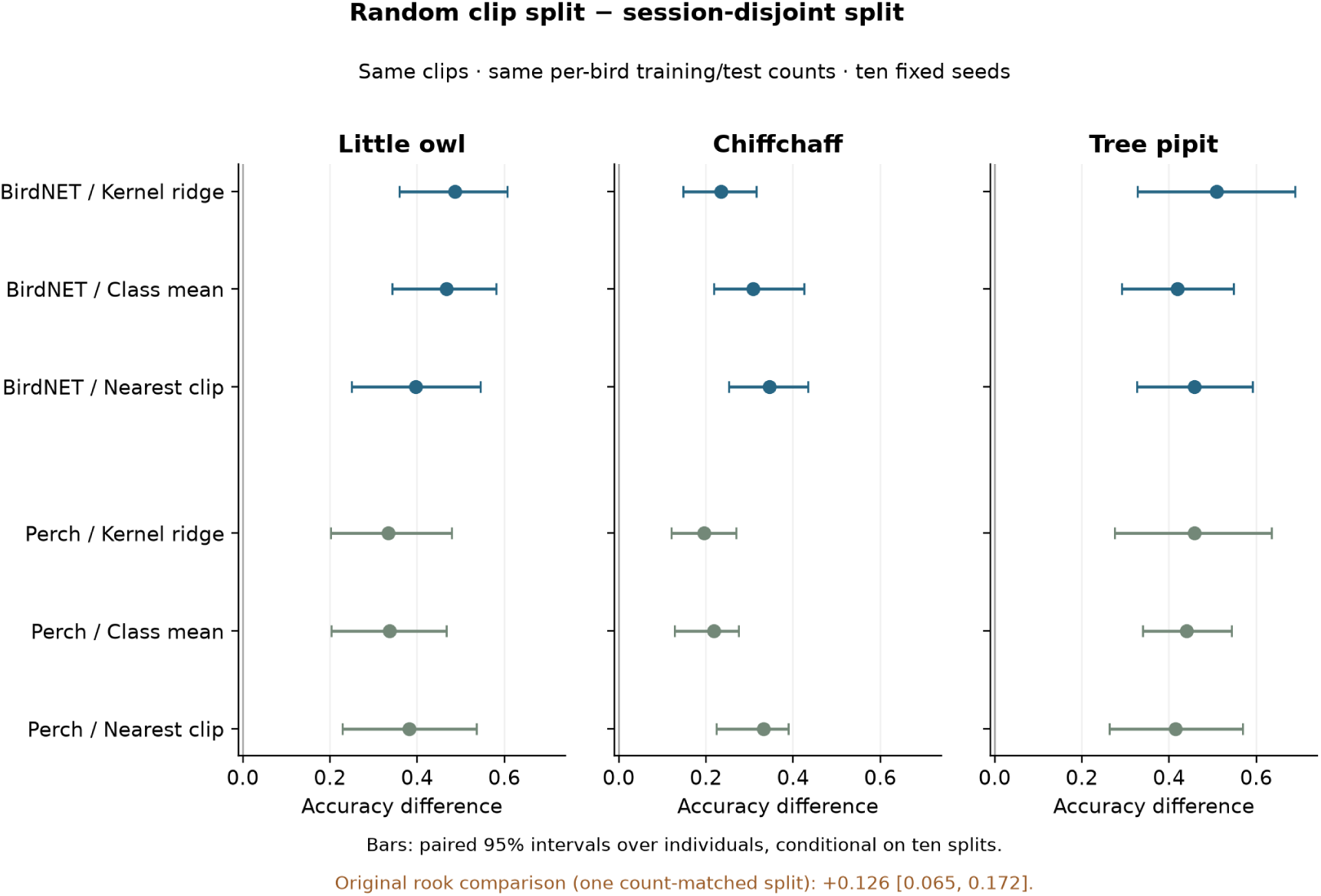
Count-matched random clip splits versus session-disjoint splits. Points show the mean accuracy difference across ten predetermined random splits, for three species, two published embeddings and three classifiers. Both arms use identical clips and retain each individual’s enrolment/test counts. Bars show paired 95% individual-bootstrap intervals, conditional on those ten splits; seed ranges are reported in Appendix S2. The orange rook annotation retains the original single count-matched random split. Positive differences describe the easier testing setting, not improved recognition in unseen sessions.

### 3.3 | Diagnostic 2: the recording site identified some individuals

Each recording held one animal in the great tit (NMI 1.000) and all but 12 of 7,109 recordings held one bat in the fruit bat (0.998), and recording day was closely tied to identity in the zebra finch recorded singly (0.827; Table 1, Table S1). Great tits were nevertheless tested at nest boxes 52 to 229 m (median 84.9 m) from their enrolment boxes, with no nest box or recording shared between enrolment and test (Table S2). In the 50-bird great tit dataset, no recording was shared, but 21 of the 50 birds were enrolled and tested at the same nest box (median distance 29.2 m, range 0 to 733 m).

For the chiffchaff within one year, a classifier fitted and tested on background recordings, which contain no bird, re-identified birds at 0.737 (0.599 to 0.813), against 0.765 for calls (uniform chance 0.077, p = 0.0001; Table 4, Figure 4). Balanced accuracy, which gives each bird equal weight, was 0.714 for backgrounds and 0.791 for calls. The corresponding query-majority reference rates were 0.395 and 0.393; these describe the most common test identity, rather than a classifier fitted on enrolment data. Thus the background result remains substantial after accounting for imbalance, but the balanced call and background accuracies should not be described as equivalent. Both crossed pairings also exceeded chance: calls-trained classifiers named birds from backgrounds at 0.456 (p = 0.0001), and backgrounds-trained classifiers named them from calls at 0.508 (p = 0.0001). When each test call was mixed with a background 10 dB louder than itself, it was named correctly 0.553 of the time with its own bird’s background and 0.217 with another bird’s (difference in balanced accuracy 0.325, 0.194 to 0.465; Table 5, Figure 5). In an exploratory analysis, 24.9% of BirdNET’s errors named the bird whose background had been added, against 1.9% of its errors on the same clips with no background added (Table S7). This result did not depend on the neural network. Backgrounds alone identified chiffchaffs within a year for 35 of the 38 embeddings and controls, all but AudioMAE and the two controls, also after Holm’s correction across the 38, and adding another bird’s background moved answers towards that bird (interval above zero with the background 10 dB louder) for 26 of 38 (Tables S5, S6). The four networks with the highest call-based accuracy on this dataset, Perch 2.0 (0.831), AudioProtoPNet (0.806), ConvNeXt (0.805) and Perch (0.778), were also the four with the highest background-only accuracy (0.787, 0.815, 0.835 and 0.795; Table S5).

**Figure 4.**
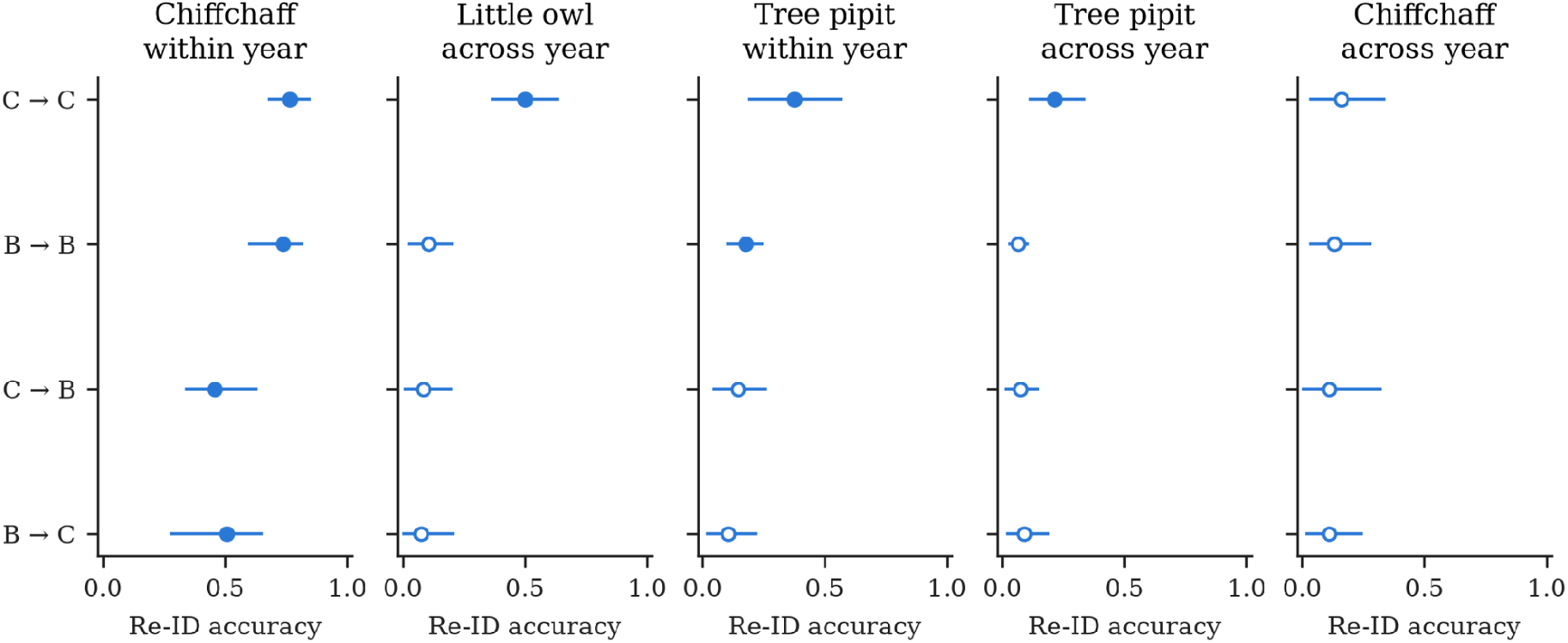
Diagnostic 2: re-ID from calls and from ambient background recordings with BirdNET on the five datasets that include backgrounds. Each panel shows four pairings of what the classifier was fitted on and tested on (C, calls; B, backgrounds; C→B, fitted on calls and tested on backgrounds). Filled circles: p < 0.05 in the permutation test; open circles: not. Bars: 95% interval over individuals.

**Figure 5.**
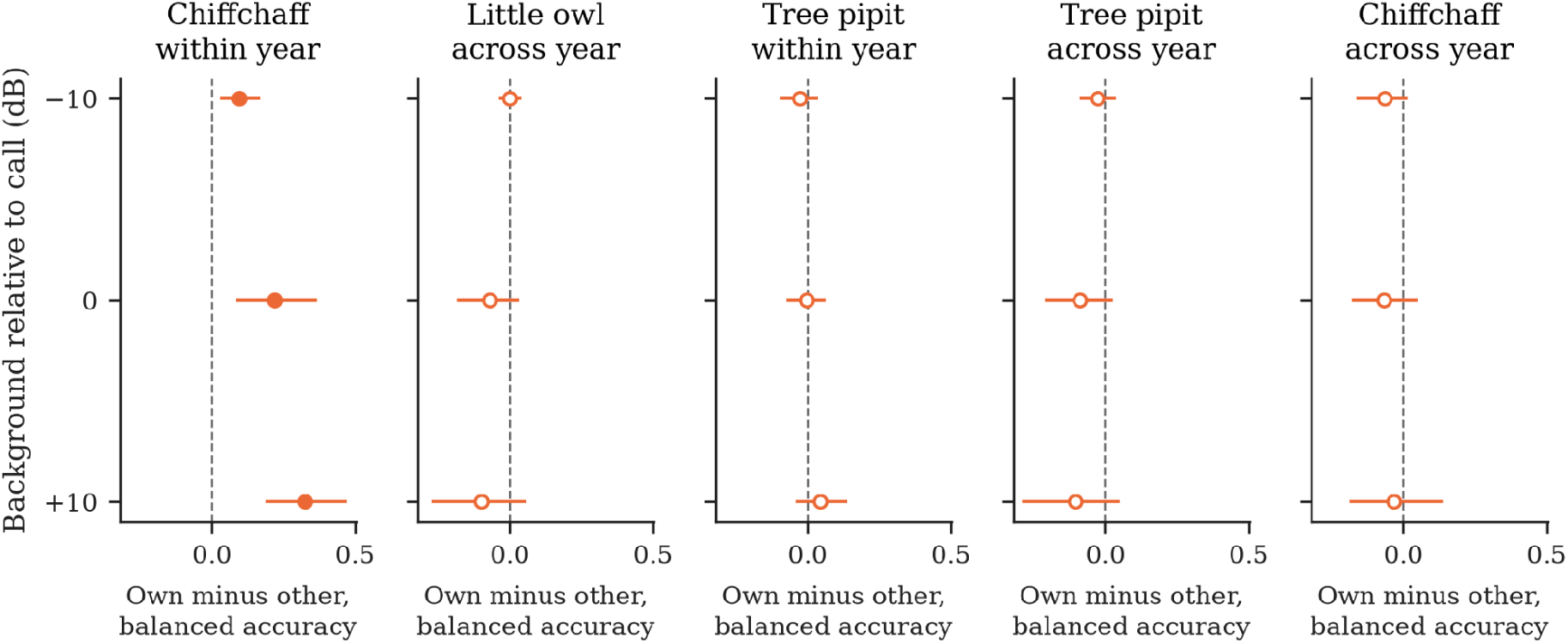
Diagnostic 2: effect of adding another animal’s background, with BirdNET. Difference in balanced accuracy between test clips mixed with a background from their own individual and from another individual, at three levels of the added background relative to the call (+10 dB: background 10 dB louder than the call), as in Table 5. Positive values mean the classifier’s answers follow the background. Bars: 95% interval over individuals; filled circles: intervals excluding zero.

**Figure 6.**
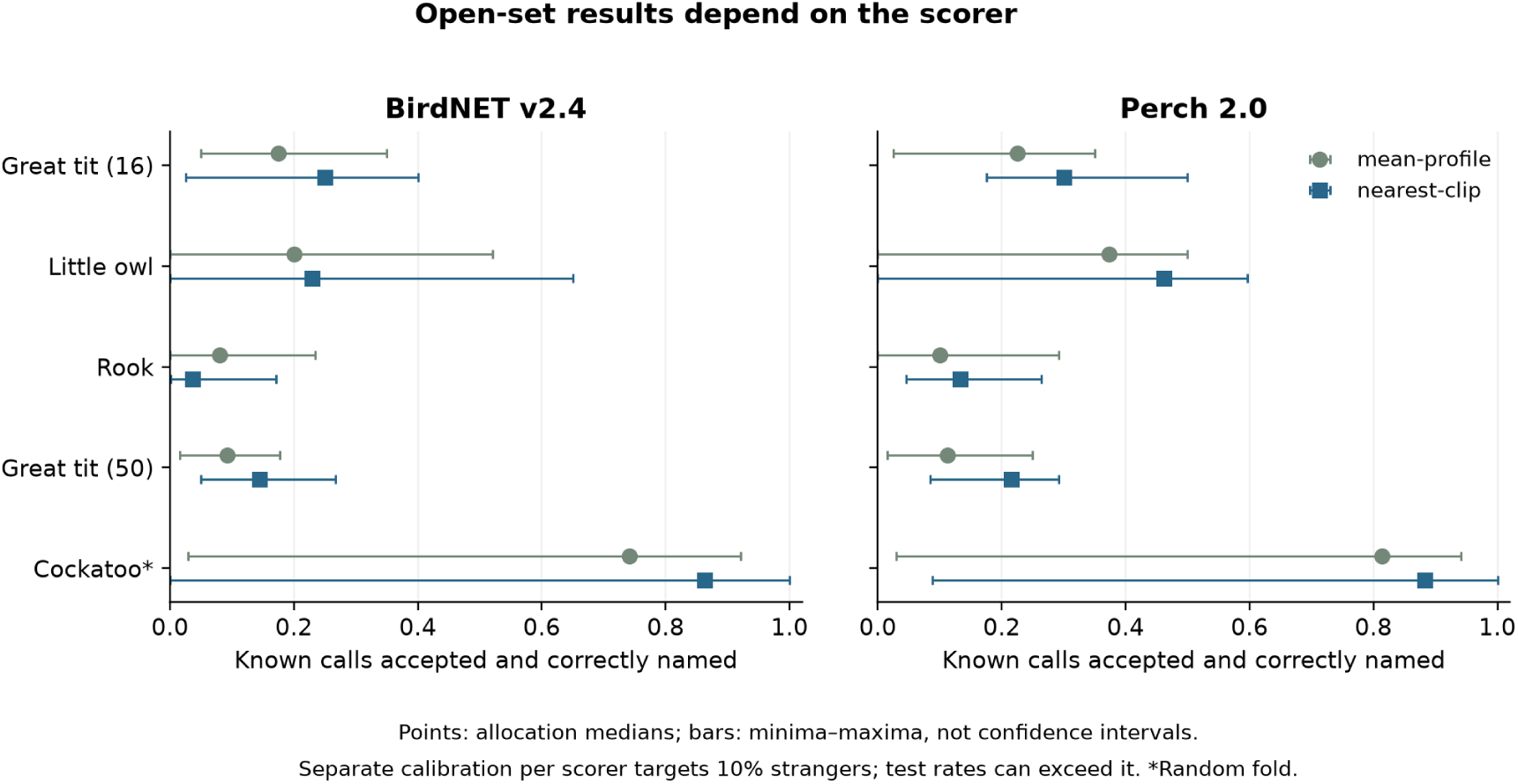
Diagnostic 4: comparison of mean-profile and nearest-enrolment-clip open-set scoring for BirdNET and Perch 2.0. Points show the median proportion of known birds’ calls accepted and correctly named over the same 16 role allocations; lines show allocation minima and maxima, not confidence intervals. Each scorer sets its own threshold using separate calibration strangers with a 10% target. Test-stranger acceptance can exceed that target; Appendix S2 reports the realised rates. The cockatoo uses a published random fold; the other endpoints keep recording sessions apart. Table 6 retains the original mean-profile diagnostics.

**Table 4.**
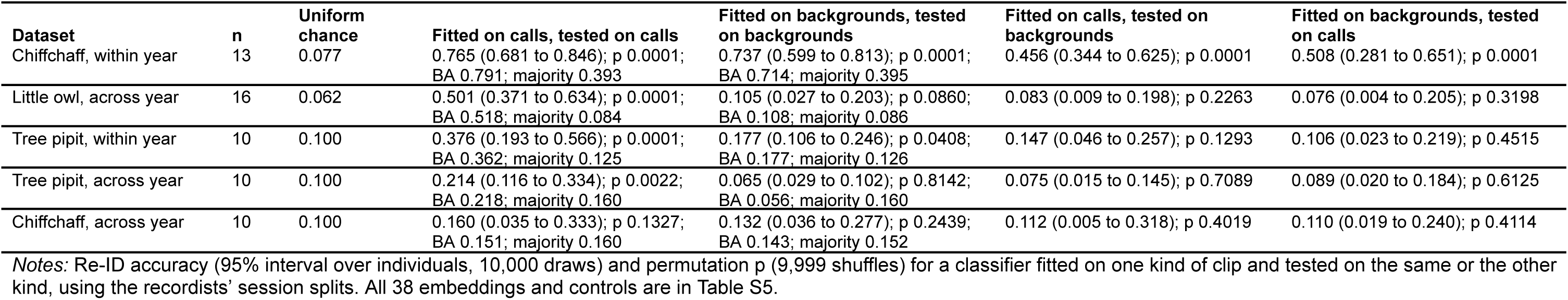
Re-ID from calls and from ambient background recordings, with BirdNET. BA denotes balanced accuracy (mean per-bird recall). Majority denotes the proportion of test clips belonging to the commonest test identity, separately for calls and backgrounds; it is a descriptive reference, not a trained baseline. Appendix S2 also reports the accuracy of always naming the enrolment-majority identity.

**Table 5.**
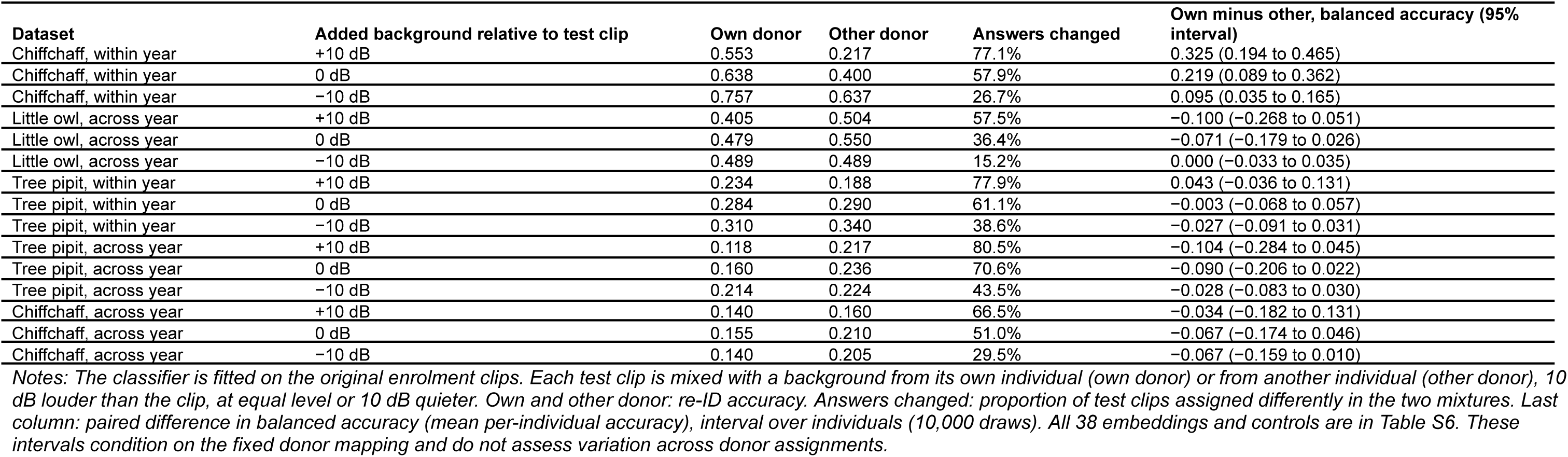
Effect of adding a background recording to test clips, with BirdNET.

**Table 6.** Re-ID with unfamiliar animals (strangers) present using the original mean-profile scorer: medians over 16 allocations of individuals to roles, with ranges. Figure 6 and Appendix S2 report the alternative nearest-clip scorer.

| Dataset | Embedding | Names correctly | Chance for naming | Separation | Equal error rate | Accepted and named, test-derived threshold | Accepted and named, calibrated threshold | Strangers accepted, calibrated threshold |
| --- | --- | --- | --- | --- | --- | --- | --- | --- |
| Great tit, across year | BirdNET | 0.600 (0.375 to 0.700) | 0.250 | 0.617 (0.466 to 0.733) | 0.413 | 0.200 (0.150 to 0.300) | 0.175 (0.050 to 0.350) | 0.087 (0.000 to 0.350) |
| Great tit, across year | Perch 2.0 | 0.625 (0.400 to 0.700) | 0.250 | 0.616 (0.497 to 0.667) | 0.425 | 0.213 (0.100 to 0.350) | 0.225 (0.025 to 0.350) | 0.087 (0.000 to 0.275) |
| Great tit, across year | BirdNET, song removed | 0.512 (0.225 to 0.700) | 0.250 | 0.548 (0.374 to 0.744) | 0.475 | 0.100 (0.050 to 0.275) | 0.100 (0.025 to 0.350) | 0.087 (0.000 to 0.325) |
| Great tit, across year | Year and nest box only, no audio | 0.500 (0.500 to 1.000) | 0.250 | 0.625 (0.375 to 1.000) | 0.250 | 0.000 (0.000 to 0.500) | 0.125 (0.000 to 0.750) | 0.250 (0.000 to 1.000) |
| Great tit, 50 birds, across year | BirdNET | 0.348 (0.275 to 0.450) | 0.077 to 0.083 | 0.545 (0.414 to 0.740) | 0.471 | 0.092 (0.008 to 0.154) | 0.092 (0.015 to 0.177) | 0.092 (0.025 to 0.275) |
| Great tit, 50 birds, across year | Perch 2.0 | 0.436 (0.223 to 0.508) | 0.077 to 0.083 | 0.549 (0.430 to 0.718) | 0.460 | 0.117 (0.031 to 0.285) | 0.112 (0.015 to 0.250) | 0.103 (0.025 to 0.277) |
| Little owl, across year | BirdNET | 0.659 (0.480 to 0.892) | 0.250 | 0.691 (0.436 to 0.881) | 0.332 | 0.159 (0.009 to 0.634) | 0.200 (0.000 to 0.520) | 0.084 (0.000 to 0.697) |
| Little owl, across year | Perch 2.0 | 0.751 (0.500 to 0.957) | 0.250 | 0.743 (0.468 to 0.880) | 0.330 | 0.287 (0.047 to 0.743) | 0.374 (0.000 to 0.500) | 0.100 (0.000 to 0.587) |
| Rook, across year | BirdNET | 0.611 (0.429 to 0.879) | 0.333 to 0.500 | 0.523 (0.408 to 0.679) | 0.495 | 0.071 (0.006 to 0.330) | 0.080 (0.000 to 0.234) | 0.100 (0.000 to 0.503) |
| Rook, across year | Perch 2.0 | 0.575 (0.401 to 0.842) | 0.333 to 0.500 | 0.610 (0.417 to 0.904) | 0.412 | 0.094 (0.003 to 0.733) | 0.101 (0.000 to 0.291) | 0.100 (0.000 to 0.403) |
| Red-tailed black cockatoo, random split | BirdNET | 0.980 (0.899 to 1.000) | 0.250 | 0.850 (0.632 to 0.997) | 0.239 | 0.599 (0.231 to 0.936) | 0.741 (0.029 to 0.921) | 0.082 (0.000 to 0.584) |
| Red-tailed black cockatoo, random split | Perch 2.0 | 0.968 (0.879 to 1.000) | 0.250 | 0.908 (0.642 to 1.000) | 0.162 | 0.761 (0.198 to 0.963) | 0.814 (0.029 to 0.941) | 0.085 (0.000 to 0.632) |

**Notes:** Names correctly: proportion of known individuals' test calls assigned to the correct individual among the known individuals of the allocation, with no strangers present. Chance for naming: one over the number of known individuals in an allocation (a range where allocations differ). Separation: probability that a known individual's call outscores a stranger's (0.5, none). Accepted and named: proportion of known individuals' test calls accepted and assigned to the correct individual. Test-derived threshold: the best value over all thresholds that accept at most one test stranger in ten, set with the test strangers themselves, so it is an upper limit only when no more than 10% of strangers are accepted. Calibrated threshold: set on other strangers to accept one in ten, as would be done in practice; the last column gives the proportion of test strangers it actually accepted. Perch 2.0 had the best average rank in the closed-set comparison. The great tit rows without audio and with the song removed come from the same allocations. All 36 neural networks are in Table S15; duration and loudness controls are excluded from this cosine-based analysis.

For the little owl, no pairing involving backgrounds exceeded its permutation test with BirdNET (background-only 0.105, p = 0.0860), and an exploratory equivalence test failed: the upper 95% bound of background-only accuracy was 0.182, above the bound of 0.1065. After Holm’s correction across the 38 embeddings and controls, background-only accuracy exceeded its permutation test for 1 on the little owl (SurfPerch), for none on the across-year chiffchaff and tree pipit, and for 8 on the within-year tree pipit, including Perch 2.0 (0.256) but not BirdNET (0.177, unadjusted p = 0.0408). On these four datasets, adding another bird’s background moved answers towards that bird for at most 3 of 38 embeddings at any level (Table S6). Clips mixed with their own bird’s background were named less accurately than with another’s for 5 to 7 of 38 embeddings on the little owl and up to 9 on the across-year tree pipit; these backgrounds came from the test year, and a classifier fitted on enrolment calls did not identify little owls from them (0.083, p = 0.2263; Table 4).

A failed background test does not show that no individual signal exists. When calls of ten chiffchaffs were planted into the across-year backgrounds, one bird per territory, BirdNET re-identified them at 0.641 (0.562 to 0.825, p = 0.0001) with the call 10 dB above the background, 0.362 (0.231 to 0.646, p = 0.0003) at equal level, below the majority-class rate of 0.436, and 0.171 (0.055 to 0.448, p = 0.1208) with the call 10 dB below the background (Table S8). Because the planted calls carry their own territory’s sound as well as the bird, these values describe detection of call and original site together.

On the right whale data of Tolkova et al. (2026), BirdNET embeddings of tag noise identified the 11 whales within their own tag recordings at 0.556 to 0.750 (p = 0.0001 at every frequency shift); at the shift chosen in advance neither crossed pairing exceeded chance, and after Holm’s correction over the 11 shifts the pairing fitted on noise and tested on calls did so at 9 (Table S17).

None of the remedies applied after recording removed the site’s contribution. Background augmentation raised call-based re-ID for the within-year chiffchaff from 0.765 to 0.814, and lowered the accuracy with which calls-trained classifiers named birds from backgrounds from 0.456 to 0.260, still above chance (p = 0.0001; Table S9); it also raised call-based re-ID for the little owl (0.501 to 0.560) and the within-year tree pipit (0.376 to 0.587). Spectral subtraction, which uses each test clip’s true identity, raised call-based re-ID on four of five datasets, for example from 0.214 to 0.358 for the tree pipit across a year, and lowered it on the within-year chiffchaff (0.765 to 0.721); these changes cannot be credited to the method, and on the within-year chiffchaff background-only re-ID remained 0.681 after it (Table S9). Across the neural networks, score normalisation often lowered call-based re-ID on the within-year chiffchaff while raising background-only re-ID (Table S10). The raw duration and loudness controls are excluded from this cosine-based analysis. In the fruit bat, no session-compensation setting removed day information while keeping identity: settings that kept re-ID accuracy above 0.33 left day prediction at 0.26 to 0.28 (majority rate 0.166), and the descriptive Spearman correlation across the 16 distinct settings was 0.616 (p = 0.0111), falling to 0.477 (p = 0.0843) when the two session-mean settings were excluded (Table S11). The significance therefore depends on settings whose day-prediction behaviour is partly imposed by their construction; these settings share the same bats and are not independent biological replicates.

### 3.4 | Diagnostic 3: holding the recording fixed

Most recordings contained two or more animals in the shared-microphone datasets (7 of 7 sessions in the group of four zebra finches, 6 of 7 in the group of eight, 67 of 81 rook recordings). In the group of four, re-ID accuracy (0.807, 0.669 to 0.963) exceeded the within-recording 95th percentile (0.587, p = 0.0001), and animal pairs were more similar than recording pairs (standardised difference 1.675), enough to tell single pairs apart with an equal error rate of 0.173. In the group of eight and the rook, re-ID also exceeded the within-recording 95th percentile (0.498 against 0.362; 0.375 against 0.224), but animal and recording pairs were equally similar in the group of eight (standardised −0.033) and only slightly different in the rook (0.260; equal error rate 0.449). Across days within 2020 the rook’s standardised difference was 0.036, and changing the microphone channel while keeping calls identical did not lower re-ID accuracy (0.405 against 0.395; Table S4). Of the 38 embeddings and controls, 33 exceeded the within-recording test in the group of four, 31 in the group of eight and 25 in the rook (Table S21). A follow-up descriptive timing check used every eligible pair of retained BirdNET test embeddings (Appendix S2). All same-bird, different-recording pairs in the two BirdPark groups were more than seven days apart, so a same-day control cannot be constructed from these clips. In the rook, same-day animal pairs had mean cosine similarity 0.748, compared with 0.731 for recording pairs. These dependent-pair means do not supply an independent significance test or remove all temporal and behavioural differences.

For the zebra finch recorded one bird at a time, labels could be shuffled only among days with more than one bird (6 of 15 enrolment days); re-ID accuracy of 0.967 exceeded the within-day 95th percentile of 0.856. In the fruit bat, only 8 of 6,188 enrolment recordings held more than one bat, so the within-recording test could shuffle only 16 of 6,252 enrolment clips, and only five pairs of test clips came from two bats in one recording, too few for the paired-clip test. The little penguin’s test clips record only a date, and the birds of one date were recorded at different nests, so no pair of its clips shares a recording. Diagnostic 3 therefore could not be applied to the fruit bat or the little penguin.

### 3.5 | Diagnostic 4: few calls were accepted and correctly named with strangers present

With the original mean-profile scorer and thresholds calibrated on other animals to admit one stranger in ten, a median of 0.175 of known great tits’ calls, 0.200 of little owls’ and 0.080 of rooks’ were accepted and correctly named by BirdNET, although among the four known birds of an allocation alone (two or three for the rook; chance 0.25, and 0.33 to 0.50 for the rook) the correct bird was named 0.600, 0.659 and 0.611 of the time (Table 6; mean-profile points in Figure 6). Perch 2.0, the neural network with the best average rank in the closed-set comparison (Section 3.6), accepted and correctly named 0.225, 0.374 and 0.101. Across the 36 neural networks, the highest medians on these three datasets were 0.225, 0.374 and 0.119 (Table S15). The calibrated thresholds accepted a median of 8% to 10% of test strangers, but up to 70% in single allocations (little owl, BirdNET), so a threshold set on one group of strangers did not transfer reliably to another, and results varied widely between allocations (little owl, BirdNET: 0.000 to 0.520). The test-derived threshold is an upper limit only where the calibrated threshold keeps within the same stranger budget. A calibrated threshold that exceeds that budget can accept and correctly name more known calls while also admitting more strangers. On the four datasets split by recording session, known birds’ calls outscored strangers’ in 0.52 to 0.74 of trials for BirdNET and Perch 2.0. Balanced accuracy on strangers was high (0.87 to 0.92) because most strangers were rejected, while balanced accuracy on known birds was low (0.06 to 0.37), giving geometric means of 0.22 to 0.48 (Table S15). Only on the cockatoo, divided at random, were more than half of known birds’ calls accepted and correctly named (BirdNET 0.741, Perch 2.0 0.814). In the 50-bird great tit dataset, with 12 or 13 known birds (chance about 0.08), 0.092 (BirdNET) and 0.112 (Perch 2.0) were accepted and correctly named. For the great tit, BirdNET applied to clips with the song removed still named the correct bird 0.512 of the time, and year and nest-box position alone, without audio, separated known birds from strangers better than the audio did (equal error rate 0.250 against 0.413).

With nearest-clip scoring, medians for BirdNET were 0.250 (16-bird great tit), 0.229 (little owl), 0.037 (rook), 0.144 (50-bird great tit) and 0.863 (cockatoo); corresponding Perch 2.0 medians were 0.300, 0.462, 0.133, 0.216 and 0.883 (Figure 6, Appendix S2). Across both tested scorers and all 36 distinct neural networks, the highest median on the wild session-disjoint endpoints was 0.462, for little owl with Perch 2.0 and nearest-clip scoring (allocation range 0.000 to 0.596). Its actual test-stranger acceptance had median 0.099 and range 0.000 to 0.615. The scorer therefore matters, and these results are not a ceiling on possible open-set performance.

The nested calibration-size check did not show monotonic improvement or reliable transfer of the 10% budget: every size exceeded it in some allocations (Appendix S2).

### 3.6 | Choosing a neural network

Under the Fisher rule, of 481 comparisons between the 37 other embeddings and controls and BirdNET on 13 datasets, 9 were above BirdNET after correction, 183 below and 289 not distinguishable (Figure 7, Table S13). Perch 2.0 had the best average rank, 4.1 of 38, and was above BirdNET on three datasets: the tree pipit within a year (0.548 against 0.376) and across a year (0.447 against 0.214) and the rook (0.418 against 0.375). It was not detectably below BirdNET on any dataset. Perch was above BirdNET for the little owl (0.646 against 0.501) and the tree pipit across a year (0.351 against 0.214). Speech neural networks were above BirdNET only for the little penguin, where two data2vec networks, WavLM Base+ and XEUS scored 0.794 to 0.798 against 0.693; for the speech networks the Fisher rule chose layer 0, the input to the first transformer layer, in 125 of 169 cases. The ranking was moderately consistent across datasets (Friedman χ² = 224.7, p < 0.0001; Kendall’s W = 0.47). BirdNET ranked 6.3 on average. With Nemenyi’s critical difference of 16.8 ranks, the rank test separated BirdNET only from the ten lowest-ranked entries, which included both controls, Bird-MAE, AudioMAE and VGGish (Table S14).

**Figure 7.**
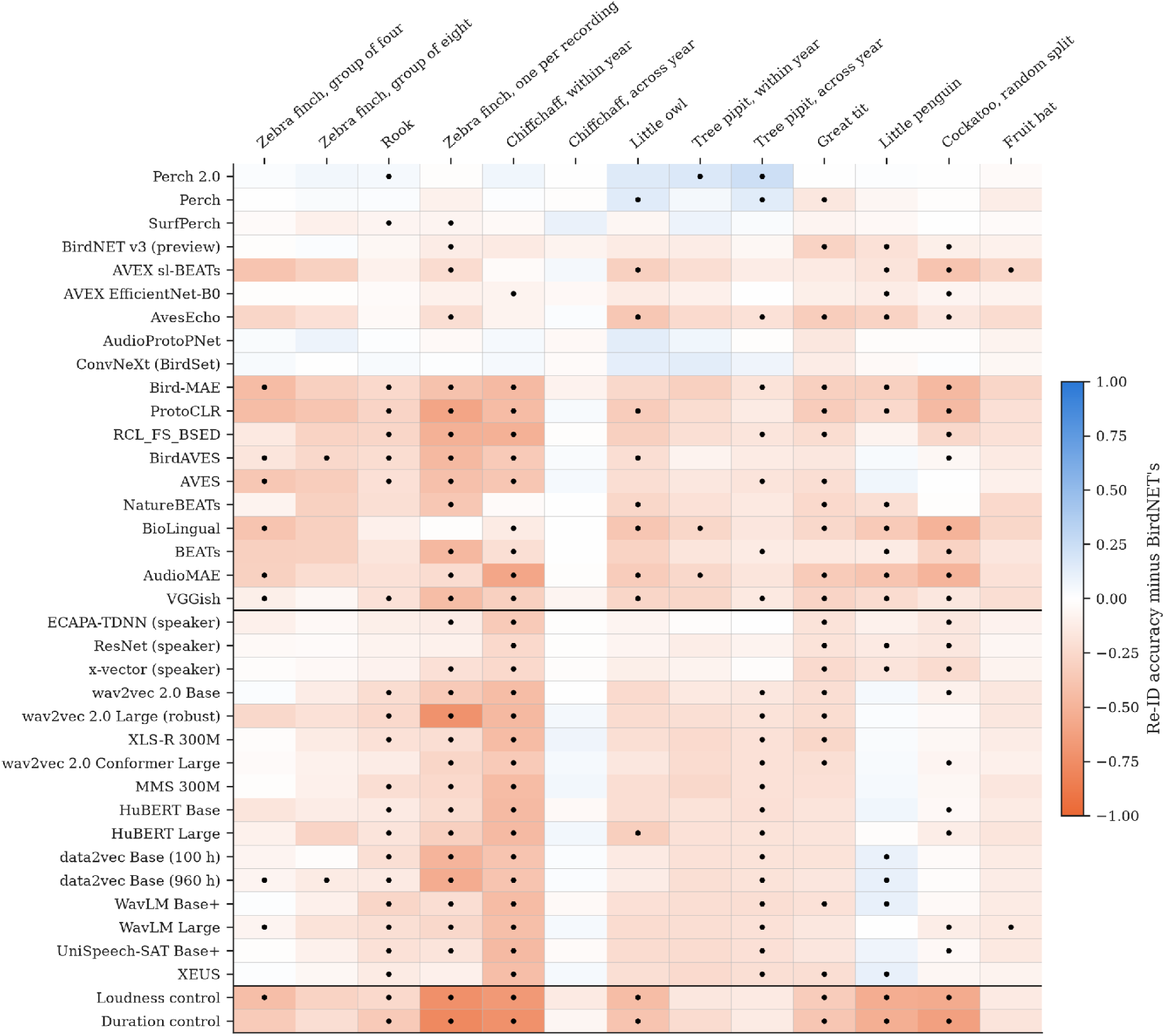
Thirty-five neural networks and two controls compared with BirdNET on 13 datasets, through the same classifier, at the layer (and, for speech networks, playback speed) chosen on enrolment clips by the Fisher rule. Colour: re-ID accuracy minus BirdNET’s. Dots: differences that remain after Holm’s correction across the 37 comparisons on that dataset. Lines separate networks trained on animal or general sound, speech networks and the two controls.

On the six datasets where the held-out rule applies, choosing each layer on enrolment sessions held out from the classifier changed the chosen layer in 130 of 222 comparisons; re-ID accuracy rose in 82, fell in 45 and was unchanged in 3 (median change +0.025). Under that rule 1 of the 222 comparisons was above BirdNET (Perch 2.0 on the rook) and 79 below, and Perch 2.0 (3.7) and BirdNET (4.0) had the best average ranks (Table S14). Reading each network at its own final output, an exploratory rule specified after review, raised re-ID accuracy for the ten animal networks with several layers in 90 of 130 network-dataset pairs (median change +0.069), and lowered it for the speech networks with several layers in 114 of 169 (median −0.041; Table S22). AVEX sl-BEATs, AvesEcho and NatureBEATs rose from average ranks of 22 to 25 to ranks of 9 to 10, but none was above BirdNET on any dataset. Under this rule 5 of 481 comparisons were above BirdNET, all Perch 2.0 and Perch, and 200 below; Perch 2.0 (3.4), BirdNET (5.6) and ConvNeXt (5.8) ranked highest, and no speech network exceeded BirdNET on the little penguin (Table S14). Within BirdNET, the internal layer chosen by the Fisher rule gave lower re-ID accuracy than the final embedding on 8 of 9 datasets (Table S16).

## 4 | DISCUSSION

Acoustic re-identification (re-ID) with machine learning (ML) is often reported to reach high accuracy. Applying four diagnostics to 13 datasets from nine species and seven studies, with 38 embeddings and controls, we found that recording each animal under its own conditions and randomly splitting individual clips can both inflate that accuracy substantially, confirming across species and studies effects previously reported in single species (Stowell et al. 2019; Arnaud et al. 2023). Within one year, background sound alone carried substantial information about the labelled chiffchaff identities: BirdNET balanced accuracy was 0.714 for backgrounds and 0.791 for songs. Background identification exceeded chance for 35 of the 38 embeddings and controls, so the effect was not confined to one neural network.

Count-matched random clip splits raised mean accuracy by up to 0.51. With unfamiliar animals present, the highest median accepted-and-correctly-named rate on wild session-disjoint endpoints was 0.374 with mean profiles and 0.462 with nearest-clip scoring. Both used independently calibrated thresholds targeting 10% stranger acceptance; that target was not guaranteed on new strangers. Only one dataset, a captive group of four zebra finches sharing one microphone, gave clear evidence that re-ID rested on the animal rather than the recording.

### 4.1 | What the diagnostics revealed

Diagnostic 1 showed that mixing clips across the recordists’ session boundary raises measured accuracy even when each bird contributes the same number of enrolment and test clips. For little owl with BirdNET and kernel ridge, the count-matched mean was 0.963 rather than 0.477. This does not show that randomisation improves a model’s ability to recognise birds in new sessions. Training on more varied recordings may genuinely help, but that improvement must be evaluated on further, untouched sessions. The present comparison changes session coverage and the clips tested; it does not isolate the effects of shared background, vocal drift, recording conditions or near-duplicate calls. The rook result likewise includes any change between the two years it mixes. A nearest-clip classifier need not show the largest gain, because its starting accuracy, ceiling and sensitivity to these factors differ from those of other classifiers.

Diagnostic 2 gave a different answer for each dataset, and within a dataset the answer was much the same for every neural network. For the within-year chiffchaff, the tests agreed that the classifier did not distinguish call from place, whichever embedding it used, so its re-ID accuracy cannot be read as a measure of the animal. For the little owl and both across-year datasets from the same study, backgrounds identified birds for at most one embedding after correction, while the best neural networks identified little owls from their calls well above BirdNET. On these datasets the backgrounds of the test year did not point to each bird, consistent with a change between years in each territory’s sound or in how it was recorded. Adding a bird’s own background sometimes lowered re-ID accuracy more than adding another bird’s, which we cannot yet explain. The background tests set only a lower bound, because a call also carries a site’s echoes and distance filtering, which an ambient clip does not capture; conversely, a test with ten birds could not detect planted calls 10 dB below their background, although those calls also carried their original sites, so the failed across-year chiffchaff test does not show that no individual signal exists. The great tit illustrates the first limit: although the split moved every bird to a new nest box, removing the song left most of BirdNET’s naming intact, and nest-box position alone rejected strangers better than the audio. In the 50-bird great tit dataset, 21 birds were enrolled and tested at the same nest box, so the larger sample does not carry the place control of the smaller one.

Diagnostic 3 was the most discriminating where it could be applied. A shared microphone removes site background and equipment but not each animal’s habitual position relative to the microphone; the loudness control, a crude indicator of distance, did not exceed the majority-class rate in the two zebra finch groups (Table S13). Passing the within-recording permutation test was not sufficient: the group of eight and the rook passed it, yet their paired-clip tests showed little or no difference between animal and recording pairs. All eligible same-bird, different-recording pairs in both zebra finch groups were more than seven days apart; the weak paired-clip result in the group of eight may therefore partly reflect changes over time in calls or recording conditions, which these data cannot separate from recording effects. In the fruit bat, recordings almost never held two bats, so neither test could be applied; there, identity was also tied to the experimental treatment (NMI 0.440) and the microphone channel (0.398), recording variables that a classifier can use. Where recordings contain several animals, passing an ordinary permutation test is not evidence that a classifier re-identifies animals, and where they do not, Diagnostic 3 cannot separate the two.

### 4.2 | Choosing a neural network

No neural network was a better choice than BirdNET on every dataset. Perch 2.0, which offers one embedding, was not detectably below BirdNET on any dataset and was above it on three; with 4 to 16 individuals per dataset, small differences would not be detected. On the rook, where every bird shares the microphone, Perch 2.0 passed the within-recording test (Table S21), but on this dataset passing that test did not show that re-ID rested on the animal, so its gain there may still come from the recording. On the tree pipit within a year, its backgrounds also identified birds after correction, so part of its gain there may come from the site; across years they did not.

The ranking of other networks depended on how their layer was chosen. The Fisher rule, which chooses the layer that best separates individuals in the enrolment clips, chose worse layers than each animal network’s own output in most comparisons, and chose the input layer of most speech networks, which scored better there than at their output. Networks with a single output, trained on bird or other animal sound, filled the first six places under every rule, but at their own outputs several animal networks with many layers ranked just behind them. Nine animal networks and every speech network analyse sound only up to 8 kHz at the recording’s own speed, and only the speech networks were also given slowed playback; the speech networks could not use absolute sound level, which BirdNET and the animal networks received. These differences favour some groups over others on some datasets, so the comparison supports networks trained on bird sound as a starting point, but not a ranking of network designs.

A higher re-ID accuracy is not, however, a better measure of the animal. On the within-year chiffchaff, where call and site are confounded, the four networks with the highest call-based accuracy were also the four with the highest background-only accuracy. Choosing a layer on enrolment sessions held out from the classifier changed the chosen layer in 130 of 222 comparisons and raised re-ID accuracy in 82 of them. That rule needs at least two recording sessions per animal in the enrolment clips, which seven of the 13 datasets lack, and studies planning enrolment recordings can provide them.

### 4.3 | Implications for population monitoring

Monitoring depends on open-set performance, because most animals heard at a site will not have been enrolled. Across the 36 neural networks on the wild session-disjoint endpoints, the highest median share of known birds’ calls accepted and correctly named rose from 37% with mean-profile scoring to 46% with nearest-clip scoring, while on the great tit and little owl BirdNET and Perch 2.0 named 60% to 75% of known birds’ calls correctly among four known birds (chance 25%), before any threshold was applied. These are outcomes of specified scoring and calibration procedures, not limits on what future models or better training data can achieve. Thresholds set on one group of strangers accepted up to 70% of another in the original mean-profile comparison, and results depended strongly on which birds were enrolled. Capture-recapture estimators that ignore misidentification can seriously overestimate population size (Yoshizaki et al. 2009; Link et al. 2010). Increasing the number of calibration strangers from 4 to 8 to 12 did not consistently improve the controlled great-tit results or ensure that the stranger budget transferred (Appendix S2). Because 6.7% of studies test the open-set case (Knight et al. 2024), this quantity is unknown for most published values.

### 4.4 | Recommendations

The diagnostics in Box 1 need no new data or models. We recommend that studies of acoustic re-ID report the split unit, its relation to the intended use, and count-matched comparisons with multiple seeds where random-clip effects are investigated; apply background tests where ambient recordings exist and record them where they do not; apply within-recording and paired-clip tests where animals share recordings; and report open-set performance with thresholds calibrated on other animals, together with the proportion of strangers those thresholds accept. Where the recording design can be chosen, recording several animals through one microphone, or one animal through several microphones, removes site background and equipment as explanations, although not each animal’s position. Removing the site after recording did not work: background augmentation raised call-based re-ID on three datasets but left backgrounds identifying birds on the within-year chiffchaff, spectral subtraction needs each test clip’s identity and so cannot be applied to unknown calls, score normalisation on the within-year chiffchaff lowered call-based re-ID for most embeddings while raising background-only re-ID, and session compensation did not remove recording day without also removing identity. When comparing neural networks, report the layer rule and each network’s frequency range. None replaces good design.

### 4.5 | Limitations and future directions

The datasets contain 4 to 16 individuals in the comparisons across datasets, typical of the field (Knight et al. 2024) but too few for precise estimates, so our conclusions rest on the direction and consistency of effects across many comparisons. The shared-microphone datasets are captive, background recordings exist in only one study, and no dataset lets a wild animal be heard by several recorders at once, which would separate animal from place in the field. The fruit bat analysis was limited to sound below 24 kHz, and networks with a 16 kHz input to sound below 8 kHz. Neural networks were used without retraining and with one classifier; fine-tuned networks and models of call sequences may perform better, and the diagnostics here are the means to test whether they gain on the animal or on the recording. Stable individual vocal differences across years have been reported with other methods (Klenova et al. 2012; Takagi 2020; Calcari et al. 2021) and with automated re-ID in ovenbirds (Lapp et al. 2025); applying the four diagnostics to such long-term wild datasets is the most important next step.

## 5 | CONCLUSIONS

Individual animals can be told apart from their calls with pretrained neural networks and a simple classifier, but whether a reported re-ID accuracy reflects the animal depends on how the data were collected and split. The four diagnostics presented here separate the animal from the recording, quantify the effect of the split and measure performance under monitoring conditions, and should accompany any re-ID accuracy used to justify acoustic population monitoring.

## Supporting information

Graphical abstract

## ACKNOWLEDGEMENTS

We thank the authors of the public datasets used here, and in particular Stowell et al. for recording an ambient background beside every call.

## FUNDING AND COMPETING INTERESTS

This research was funded by Xeroth AI Limited. Graham Wallington is its Chief Executive Officer and Jackie Lighten is its Chief Operating Officer; both are shareholders. The funder’s CEO and COO are the authors of this study and participated in the work as described in Author contributions.

## AUTHOR CONTRIBUTIONS

Graham Wallington conceived the study, implemented the measurements and analysed the data. Both authors designed the analyses, wrote and approved the final manuscript.

## DATA AVAILABILITY STATEMENT

All recordings analysed are publicly available. The zebra finch recordings of Elie & Theunissen (2020), the fruit bat recordings of Prat et al. (2017c) and the great tit recordings of Merino Recalde (2023) are cited with their dataset identifiers; the BirdPark (Rüttimann et al. 2024), rook (Martin et al. 2022a), little penguin and cockatoo (Huang et al. 2025b) and chiffchaff, little owl and tree pipit (Stowell et al. 2018) releases are cited in the reference list. The right whale recordings of Tolkova et al. (2026) were taken from github.com/avokloti/narw-acoustic-identification at commit 3cad65d, which declares no licence; no clip from it is redistributed. Analysis software version 0.3.0 is archived at https://doi.org/10.5281/zenodo.23167850 and developed at https://github.com/OpenXeroth/acoustic-re-id-research. The v10 evidence bundle is archived at https://doi.org/10.5281/zenodo.23174588. It contains aggregate projections of all 224 required result files, original and public-file checksums, anonymous split membership, original package and weight inventories, and the verification audit, together with aggregate projections and provenance for the 194 follow-up result files described in Appendix S2. All 224 original result files and all 194 new follow-up outputs have checksum-verified independent content-addressed archive copies. The projections omit recordings, embeddings, real identity annotations and per-clip observations; their hashes therefore differ from the originals. Numerical aggregates can be inspected independently; a fresh inference run additionally requires the original datasets and model weights, locally reconstructed annotations and the recorded environments. These resources remain subject to their publishers’ terms and are not relicensed by this release. The analysis software is available under the Apache 2.0 open-source licence. Third-party restrictions, including BirdNET v2.4’s CC BY-NC-SA 4.0 terms, continue to apply where applicable.

## SUPPORTING INFORMATION

Additional supporting information can be found in the separate file: Appendix S1 (methods detail), Tables S1 to S22, and Appendix S2 (follow-up sensitivity analyses and original split comparisons).\

## Notes

https://doi.org/10.5281/zenodo.23174588

https://doi.org/10.5281/zenodo.23167850

