## Supporting information: Appendices S1-S2 and supplementary tables for "Recording context and data splitting can inflate acoustic re-identification accuracy across species and studies"

### SUPPORTING INFORMATION: AUTHOR MANUSCRIPT v10

**Recording context and data splitting can inflate acoustic re-identification accuracy across species and studies.** Graham Wallington and Jackie Lighten. Version 10; prepared for bioRxiv submission. Licensing and access are stated in Appendix S2.

#### Appendix S1. Methods detail

##### S1.1 Reading BirdNET

BirdNET v2.4 was read at the layer before its species predictions, which gives 1,024 numbers for each 3 s of sound at 48 kHz. Sound was read as whole numbers and scaled, mixed to one channel and cut into 3-s segments at the recording's own sample rate. Each segment was then resampled to 48 kHz by the Fourier method, and the last segment was padded with silence. The embedding of a clip is the mean of its segment embeddings. This order matters: the Fourier method gives a different result if the whole recording is resampled before it is cut. Our reader reproduced the embeddings of birdnet==1.1.0 to within $3 \times 10^{-6}$ on eight great tit clips. BirdNET was given the sound at its recorded level, except that BirdPark clips carry a common endpoint gain (0.95 divided by the endpoint peak), preserving relative but not absolute levels.

Seven layers inside BirdNET can be read: the outputs of four processing stages of falling time resolution, two layers after the last stage and the final embedding. Each was averaged over time, and each was also summarised by its mean and spread over time, giving 14 candidate embeddings. These were used only in the analysis of BirdNET's internal layers (S1.13).

##### S1.2 Neural networks trained on animal or general sound

The 19 networks (Table S12) were loaded through the Bacpipe 1.3.5 library (Kather et al. 2026), with the weights it supplies, except the two AVEX networks, which were loaded through the Earth Species Project's avex library. A network entered the comparison only after it had loaded on our computer and produced an embedding for a real clip. The retained BirdNET v3 preview ONNX file has SHA-256 6f58d7ffa4c33bf49c8c67ac27bc5265a940a139cb67254477997bad41efc16d and differs from the preview3 file in Zenodo record 18247420; that record is not the exact checkpoint used here. Original package inventories and weight checksums are in the v8 evidence deposit. Each clip was resampled to the network's own sample rate with an anti-aliasing filter and cut into windows of the network's own input length (for AVEX, its 5-s training window); the last window, and any clip shorter than one window, was padded with silence. Each window passed through the network's published preprocessing, and the clip's embedding was the mean over its windows. The sound was given to the network at its recorded level, apart from the common BirdPark endpoint gain described in S1.1 and the network's own preprocessing.

For networks built from a stack of repeated transformer blocks (AvesEcho, Bird-MAE, ProtoCLR, BirdAVES, AVES, NatureBEATs, BioLingual, BEATs, AudioMAE and AVEX sl-BEATs-all), the output of every block, averaged over its time steps and then over windows, was a candidate embedding alongside the network's own embedding, which is the output read under the own-final-output rule (S1.5). The convolutional networks (Perch 2.0, Perch, BirdNET v3, SurfPerch, AVEX EfficientNet-B0, AudioProtoPNet, ConvNeXt, RCL_FS_BSED and VGGish) offer their own embedding only; Perch 2.0 and BirdNET v3 are distributed in a format whose internal layers cannot be read. For each network, the sample rate, window length and the named blocks that were read are listed in Table S12, from a record made by reloading each network and checking that it produced the same candidate names as the measurements. Every weight file was checked against its recorded checksum before that reload.

##### S1.3 Speech neural networks

Seventeen speech network checkpoints were measured: sixteen in an earlier run and again for this study, and XEUS. One of the sixteen duplicates another (below), so 16 speech networks are compared. The input to the first transformer block and the output of every block were candidates, at three playback speeds: the recording's own speed, one half and one third. Slowing was applied by dividing the recording's sample rate, so a 250 kHz bat clip was treated as 250, 125 or 83.3 kHz, and the highest original frequency reaching the 8 kHz range of these networks was 24 kHz. Sound was brought to 16 kHz by polyphase resampling with an anti-aliasing filter. Unlike BirdNET and the networks in S1.2, the speech networks were given each clip with its mean removed and scaled to a peak of one, so they could not use a clip's absolute level; the loudness control could. A clip shorter than a network accepts was padded with silence to the shortest length it accepts. On the chiffchaff and tree pipit datasets, the WavLM combinations that exceeded the 16 GB graphics card were read in 60-s windows, each layer averaged over every frame of every window.

The speaker-verification version of WavLM Base+ is loaded through the same interface as the other networks, which reads the transformer and not the speaker-recognition layers; its weights are identical to those of WavLM Base+, so it gives the same values at every layer and speed (it was run at the card's reduced precision on the across-year tree pipit, which changed Fisher ratios in the sixth significant figure and no choice or accuracy). It was measured because it was specified in advance, but it is left out of every count, rank and correction.

The 16 networks of the earlier run were run again on every dataset, keeping every candidate embedding, so that both layer rules could be applied to the same embeddings. Each candidate's re-ID accuracy and its right or wrong answer on every test clip were compared with the earlier run. Of 8,297 candidates on the 12 datasets of the earlier run, 8,295 were identical. The other two were layers 16 and 20 of the wav2vec 2.0 conformer at one-third speed on the across-year chiffchaff (0.120 against 0.125, and 0.150 against 0.155), where the graphics card's matrix arithmetic differed between the runs. Neither was chosen by either rule, and every value reported from that run was identical on every dataset (Table S19). The graphics card's matrix arithmetic had not been recorded in the earlier run. For 28 combinations of network and dataset (24 at full 32-bit precision and 4 at the card's reduced TF32 precision, on the great tit, the within-year and across-year chiffchaff and the across-year tree pipit), the rerun reproduced the earlier run only with the arithmetic that run had used; each of these networks was rerun whole on that dataset with that setting, which is recorded in its result file.

**S1.4 Classifier**

Embeddings were scaled to unit length and centred on the mean of the enrolment embeddings. Kernel ridge regression used a linear kernel and a fixed penalty of 1.0, and predicted a one-hot code for each individual; a test clip was assigned to the individual with the highest predicted value. Embeddings of three or fewer numbers (the controls) were standardised on the enrolment clips instead of scaled to unit length.

**S1.5 The layer rules**

*Fisher rule.* For each network, the candidate (layer and, for speech networks, playback speed) with the highest Fisher ratio on the enrolment clips was chosen: the ratio of variation between individuals to variation within individuals, averaged over the embedding's dimensions. The Fisher ratio was computed on raw candidate embeddings, using unweighted means over individuals; the classifier instead used the feature preparation in S1.4. The test clips were not used.

*Held-out-session rule.* Each individual's enrolment sessions were dealt into five groups by a fixed hash. For each group in turn, the classifier was fitted on the enrolment clips of the other four and scored on that group, and the candidate with the highest accuracy on the held-out sessions was chosen; ties went to the Fisher ratio. Individuals with one enrolment session, or whose sessions are not recorded, were used to fit the classifier but were never scored. The rule could be applied to seven datasets, with sessions as defined in Table 1: the fruit bat, the zebra finch recorded singly, both BirdPark groups, both great tit datasets and the rook. In the Stowell et al. datasets the enrolment clips carry no session within a bird, all little penguin enrolment clips come from one night, and the cockatoo release records no session; on these datasets the rule can only choose what the Fisher rule chose, so it is summarised on the six primary datasets where it applies.

*Own final output.* Specified after the fourth round of review and exploratory: each network was read at the output its authors provide, the network's own embedding for networks trained on animal or general sound, the last transformer layer at the recording's own speed for speech networks (layer 12 of base and layer 24 of large networks, layer 18 of XEUS), and the output at the recording's own speed for the speaker networks. Networks with one embedding, BirdNET and the controls are unchanged. The accuracy and right or wrong answer of every candidate were already stored, so no network was run again. Initial candidate-sweep diagnostics used 999 label permutations and 2,000 individual-bootstrap draws, confirmed from the byte-matched source for all 108 retained sweep, selection-refresh and replay records. Final paired network comparisons instead used 10,000 shared draws; headline and background nulls used 9,999 permutations, as stated in their tables.

**S1.6 Datasets**

*BirdPark.* From the BirdPark release (Rüttimann et al. 2024) we kept vocal segments that had been attributed to one bird by its transmitter, did not overlap another bird's syllable, and were recorded on the shared wall microphone. Each group was divided by recording file, with the three files of lowest hash as the test side.

*Rook.* Each rook recording stores several microphones as channels, and the channel index is not the same physical microphone on every day. We used channel 0 throughout, so that each call was counted once. Birds with at least 20 calls in each year were kept (11 of 15), recordings from 2020 were used for enrolment and from 2021 for testing, and enrolment was capped at 90 clips per bird, drawn with a fixed seed. For Diagnostic 3, the 2020 recordings were also divided by day, and the same calls were enrolled on channel 0 and tested on channel 1 (Table S4).

*Great tit with 50 birds.* From the release of Merino Recalde et al. (2024), a bird was included when its ring was recorded as the father of a nest attempt and also as a male great tit in the morphometric records, and when songs from his nest attempts existed in two or more years. Ten songs per bird and year were drawn by a hash of the file name; songs from the first year were used for enrolment and from the next for testing. As in the 16-bird dataset, a song is attributed to a bird through the nest attempt, not through an observation of the singer.

##### S1.7 Beecher's information statistic

On every embedding wider than three numbers, embeddings were scaled to unit length and reduced to principal components fitted on all call clips of the dataset, with at most one fewer component than individuals. For each component, a one-way analysis of variance gave F, and the component contributed $\log_2\sqrt{(F + n_0 - 1)/n_0}$ bits, where $n_0$ is the effective group size of the unbalanced design; components with F below one contributed nothing (Beecher 1989).

##### S1.8 Paired-clip test

All pairs of test clips were formed, or 800,000 drawn at random with a fixed seed where more exist, and scored by the cosine similarity of their unit-length embeddings. Each pair was classed by whether the two clips came from the same individual and from the same recording. The reported difference is the mean similarity of animal pairs (same individual, different recordings) minus that of recording pairs (different individuals, same recording). The standardised difference divides it by the square root of the mean of the two population variances. The equal error rate approximates the crossing of false-accept and false-reject rates. A preprint audit found that the original implementation could split tied scores according to their labels. The corrected implementation groups equal scores and includes rejection of all scores, then reports the mean of the two error rates at the attainable threshold closest to equality. The pre-submission audit recomputed these statistics from the original cached embeddings, matching manifest hashes, query order and pair summaries. All paired-clip equal error rates in Tables S3 and S4 were unchanged at their reported precision. Nineteen headline and rook runs also reproduced per-query classification and classifier AUC exactly; classifier EER was unchanged, with only floating-point rounding below $10^{-15}$ in some true-accept rates. The original and corrected values, source and cache digests, and audit code are in the v8 evidence deposit (https://doi.org/10.5281/zenodo.23159989).

**S1.9 Within-recording permutation test**

Enrolment labels were shuffled 9,999 times, each time only among clips from the same recording, and the classifier was refitted. A recording holding one animal keeps its label, so the test can only shuffle clips from recordings with two or more animals. In the fruit bat, the unit is the recording file. Only 8 of 6,188 bat enrolment recordings held two or more bats, so 16 of 6,252 enrolment clips could be shuffled. In the zebra finch recorded singly, the unit is the day, and 6 of the 15 enrolment days held more than one bird.

**S1.10 Background tests**

*Four pairings.* The classifier was fitted on calls or on background recordings and tested on calls or on backgrounds, with the same individuals, splits and seed as the call-only test. Each pairing had its own label-permutation test (9,999 shuffles) and interval over individuals (10,000 draws).

*Added background.* Each test clip was mixed with one background from its own individual and one from another individual (the donor), at 10 dB above, equal to and 10 dB below the clip's own level. Each individual's donor was the next individual in sorted order, and a hash of the test clip's name chose which of the own and donor backgrounds was used, from the test side of the split, and the classifier fitted on the original enrolment clips was applied. We report the difference in balanced accuracy between the two mixtures, with a paired interval over individuals, and the proportion of answers that the donor's background changed. Each mixture was peak-scaled to 0.95, whereas the original arm was unscaled. Own-versus-other mixtures share that scaling rule; comparisons with the original arm, including the donor-naming baseline, also differ in level preprocessing. The mixed clips were written once per dataset and read by every network.

*Donor naming.* When BirdNET named the wrong individual for a clip mixed with a donor's background, we counted how often the wrong answer was the donor, and compared that share with the same share among BirdNET's errors on the unmixed test clips, which measures how often it already confused each individual with its fixed donor, and with 9,999 random reassignments of donors that kept each clip’s own individual excluded. This analysis was specified after results were seen and is exploratory.

*Planted-signal test.* Calls from the ten within-year chiffchaffs with the most enrolment calls were mixed into the across-year chiffchaff background recordings, each bird into the territory of one across-year bird, enrolment calls into enrolment-year backgrounds and test calls into test-year backgrounds, at 10 dB above, equal to and 10 dB below the background. Each planted bird therefore had its own site, as in the real data, while the call it carried was known to differ between birds. The three datasets were scored with BirdNET as in Table 2.

*Right whale.* The four pairings were also run with BirdNET on 234 upcalls and 234 paired recordings of tag noise from 11 North Atlantic right whales carrying suction-cup tags, released with Tolkova et al. (2026); these data are not among the 13 datasets. Each whale carried one tag deployment, so no division by session exists. Within each whale, a call and its paired noise went to the test side when the first byte of a SHA-256 hash of a fixed salt followed by the clip stem was below 77 and to enrolment otherwise, putting about three in ten clips on the test side and keeping each call with its own noise. Individuals with no test clip after this split were omitted. The original manifests confirm 11 retained whales at every shift, with 162 enrolment and 72 test calls and the same numbers of paired noise clips. Before BirdNET read each clip, the 50 to 500 Hz band was shifted upward by 0 to 10,000 Hz in steps of 1,000 Hz, as in Tolkova et al. (2026), identically for calls and noise. The shift for the result was specified in advance as the one with the highest Fisher ratio on the enrolment calls, which was 10,000 Hz.

*Equivalence test on the little owl.* We asked whether background-only accuracy lay below chance plus one tenth of the call’s excess over chance (0.044 above chance), passing if the 95th percentile of the interval over individuals fell below that bound. The margin was set after the background estimate was seen but derived from the call result alone.

**S1.11 Remedies after recording**

*Background augmentation* (Stowell et al. 2019) mixed every enrolment call with a background from each other individual, at the clips' own levels, and added the mixtures to the enrolment clips; test clips were not changed. It was applied with BirdNET to the within-year chiffchaff, the little owl and the within-year tree pipit.

*Spectral subtraction.* For each individual within each side of the split, the mean short-time magnitude spectrum (Hann window of 1,024 samples, step 256) of its background recordings was computed. That spectrum, times one or times two, was subtracted from every call and background clip of the individual, keeping at least 5% of each clip's own magnitude in every frequency bin and keeping the phase. Because it uses each individual's own backgrounds, it asks whether the site can be removed when the site of every test clip is known.

*Adaptive score normalisation* (Matějka et al. 2017). Scores were the class-mean classifier's cosine similarities on unit-scaled, enrolment-centred embeddings. Duration and loudness controls are excluded because cosine unit normalisation removes their magnitude. The comparison set was every background recording on the enrolment side. For each enrolled individual and each test clip, the mean and spread of its 50 highest scores against that set were computed, and each score was replaced by the average of its two standardised values.

*Session compensation.* Four methods from human speaker recognition were applied to the fruit bat's BirdNET embeddings: subtracting each session's mean computed over all clips of that day on the same side of the split, every bat included, subtracting a running session mean that uses only the clips already heard (using the global enrolment mean until five clips have been heard), within-class covariance normalisation (Hatch et al. 2006) at eight levels of shrinkage, and removing 1 to 32 directions of variation between sessions. How well the recording day could still be predicted was measured by a classifier fitted on all but one bat and tested on the one left out, for each bat in turn. Because a day's mean includes every bat, subtracting it places each bat's clips opposite the other bats' clips of that day, so this classifier predicts that bat's days systematically wrongly after mean subtraction. Within-class covariance normalisation was also applied to the little owl.

##### S1.12 Open-set allocation

Within each of four groups of individuals, individuals were ordered by a hash of a fixed salt, the group and the individual, dealt round the four roles (known and strangers, each for calibration and for testing), and rotated so that every individual held every role once per group, giving 16 allocations. Each role held four individuals, two or three on the rook (11 birds) and 12 or 13 on the 50-bird great tit. The test gallery holds only the test-known individuals, so naming the correct individual is a choice among four (chance 0.25). The 36 neural-network embeddings were scaled to unit length, without per-dimension standardisation or enrolment centring, and scored by cosine similarity. Duration and loudness controls are excluded from this analysis because unit normalisation removes their magnitude. The separately reported acquisition-context control retains its own calibration-only standardisation. For each stranger budget, the calibrated threshold was the lowest score at which no more than that proportion of the calibration strangers' test clips was accepted; thresholds were set before the test individuals were scored. The test-derived threshold is the best of all thresholds that accept at most the budgeted proportion of the test strangers, found with the test strangers themselves. A calibrated threshold may accept more test strangers than the budget, and the proportion it accepted is reported. Balanced accuracy on known individuals is the mean over known individuals of the proportion of their clips accepted and correctly named; balanced accuracy on strangers is the mean over strangers of each stranger's proportion of clips rejected; their geometric mean is the AnimalCLEF 2025 score (Adam et al. 2025).

##### S1.13 BirdNET's internal layers, learned combination and sequence pooling

Of the 14 candidates in S1.1, the one with the highest enrolment Fisher ratio was chosen, and its permutation p was multiplied by the number of distinct candidates and capped at one. Two of the 14 repeat others (the post-convolution mean is the final embedding, and the final embedding's spread over time is zero), so the multiplier was 12. The learned combination took each of the seven layers averaged over time, scaled and centred it, projected it onto at most 16 directions that best separate individuals relative to variation within them, and joined the projections, all fitted on enrolment clips only; its permutation test refitted the projection for every shuffle. Learned sequence pooling weighted each time step of one of BirdNET's five time-resolved layers by a learned attention vector and mapped the weighted mean and spread to 64 numbers, trained on about two fifths of the individuals and scored on the others, which it never saw; with 3 to 7 individuals scored per dataset, its values are not comparable in scale with Table 2.

##### S1.14 Provenance and archive

Each result file records its code version, clip lists and their checksums, splits, random seeds and computing environment. Every run was made on one computer under a resource lease that records the job. The statistics added in this version (Tables S13, S14, S21 and S22 and the counts in the text) were computed from the same result files on a second computer, after every table and figure of the previous version had been reproduced there byte for byte. Every result file is archived in cloud storage under the SHA-256 checksum of its contents, and an archived copy is accepted only when its size and MD5 match the local file. Before the original tables were written, an evidence gate checked the presence, provenance and role-specific contracts of all 224 result files. It checked the stated resampling counts for most roles, but did not enforce them for corrected-sweep, speech, selection-refresh and speech-replay results. The separate pre-submission source audit recovered the initial-sweep settings for all 108 affected records: 999 permutations and 2,000 bootstrap draws, distinguished from the final analyses in S1.5. Every table and figure was then produced from those files by scripts that record the file and checksum behind every value.

#### Tables

**Table S1.** How closely identity is tied to each recording variable stored with the clips, over all clips of each dataset.

| Dataset | Recording variable | Levels | NMI | Cramér's V | Levels holding two or more individuals | Individuals on two or more levels |
|---|---|---|---|---|---|---|
| Zebra finch, group of four | microphone | 1 | 0.000 | 0.000 | 1 of 1 | 0 of 4 |
| Zebra finch, group of four | session | 7 | 0.473 | 0.644 | 7 of 7 | 4 of 4 |
| Zebra finch, group of eight | microphone | 1 | 0.000 | 0.000 | 1 of 1 | 0 of 8 |
| Zebra finch, group of eight | session | 7 | 0.407 | 0.494 | 6 of 7 | 8 of 8 |
| Rook, across year | microphone channel | 1 | 0.000 | 0.000 | 1 of 1 | 0 of 11 |
| Rook, across year | calling event | 327 | 0.515 | 0.629 | 50 of 327 | 11 of 11 |
| Rook, across year | recording time | 23 | 0.383 | 0.496 | 23 of 23 | 11 of 11 |
| Rook, across year | recording | 81 | 0.549 | 0.649 | 67 of 81 | 11 of 11 |
| Rook, across year | channels stored | 3 | 0.120 | 0.271 | 3 of 3 | 11 of 11 |
| Rook, across year | year | 2 | 0.185 | 0.476 | 2 of 2 | 11 of 11 |
| Zebra finch, one bird per recording | date | 24 | 0.827 | 0.863 | 7 of 24 | 9 of 9 |
| Zebra finch, one bird per recording | day | 24 | 0.827 | 0.863 | 7 of 24 | 9 of 9 |
| Chiffchaff, within year | session span | 1 | 0.000 | 0.000 | 1 of 1 | 0 of 13 |
| Chiffchaff, across year | session span | 1 | 0.000 | 0.000 | 1 of 1 | 0 of 10 |
| Little owl, across year | session span | 1 | 0.000 | 0.000 | 1 of 1 | 0 of 16 |
| Tree pipit, within year | session span | 1 | 0.000 | 0.000 | 1 of 1 | 0 of 10 |
| Tree pipit, across year | session span | 1 | 0.000 | 0.000 | 1 of 1 | 0 of 10 |
| Great tit, across year | cohort | 3 | 1.000 | 1.000 | 3 of 3 | 0 of 16 |
| Great tit, across year | nest box | 32 | 1.000 | 1.000 | 0 of 32 | 16 of 16 |
| Great tit, across year | recording | 215 | 1.000 | 1.000 | 0 of 215 | 16 of 16 |
| Great tit, across year | year | 3 | 0.358 | 0.500 | 3 of 3 | 16 of 16 |
| Little penguin, across night | call type | 1 | 0.000 | 0.000 | 1 of 1 | 0 of 15 |
| Little penguin, across night | night | 2 | 0.027 | 0.138 | 2 of 2 | 13 of 15 |
| Little penguin, across night | recording time | 3 | 0.493 | 0.576 | 2 of 3 | 5 of 13 |
| Little penguin, across night | day | 3 | 0.493 | 0.576 | 2 of 3 | 5 of 13 |
| Egyptian fruit bat, across year | microphone channel | 4 | 0.398 | 0.498 | 4 of 4 | 6 of 6 |
| Egyptian fruit bat, across year | folder | 6 | 0.057 | 0.186 | 6 of 6 | 6 of 6 |
| Egyptian fruit bat, across year | recording time | 7099 | 0.998 | 0.998 | 18 of 7099 | 6 of 6 |
| Egyptian fruit bat, across year | day | 76 | 0.136 | 0.285 | 75 of 76 | 6 of 6 |
| Egyptian fruit bat, across year | recording | 7109 | 0.998 | 0.998 | 12 of 7109 | 6 of 6 |
| Egyptian fruit bat, across year | treatment | 5 | 0.440 | 0.489 | 5 of 5 | 6 of 6 |

*Notes:* NMI, mutual information divided by the smaller of the two entropies: 0, the variable is unrelated to identity; 1, either determines the other. It is 1 both when every level holds one individual (each recording one animal) and when every individual sits on one level (each bird in one year cohort); the last two columns show which. A variable with one level carries no information. The Stowell et al. datasets store no recording variable within a split.

**Table S2.** Great tit place control: for each bird of the 16-bird dataset, whether any nest box or recording contributed to both its enrolment and test clips, whether the years differ, and the distance between its enrolment and test nest boxes.

| Bird | Same nest box on both sides | Same recording on both sides | Years differ | Distance between nest boxes (m) |
|---|---|---|---|---|
| GT-01 | no | no | yes | 59.4 |
| GT-02 | no | no | yes | 77.8 |
| GT-03 | no | no | yes | 65.0 |
| GT-04 | no | no | yes | 54.5 |
| GT-05 | no | no | yes | 79.8 |
| GT-06 | no | no | yes | 115.4 |
| GT-07 | no | no | yes | 57.1 |
| GT-08 | no | no | yes | 228.5 |
| GT-09 | no | no | yes | 194.3 |
| GT-10 | no | no | yes | 151.6 |
| GT-11 | no | no | yes | 145.6 |
| GT-12 | no | no | yes | 52.0 |
| GT-13 | no | no | yes | 174.5 |
| GT-14 | no | no | yes | 63.9 |
| GT-15 | no | no | yes | 113.3 |
| GT-16 | no | no | yes | 90.0 |

*Notes:* Bird codes are arbitrary; ring numbers are not published. Distances ranged from 52.0 to 228.5 m (median 84.9 m). In the 50-bird dataset, no recording contributed to both sides, but 21 of 50 birds were enrolled and tested at the same nest box; distances ranged from 0.0 to 733.4 m (median 29.2 m).

**Table S3.** Closed-set re-ID with BirdNET and the kernel ridge classifier on every dataset, all quantities.

| Dataset | n | Enrolment / test clips | Re-ID accuracy (95% interval) | Three-stage interval | Uniform chance | Majority class | Permutation 95th percentile | p | Within-recording 95th percentile (p) | Paired-clip difference | Standardised difference | Equal error rate |
|---|---|---|---|---|---|---|---|---|---|---|---|---|
| Zebra finch, group of four | 4 | 199 / 218 | 0.807 (0.669 to 0.963) | 0.613 to 0.963 | 0.250 | 0.445 | 0.491 | 0.0001 | 0.587 (0.0001) | 0.161 | 1.675 | 0.173 |
| Zebra finch, group of eight | 8 | 260 / 229 | 0.498 (0.176 to 0.725) | 0.124 to 0.754 | 0.125 | 0.358 | 0.293 | 0.0002 | 0.362 (0.0002) | −0.003 | −0.033 | 0.524 |
| Rook, across year | 11 | 990 / 4,417 | 0.375 (0.260 to 0.510) | 0.250 to 0.529 | 0.091 | 0.241 | 0.127 | 0.0001 | 0.224 (0.0001) | 0.030 | 0.260 | 0.449 |
| Zebra finch, one bird per recording | 9 | 90 / 90 | 0.967 (0.933 to 1.000) | 0.906 to 1.000 | 0.111 | 0.111 | 0.244 | 0.0001 | 0.856 (0.0002) | n/a | n/a | n/a |
| Chiffchaff, within year | 13 | 5,107 / 1,131 | 0.765 (0.681 to 0.846) | n/a | 0.077 | 0.393 | 0.154 | 0.0001 | n/a | n/a | n/a | n/a |
| Chiffchaff, across year | 10 | 324 / 200 | 0.160 (0.035 to 0.333) | n/a | 0.100 | 0.160 | 0.190 | 0.1327 | n/a | n/a | n/a | n/a |
| Little owl, across year | 16 | 545 / 407 | 0.501 (0.371 to 0.634) | n/a | 0.062 | 0.084 | 0.123 | 0.0001 | n/a | n/a | n/a | n/a |
| Tree pipit, within year | 10 | 409 / 303 | 0.376 (0.193 to 0.566) | n/a | 0.100 | 0.125 | 0.162 | 0.0001 | n/a | n/a | n/a | n/a |
| Tree pipit, across year | 10 | 409 / 313 | 0.214 (0.116 to 0.334) | n/a | 0.100 | 0.160 | 0.157 | 0.0022 | n/a | n/a | n/a | n/a |
| Great tit, across year | 16 | 160 / 160 | 0.444 (0.319 to 0.575) | 0.297 to 0.594 | 0.062 | 0.062 | 0.106 | 0.0001 | n/a | n/a | n/a | n/a |
| Little penguin, across night | 15 | 2,429 / 792 | 0.693 (0.604 to 0.772) | 0.594 to 0.782 | 0.067 | 0.136 | 0.115 | 0.0001 | n/a | n/a | n/a | n/a |
| Red-tailed black cockatoo, random split | 16 | 1,428 / 357 | 0.961 (0.929 to 0.980) | n/a | 0.062 | 0.216 | 0.182 | 0.0001 | n/a | n/a | n/a | n/a |
| Egyptian fruit bat, across year | 6 | 6,252 / 928 | 0.381 (0.189 to 0.569) | 0.188 to 0.565 | 0.167 | 0.297 | 0.239 | 0.0001 | n/a | n/a | n/a | n/a |
| Great tit, 50 birds, across year | 50 | 500 / 500 | 0.270 (0.202 to 0.342) | 0.188 to 0.357 | 0.020 | 0.020 | 0.036 | 0.0001 | n/a | n/a | n/a | n/a |

*Notes: As Table 2. Paired-clip test: mean cosine similarity of animal pairs (one individual, different recordings) minus that of recording pairs (different individuals, one recording), its standardised value and the equal error rate for telling the two kinds of pair apart from a single pair. n/a: the dataset has no recording holding two individuals, or does not identify recordings. The fruit bat has only five pairs of test clips from two bats in one recording and 16 enrolment clips that the within-recording test can shuffle, so neither test is reported. The little penguin's test clips record only a calendar date, across nests, so a pair from one date is not a pair from one recording. The equal error rates were verified from the original cached embeddings with corrected tie handling; see S1.8.*

**Table S4.** The rook under five divisions, with BirdNET.

| Division | n | Enrolment / test clips | Re-ID accuracy (95% interval) | Three-stage interval | Permutation 95th percentile | p | Within-recording p | Standardised paired-clip difference | Equal error rate |
|---|---|---|---|---|---|---|---|---|---|
| Enrolled in 2020, tested in 2021 | 11 | 8,226 / 4,417 | 0.422 (0.251 to 0.561) | 0.247 to 0.578 | 0.137 | 0.0001 | 0.0001 | 0.260 | 0.449 |
| The same, enrolment capped at 90 clips per bird | 11 | 990 / 4,417 | 0.375 (0.260 to 0.510) | 0.250 to 0.529 | 0.127 | 0.0001 | 0.0001 | 0.260 | 0.449 |
| The same clips, divided at random within each bird | 11 | 8,226 / 4,417 | 0.547 (0.351 to 0.683) | 0.343 to 0.694 | 0.127 | 0.0001 | 0.0001 | 0.102 | 0.482 |
| Enrolled and tested on different 2020 days | 14 | 5,656 / 4,143 | 0.395 (0.248 to 0.512) | 0.233 to 0.599 | 0.148 | 0.0001 | 0.0001 | 0.036 | 0.496 |
| The same calls, enrolled on channel 0, tested on channel 1 | 14 | 5,656 / 4,143 | 0.405 (0.259 to 0.515) | 0.238 to 0.607 | 0.156 | 0.0001 | 0.0001 | −0.045 | 0.510 |

*Notes: Channel 0 throughout unless stated. The channel index is not the same physical microphone on every recording day. The equal error rates were verified from the original cached embeddings with corrected tie handling; see S1.8.*

**Table S5.** Re-ID from calls and from background recordings for all 38 embeddings and controls, each at its layer chosen by the Fisher rule.

| Dataset | Embedding | C→C | B→B | C→B | B→C |
|---|---|---|---|---|---|
| Chiffchaff, within year | BirdNET v2.4 | 0.765 (0.681 to 0.846); 0.0001 | 0.737 (0.599 to 0.813); 0.0001 | 0.456 (0.344 to 0.625); 0.0001 | 0.508 (0.281 to 0.651); 0.0001 |
| Chiffchaff, within year | Perch 2.0 | 0.831 (0.756 to 0.885); 0.0001 | 0.787 (0.674 to 0.861); 0.0001 | 0.536 (0.457 to 0.649); 0.0001 | 0.599 (0.368 to 0.724); 0.0001 |
| Chiffchaff, within year | Perch | 0.778 (0.656 to 0.858); 0.0001 | 0.795 (0.663 to 0.865); 0.0001 | 0.553 (0.478 to 0.670); 0.0001 | 0.462 (0.334 to 0.583); 0.0001 |
| Chiffchaff, within year | SurfPerch | 0.729 (0.633 to 0.812); 0.0001 | 0.615 (0.501 to 0.695); 0.0001 | 0.239 (0.141 to 0.398); 0.0042 | 0.147 (0.086 to 0.226); 0.1438 |
| Chiffchaff, within year | BirdNET v3 (preview) | 0.624 (0.372 to 0.767); 0.0001 | 0.620 (0.416 to 0.731); 0.0001 | 0.545 (0.294 to 0.681); 0.0001 | 0.332 (0.233 to 0.504); 0.0001 |
| Chiffchaff, within year | AVEX sl-BEATs | 0.737 (0.588 to 0.818); 0.0001 | 0.685 (0.526 to 0.765); 0.0001 | 0.494 (0.341 to 0.609); 0.0001 | 0.378 (0.286 to 0.496); 0.0003 |
| Chiffchaff, within year | AVEX EfficientNet-B0 | 0.691 (0.576 to 0.761); 0.0001 | 0.608 (0.446 to 0.698); 0.0001 | 0.262 (0.136 to 0.382); 0.0012 | 0.207 (0.132 to 0.385); 0.0532 |
| Chiffchaff, within year | AvesEcho | 0.691 (0.522 to 0.774); 0.0001 | 0.655 (0.486 to 0.733); 0.0001 | 0.402 (0.247 to 0.506); 0.0001 | 0.589 (0.362 to 0.709); 0.0001 |
| Chiffchaff, within year | AudioProtoPNet | 0.806 (0.649 to 0.889); 0.0001 | 0.815 (0.687 to 0.880); 0.0001 | 0.675 (0.516 to 0.746); 0.0001 | 0.574 (0.377 to 0.686); 0.0001 |
| Chiffchaff, within year | ConvNeXt (BirdSet) | 0.805 (0.688 to 0.866); 0.0001 | 0.835 (0.693 to 0.902); 0.0001 | 0.671 (0.498 to 0.751); 0.0001 | 0.559 (0.379 to 0.665); 0.0001 |
| Chiffchaff, within year | Bird-MAE | 0.319 (0.110 to 0.446); 0.0001 | 0.270 (0.150 to 0.428); 0.0002 | 0.363 (0.095 to 0.519); 0.0001 | 0.126 (0.025 to 0.327); 0.2143 |
| Chiffchaff, within year | ProtoCLR | 0.325 (0.211 to 0.477); 0.0001 | 0.285 (0.190 to 0.461); 0.0009 | 0.185 (0.103 to 0.321); 0.0832 | 0.111 (0.023 to 0.293); 0.2838 |
| Chiffchaff, within year | RCL_FS_BSED | 0.256 (0.119 to 0.496); 0.0154 | 0.283 (0.171 to 0.485); 0.0060 | 0.234 (0.107 to 0.468); 0.0660 | 0.134 (0.018 to 0.382); 0.1790 |
| Chiffchaff, within year | BirdAVES | 0.420 (0.329 to 0.589); 0.0001 | 0.363 (0.249 to 0.553); 0.0001 | 0.311 (0.188 to 0.487); 0.0029 | 0.172 (0.057 to 0.359); 0.1106 |
| Chiffchaff, within year | AVES | 0.397 (0.310 to 0.555); 0.0001 | 0.345 (0.236 to 0.518); 0.0001 | 0.357 (0.173 to 0.497); 0.0005 | 0.159 (0.046 to 0.357); 0.1242 |
| Chiffchaff, within year | NatureBEATs | 0.741 (0.579 to 0.816); 0.0001 | 0.682 (0.530 to 0.756); 0.0001 | 0.511 (0.330 to 0.612); 0.0001 | 0.553 (0.399 to 0.629); 0.0001 |
| Chiffchaff, within year | BioLingual | 0.637 (0.451 to 0.747); 0.0001 | 0.583 (0.398 to 0.695); 0.0001 | 0.200 (0.078 to 0.453); 0.0766 | 0.254 (0.130 to 0.485); 0.0597 |
| Chiffchaff, within year | BEATs | 0.558 (0.343 to 0.672); 0.0001 | 0.617 (0.435 to 0.713); 0.0001 | 0.494 (0.243 to 0.635); 0.0001 | 0.263 (0.134 to 0.493); 0.0072 |
| Chiffchaff, within year | AudioMAE | 0.184 (0.124 to 0.280); 0.0882 | 0.185 (0.053 to 0.471); 0.0912 | 0.142 (0.047 to 0.267); 0.1817 | 0.077 (0.021 to 0.186); 0.5387 |
| Chiffchaff, within year | VGGish | 0.458 (0.256 to 0.560); 0.0001 | 0.417 (0.230 to 0.511); 0.0001 | 0.209 (0.117 to 0.374); 0.0498 | 0.399 (0.125 to 0.556); 0.0001 |
| Chiffchaff, within year | ECAPA-TDNN (speaker) | 0.416 (0.303 to 0.547); 0.0001 | 0.498 (0.347 to 0.587); 0.0001 | 0.198 (0.101 to 0.386); 0.0078 | 0.309 (0.189 to 0.383); 0.0002 |
| Chiffchaff, within year | ResNet (speaker) | 0.523 (0.326 to 0.633); 0.0001 | 0.455 (0.310 to 0.550); 0.0001 | 0.228 (0.140 to 0.416); 0.0009 | 0.414 (0.158 to 0.539); 0.0001 |
| Chiffchaff, within year | x-vector (speaker) | 0.517 (0.325 to 0.634); 0.0001 | 0.536 (0.333 to 0.639); 0.0001 | 0.177 (0.076 to 0.397); 0.0779 | 0.294 (0.148 to 0.366); 0.0079 |
| Chiffchaff, within year | wav2vec 2.0 Base | 0.331 (0.234 to 0.528); 0.0001 | 0.323 (0.217 to 0.516); 0.0001 | 0.271 (0.142 to 0.392); 0.0063 | 0.185 (0.074 to 0.381); 0.0831 |
| Chiffchaff, within year | wav2vec 2.0 Large (robust) | 0.305 (0.208 to 0.444); 0.0004 | 0.289 (0.187 to 0.445); 0.0008 | 0.285 (0.140 to 0.442); 0.0030 | 0.141 (0.032 to 0.340); 0.1609 |
| Chiffchaff, within year | XLS-R 300M | 0.321 (0.214 to 0.444); 0.0001 | 0.285 (0.189 to 0.425); 0.0012 | 0.301 (0.137 to 0.445); 0.0010 | 0.160 (0.060 to 0.332); 0.1286 |
| Chiffchaff, within year | wav2vec 2.0 Conformer Large | 0.414 (0.306 to 0.577); 0.0001 | 0.425 (0.299 to 0.561); 0.0001 | 0.268 (0.157 to 0.492); 0.0046 | 0.221 (0.137 to 0.362); 0.0619 |
| Chiffchaff, within year | MMS 300M | 0.324 (0.225 to 0.473); 0.0001 | 0.278 (0.184 to 0.428); 0.0005 | 0.300 (0.148 to 0.439); 0.0004 | 0.161 (0.055 to 0.344); 0.1289 |
| Chiffchaff, within year | HuBERT Base | 0.314 (0.215 to 0.504); 0.0002 | 0.322 (0.225 to 0.497); 0.0002 | 0.251 (0.122 to 0.364); 0.0141 | 0.155 (0.040 to 0.358); 0.1306 |
| Chiffchaff, within year | HuBERT Large | 0.316 (0.214 to 0.444); 0.0003 | 0.283 (0.188 to 0.435); 0.0032 | 0.311 (0.144 to 0.474); 0.0023 | 0.135 (0.031 to 0.330); 0.1674 |
| Chiffchaff, within year | data2vec Base (100 h) | 0.382 (0.278 to 0.505); 0.0001 | 0.375 (0.227 to 0.471); 0.0001 | 0.229 (0.117 to 0.443); 0.0250 | 0.123 (0.031 to 0.297); 0.2099 |
| Chiffchaff, within year | data2vec Base (960 h) | 0.378 (0.270 to 0.494); 0.0001 | 0.330 (0.208 to 0.440); 0.0001 | 0.270 (0.168 to 0.461); 0.0058 | 0.161 (0.060 to 0.333); 0.1227 |
| Chiffchaff, within year | WavLM Base+ | 0.327 (0.232 to 0.518); 0.0001 | 0.319 (0.226 to 0.499); 0.0002 | 0.284 (0.172 to 0.415); 0.0031 | 0.164 (0.063 to 0.340); 0.1141 |
| Chiffchaff, within year | WavLM Large | 0.347 (0.226 to 0.472); 0.0002 | 0.299 (0.198 to 0.434); 0.0014 | 0.309 (0.142 to 0.444); 0.0019 | 0.141 (0.040 to 0.330); 0.1615 |
| Chiffchaff, within year | UniSpeech-SAT Base+ | 0.317 (0.219 to 0.504); 0.0001 | 0.304 (0.210 to 0.477); 0.0004 | 0.265 (0.155 to 0.401); 0.0092 | 0.193 (0.081 to 0.394); 0.0869 |
| Chiffchaff, within year | XEUS | 0.341 (0.236 to 0.467); 0.0001 | 0.334 (0.216 to 0.473); 0.0002 | 0.313 (0.181 to 0.459); 0.0018 | 0.154 (0.056 to 0.327); 0.1395 |
| Chiffchaff, within year | Loudness control | 0.094 (0.012 to 0.269); 0.4053 | 0.118 (0.016 to 0.307); 0.2596 | 0.073 (0.002 to 0.255); 0.5224 | 0.116 (0.029 to 0.281); 0.2341 |
| Chiffchaff, within year | Duration control | 0.040 (0.000 to 0.146); 0.8848 | 0.088 (0.000 to 0.304); 0.4116 | 0.050 (0.000 to 0.191); 0.6576 | 0.047 (0.000 to 0.161); 0.8036 |
| Little owl, across year | BirdNET v2.4 | 0.501 (0.371 to 0.634); 0.0001 | 0.105 (0.027 to 0.203); 0.0860 | 0.083 (0.009 to 0.198); 0.2263 | 0.076 (0.004 to 0.205); 0.3198 |
| Little owl, across year | Perch 2.0 | 0.651 (0.489 to 0.785); 0.0001 | 0.029 (0.008 to 0.054); 0.9597 | 0.108 (0.017 to 0.245); 0.0474 | 0.052 (0.000 to 0.127); 0.6445 |
| Little owl, across year | Perch | 0.646 (0.499 to 0.776); 0.0001 | 0.064 (0.024 to 0.114); 0.4603 | 0.115 (0.037 to 0.215); 0.0127 | 0.052 (0.000 to 0.147); 0.6578 |
| Little owl, across year | SurfPerch | 0.442 (0.290 to 0.591); 0.0001 | 0.147 (0.068 to 0.238); 0.0005 | 0.120 (0.028 to 0.232); 0.0133 | 0.049 (0.003 to 0.126); 0.6800 |
| Little owl, across year | BirdNET v3 (preview) | 0.373 (0.233 to 0.530); 0.0001 | 0.086 (0.012 to 0.194); 0.1735 | 0.156 (0.018 to 0.332); 0.0004 | 0.005 (0.000 to 0.016); 0.9977 |
| Little owl, across year | AVEX sl-BEATs | 0.194 (0.081 to 0.329); 0.0005 | 0.149 (0.031 to 0.307); 0.0122 | 0.068 (0.000 to 0.210); 0.4289 | 0.054 (0.000 to 0.159); 0.6706 |

| Dataset | Embedding | C→C | B→B | C→B | B→C |
|---|---|---|---|---|---|
| Little owl, across year | AVEX EfficientNet-B0 | 0.373 (0.234 to 0.522); 0.0001 | 0.098 (0.025 to 0.191); 0.0626 | 0.098 (0.017 to 0.200); 0.0727 | 0.084 (0.012 to 0.189); 0.2328 |
| Little owl, across year | AvesEcho | 0.138 (0.028 to 0.264); 0.0439 | 0.086 (0.023 to 0.173); 0.2243 | 0.066 (0.000 to 0.202); 0.4412 | 0.091 (0.000 to 0.224); 0.1555 |
| Little owl, across year | AudioProtoPNet | 0.631 (0.469 to 0.781); 0.0001 | 0.068 (0.027 to 0.126); 0.3611 | 0.088 (0.011 to 0.188); 0.1376 | 0.047 (0.000 to 0.147); 0.7284 |
| Little owl, across year | ConvNeXt (BirdSet) | 0.624 (0.477 to 0.751); 0.0001 | 0.061 (0.023 to 0.106); 0.4979 | 0.068 (0.013 to 0.148); 0.4036 | 0.074 (0.005 to 0.174); 0.3434 |
| Little owl, across year | Bird-MAE | 0.251 (0.102 to 0.407); 0.0003 | 0.098 (0.010 to 0.221); 0.1489 | 0.066 (0.000 to 0.207); 0.4540 | 0.044 (0.000 to 0.145); 0.8725 |
| Little owl, across year | ProtoCLR | 0.172 (0.065 to 0.305); 0.0077 | 0.054 (0.000 to 0.147); 0.5654 | 0.110 (0.000 to 0.272); 0.0287 | 0.044 (0.000 to 0.145); 0.8910 |
| Little owl, across year | RCL_FS_BSED | 0.189 (0.068 to 0.328); 0.0015 | 0.064 (0.004 to 0.169); 0.4495 | 0.103 (0.000 to 0.254); 0.1423 | 0.059 (0.000 to 0.149); 0.5083 |
| Little owl, across year | BirdAVES | 0.285 (0.158 to 0.430); 0.0001 | 0.076 (0.012 to 0.184); 0.2973 | 0.095 (0.000 to 0.248); 0.0941 | 0.101 (0.000 to 0.244); 0.1078 |
| Little owl, across year | AVES | 0.285 (0.162 to 0.410); 0.0001 | 0.108 (0.021 to 0.214); 0.0989 | 0.076 (0.000 to 0.203); 0.3371 | 0.143 (0.021 to 0.299); 0.0110 |
| Little owl, across year | NatureBEATs | 0.241 (0.134 to 0.357); 0.0002 | 0.132 (0.025 to 0.272); 0.0295 | 0.066 (0.000 to 0.207); 0.4569 | 0.066 (0.000 to 0.151); 0.4489 |
| Little owl, across year | BioLingual | 0.133 (0.023 to 0.254); 0.0578 | 0.064 (0.000 to 0.186); 0.4593 | 0.064 (0.000 to 0.199); 0.5268 | 0.044 (0.000 to 0.145); 0.9168 |
| Little owl, across year | BEATs | 0.216 (0.088 to 0.369); 0.0001 | 0.149 (0.034 to 0.307); 0.0177 | 0.066 (0.000 to 0.207); 0.4734 | 0.054 (0.000 to 0.174); 0.7038 |
| Little owl, across year | AudioMAE | 0.170 (0.066 to 0.288); 0.0087 | 0.081 (0.000 to 0.214); 0.3146 | 0.066 (0.000 to 0.207); 0.4807 | 0.044 (0.000 to 0.145); 0.9251 |
| Little owl, across year | VGGish | 0.231 (0.098 to 0.379); 0.0001 | 0.037 (0.011 to 0.074); 0.8080 | 0.034 (0.000 to 0.089); 0.8451 | 0.091 (0.014 to 0.214); 0.1602 |
| Little owl, across year | ECAPA-TDNN (speaker) | 0.437 (0.278 to 0.598); 0.0001 | 0.071 (0.015 to 0.163); 0.3360 | 0.122 (0.000 to 0.261); 0.0103 | 0.076 (0.009 to 0.173); 0.3076 |
| Little owl, across year | ResNet (speaker) | 0.437 (0.280 to 0.591); 0.0001 | 0.098 (0.037 to 0.165); 0.0658 | 0.081 (0.000 to 0.216); 0.2028 | 0.071 (0.000 to 0.203); 0.3688 |
| Little owl, across year | x-vector (speaker) | 0.393 (0.252 to 0.537); 0.0001 | 0.105 (0.027 to 0.204); 0.0683 | 0.120 (0.000 to 0.299); 0.0062 | 0.167 (0.005 to 0.352); 0.0021 |
| Little owl, across year | wav2vec 2.0 Base | 0.263 (0.105 to 0.444); 0.0002 | 0.132 (0.035 to 0.246); 0.0265 | 0.103 (0.002 to 0.250); 0.1012 | 0.150 (0.027 to 0.302); 0.0135 |
| Little owl, across year | wav2vec 2.0 Large (robust) | 0.270 (0.132 to 0.419); 0.0002 | 0.090 (0.007 to 0.188); 0.2107 | 0.110 (0.004 to 0.250); 0.0883 | 0.098 (0.013 to 0.235); 0.1600 |
| Little owl, across year | XLS-R 300M | 0.292 (0.138 to 0.458); 0.0001 | 0.081 (0.004 to 0.176); 0.2789 | 0.115 (0.004 to 0.265); 0.0671 | 0.103 (0.010 to 0.233); 0.1391 |
| Little owl, across year | wav2vec 2.0 Conformer Large | 0.349 (0.198 to 0.507); 0.0001 | 0.117 (0.034 to 0.220); 0.0748 | 0.115 (0.005 to 0.264); 0.0699 | 0.081 (0.000 to 0.223); 0.2878 |
| Little owl, across year | MMS 300M | 0.290 (0.137 to 0.459); 0.0001 | 0.090 (0.014 to 0.188); 0.1986 | 0.120 (0.004 to 0.272); 0.0501 | 0.115 (0.012 to 0.262); 0.0833 |
| Little owl, across year | HuBERT Base | 0.290 (0.151 to 0.440); 0.0001 | 0.152 (0.041 to 0.287); 0.0133 | 0.086 (0.005 to 0.222); 0.2233 | 0.108 (0.010 to 0.253); 0.1171 |
| Little owl, across year | HuBERT Large | 0.187 (0.089 to 0.309); 0.0073 | 0.110 (0.021 to 0.223); 0.1099 | 0.088 (0.000 to 0.234); 0.1473 | 0.059 (0.000 to 0.167); 0.5474 |
| Little owl, across year | data2vec Base (100 h) | 0.361 (0.214 to 0.521); 0.0001 | 0.108 (0.031 to 0.201); 0.1049 | 0.117 (0.005 to 0.266); 0.0587 | 0.091 (0.000 to 0.231); 0.2006 |
| Little owl, across year | data2vec Base (960 h) | 0.378 (0.222 to 0.551); 0.0001 | 0.098 (0.025 to 0.189); 0.1535 | 0.117 (0.005 to 0.271); 0.0579 | 0.079 (0.000 to 0.216); 0.2981 |
| Little owl, across year | WavLM Base+ | 0.305 (0.162 to 0.455); 0.0001 | 0.149 (0.044 to 0.278); 0.0137 | 0.117 (0.000 to 0.280); 0.0558 | 0.064 (0.000 to 0.201); 0.4642 |
| Little owl, across year | WavLM Large | 0.268 (0.131 to 0.414); 0.0001 | 0.086 (0.013 to 0.176); 0.2450 | 0.110 (0.000 to 0.264); 0.0891 | 0.101 (0.005 to 0.247); 0.1644 |
| Little owl, across year | UniSpeech-SAT Base+ | 0.270 (0.130 to 0.427); 0.0001 | 0.161 (0.050 to 0.295); 0.0062 | 0.110 (0.005 to 0.267); 0.0801 | 0.098 (0.005 to 0.241); 0.1630 |
| Little owl, across year | XEUS | 0.297 (0.155 to 0.454); 0.0001 | 0.112 (0.024 to 0.234); 0.0891 | 0.117 (0.000 to 0.274); 0.0642 | 0.088 (0.003 to 0.230); 0.2368 |
| Little owl, across year | Loudness control | 0.069 (0.014 to 0.136); 0.4105 | 0.020 (0.000 to 0.048); 0.9465 | 0.020 (0.003 to 0.041); 0.9470 | 0.059 (0.000 to 0.151); 0.5832 |
| Little owl, across year | Duration control | 0.115 (0.000 to 0.294); 0.0945 | 0.054 (0.000 to 0.172); 0.5617 | 0.068 (0.000 to 0.214); 0.3975 | 0.044 (0.000 to 0.145); 0.9470 |
| Tree pipit, within year | BirdNET v2.4 | 0.376 (0.193 to 0.566); 0.0001 | 0.177 (0.106 to 0.246); 0.0408 | 0.147 (0.046 to 0.257); 0.1293 | 0.106 (0.023 to 0.219); 0.4515 |
| Tree pipit, within year | Perch 2.0 | 0.548 (0.357 to 0.736); 0.0001 | 0.256 (0.109 to 0.426); 0.0004 | 0.150 (0.016 to 0.379); 0.0928 | 0.083 (0.000 to 0.262); 0.7466 |
| Tree pipit, within year | Perch | 0.426 (0.225 to 0.628); 0.0001 | 0.242 (0.111 to 0.393); 0.0004 | 0.225 (0.053 to 0.436); 0.0008 | 0.129 (0.027 to 0.284); 0.1858 |
| Tree pipit, within year | SurfPerch | 0.455 (0.261 to 0.639); 0.0001 | 0.280 (0.140 to 0.436); 0.0001 | 0.188 (0.040 to 0.415); 0.0042 | 0.165 (0.050 to 0.295); 0.0263 |
| Tree pipit, within year | BirdNET v3 (preview) | 0.257 (0.127 to 0.414); 0.0001 | 0.259 (0.147 to 0.378); 0.0004 | 0.215 (0.076 to 0.399); 0.0024 | 0.086 (0.000 to 0.266); 0.7436 |
| Tree pipit, within year | AVEX sl-BEATs | 0.162 (0.054 to 0.285); 0.0515 | 0.119 (0.040 to 0.213); 0.3006 | 0.099 (0.004 to 0.275); 0.5300 | 0.145 (0.043 to 0.269); 0.0987 |
| Tree pipit, within year | AVEX EfficientNet-B0 | 0.281 (0.108 to 0.473); 0.0001 | 0.198 (0.077 to 0.329); 0.0023 | 0.154 (0.027 to 0.354); 0.0615 | 0.191 (0.061 to 0.341); 0.0047 |
| Tree pipit, within year | AvesEcho | 0.135 (0.015 to 0.327); 0.1992 | 0.089 (0.007 to 0.212); 0.5810 | 0.116 (0.000 to 0.334); 0.3580 | 0.102 (0.013 to 0.240); 0.4698 |
| Tree pipit, within year | AudioProtoPNet | 0.442 (0.234 to 0.655); 0.0001 | 0.253 (0.129 to 0.409); 0.0005 | 0.246 (0.089 to 0.441); 0.0004 | 0.089 (0.000 to 0.269); 0.6622 |
| Tree pipit, within year | ConvNeXt (BirdSet) | 0.502 (0.290 to 0.705); 0.0001 | 0.246 (0.122 to 0.404); 0.0004 | 0.225 (0.081 to 0.410); 0.0008 | 0.102 (0.016 to 0.245); 0.5015 |
| Tree pipit, within year | Bird-MAE | 0.069 (0.019 to 0.131); 0.8566 | 0.109 (0.023 to 0.233); 0.4521 | 0.085 (0.000 to 0.207); 0.6953 | 0.063 (0.007 to 0.130); 0.9126 |
| Tree pipit, within year | ProtoCLR | 0.152 (0.026 to 0.309); 0.0891 | 0.137 (0.033 to 0.271); 0.1454 | 0.143 (0.013 to 0.365); 0.0735 | 0.102 (0.000 to 0.272); 0.4982 |
| Tree pipit, within year | RCL_FS_BSED | 0.172 (0.053 to 0.329); 0.1120 | 0.092 (0.013 to 0.216); 0.5313 | 0.212 (0.019 to 0.420); 0.0282 | 0.096 (0.021 to 0.194); 0.5178 |
| Tree pipit, within year | BirdAVES | 0.300 (0.104 to 0.503); 0.0003 | 0.218 (0.076 to 0.409); 0.0173 | 0.218 (0.033 to 0.446); 0.0095 | 0.228 (0.048 to 0.445); 0.0043 |

| Dataset | Embedding | C→C | B→B | C→B | B→C |
|---|---|---|---|---|---|
| Tree pipit, within year | AVES | 0.228 (0.064 to 0.424); 0.0065 | 0.212 (0.083 to 0.385); 0.0210 | 0.188 (0.017 to 0.426); 0.0401 | 0.277 (0.069 to 0.503); 0.0006 |
| Tree pipit, within year | NatureBEATs | 0.182 (0.083 to 0.283); 0.0242 | 0.113 (0.039 to 0.198); 0.3616 | 0.109 (0.011 to 0.255); 0.4074 | 0.158 (0.050 to 0.280); 0.0571 |
| Tree pipit, within year | BioLingual | 0.112 (0.014 to 0.227); 0.3898 | 0.109 (0.037 to 0.199); 0.4210 | 0.099 (0.007 to 0.257); 0.5148 | 0.086 (0.010 to 0.210); 0.6694 |
| Tree pipit, within year | BEATs | 0.211 (0.095 to 0.329); 0.0052 | 0.140 (0.060 to 0.229); 0.1677 | 0.126 (0.048 to 0.224); 0.2568 | 0.129 (0.021 to 0.259); 0.2320 |
| Tree pipit, within year | AudioMAE | 0.125 (0.024 to 0.240); 0.2976 | 0.130 (0.000 to 0.292); 0.2772 | 0.167 (0.000 to 0.396); 0.0485 | 0.112 (0.000 to 0.288); 0.3141 |
| Tree pipit, within year | VGGish | 0.116 (0.028 to 0.218); 0.3378 | 0.164 (0.060 to 0.292); 0.0800 | 0.068 (0.007 to 0.152); 0.8217 | 0.033 (0.000 to 0.072); 0.9801 |
| Tree pipit, within year | ECAPA-TDNN (speaker) | 0.356 (0.166 to 0.565); 0.0001 | 0.222 (0.103 to 0.350); 0.0010 | 0.171 (0.018 to 0.400); 0.0285 | 0.195 (0.050 to 0.375); 0.0080 |
| Tree pipit, within year | ResNet (speaker) | 0.264 (0.127 to 0.426); 0.0001 | 0.208 (0.110 to 0.320); 0.0011 | 0.160 (0.014 to 0.368); 0.0310 | 0.162 (0.048 to 0.306); 0.0360 |
| Tree pipit, within year | x-vector (speaker) | 0.304 (0.117 to 0.512); 0.0001 | 0.184 (0.058 to 0.334); 0.0348 | 0.130 (0.007 to 0.338); 0.2151 | 0.155 (0.034 to 0.301); 0.0796 |
| Tree pipit, within year | wav2vec 2.0 Base | 0.231 (0.060 to 0.419); 0.0053 | 0.195 (0.062 to 0.380); 0.0290 | 0.082 (0.024 to 0.150); 0.6838 | 0.155 (0.022 to 0.343); 0.1023 |
| Tree pipit, within year | wav2vec 2.0 Large (robust) | 0.168 (0.004 to 0.388); 0.1049 | 0.137 (0.023 to 0.328); 0.2251 | 0.109 (0.015 to 0.222); 0.4228 | 0.205 (0.010 to 0.444); 0.0252 |
| Tree pipit, within year | XLS-R 300M | 0.135 (0.014 to 0.327); 0.2424 | 0.150 (0.022 to 0.343); 0.1641 | 0.167 (0.004 to 0.411); 0.0754 | 0.201 (0.025 to 0.424); 0.0307 |
| Tree pipit, within year | wav2vec 2.0 Conformer Large | 0.149 (0.020 to 0.342); 0.1782 | 0.174 (0.051 to 0.366); 0.0837 | 0.191 (0.010 to 0.418); 0.0270 | 0.231 (0.022 to 0.461); 0.0099 |
| Tree pipit, within year | MMS 300M | 0.125 (0.016 to 0.316); 0.2906 | 0.143 (0.020 to 0.336); 0.1920 | 0.150 (0.007 to 0.362); 0.1362 | 0.201 (0.029 to 0.414); 0.0292 |
| Tree pipit, within year | HuBERT Base | 0.191 (0.035 to 0.384); 0.0429 | 0.150 (0.040 to 0.321); 0.1522 | 0.143 (0.033 to 0.280); 0.1515 | 0.198 (0.017 to 0.413); 0.0304 |
| Tree pipit, within year | HuBERT Large | 0.188 (0.016 to 0.419); 0.0624 | 0.150 (0.032 to 0.342); 0.1625 | 0.078 (0.011 to 0.178); 0.7111 | 0.221 (0.019 to 0.480); 0.0113 |
| Tree pipit, within year | data2vec Base (100 h) | 0.182 (0.038 to 0.374); 0.0644 | 0.191 (0.059 to 0.386); 0.0463 | 0.116 (0.028 to 0.218); 0.3642 | 0.224 (0.039 to 0.440); 0.0102 |
| Tree pipit, within year | data2vec Base (960 h) | 0.172 (0.041 to 0.356); 0.0870 | 0.177 (0.049 to 0.372); 0.0690 | 0.102 (0.028 to 0.188); 0.4934 | 0.224 (0.047 to 0.439); 0.0094 |
| Tree pipit, within year | WavLM Base+ | 0.158 (0.025 to 0.350); 0.1245 | 0.184 (0.053 to 0.360); 0.0512 | 0.167 (0.031 to 0.351); 0.0637 | 0.198 (0.010 to 0.426); 0.0310 |
| Tree pipit, within year | WavLM Large | 0.165 (0.009 to 0.374); 0.1156 | 0.164 (0.044 to 0.353); 0.1106 | 0.085 (0.017 to 0.177); 0.6563 | 0.221 (0.010 to 0.486); 0.0113 |
| Tree pipit, within year | UniSpeech-SAT Base+ | 0.145 (0.018 to 0.325); 0.1816 | 0.167 (0.054 to 0.322); 0.0885 | 0.113 (0.027 to 0.233); 0.3851 | 0.162 (0.013 to 0.372); 0.1059 |
| Tree pipit, within year | XEUS | 0.139 (0.010 to 0.329); 0.2213 | 0.177 (0.048 to 0.372); 0.0718 | 0.160 (0.023 to 0.332); 0.0948 | 0.218 (0.020 to 0.449); 0.0157 |
| Tree pipit, within year | Loudness control | 0.228 (0.032 to 0.451); 0.0160 | 0.246 (0.052 to 0.475); 0.0063 | 0.215 (0.026 to 0.443); 0.0190 | 0.185 (0.045 to 0.378); 0.0603 |
| Tree pipit, within year | Duration control | 0.145 (0.000 to 0.336); 0.1041 | 0.126 (0.000 to 0.310); 0.2911 | 0.126 (0.000 to 0.355); 0.2220 | 0.116 (0.000 to 0.328); 0.2804 |
| Tree pipit, across year | BirdNET v2.4 | 0.214 (0.116 to 0.334); 0.0022 | 0.065 (0.029 to 0.102); 0.8142 | 0.075 (0.015 to 0.145); 0.7089 | 0.089 (0.020 to 0.184); 0.6125 |
| Tree pipit, across year | Perch 2.0 | 0.447 (0.293 to 0.624); 0.0001 | 0.163 (0.085 to 0.247); 0.0389 | 0.105 (0.020 to 0.275); 0.4218 | 0.093 (0.000 to 0.302); 0.5698 |
| Tree pipit, across year | Perch | 0.351 (0.219 to 0.518); 0.0001 | 0.078 (0.022 to 0.142); 0.7554 | 0.075 (0.033 to 0.128); 0.7816 | 0.086 (0.004 to 0.224); 0.6723 |
| Tree pipit, across year | SurfPerch | 0.243 (0.151 to 0.358); 0.0001 | 0.072 (0.039 to 0.105); 0.8485 | 0.072 (0.004 to 0.171); 0.8377 | 0.083 (0.031 to 0.139); 0.6997 |
| Tree pipit, across year | BirdNET v3 (preview) | 0.160 (0.095 to 0.227); 0.0388 | 0.150 (0.066 to 0.235); 0.0915 | 0.160 (0.036 to 0.294); 0.0677 | 0.093 (0.004 to 0.278); 0.5612 |
| Tree pipit, across year | AVEX sl-BEATs | 0.096 (0.060 to 0.130); 0.5067 | 0.039 (0.009 to 0.081); 0.9857 | 0.049 (0.007 to 0.115); 0.9577 | 0.042 (0.006 to 0.094); 0.9651 |
| Tree pipit, across year | AVEX EfficientNet-B0 | 0.211 (0.112 to 0.323); 0.0003 | 0.131 (0.067 to 0.207); 0.1525 | 0.101 (0.026 to 0.218); 0.4591 | 0.093 (0.029 to 0.172); 0.5766 |
| Tree pipit, across year | AvesEcho | 0.019 (0.000 to 0.039); 0.9988 | 0.033 (0.004 to 0.076); 0.9804 | 0.078 (0.000 to 0.244); 0.7513 | 0.099 (0.000 to 0.279); 0.4976 |
| Tree pipit, across year | AudioProtoPNet | 0.201 (0.103 to 0.315); 0.0023 | 0.095 (0.039 to 0.159); 0.5396 | 0.098 (0.019 to 0.210); 0.4980 | 0.086 (0.000 to 0.274); 0.6571 |
| Tree pipit, across year | ConvNeXt (BirdSet) | 0.288 (0.141 to 0.464); 0.0001 | 0.098 (0.037 to 0.155); 0.4963 | 0.072 (0.007 to 0.147); 0.7903 | 0.102 (0.000 to 0.275); 0.4558 |
| Tree pipit, across year | Bird-MAE | 0.032 (0.000 to 0.096); 0.9861 | 0.059 (0.000 to 0.169); 0.9225 | 0.065 (0.000 to 0.195); 0.8727 | 0.010 (0.000 to 0.020); 1.0000 |
| Tree pipit, across year | ProtoCLR | 0.083 (0.029 to 0.159); 0.6308 | 0.075 (0.025 to 0.139); 0.7971 | 0.105 (0.018 to 0.256); 0.4003 | 0.089 (0.008 to 0.246); 0.5785 |
| Tree pipit, across year | RCL_FS_BSED | 0.045 (0.016 to 0.072); 0.8746 | 0.013 (0.000 to 0.038); 0.9792 | 0.003 (0.000 to 0.011); 0.9956 | 0.099 (0.000 to 0.269); 0.4667 |
| Tree pipit, across year | BirdAVES | 0.083 (0.041 to 0.116); 0.6309 | 0.029 (0.000 to 0.078); 0.9714 | 0.023 (0.000 to 0.060); 0.9834 | 0.115 (0.020 to 0.198); 0.3404 |
| Tree pipit, across year | AVES | 0.038 (0.016 to 0.060); 0.9660 | 0.013 (0.000 to 0.035); 0.9982 | 0.042 (0.000 to 0.139); 0.9059 | 0.102 (0.014 to 0.192); 0.4460 |
| Tree pipit, across year | NatureBEATs | 0.070 (0.036 to 0.100); 0.7896 | 0.029 (0.005 to 0.059); 0.9965 | 0.049 (0.007 to 0.114); 0.9513 | 0.038 (0.006 to 0.081); 0.9745 |
| Tree pipit, across year | BioLingual | 0.058 (0.000 to 0.135); 0.8599 | 0.042 (0.013 to 0.088); 0.9608 | 0.078 (0.004 to 0.230); 0.7020 | 0.070 (0.000 to 0.224); 0.7916 |
| Tree pipit, across year | BEATs | 0.086 (0.045 to 0.130); 0.5970 | 0.072 (0.012 to 0.152); 0.7487 | 0.046 (0.016 to 0.083); 0.9355 | 0.054 (0.000 to 0.123); 0.8877 |
| Tree pipit, across year | AudioMAE | 0.045 (0.000 to 0.140); 0.9118 | 0.039 (0.000 to 0.098); 0.9699 | 0.052 (0.000 to 0.141); 0.9422 | 0.067 (0.000 to 0.219); 0.8696 |
| Tree pipit, across year | VGGish | 0.070 (0.034 to 0.110); 0.7720 | 0.114 (0.040 to 0.225); 0.3392 | 0.039 (0.010 to 0.077); 0.9427 | 0.112 (0.036 to 0.201); 0.3590 |
| Tree pipit, across year | ECAPA-TDNN (speaker) | 0.214 (0.114 to 0.344); 0.0006 | 0.118 (0.072 to 0.176); 0.2628 | 0.101 (0.008 to 0.262); 0.4581 | 0.147 (0.060 to 0.235); 0.0635 |
| Tree pipit, across year | ResNet (speaker) | 0.137 (0.066 to 0.231); 0.0940 | 0.072 (0.030 to 0.118); 0.8537 | 0.085 (0.010 to 0.220); 0.6758 | 0.153 (0.054 to 0.241); 0.0397 |

| Dataset | Embedding | C→C | B→B | C→B | B→C |
|---|---|---|---|---|---|
| Tree pipit, across year | x-vector (speaker) | 0.195 (0.080 to 0.354); 0.0064 | 0.069 (0.031 to 0.124); 0.8089 | 0.105 (0.013 to 0.275); 0.4229 | 0.089 (0.016 to 0.193); 0.6015 |
| Tree pipit, across year | wav2vec 2.0 Base | 0.064 (0.000 to 0.170); 0.7734 | 0.020 (0.004 to 0.037); 0.9893 | 0.069 (0.016 to 0.143); 0.7453 | 0.147 (0.000 to 0.389); 0.1706 |
| Tree pipit, across year | wav2vec 2.0 Large (robust) | 0.032 (0.006 to 0.062); 0.9397 | 0.026 (0.000 to 0.054); 0.9487 | 0.092 (0.000 to 0.216); 0.5296 | 0.099 (0.000 to 0.208); 0.4728 |
| Tree pipit, across year | XLS-R 300M | 0.022 (0.000 to 0.053); 0.9742 | 0.020 (0.000 to 0.038); 0.9720 | 0.082 (0.000 to 0.193); 0.6110 | 0.086 (0.000 to 0.167); 0.5659 |
| Tree pipit, across year | wav2vec 2.0 Conformer Large | 0.026 (0.000 to 0.056); 0.9657 | 0.029 (0.000 to 0.071); 0.9337 | 0.033 (0.000 to 0.098); 0.9308 | 0.115 (0.000 to 0.260); 0.3624 |
| Tree pipit, across year | MMS 300M | 0.019 (0.000 to 0.044); 0.9828 | 0.023 (0.000 to 0.046); 0.9656 | 0.065 (0.000 to 0.159); 0.7542 | 0.089 (0.000 to 0.173); 0.5453 |
| Tree pipit, across year | HuBERT Base | 0.013 (0.000 to 0.027); 0.9964 | 0.029 (0.000 to 0.061); 0.9587 | 0.000 (0.000 to 0.000); 1.0000 | 0.134 (0.000 to 0.355); 0.2332 |
| Tree pipit, across year | HuBERT Large | 0.032 (0.000 to 0.069); 0.9326 | 0.033 (0.007 to 0.056); 0.9168 | 0.078 (0.000 to 0.207); 0.6415 | 0.102 (0.000 to 0.213); 0.4550 |
| Tree pipit, across year | data2vec Base (100 h) | 0.029 (0.004 to 0.058); 0.9654 | 0.010 (0.000 to 0.027); 0.9967 | 0.029 (0.000 to 0.095); 0.9532 | 0.093 (0.000 to 0.205); 0.5158 |
| Tree pipit, across year | data2vec Base (960 h) | 0.029 (0.009 to 0.051); 0.9674 | 0.010 (0.000 to 0.024); 0.9965 | 0.033 (0.000 to 0.105); 0.9414 | 0.073 (0.000 to 0.150); 0.6765 |
| Tree pipit, across year | WavLM Base+ | 0.000 (0.000 to 0.000); 1.0000 | 0.026 (0.004 to 0.055); 0.9676 | 0.000 (0.000 to 0.000); 1.0000 | 0.118 (0.000 to 0.318); 0.3259 |
| Tree pipit, across year | WavLM Large | 0.026 (0.000 to 0.053); 0.9647 | 0.020 (0.004 to 0.036); 0.9693 | 0.059 (0.000 to 0.166); 0.7961 | 0.099 (0.011 to 0.208); 0.4818 |
| Tree pipit, across year | UniSpeech-SAT Base+ | 0.016 (0.003 to 0.031); 0.9935 | 0.026 (0.004 to 0.055); 0.9682 | 0.007 (0.000 to 0.021); 0.9989 | 0.131 (0.000 to 0.354); 0.2512 |
| Tree pipit, across year | XEUS | 0.010 (0.000 to 0.019); 0.9981 | 0.016 (0.000 to 0.037); 0.9837 | 0.007 (0.000 to 0.021); 0.9964 | 0.115 (0.009 to 0.254); 0.3542 |
| Tree pipit, across year | Loudness control | 0.073 (0.019 to 0.139); 0.7154 | 0.036 (0.000 to 0.086); 0.9528 | 0.075 (0.014 to 0.147); 0.6927 | 0.121 (0.000 to 0.338); 0.2985 |
| Tree pipit, across year | Duration control | 0.083 (0.000 to 0.242); 0.6719 | 0.072 (0.000 to 0.188); 0.8394 | 0.092 (0.000 to 0.253); 0.5670 | 0.073 (0.000 to 0.247); 0.9383 |
| Chiffchaff, across year | BirdNET v2.4 | 0.160 (0.035 to 0.333); 0.1327 | 0.132 (0.036 to 0.277); 0.2439 | 0.112 (0.005 to 0.318); 0.4019 | 0.110 (0.019 to 0.240); 0.4114 |
| Chiffchaff, across year | Perch 2.0 | 0.160 (0.043 to 0.293); 0.0916 | 0.127 (0.036 to 0.262); 0.2586 | 0.147 (0.017 to 0.337); 0.1542 | 0.125 (0.000 to 0.262); 0.2596 |
| Chiffchaff, across year | Perch | 0.150 (0.046 to 0.279); 0.1178 | 0.137 (0.043 to 0.281); 0.1783 | 0.173 (0.036 to 0.362); 0.0552 | 0.075 (0.011 to 0.156); 0.7837 |
| Chiffchaff, across year | SurfPerch | 0.255 (0.104 to 0.428); 0.0009 | 0.193 (0.050 to 0.387); 0.0152 | 0.188 (0.036 to 0.380); 0.0163 | 0.190 (0.006 to 0.445); 0.0122 |
| Chiffchaff, across year | BirdNET v3 (preview) | 0.085 (0.033 to 0.150); 0.6650 | 0.071 (0.026 to 0.132); 0.7682 | 0.096 (0.000 to 0.214); 0.5231 | 0.005 (0.000 to 0.016); 1.0000 |
| Chiffchaff, across year | AVEX sl-BEATs | 0.210 (0.073 to 0.364); 0.0133 | 0.173 (0.040 to 0.342); 0.0602 | 0.173 (0.000 to 0.360); 0.0580 | 0.130 (0.017 to 0.291); 0.2144 |
| Chiffchaff, across year | AVEX EfficientNet-B0 | 0.120 (0.035 to 0.236); 0.3143 | 0.147 (0.032 to 0.303); 0.1095 | 0.137 (0.031 to 0.292); 0.1738 | 0.085 (0.006 to 0.231); 0.6945 |
| Chiffchaff, across year | AvesEcho | 0.180 (0.015 to 0.383); 0.0780 | 0.086 (0.000 to 0.274); 0.5860 | 0.188 (0.000 to 0.391); 0.0755 | 0.120 (0.000 to 0.347); 0.3371 |
| Chiffchaff, across year | AudioProtoPNet | 0.115 (0.047 to 0.200); 0.3617 | 0.107 (0.024 to 0.236); 0.4211 | 0.142 (0.027 to 0.337); 0.1693 | 0.045 (0.000 to 0.114); 0.9732 |
| Chiffchaff, across year | ConvNeXt (BirdSet) | 0.135 (0.035 to 0.266); 0.2006 | 0.137 (0.027 to 0.296); 0.1908 | 0.137 (0.019 to 0.348); 0.2069 | 0.105 (0.009 to 0.236); 0.4655 |
| Chiffchaff, across year | Bird-MAE | 0.080 (0.014 to 0.178); 0.7886 | 0.081 (0.029 to 0.144); 0.6905 | 0.091 (0.025 to 0.180); 0.6050 | 0.120 (0.000 to 0.267); 0.2755 |
| Chiffchaff, across year | ProtoCLR | 0.195 (0.041 to 0.385); 0.0121 | 0.127 (0.021 to 0.332); 0.2535 | 0.117 (0.025 to 0.242); 0.3413 | 0.105 (0.034 to 0.190); 0.4577 |
| Chiffchaff, across year | RCL_FS_BSED | 0.150 (0.016 to 0.318); 0.2351 | 0.107 (0.000 to 0.274); 0.4498 | 0.142 (0.031 to 0.268); 0.2606 | 0.170 (0.000 to 0.416); 0.1482 |
| Chiffchaff, across year | BirdAVES | 0.190 (0.021 to 0.399); 0.0691 | 0.107 (0.000 to 0.313); 0.4261 | 0.188 (0.034 to 0.369); 0.0671 | 0.105 (0.000 to 0.330); 0.4518 |
| Chiffchaff, across year | AVES | 0.205 (0.000 to 0.414); 0.0508 | 0.102 (0.000 to 0.312); 0.4544 | 0.203 (0.023 to 0.402); 0.0426 | 0.155 (0.000 to 0.397); 0.1568 |
| Chiffchaff, across year | NatureBEATs | 0.160 (0.032 to 0.304); 0.1046 | 0.107 (0.015 to 0.271); 0.4273 | 0.188 (0.000 to 0.384); 0.0373 | 0.105 (0.010 to 0.269); 0.4491 |
| Chiffchaff, across year | BioLingual | 0.160 (0.045 to 0.290); 0.0988 | 0.173 (0.064 to 0.316); 0.0415 | 0.137 (0.029 to 0.314); 0.1900 | 0.120 (0.000 to 0.302); 0.2969 |
| Chiffchaff, across year | BEATs | 0.155 (0.042 to 0.289); 0.1273 | 0.117 (0.019 to 0.271); 0.3498 | 0.147 (0.023 to 0.274); 0.1735 | 0.150 (0.000 to 0.368); 0.1404 |
| Chiffchaff, across year | AudioMAE | 0.150 (0.045 to 0.281); 0.1418 | 0.173 (0.021 to 0.374); 0.0523 | 0.157 (0.000 to 0.360); 0.0914 | 0.155 (0.000 to 0.379); 0.0916 |
| Chiffchaff, across year | VGGish | 0.095 (0.000 to 0.278); 0.5181 | 0.066 (0.000 to 0.210); 0.7368 | 0.046 (0.000 to 0.117); 0.8757 | 0.155 (0.000 to 0.369); 0.1480 |
| Chiffchaff, across year | ECAPA-TDNN (speaker) | 0.150 (0.056 to 0.280); 0.1265 | 0.147 (0.012 to 0.306); 0.1281 | 0.147 (0.027 to 0.354); 0.1550 | 0.170 (0.012 to 0.384); 0.0415 |
| Chiffchaff, across year | ResNet (speaker) | 0.145 (0.046 to 0.290); 0.1644 | 0.112 (0.022 to 0.226); 0.3711 | 0.086 (0.022 to 0.168); 0.6370 | 0.130 (0.022 to 0.273); 0.2330 |
| Chiffchaff, across year | x-vector (speaker) | 0.140 (0.032 to 0.318); 0.2328 | 0.168 (0.000 to 0.389); 0.1158 | 0.122 (0.011 to 0.319); 0.3304 | 0.110 (0.016 to 0.229); 0.4192 |
| Chiffchaff, across year | wav2vec 2.0 Base | 0.165 (0.000 to 0.389); 0.1430 | 0.132 (0.016 to 0.322); 0.2649 | 0.223 (0.000 to 0.478); 0.0144 | 0.175 (0.000 to 0.395); 0.0945 |
| Chiffchaff, across year | wav2vec 2.0 Large (robust) | 0.210 (0.016 to 0.433); 0.0486 | 0.102 (0.000 to 0.283); 0.4563 | 0.178 (0.039 to 0.322); 0.0880 | 0.170 (0.000 to 0.418); 0.1285 |
| Chiffchaff, across year | XLS-R 300M | 0.235 (0.056 to 0.441); 0.0213 | 0.117 (0.000 to 0.321); 0.3510 | 0.193 (0.033 to 0.359); 0.0548 | 0.175 (0.000 to 0.429); 0.1096 |
| Chiffchaff, across year | wav2vec 2.0 Conformer Large | 0.205 (0.000 to 0.389); 0.0497 | 0.107 (0.000 to 0.286); 0.4178 | 0.183 (0.023 to 0.353); 0.0690 | 0.160 (0.000 to 0.394); 0.1511 |
| Chiffchaff, across year | MMS 300M | 0.220 (0.032 to 0.436); 0.0347 | 0.107 (0.000 to 0.295); 0.4213 | 0.198 (0.029 to 0.374); 0.0448 | 0.165 (0.000 to 0.410); 0.1395 |
| Chiffchaff, across year | HuBERT Base | 0.125 (0.000 to 0.348); 0.3242 | 0.112 (0.000 to 0.313); 0.3894 | 0.193 (0.017 to 0.440); 0.0392 | 0.155 (0.000 to 0.393); 0.1620 |

| Dataset | Embedding | C→C | B→B | C→B | B→C |
|---|---|---|---|---|---|
| Chiffchaff, across year | HuBERT Large | 0.215 (0.000 to 0.445); 0.0422 | 0.127 (0.000 to 0.339); 0.2895 | 0.208 (0.043 to 0.380); 0.0379 | 0.180 (0.000 to 0.438); 0.0956 |
| Chiffchaff, across year | data2vec Base (100 h) | 0.180 (0.000 to 0.398); 0.0995 | 0.107 (0.000 to 0.308); 0.4238 | 0.152 (0.000 to 0.321); 0.1637 | 0.175 (0.000 to 0.414); 0.1020 |
| Chiffchaff, across year | data2vec Base (960 h) | 0.180 (0.000 to 0.394); 0.1000 | 0.096 (0.000 to 0.286); 0.4923 | 0.152 (0.000 to 0.330); 0.1596 | 0.155 (0.000 to 0.379); 0.1647 |
| Chiffchaff, across year | WavLM Base+ | 0.125 (0.000 to 0.348); 0.3240 | 0.102 (0.000 to 0.303); 0.4601 | 0.218 (0.034 to 0.451); 0.0193 | 0.150 (0.016 to 0.367); 0.1906 |
| Chiffchaff, across year | WavLM Large | 0.200 (0.024 to 0.386); 0.0611 | 0.107 (0.000 to 0.291); 0.4201 | 0.168 (0.030 to 0.315); 0.1099 | 0.185 (0.000 to 0.451); 0.0810 |
| Chiffchaff, across year | UniSpeech-SAT Base+ | 0.130 (0.000 to 0.349); 0.2931 | 0.102 (0.000 to 0.293); 0.4630 | 0.234 (0.052 to 0.467); 0.0093 | 0.150 (0.000 to 0.389); 0.1823 |
| Chiffchaff, across year | XEUS | 0.205 (0.016 to 0.407); 0.0551 | 0.107 (0.006 to 0.292); 0.4219 | 0.218 (0.016 to 0.412); 0.0258 | 0.135 (0.000 to 0.339); 0.2592 |
| Chiffchaff, across year | Loudness control | 0.025 (0.000 to 0.058); 0.9672 | 0.036 (0.000 to 0.077); 0.9682 | 0.086 (0.000 to 0.238); 0.6503 | 0.030 (0.000 to 0.094); 0.9676 |
| Chiffchaff, across year | Duration control | 0.105 (0.000 to 0.297); 0.4988 | 0.117 (0.000 to 0.324); 0.3154 | 0.041 (0.000 to 0.138); 0.9426 | 0.085 (0.000 to 0.240); 0.6863 |

*Notes:* C→B: classifier fitted on calls and tested on backgrounds; the other columns likewise. Each cell: re-ID accuracy (95% interval over individuals, 10,000 draws); permutation p (9,999 shuffles), unadjusted. Entries whose B→B p is below 0.05 after Holm's correction over the 38 entries of a dataset: Chiffchaff, within year 35; Little owl, across year 1; Tree pipit, within year 8; Tree pipit, across year 0; Chiffchaff, across year 0 (unadjusted: 35; 8; 16; 1; 2; about 2 of 38 expected by chance).

**Table S6.** Effect of adding a background recording to test clips, for all 38 embeddings and controls: own-donor minus other-donor balanced accuracy.

| Dataset | Embedding | Background +10 dB | Background 0 dB | Background −10 dB |
|---|---|---|---|---|
| Chiffchaff, within year | BirdNET v2.4 | 0.325 (0.194 to 0.465) | 0.219 (0.089 to 0.362) | 0.095 (0.035 to 0.165) |
| Chiffchaff, within year | Perch 2.0 | 0.450 (0.300 to 0.591) | 0.337 (0.221 to 0.461) | 0.124 (0.054 to 0.204) |
| Chiffchaff, within year | Perch | 0.458 (0.305 to 0.613) | 0.329 (0.183 to 0.492) | 0.151 (0.060 to 0.268) |
| Chiffchaff, within year | SurfPerch | 0.215 (0.103 to 0.343) | 0.164 (0.078 to 0.269) | 0.067 (0.028 to 0.106) |
| Chiffchaff, within year | BirdNET v3 (preview) | 0.311 (0.188 to 0.457) | 0.250 (0.144 to 0.378) | 0.107 (0.039 to 0.176) |
| Chiffchaff, within year | AVEX sl-BEATs | 0.215 (0.090 to 0.351) | 0.184 (0.080 to 0.303) | 0.112 (0.066 to 0.161) |
| Chiffchaff, within year | AVEX EfficientNet-B0 | 0.243 (0.124 to 0.371) | 0.225 (0.129 to 0.330) | 0.098 (0.049 to 0.148) |
| Chiffchaff, within year | AvesEcho | 0.105 (0.031 to 0.185) | 0.053 (−0.014 to 0.123) | 0.072 (0.019 to 0.127) |
| Chiffchaff, within year | AudioProtoPNet | 0.429 (0.278 to 0.582) | 0.348 (0.208 to 0.498) | 0.215 (0.104 to 0.340) |
| Chiffchaff, within year | ConvNeXt (BirdSet) | 0.417 (0.250 to 0.585) | 0.314 (0.181 to 0.451) | 0.200 (0.120 to 0.291) |
| Chiffchaff, within year | Bird-MAE | 0.082 (0.0003 to 0.192) | 0.073 (−0.001 to 0.172) | 0.031 (−0.002 to 0.073) |
| Chiffchaff, within year | ProtoCLR | 0.118 (0.053 to 0.192) | 0.130 (0.041 to 0.250) | 0.080 (0.028 to 0.147) |
| Chiffchaff, within year | RCL_FS_BSED | 0.018 (−0.014 to 0.064) | 0.022 (−0.002 to 0.067) | 0.029 (0.000 to 0.084) |
| Chiffchaff, within year | BirdAVES | 0.105 (0.003 to 0.221) | 0.080 (0.005 to 0.161) | 0.075 (0.027 to 0.138) |
| Chiffchaff, within year | AVES | 0.058 (−0.016 to 0.142) | 0.050 (0.009 to 0.108) | 0.038 (−0.009 to 0.089) |
| Chiffchaff, within year | NatureBEATs | 0.113 (0.033 to 0.214) | 0.109 (0.029 to 0.217) | 0.107 (0.040 to 0.188) |
| Chiffchaff, within year | BioLingual | 0.250 (0.124 to 0.398) | 0.157 (0.044 to 0.290) | 0.074 (0.025 to 0.118) |
| Chiffchaff, within year | BEATs | 0.229 (0.095 to 0.371) | 0.210 (0.075 to 0.346) | 0.155 (0.073 to 0.246) |
| Chiffchaff, within year | AudioMAE | −0.013 (−0.081 to 0.031) | 0.018 (−0.003 to 0.048) | 0.001 (−0.007 to 0.009) |
| Chiffchaff, within year | VGGish | 0.066 (−0.028 to 0.166) | 0.071 (0.012 to 0.139) | 0.046 (−0.020 to 0.106) |
| Chiffchaff, within year | ECAPA-TDNN (speaker) | 0.178 (0.074 to 0.283) | 0.172 (0.073 to 0.289) | 0.085 (0.011 to 0.172) |
| Chiffchaff, within year | ResNet (speaker) | 0.222 (0.117 to 0.341) | 0.187 (0.093 to 0.294) | 0.110 (0.047 to 0.179) |
| Chiffchaff, within year | x-vector (speaker) | 0.198 (0.081 to 0.331) | 0.195 (0.083 to 0.317) | 0.092 (0.041 to 0.148) |
| Chiffchaff, within year | wav2vec 2.0 Base | 0.140 (0.025 to 0.265) | 0.144 (0.041 to 0.259) | 0.046 (−0.018 to 0.122) |
| Chiffchaff, within year | wav2vec 2.0 Large (robust) | 0.088 (−0.008 to 0.196) | 0.061 (0.002 to 0.145) | 0.044 (−0.009 to 0.106) |
| Chiffchaff, within year | XLS-R 300M | 0.049 (−0.071 to 0.166) | 0.087 (0.020 to 0.176) | 0.052 (0.004 to 0.108) |
| Chiffchaff, within year | wav2vec 2.0 Conformer Large | 0.189 (0.032 to 0.352) | 0.224 (0.100 to 0.363) | 0.096 (0.025 to 0.169) |
| Chiffchaff, within year | MMS 300M | 0.066 (−0.014 to 0.159) | 0.085 (0.019 to 0.176) | 0.042 (−0.008 to 0.103) |
| Chiffchaff, within year | HuBERT Base | 0.098 (−0.020 to 0.224) | 0.133 (0.023 to 0.262) | 0.053 (−0.019 to 0.149) |
| Chiffchaff, within year | HuBERT Large | 0.086 (−0.021 to 0.221) | 0.093 (−0.002 to 0.212) | 0.036 (−0.019 to 0.111) |
| Chiffchaff, within year | data2vec Base (100 h) | 0.125 (0.037 to 0.218) | 0.113 (0.015 to 0.213) | 0.059 (−0.014 to 0.134) |
| Chiffchaff, within year | data2vec Base (960 h) | 0.150 (0.049 to 0.266) | 0.140 (0.063 to 0.231) | 0.057 (−0.017 to 0.136) |
| Chiffchaff, within year | WavLM Base+ | 0.142 (0.012 to 0.274) | 0.155 (0.025 to 0.304) | 0.091 (0.010 to 0.194) |
| Chiffchaff, within year | WavLM Large | 0.078 (−0.040 to 0.208) | 0.090 (0.004 to 0.197) | 0.049 (−0.007 to 0.118) |
| Chiffchaff, within year | UniSpeech-SAT Base+ | 0.141 (0.026 to 0.253) | 0.115 (0.007 to 0.244) | 0.059 (−0.008 to 0.146) |
| Chiffchaff, within year | XEUS | 0.129 (0.039 to 0.228) | 0.122 (0.036 to 0.217) | 0.033 (−0.021 to 0.093) |
| Chiffchaff, within year | Loudness control | 0.004 (0.000 to 0.013) | −0.002 (−0.007 to 0.000) | 0.000 (0.000 to 0.000) |
| Chiffchaff, within year | Duration control | 0.000 (0.000 to 0.000) | 0.000 (0.000 to 0.000) | 0.000 (0.000 to 0.000) |
| Little owl, across year | BirdNET v2.4 | −0.100 (−0.268 to 0.051) | −0.071 (−0.179 to 0.026) | 0.000 (−0.033 to 0.035) |
| Little owl, across year | Perch 2.0 | −0.069 (−0.197 to 0.047) | −0.033 (−0.076 to 0.001) | 0.003 (−0.034 to 0.036) |
| Little owl, across year | Perch | −0.052 (−0.133 to 0.020) | −0.038 (−0.085 to 0.010) | −0.007 (−0.035 to 0.022) |
| Little owl, across year | SurfPerch | −0.018 (−0.095 to 0.055) | −0.002 (−0.084 to 0.079) | −0.016 (−0.071 to 0.037) |
| Little owl, across year | BirdNET v3 (preview) | 0.021 (−0.073 to 0.116) | −0.040 (−0.107 to 0.024) | 0.022 (−0.015 to 0.064) |
| Little owl, across year | AVEX sl-BEATs | 0.026 (−0.006 to 0.083) | 0.021 (0.000 to 0.062) | 0.012 (−0.004 to 0.032) |
| Little owl, across year | AVEX EfficientNet-B0 | −0.074 (−0.167 to 0.017) | −0.025 (−0.084 to 0.035) | −0.023 (−0.047 to 0.001) |

| Dataset | Embedding | Background +10 dB | Background 0 dB | Background −10 dB |
|---|---|---|---|---|
| Little owl, across year | AvesEcho | −0.002 (−0.013 to 0.007) | −0.002 (−0.013 to 0.007) | −0.006 (−0.019 to 0.000) |
| Little owl, across year | AudioProtoPNet | −0.103 (−0.211 to −0.003) | −0.078 (−0.154 to −0.011) | −0.047 (−0.092 to −0.011) |
| Little owl, across year | ConvNeXt (BirdSet) | −0.041 (−0.119 to 0.044) | −0.044 (−0.112 to 0.021) | −0.019 (−0.062 to 0.018) |
| Little owl, across year | Bird-MAE | −0.047 (−0.119 to 0.008) | −0.037 (−0.086 to 0.007) | −0.010 (−0.027 to 0.006) |
| Little owl, across year | ProtoCLR | 0.049 (−0.015 to 0.158) | 0.014 (−0.039 to 0.092) | 0.013 (−0.023 to 0.056) |
| Little owl, across year | RCL_FS_BSED | −0.016 (−0.047 to 0.000) | −0.025 (−0.074 to 0.000) | −0.007 (−0.020 to 0.000) |
| Little owl, across year | BirdAVES | −0.039 (−0.128 to 0.034) | −0.033 (−0.094 to 0.015) | −0.020 (−0.063 to 0.018) |
| Little owl, across year | AVES | −0.007 (−0.076 to 0.054) | −0.028 (−0.089 to 0.017) | −0.024 (−0.067 to 0.009) |
| Little owl, across year | NatureBEATs | 0.007 (−0.022 to 0.042) | 0.003 (0.000 to 0.010) | 0.000 (0.000 to 0.000) |
| Little owl, across year | BioLingual | 0.000 (0.000 to 0.000) | 0.000 (0.000 to 0.000) | 0.000 (0.000 to 0.000) |
| Little owl, across year | BEATs | 0.000 (0.000 to 0.000) | 0.000 (0.000 to 0.000) | −0.002 (−0.006 to 0.000) |
| Little owl, across year | AudioMAE | 0.026 (0.004 to 0.052) | −0.009 (−0.033 to 0.013) | 0.000 (0.000 to 0.000) |
| Little owl, across year | VGGish | −0.009 (−0.082 to 0.083) | 0.027 (−0.021 to 0.106) | 0.016 (−0.006 to 0.043) |
| Little owl, across year | ECAPA-TDNN (speaker) | 0.008 (−0.113 to 0.115) | −0.003 (−0.058 to 0.044) | −0.005 (−0.044 to 0.036) |
| Little owl, across year | ResNet (speaker) | −0.045 (−0.143 to 0.038) | −0.058 (−0.153 to 0.012) | −0.031 (−0.063 to −0.001) |
| Little owl, across year | x-vector (speaker) | 0.010 (−0.068 to 0.080) | −0.029 (−0.093 to 0.028) | −0.033 (−0.054 to −0.013) |
| Little owl, across year | wav2vec 2.0 Base | 0.001 (−0.037 to 0.041) | −0.010 (−0.065 to 0.035) | −0.035 (−0.061 to −0.009) |
| Little owl, across year | wav2vec 2.0 Large (robust) | −0.041 (−0.128 to 0.032) | −0.077 (−0.181 to 0.024) | −0.004 (−0.041 to 0.036) |
| Little owl, across year | XLS-R 300M | −0.080 (−0.158 to −0.008) | −0.079 (−0.187 to 0.028) | −0.018 (−0.061 to 0.025) |
| Little owl, across year | wav2vec 2.0 Conformer Large | −0.084 (−0.191 to 0.009) | −0.059 (−0.138 to 0.016) | −0.021 (−0.066 to 0.032) |
| Little owl, across year | MMS 300M | −0.096 (−0.170 to −0.023) | −0.122 (−0.242 to −0.013) | −0.029 (−0.074 to 0.013) |
| Little owl, across year | HuBERT Base | −0.006 (−0.127 to 0.100) | −0.038 (−0.120 to 0.037) | −0.022 (−0.066 to 0.015) |
| Little owl, across year | HuBERT Large | −0.102 (−0.194 to −0.022) | −0.074 (−0.157 to 0.011) | −0.026 (−0.059 to 0.003) |
| Little owl, across year | data2vec Base (100 h) | −0.024 (−0.142 to 0.079) | −0.090 (−0.153 to −0.033) | −0.045 (−0.082 to −0.007) |
| Little owl, across year | data2vec Base (960 h) | −0.045 (−0.152 to 0.049) | −0.109 (−0.175 to −0.051) | −0.071 (−0.154 to −0.009) |
| Little owl, across year | WavLM Base+ | −0.026 (−0.122 to 0.068) | −0.076 (−0.155 to −0.009) | −0.017 (−0.058 to 0.022) |
| Little owl, across year | WavLM Large | −0.101 (−0.183 to −0.032) | −0.106 (−0.209 to −0.010) | −0.050 (−0.109 to −0.001) |
| Little owl, across year | UniSpeech-SAT Base+ | 0.004 (−0.107 to 0.121) | −0.047 (−0.146 to 0.044) | −0.030 (−0.078 to 0.014) |
| Little owl, across year | XEUS | −0.023 (−0.135 to 0.086) | −0.061 (−0.144 to 0.030) | −0.020 (−0.066 to 0.028) |
| Little owl, across year | Loudness control | 0.012 (0.000 to 0.033) | 0.020 (0.000 to 0.054) | 0.005 (−0.011 to 0.026) |
| Little owl, across year | Duration control | 0.000 (0.000 to 0.000) | 0.000 (0.000 to 0.000) | 0.000 (0.000 to 0.000) |
| Tree pipit, within year | BirdNET v2.4 | 0.043 (−0.036 to 0.131) | −0.003 (−0.068 to 0.057) | −0.027 (−0.091 to 0.031) |
| Tree pipit, within year | Perch 2.0 | −0.012 (−0.172 to 0.134) | −0.001 (−0.117 to 0.110) | −0.004 (−0.102 to 0.072) |
| Tree pipit, within year | Perch | 0.058 (−0.101 to 0.207) | 0.031 (−0.083 to 0.141) | −0.025 (−0.112 to 0.035) |
| Tree pipit, within year | SurfPerch | 0.026 (−0.110 to 0.162) | −0.027 (−0.158 to 0.096) | −0.075 (−0.151 to −0.007) |
| Tree pipit, within year | BirdNET v3 (preview) | 0.066 (0.007 to 0.152) | 0.063 (0.003 to 0.142) | 0.009 (−0.030 to 0.065) |
| Tree pipit, within year | AVEX sl-BEATs | 0.025 (−0.025 to 0.079) | 0.005 (−0.054 to 0.064) | −0.026 (−0.076 to 0.017) |
| Tree pipit, within year | AVEX EfficientNet-B0 | 0.010 (−0.062 to 0.073) | 0.045 (−0.010 to 0.100) | 0.019 (0.006 to 0.033) |
| Tree pipit, within year | AvesEcho | 0.015 (−0.020 to 0.064) | 0.012 (−0.017 to 0.043) | −0.022 (−0.061 to 0.004) |
| Tree pipit, within year | AudioProtoPNet | 0.125 (−0.043 to 0.264) | 0.072 (−0.092 to 0.195) | −0.026 (−0.139 to 0.052) |
| Tree pipit, within year | ConvNeXt (BirdSet) | 0.137 (0.008 to 0.271) | 0.090 (−0.012 to 0.196) | 0.012 (−0.091 to 0.099) |
| Tree pipit, within year | Bird-MAE | 0.005 (−0.013 to 0.027) | 0.003 (0.000 to 0.009) | 0.006 (−0.011 to 0.027) |
| Tree pipit, within year | ProtoCLR | −0.024 (−0.057 to 0.006) | −0.021 (−0.053 to 0.008) | 0.006 (−0.008 to 0.026) |
| Tree pipit, within year | RCL_FS_BSED | 0.105 (0.000 to 0.292) | 0.070 (−0.014 to 0.225) | 0.029 (−0.034 to 0.121) |
| Tree pipit, within year | BirdAVES | 0.024 (−0.124 to 0.190) | 0.011 (−0.140 to 0.157) | −0.014 (−0.081 to 0.053) |
| Tree pipit, within year | AVES | 0.087 (−0.039 to 0.274) | 0.040 (−0.090 to 0.214) | −0.038 (−0.089 to 0.003) |
| Tree pipit, within year | NatureBEATs | 0.018 (−0.014 to 0.055) | −0.003 (−0.063 to 0.058) | 0.005 (−0.033 to 0.045) |

| Dataset | Embedding | Background +10 dB | Background 0 dB | Background −10 dB |
|---|---|---|---|---|
| Tree pipit, within year | BioLingual | 0.003 (−0.013 to 0.024) | −0.012 (−0.024 to 0.000) | −0.000 (−0.021 to 0.016) |
| Tree pipit, within year | BEATs | 0.087 (0.001 to 0.206) | 0.070 (−0.001 to 0.152) | 0.012 (−0.030 to 0.050) |
| Tree pipit, within year | AudioMAE | 0.028 (−0.032 to 0.114) | −0.004 (−0.050 to 0.037) | −0.004 (−0.018 to 0.011) |
| Tree pipit, within year | VGGish | −0.012 (−0.103 to 0.104) | −0.055 (−0.109 to −0.005) | −0.024 (−0.070 to 0.017) |
| Tree pipit, within year | ECAPA-TDNN (speaker) | 0.001 (−0.072 to 0.065) | −0.047 (−0.134 to 0.031) | −0.041 (−0.102 to 0.015) |
| Tree pipit, within year | ResNet (speaker) | 0.039 (−0.026 to 0.111) | −0.015 (−0.057 to 0.023) | −0.051 (−0.119 to 0.010) |
| Tree pipit, within year | x-vector (speaker) | −0.037 (−0.124 to 0.027) | −0.031 (−0.092 to 0.018) | −0.005 (−0.038 to 0.028) |
| Tree pipit, within year | wav2vec 2.0 Base | 0.015 (−0.089 to 0.135) | 0.016 (−0.164 to 0.212) | 0.010 (−0.061 to 0.108) |
| Tree pipit, within year | wav2vec 2.0 Large (robust) | 0.028 (−0.161 to 0.279) | −0.007 (−0.200 to 0.244) | 0.005 (−0.115 to 0.167) |
| Tree pipit, within year | XLS-R 300M | 0.045 (−0.156 to 0.292) | −0.004 (−0.197 to 0.244) | −0.020 (−0.107 to 0.067) |
| Tree pipit, within year | wav2vec 2.0 Conformer Large | 0.042 (−0.144 to 0.286) | −0.029 (−0.238 to 0.230) | −0.021 (−0.131 to 0.104) |
| Tree pipit, within year | MMS 300M | 0.043 (−0.157 to 0.289) | 0.001 (−0.191 to 0.240) | −0.021 (−0.115 to 0.077) |
| Tree pipit, within year | HuBERT Base | −0.020 (−0.209 to 0.216) | −0.011 (−0.206 to 0.226) | 0.047 (−0.087 to 0.199) |
| Tree pipit, within year | HuBERT Large | 0.035 (−0.118 to 0.225) | 0.031 (−0.145 to 0.263) | 0.021 (−0.100 to 0.169) |
| Tree pipit, within year | data2vec Base (100 h) | 0.014 (−0.169 to 0.258) | 0.011 (−0.170 to 0.247) | 0.041 (−0.071 to 0.207) |
| Tree pipit, within year | data2vec Base (960 h) | 0.009 (−0.189 to 0.252) | −0.005 (−0.187 to 0.227) | 0.017 (−0.095 to 0.167) |
| Tree pipit, within year | WavLM Base+ | 0.000 (−0.214 to 0.257) | −0.007 (−0.245 to 0.252) | 0.029 (−0.080 to 0.155) |
| Tree pipit, within year | WavLM Large | 0.034 (−0.106 to 0.231) | 0.004 (−0.185 to 0.239) | −0.012 (−0.130 to 0.133) |
| Tree pipit, within year | UniSpeech-SAT Base+ | 0.037 (−0.171 to 0.282) | 0.001 (−0.209 to 0.236) | 0.031 (−0.078 to 0.151) |
| Tree pipit, within year | XEUS | 0.002 (−0.245 to 0.279) | −0.027 (−0.274 to 0.240) | −0.003 (−0.140 to 0.164) |
| Tree pipit, within year | Loudness control | 0.078 (0.000 to 0.172) | 0.033 (−0.003 to 0.079) | 0.003 (0.000 to 0.010) |
| Tree pipit, within year | Duration control | 0.000 (0.000 to 0.000) | 0.000 (0.000 to 0.000) | 0.000 (0.000 to 0.000) |
| Tree pipit, across year | BirdNET v2.4 | −0.104 (−0.284 to 0.045) | −0.090 (−0.206 to 0.022) | −0.028 (−0.083 to 0.030) |
| Tree pipit, across year | Perch 2.0 | −0.003 (−0.140 to 0.118) | 0.024 (−0.039 to 0.086) | −0.013 (−0.070 to 0.047) |
| Tree pipit, across year | Perch | −0.095 (−0.239 to 0.049) | −0.052 (−0.179 to 0.079) | −0.047 (−0.154 to 0.052) |
| Tree pipit, across year | SurfPerch | −0.103 (−0.251 to 0.012) | −0.038 (−0.148 to 0.045) | 0.009 (−0.040 to 0.064) |
| Tree pipit, across year | BirdNET v3 (preview) | 0.002 (−0.074 to 0.072) | −0.011 (−0.068 to 0.035) | −0.004 (−0.032 to 0.021) |
| Tree pipit, across year | AVEX sl-BEATs | −0.137 (−0.320 to −0.013) | −0.096 (−0.250 to 0.004) | −0.055 (−0.172 to 0.022) |
| Tree pipit, across year | AVEX EfficientNet-B0 | −0.056 (−0.189 to 0.049) | −0.050 (−0.111 to 0.007) | −0.009 (−0.054 to 0.049) |
| Tree pipit, across year | AvesEcho | −0.055 (−0.126 to 0.004) | −0.049 (−0.120 to 0.001) | −0.011 (−0.040 to 0.015) |
| Tree pipit, across year | AudioProtoPNet | −0.070 (−0.234 to 0.078) | −0.052 (−0.201 to 0.081) | −0.054 (−0.152 to 0.029) |
| Tree pipit, across year | ConvNeXt (BirdSet) | −0.096 (−0.271 to 0.044) | −0.110 (−0.259 to 0.016) | −0.061 (−0.143 to 0.019) |
| Tree pipit, across year | Bird-MAE | −0.033 (−0.080 to 0.000) | −0.023 (−0.063 to 0.000) | −0.016 (−0.037 to 0.000) |
| Tree pipit, across year | ProtoCLR | −0.015 (−0.040 to 0.000) | −0.004 (−0.025 to 0.012) | −0.004 (−0.012 to 0.004) |
| Tree pipit, across year | RCL_FS_BSED | −0.143 (−0.337 to 0.000) | −0.080 (−0.213 to 0.000) | −0.053 (−0.160 to 0.000) |
| Tree pipit, across year | BirdAVES | −0.181 (−0.409 to −0.010) | −0.161 (−0.368 to 0.000) | −0.107 (−0.256 to 0.002) |
| Tree pipit, across year | AVES | −0.194 (−0.434 to −0.013) | −0.167 (−0.387 to 0.000) | −0.084 (−0.197 to 0.000) |
| Tree pipit, across year | NatureBEATs | −0.108 (−0.293 to 0.013) | −0.099 (−0.271 to 0.003) | −0.067 (−0.227 to 0.023) |
| Tree pipit, across year | BioLingual | −0.047 (−0.137 to 0.007) | −0.023 (−0.080 to 0.010) | −0.017 (−0.043 to 0.000) |
| Tree pipit, across year | BEATs | −0.129 (−0.292 to −0.008) | −0.076 (−0.213 to 0.016) | −0.043 (−0.117 to 0.000) |
| Tree pipit, across year | AudioMAE | −0.010 (−0.030 to 0.000) | −0.007 (−0.020 to 0.000) | −0.003 (−0.010 to 0.000) |
| Tree pipit, across year | VGGish | −0.070 (−0.197 to 0.000) | −0.006 (−0.031 to 0.024) | −0.012 (−0.047 to 0.017) |
| Tree pipit, across year | ECAPA-TDNN (speaker) | −0.053 (−0.099 to −0.010) | 0.021 (−0.051 to 0.099) | −0.033 (−0.078 to 0.014) |
| Tree pipit, across year | ResNet (speaker) | −0.000 (−0.072 to 0.086) | −0.010 (−0.089 to 0.068) | 0.003 (−0.027 to 0.034) |
| Tree pipit, across year | x-vector (speaker) | −0.030 (−0.057 to −0.007) | 0.024 (−0.024 to 0.075) | −0.043 (−0.082 to −0.004) |
| Tree pipit, across year | wav2vec 2.0 Base | −0.107 (−0.285 to 0.018) | −0.062 (−0.140 to 0.005) | −0.034 (−0.063 to −0.011) |
| Tree pipit, across year | wav2vec 2.0 Large (robust) | −0.075 (−0.274 to 0.045) | −0.062 (−0.199 to 0.013) | 0.001 (−0.021 to 0.022) |

| Dataset | Embedding | Background +10 dB | Background 0 dB | Background −10 dB |
|---|---|---|---|---|
| Tree pipit, across year | XLS-R 300M | −0.074 (−0.274 to 0.043) | −0.054 (−0.177 to 0.013) | 0.010 (−0.013 to 0.033) |
| Tree pipit, across year | wav2vec 2.0 Conformer Large | −0.098 (−0.279 to 0.000) | −0.086 (−0.251 to 0.007) | −0.006 (−0.052 to 0.034) |
| Tree pipit, across year | MMS 300M | −0.076 (−0.276 to 0.041) | −0.059 (−0.190 to 0.016) | 0.007 (−0.013 to 0.027) |
| Tree pipit, across year | HuBERT Base | −0.224 (−0.454 to −0.028) | −0.111 (−0.226 to −0.020) | −0.022 (−0.046 to −0.001) |
| Tree pipit, across year | HuBERT Large | −0.068 (−0.248 to 0.040) | −0.067 (−0.210 to 0.010) | −0.019 (−0.041 to 0.000) |
| Tree pipit, across year | data2vec Base (100 h) | −0.082 (−0.274 to 0.035) | −0.067 (−0.187 to 0.011) | −0.006 (−0.042 to 0.024) |
| Tree pipit, across year | data2vec Base (960 h) | −0.092 (−0.276 to 0.023) | −0.064 (−0.191 to 0.016) | −0.007 (−0.042 to 0.022) |
| Tree pipit, across year | WavLM Base+ | −0.235 (−0.482 to −0.039) | −0.120 (−0.253 to −0.018) | −0.012 (−0.034 to 0.004) |
| Tree pipit, across year | WavLM Large | −0.080 (−0.274 to 0.040) | −0.064 (−0.194 to 0.007) | −0.004 (−0.020 to 0.007) |
| Tree pipit, across year | UniSpeech-SAT Base+ | −0.175 (−0.383 to −0.021) | −0.060 (−0.150 to −0.004) | −0.009 (−0.034 to 0.010) |
| Tree pipit, across year | XEUS | −0.082 (−0.268 to 0.019) | −0.075 (−0.251 to 0.025) | −0.019 (−0.070 to 0.013) |
| Tree pipit, across year | Loudness control | −0.018 (−0.100 to 0.048) | −0.015 (−0.084 to 0.043) | −0.009 (−0.028 to 0.000) |
| Tree pipit, across year | Duration control | 0.000 (0.000 to 0.000) | 0.000 (0.000 to 0.000) | 0.000 (0.000 to 0.000) |
| Chiffchaff, across year | BirdNET v2.4 | −0.034 (−0.182 to 0.131) | −0.067 (−0.174 to 0.046) | −0.067 (−0.159 to 0.010) |
| Chiffchaff, across year | Perch 2.0 | −0.083 (−0.222 to 0.064) | −0.048 (−0.132 to 0.021) | −0.003 (−0.051 to 0.046) |
| Chiffchaff, across year | Perch | −0.051 (−0.261 to 0.147) | −0.066 (−0.216 to 0.066) | −0.018 (−0.068 to 0.039) |
| Chiffchaff, across year | SurfPerch | 0.048 (−0.022 to 0.136) | 0.058 (−0.007 to 0.137) | 0.008 (−0.023 to 0.044) |
| Chiffchaff, across year | BirdNET v3 (preview) | −0.073 (−0.211 to 0.065) | −0.037 (−0.105 to 0.036) | −0.046 (−0.091 to −0.001) |
| Chiffchaff, across year | AVEX sl-BEATs | 0.005 (−0.085 to 0.085) | 0.029 (−0.047 to 0.140) | 0.041 (0.003 to 0.083) |
| Chiffchaff, across year | AVEX EfficientNet-B0 | 0.000 (−0.101 to 0.094) | −0.032 (−0.103 to 0.048) | 0.006 (−0.033 to 0.049) |
| Chiffchaff, across year | AvesEcho | −0.032 (−0.272 to 0.149) | 0.041 (−0.027 to 0.116) | 0.014 (−0.009 to 0.039) |
| Chiffchaff, across year | AudioProtoPNet | −0.075 (−0.214 to 0.074) | −0.065 (−0.161 to 0.033) | 0.002 (−0.035 to 0.042) |
| Chiffchaff, across year | ConvNeXt (BirdSet) | −0.021 (−0.166 to 0.157) | −0.010 (−0.088 to 0.093) | 0.016 (−0.017 to 0.053) |
| Chiffchaff, across year | Bird-MAE | 0.006 (−0.037 to 0.061) | −0.010 (−0.065 to 0.052) | −0.015 (−0.048 to 0.010) |
| Chiffchaff, across year | ProtoCLR | −0.030 (−0.124 to 0.069) | −0.042 (−0.145 to 0.066) | −0.054 (−0.131 to 0.011) |
| Chiffchaff, across year | RCL_FS_BSED | 0.045 (−0.068 to 0.188) | 0.005 (−0.082 to 0.106) | −0.023 (−0.056 to 0.000) |
| Chiffchaff, across year | BirdAVES | 0.036 (−0.071 to 0.207) | −0.012 (−0.086 to 0.055) | −0.003 (−0.058 to 0.041) |
| Chiffchaff, across year | AVES | 0.040 (−0.110 to 0.237) | 0.025 (−0.029 to 0.086) | −0.007 (−0.079 to 0.063) |
| Chiffchaff, across year | NatureBEATs | −0.040 (−0.115 to 0.030) | 0.005 (−0.009 to 0.021) | −0.001 (−0.025 to 0.024) |
| Chiffchaff, across year | BioLingual | −0.051 (−0.092 to −0.017) | −0.015 (−0.063 to 0.046) | −0.027 (−0.067 to 0.013) |
| Chiffchaff, across year | BEATs | 0.042 (−0.097 to 0.168) | 0.037 (−0.013 to 0.101) | 0.014 (−0.047 to 0.066) |
| Chiffchaff, across year | AudioMAE | −0.023 (−0.050 to 0.003) | 0.003 (−0.021 to 0.028) | 0.032 (0.005 to 0.066) |
| Chiffchaff, across year | VGGish | 0.081 (−0.025 to 0.263) | 0.021 (−0.026 to 0.080) | 0.009 (0.000 to 0.027) |
| Chiffchaff, across year | ECAPA-TDNN (speaker) | 0.053 (−0.062 to 0.181) | 0.068 (0.009 to 0.157) | 0.025 (−0.017 to 0.072) |
| Chiffchaff, across year | ResNet (speaker) | −0.021 (−0.167 to 0.133) | −0.056 (−0.138 to 0.012) | −0.007 (−0.058 to 0.038) |
| Chiffchaff, across year | x-vector (speaker) | 0.009 (−0.079 to 0.129) | −0.023 (−0.094 to 0.060) | −0.054 (−0.101 to −0.014) |
| Chiffchaff, across year | wav2vec 2.0 Base | 0.041 (−0.094 to 0.232) | 0.003 (−0.084 to 0.094) | −0.004 (−0.025 to 0.016) |
| Chiffchaff, across year | wav2vec 2.0 Large (robust) | −0.050 (−0.245 to 0.158) | −0.034 (−0.151 to 0.077) | −0.024 (−0.050 to −0.001) |
| Chiffchaff, across year | XLS-R 300M | −0.044 (−0.281 to 0.191) | −0.062 (−0.207 to 0.064) | −0.033 (−0.084 to 0.013) |
| Chiffchaff, across year | wav2vec 2.0 Conformer Large | −0.082 (−0.310 to 0.141) | −0.038 (−0.164 to 0.108) | −0.019 (−0.072 to 0.037) |
| Chiffchaff, across year | MMS 300M | −0.006 (−0.226 to 0.213) | −0.030 (−0.175 to 0.101) | −0.005 (−0.043 to 0.029) |
| Chiffchaff, across year | HuBERT Base | −0.094 (−0.258 to 0.055) | −0.041 (−0.113 to 0.018) | −0.005 (−0.016 to 0.000) |
| Chiffchaff, across year | HuBERT Large | −0.011 (−0.242 to 0.237) | −0.033 (−0.127 to 0.060) | 0.002 (−0.032 to 0.037) |
| Chiffchaff, across year | data2vec Base (100 h) | 0.011 (−0.164 to 0.203) | −0.005 (−0.156 to 0.145) | 0.002 (−0.033 to 0.035) |
| Chiffchaff, across year | data2vec Base (960 h) | −0.010 (−0.180 to 0.174) | −0.032 (−0.177 to 0.108) | −0.004 (−0.047 to 0.034) |
| Chiffchaff, across year | WavLM Base+ | −0.096 (−0.257 to 0.051) | −0.039 (−0.109 to 0.015) | 0.000 (0.000 to 0.000) |
| Chiffchaff, across year | WavLM Large | −0.066 (−0.329 to 0.175) | −0.084 (−0.230 to 0.052) | −0.046 (−0.105 to 0.002) |

| Dataset | Embedding | Background +10 dB | Background 0 dB | Background −10 dB |
| --- | --- | --- | --- | --- |
| Chiffchaff, across year | UniSpeech-SAT Base+ | −0.078 (−0.268 to 0.104) | −0.043 (−0.120 to 0.023) | −0.010 (−0.025 to 0.000) |
| Chiffchaff, across year | XEUS | −0.020 (−0.321 to 0.273) | −0.016 (−0.210 to 0.168) | −0.033 (−0.122 to 0.050) |
| Chiffchaff, across year | Loudness control | −0.026 (−0.069 to 0.000) | −0.007 (−0.021 to 0.000) | 0.000 (0.000 to 0.000) |
| Chiffchaff, across year | Duration control | 0.000 (0.000 to 0.000) | 0.000 (0.000 to 0.000) | 0.000 (0.000 to 0.000) |

*Notes:* Background level relative to the test clip. Positive values mean the classifier's answers follow the background; negative values mean a clip mixed with its own individual's test-year background was named less accurately than one mixed with another individual's. Interval over individuals, 10,000 draws.

**Table S7.** Exploratory analysis, specified after the results were seen: when BirdNET named the wrong individual for a clip mixed with another individual's background, the share of those errors that named the donor.

| Dataset | Added background relative to test clip | Wrong answers | Named the donor, no background added | Named the donor, background added | Under reassigned donors | p |
|---|---|---|---|---|---|---|
| Chiffchaff, within year | +10 dB | 886 | 1.9% | 24.9% | 7.7% | 0.0001 |
| Chiffchaff, within year | 0 dB | 679 | 1.9% | 20.2% | 8.0% | 0.0001 |
| Chiffchaff, within year | −10 dB | 410 | 1.9% | 11.7% | 6.7% | 0.0001 |
| Little owl, across year | +10 dB | 202 | 2.0% | 1.0% | 6.3% | 1.0000 |
| Little owl, across year | 0 dB | 183 | 2.0% | 1.1% | 6.8% | 1.0000 |
| Little owl, across year | −10 dB | 208 | 2.0% | 1.4% | 5.6% | 0.9996 |
| Tree pipit, within year | +10 dB | 246 | 12.2% | 13.0% | 9.5% | 0.0402 |
| Tree pipit, within year | 0 dB | 215 | 12.2% | 15.8% | 9.4% | 0.0011 |
| Tree pipit, within year | −10 dB | 200 | 12.2% | 12.5% | 9.8% | 0.1146 |
| Tree pipit, across year | +10 dB | 245 | 14.2% | 8.2% | 11.0% | 0.9487 |
| Tree pipit, across year | 0 dB | 239 | 14.2% | 15.5% | 10.7% | 0.0114 |
| Tree pipit, across year | −10 dB | 243 | 14.2% | 16.0% | 11.0% | 0.0083 |
| Chiffchaff, across year | +10 dB | 168 | 3.6% | 12.5% | 10.5% | 0.2116 |
| Chiffchaff, across year | 0 dB | 158 | 3.6% | 10.1% | 11.2% | 0.7212 |
| Chiffchaff, across year | −10 dB | 159 | 3.6% | 2.5% | 10.4% | 1.0000 |

*Notes:* Each individual's donor is the next individual in sorted order, so the donor is fixed for all of an individual's clips; a hash chooses only which of the donor's background recordings is used. No background added: the same share among the errors on the unmixed test clips, which shows how often BirdNET already confused each individual with its donor. Reassigned donors: 9,999 random reassignments of donors among the wrong answers, never giving a clip its own individual; p is the share at or above the observed value. This comparison does not control for the fixed pairing, so the column without background is the one to compare with.

**Table S8.** Planted-signal test: calls of ten within-year chiffchaffs mixed into the across-year chiffchaff backgrounds, one bird per territory, scored with BirdNET.

| Mixture | Re-ID accuracy (95% interval) | p | Uniform chance | Majority class |
|---|---|---|---|---|
| Call 10 dB above background | 0.641 (0.562 to 0.825) | 0.0001 | 0.100 | 0.436 |
| Equal level | 0.362 (0.231 to 0.646) | 0.0003 | 0.100 | 0.436 |
| Call 10 dB below background | 0.171 (0.055 to 0.448) | 0.1208 | 0.100 | 0.436 |

*Notes:* Interval over individuals, 10,000 draws; p from 9,999 label shuffles.

**Table S9.** Background augmentation and spectral subtraction with BirdNET, scored through the four pairings.

| Dataset | Remedy | Arm | C→C | B→B | C→B | B→C |
|---|---|---|---|---|---|---|
| Chiffchaff, within year | Background augmentation | Original | 0.765 (0.683 to 0.846); 0.0001 | 0.737 (0.597 to 0.810); 0.0001 | 0.456 (0.343 to 0.627); 0.0001 | 0.508 (0.278 to 0.654); 0.0001 |
| Chiffchaff, within year | Background augmentation | Augmented | 0.814 (0.770 to 0.890); 0.0001 | 0.737 (0.597 to 0.810); 0.0001 | 0.260 (0.148 to 0.420); 0.0001 | 0.508 (0.278 to 0.654); 0.0001 |
| Little owl, across year | Background augmentation | Original | 0.501 (0.376 to 0.630); 0.0001 | 0.105 (0.028 to 0.202); 0.0832 | 0.083 (0.009 to 0.199); 0.2334 | 0.076 (0.005 to 0.204); 0.3273 |
| Little owl, across year | Background augmentation | Augmented | 0.560 (0.406 to 0.713); 0.0001 | 0.105 (0.028 to 0.202); 0.0832 | 0.059 (0.000 to 0.174); 0.5312 | 0.076 (0.005 to 0.204); 0.3273 |
| Tree pipit, within year | Background augmentation | Original | 0.376 (0.192 to 0.561); 0.0001 | 0.177 (0.106 to 0.245); 0.0407 | 0.147 (0.044 to 0.259); 0.1326 | 0.106 (0.023 to 0.221); 0.4514 |
| Tree pipit, within year | Background augmentation | Augmented | 0.587 (0.397 to 0.750); 0.0001 | 0.177 (0.106 to 0.245); 0.0407 | 0.085 (0.004 to 0.253); 0.6807 | 0.106 (0.023 to 0.221); 0.4514 |
| Chiffchaff, within year | Spectral subtraction | Original | 0.765 (0.681 to 0.846); 0.0001 | 0.737 (0.599 to 0.813); 0.0001 | 0.456 (0.344 to 0.625); 0.0001 | 0.508 (0.281 to 0.651); 0.0001 |
| Chiffchaff, within year | Spectral subtraction | Subtracted once | 0.721 (0.657 to 0.838); 0.0001 | 0.681 (0.606 to 0.753); 0.0001 | 0.438 (0.342 to 0.595); 0.0001 | 0.267 (0.184 to 0.344); 0.0194 |
| Chiffchaff, within year | Spectral subtraction | Subtracted twice | 0.736 (0.658 to 0.829); 0.0001 | 0.708 (0.599 to 0.780); 0.0001 | 0.470 (0.362 to 0.646); 0.0001 | 0.271 (0.151 to 0.345); 0.0197 |
| Little owl, across year | Spectral subtraction | Original | 0.501 (0.371 to 0.634); 0.0001 | 0.105 (0.027 to 0.203); 0.0860 | 0.083 (0.009 to 0.198); 0.2263 | 0.076 (0.004 to 0.205); 0.3198 |
| Little owl, across year | Spectral subtraction | Subtracted once | 0.545 (0.415 to 0.680); 0.0001 | 0.083 (0.021 to 0.179); 0.1875 | 0.022 (0.004 to 0.044); 0.9820 | 0.052 (0.000 to 0.154); 0.6380 |
| Little owl, across year | Spectral subtraction | Subtracted twice | 0.523 (0.390 to 0.664); 0.0001 | 0.061 (0.016 to 0.121); 0.4950 | 0.100 (0.017 to 0.206); 0.0596 | 0.106 (0.005 to 0.277); 0.0893 |
| Tree pipit, within year | Spectral subtraction | Original | 0.376 (0.193 to 0.566); 0.0001 | 0.177 (0.106 to 0.246); 0.0408 | 0.147 (0.046 to 0.257); 0.1293 | 0.106 (0.023 to 0.219); 0.4515 |
| Tree pipit, within year | Spectral subtraction | Subtracted once | 0.485 (0.308 to 0.670); 0.0001 | 0.225 (0.084 to 0.432); 0.0027 | 0.167 (0.007 to 0.406); 0.0401 | 0.096 (0.003 to 0.277); 0.5803 |
| Tree pipit, within year | Spectral subtraction | Subtracted twice | 0.502 (0.316 to 0.692); 0.0001 | 0.181 (0.074 to 0.313); 0.0200 | 0.154 (0.007 to 0.383); 0.0660 | 0.092 (0.006 to 0.267); 0.6352 |
| Tree pipit, across year | Spectral subtraction | Original | 0.214 (0.116 to 0.334); 0.0022 | 0.065 (0.029 to 0.102); 0.8142 | 0.075 (0.015 to 0.145); 0.7089 | 0.089 (0.020 to 0.184); 0.6125 |
| Tree pipit, across year | Spectral subtraction | Subtracted once | 0.358 (0.219 to 0.520); 0.0001 | 0.098 (0.054 to 0.151); 0.4966 | 0.108 (0.000 to 0.278); 0.3893 | 0.099 (0.004 to 0.298); 0.5082 |
| Tree pipit, across year | Spectral subtraction | Subtracted twice | 0.323 (0.194 to 0.473); 0.0001 | 0.075 (0.039 to 0.110); 0.7739 | 0.101 (0.000 to 0.264); 0.4536 | 0.102 (0.000 to 0.298); 0.4620 |
| Chiffchaff, across year | Spectral subtraction | Original | 0.160 (0.035 to 0.333); 0.1327 | 0.132 (0.036 to 0.277); 0.2439 | 0.112 (0.005 to 0.318); 0.4019 | 0.110 (0.019 to 0.240); 0.4114 |
| Chiffchaff, across year | Spectral subtraction | Subtracted once | 0.205 (0.050 to 0.395); 0.0202 | 0.168 (0.070 to 0.296); 0.0727 | 0.203 (0.018 to 0.448); 0.0168 | 0.115 (0.010 to 0.268); 0.3568 |
| Chiffchaff, across year | Spectral subtraction | Subtracted twice | 0.215 (0.055 to 0.422); 0.0074 | 0.137 (0.033 to 0.289); 0.1817 | 0.168 (0.000 to 0.403); 0.0607 | 0.120 (0.000 to 0.323); 0.2935 |

*Notes:* Each cell: re-ID accuracy (95% interval over individuals, 10,000 draws); permutation p (9,999 shuffles). Background augmentation changes only the enrolment calls, so the two pairings fitted on backgrounds are unchanged. Spectral subtraction removes each individual's mean background spectrum, once or twice, from all its clips. The original arm was scored with its own draws of individuals, so its interval bounds can differ slightly from those in Table 4.

**Table S10.** Adaptive score normalisation with the class-mean classifier: re-ID accuracy before and after, for all 36 neural networks.

| Dataset | Embedding | Calls tested | Backgrounds tested |
|---|---|---|---|
| Chiffchaff, within year | BirdNET v2.4 | 0.549 to 0.386 | 0.265 to 0.192 |
| Chiffchaff, within year | Perch 2.0 | 0.542 to 0.463 | 0.269 to 0.311 |
| Chiffchaff, within year | Perch | 0.448 to 0.324 | 0.237 to 0.315 |
| Chiffchaff, within year | SurfPerch | 0.321 to 0.398 | 0.106 to 0.125 |
| Chiffchaff, within year | BirdNET v3 (preview) | 0.407 to 0.445 | 0.304 to 0.395 |
| Chiffchaff, within year | AVEX sl-BEATs | 0.375 to 0.317 | 0.162 to 0.181 |
| Chiffchaff, within year | AVEX EfficientNet-B0 | 0.383 to 0.378 | 0.108 to 0.222 |
| Chiffchaff, within year | AvesEcho | 0.374 to 0.299 | 0.176 to 0.167 |
| Chiffchaff, within year | AudioProtoPNet | 0.489 to 0.274 | 0.244 to 0.339 |
| Chiffchaff, within year | ConvNeXt (BirdSet) | 0.494 to 0.270 | 0.258 to 0.240 |
| Chiffchaff, within year | Bird-MAE | 0.148 to 0.133 | 0.109 to 0.134 |
| Chiffchaff, within year | ProtoCLR | 0.094 to 0.148 | 0.068 to 0.089 |
| Chiffchaff, within year | RCL_FS_BSED | 0.131 to 0.139 | 0.166 to 0.161 |
| Chiffchaff, within year | BirdAVES | 0.199 to 0.164 | 0.152 to 0.181 |
| Chiffchaff, within year | AVES | 0.224 to 0.163 | 0.198 to 0.220 |
| Chiffchaff, within year | NatureBEATs | 0.408 to 0.341 | 0.236 to 0.268 |
| Chiffchaff, within year | BioLingual | 0.256 to 0.378 | 0.080 to 0.121 |
| Chiffchaff, within year | BEATs | 0.182 to 0.133 | 0.281 to 0.177 |
| Chiffchaff, within year | AudioMAE | 0.065 to 0.135 | 0.068 to 0.099 |
| Chiffchaff, within year | VGGish | 0.202 to 0.322 | 0.070 to 0.106 |
| Chiffchaff, within year | ECAPA-TDNN (speaker) | 0.270 to 0.224 | 0.119 to 0.136 |
| Chiffchaff, within year | ResNet (speaker) | 0.385 to 0.307 | 0.217 to 0.205 |
| Chiffchaff, within year | x-vector (speaker) | 0.292 to 0.342 | 0.116 to 0.186 |
| Chiffchaff, within year | wav2vec 2.0 Base | 0.134 to 0.144 | 0.082 to 0.123 |
| Chiffchaff, within year | wav2vec 2.0 Large (robust) | 0.163 to 0.148 | 0.100 to 0.112 |
| Chiffchaff, within year | XLS-R 300M | 0.156 to 0.138 | 0.096 to 0.115 |
| Chiffchaff, within year | wav2vec 2.0 Conformer Large | 0.220 to 0.210 | 0.145 to 0.175 |
| Chiffchaff, within year | MMS 300M | 0.178 to 0.141 | 0.115 to 0.125 |
| Chiffchaff, within year | HuBERT Base | 0.151 to 0.153 | 0.085 to 0.135 |
| Chiffchaff, within year | HuBERT Large | 0.180 to 0.159 | 0.110 to 0.146 |
| Chiffchaff, within year | data2vec Base (100 h) | 0.149 to 0.164 | 0.113 to 0.203 |
| Chiffchaff, within year | data2vec Base (960 h) | 0.183 to 0.160 | 0.195 to 0.183 |
| Chiffchaff, within year | WavLM Base+ | 0.159 to 0.134 | 0.095 to 0.151 |
| Chiffchaff, within year | WavLM Large | 0.194 to 0.169 | 0.112 to 0.151 |
| Chiffchaff, within year | UniSpeech-SAT Base+ | 0.161 to 0.139 | 0.091 to 0.157 |
| Chiffchaff, within year | XEUS | 0.210 to 0.166 | 0.186 to 0.211 |
| Little owl, across year | BirdNET v2.4 | 0.393 to 0.386 | 0.071 to 0.066 |
| Little owl, across year | Perch 2.0 | 0.526 to 0.423 | 0.073 to 0.105 |
| Little owl, across year | Perch | 0.577 to 0.501 | 0.125 to 0.115 |
| Little owl, across year | SurfPerch | 0.359 to 0.337 | 0.120 to 0.115 |
| Little owl, across year | BirdNET v3 (preview) | 0.140 to 0.143 | 0.078 to 0.049 |
| Little owl, across year | AVEX sl-BEATs | 0.182 to 0.189 | 0.012 to 0.020 |
| Little owl, across year | AVEX EfficientNet-B0 | 0.263 to 0.248 | 0.117 to 0.112 |
| Little owl, across year | AvesEcho | 0.143 to 0.177 | 0.090 to 0.088 |
| Little owl, across year | AudioProtoPNet | 0.437 to 0.297 | 0.088 to 0.081 |

| Dataset | Embedding | Calls tested | Backgrounds tested |
|---|---|---|---|
| Little owl, across year | ConvNeXt (BirdSet) | 0.479 to 0.396 | 0.100 to 0.100 |
| Little owl, across year | Bird-MAE | 0.256 to 0.312 | 0.068 to 0.044 |
| Little owl, across year | ProtoCLR | 0.263 to 0.297 | 0.093 to 0.105 |
| Little owl, across year | RCL_FS_BSED | 0.172 to 0.123 | 0.088 to 0.088 |
| Little owl, across year | BirdAVES | 0.219 to 0.214 | 0.046 to 0.046 |
| Little owl, across year | AVES | 0.241 to 0.270 | 0.051 to 0.054 |
| Little owl, across year | NatureBEATs | 0.243 to 0.265 | 0.049 to 0.032 |
| Little owl, across year | BioLingual | 0.123 to 0.219 | 0.066 to 0.059 |
| Little owl, across year | BEATs | 0.204 to 0.162 | 0.064 to 0.034 |
| Little owl, across year | AudioMAE | 0.209 to 0.258 | 0.066 to 0.061 |
| Little owl, across year | VGGish | 0.197 to 0.145 | 0.083 to 0.061 |
| Little owl, across year | ECAPA-TDNN (speaker) | 0.287 to 0.283 | 0.117 to 0.117 |
| Little owl, across year | ResNet (speaker) | 0.354 to 0.280 | 0.051 to 0.054 |
| Little owl, across year | x-vector (speaker) | 0.243 to 0.248 | 0.054 to 0.064 |
| Little owl, across year | wav2vec 2.0 Base | 0.160 to 0.120 | 0.051 to 0.044 |
| Little owl, across year | wav2vec 2.0 Large (robust) | 0.133 to 0.155 | 0.056 to 0.064 |
| Little owl, across year | XLS-R 300M | 0.147 to 0.231 | 0.059 to 0.066 |
| Little owl, across year | wav2vec 2.0 Conformer Large | 0.199 to 0.233 | 0.051 to 0.054 |
| Little owl, across year | MMS 300M | 0.160 to 0.243 | 0.054 to 0.071 |
| Little owl, across year | HuBERT Base | 0.165 to 0.155 | 0.051 to 0.051 |
| Little owl, across year | HuBERT Large | 0.214 to 0.194 | 0.076 to 0.081 |
| Little owl, across year | data2vec Base (100 h) | 0.236 to 0.251 | 0.051 to 0.078 |
| Little owl, across year | data2vec Base (960 h) | 0.265 to 0.263 | 0.051 to 0.071 |
| Little owl, across year | WavLM Base+ | 0.147 to 0.138 | 0.051 to 0.051 |
| Little owl, across year | WavLM Large | 0.167 to 0.201 | 0.051 to 0.061 |
| Little owl, across year | UniSpeech-SAT Base+ | 0.162 to 0.162 | 0.051 to 0.054 |
| Little owl, across year | XEUS | 0.204 to 0.197 | 0.051 to 0.051 |
| Tree pipit, within year | BirdNET v2.4 | 0.327 to 0.317 | 0.184 to 0.137 |
| Tree pipit, within year | Perch 2.0 | 0.469 to 0.436 | 0.116 to 0.181 |
| Tree pipit, within year | Perch | 0.300 to 0.294 | 0.184 to 0.208 |
| Tree pipit, within year | SurfPerch | 0.330 to 0.287 | 0.133 to 0.181 |
| Tree pipit, within year | BirdNET v3 (preview) | 0.218 to 0.132 | 0.218 to 0.287 |
| Tree pipit, within year | AVEX sl-BEATs | 0.125 to 0.125 | 0.075 to 0.065 |
| Tree pipit, within year | AVEX EfficientNet-B0 | 0.244 to 0.254 | 0.123 to 0.143 |
| Tree pipit, within year | AvesEcho | 0.089 to 0.089 | 0.075 to 0.075 |
| Tree pipit, within year | AudioProtoPNet | 0.337 to 0.310 | 0.225 to 0.300 |
| Tree pipit, within year | ConvNeXt (BirdSet) | 0.360 to 0.376 | 0.177 to 0.229 |
| Tree pipit, within year | Bird-MAE | 0.116 to 0.132 | 0.082 to 0.102 |
| Tree pipit, within year | ProtoCLR | 0.135 to 0.139 | 0.119 to 0.102 |
| Tree pipit, within year | RCL_FS_BSED | 0.106 to 0.116 | 0.140 to 0.140 |
| Tree pipit, within year | BirdAVES | 0.221 to 0.241 | 0.184 to 0.184 |
| Tree pipit, within year | AVES | 0.248 to 0.185 | 0.218 to 0.167 |
| Tree pipit, within year | NatureBEATs | 0.122 to 0.129 | 0.065 to 0.072 |
| Tree pipit, within year | BioLingual | 0.069 to 0.066 | 0.126 to 0.130 |
| Tree pipit, within year | BEATs | 0.149 to 0.185 | 0.092 to 0.082 |
| Tree pipit, within year | AudioMAE | 0.099 to 0.083 | 0.137 to 0.130 |
| Tree pipit, within year | VGGish | 0.089 to 0.122 | 0.061 to 0.051 |

| Dataset | Embedding | Calls tested | Backgrounds tested |
|---|---|---|---|
| Tree pipit, within year | ECAPA-TDNN (speaker) | 0.300 to 0.307 | 0.191 to 0.208 |
| Tree pipit, within year | ResNet (speaker) | 0.201 to 0.201 | 0.143 to 0.133 |
| Tree pipit, within year | x-vector (speaker) | 0.281 to 0.267 | 0.137 to 0.157 |
| Tree pipit, within year | wav2vec 2.0 Base | 0.191 to 0.158 | 0.130 to 0.137 |
| Tree pipit, within year | wav2vec 2.0 Large (robust) | 0.158 to 0.178 | 0.113 to 0.171 |
| Tree pipit, within year | XLS-R 300M | 0.155 to 0.158 | 0.109 to 0.171 |
| Tree pipit, within year | wav2vec 2.0 Conformer Large | 0.155 to 0.149 | 0.137 to 0.205 |
| Tree pipit, within year | MMS 300M | 0.142 to 0.132 | 0.099 to 0.147 |
| Tree pipit, within year | HuBERT Base | 0.195 to 0.165 | 0.171 to 0.147 |
| Tree pipit, within year | HuBERT Large | 0.162 to 0.155 | 0.113 to 0.171 |
| Tree pipit, within year | data2vec Base (100 h) | 0.158 to 0.165 | 0.102 to 0.143 |
| Tree pipit, within year | data2vec Base (960 h) | 0.155 to 0.165 | 0.102 to 0.137 |
| Tree pipit, within year | WavLM Base+ | 0.208 to 0.152 | 0.164 to 0.140 |
| Tree pipit, within year | WavLM Large | 0.152 to 0.165 | 0.116 to 0.184 |
| Tree pipit, within year | UniSpeech-SAT Base+ | 0.172 to 0.149 | 0.150 to 0.137 |
| Tree pipit, within year | XEUS | 0.175 to 0.155 | 0.106 to 0.171 |
| Tree pipit, across year | BirdNET v2.4 | 0.173 to 0.214 | 0.056 to 0.078 |
| Tree pipit, across year | Perch 2.0 | 0.345 to 0.281 | 0.075 to 0.124 |
| Tree pipit, across year | Perch | 0.195 to 0.243 | 0.062 to 0.114 |
| Tree pipit, across year | SurfPerch | 0.166 to 0.134 | 0.065 to 0.042 |
| Tree pipit, across year | BirdNET v3 (preview) | 0.141 to 0.112 | 0.176 to 0.199 |
| Tree pipit, across year | AVEX sl-BEATs | 0.121 to 0.086 | 0.114 to 0.121 |
| Tree pipit, across year | AVEX EfficientNet-B0 | 0.153 to 0.163 | 0.111 to 0.114 |
| Tree pipit, across year | AvesEcho | 0.035 to 0.070 | 0.098 to 0.085 |
| Tree pipit, across year | AudioProtoPNet | 0.137 to 0.125 | 0.078 to 0.147 |
| Tree pipit, across year | ConvNeXt (BirdSet) | 0.214 to 0.224 | 0.029 to 0.088 |
| Tree pipit, across year | Bird-MAE | 0.058 to 0.080 | 0.098 to 0.085 |
| Tree pipit, across year | ProtoCLR | 0.093 to 0.112 | 0.111 to 0.092 |
| Tree pipit, across year | RCL_FS_BSED | 0.061 to 0.093 | 0.016 to 0.007 |
| Tree pipit, across year | BirdAVES | 0.054 to 0.125 | 0.039 to 0.013 |
| Tree pipit, across year | AVES | 0.038 to 0.112 | 0.013 to 0.029 |
| Tree pipit, across year | NatureBEATs | 0.125 to 0.099 | 0.121 to 0.127 |
| Tree pipit, across year | BioLingual | 0.054 to 0.077 | 0.101 to 0.105 |
| Tree pipit, across year | BEATs | 0.128 to 0.083 | 0.137 to 0.095 |
| Tree pipit, across year | AudioMAE | 0.051 to 0.058 | 0.078 to 0.075 |
| Tree pipit, across year | VGGish | 0.102 to 0.099 | 0.023 to 0.056 |
| Tree pipit, across year | ECAPA-TDNN (speaker) | 0.195 to 0.208 | 0.108 to 0.114 |
| Tree pipit, across year | ResNet (speaker) | 0.157 to 0.141 | 0.105 to 0.118 |
| Tree pipit, across year | x-vector (speaker) | 0.163 to 0.166 | 0.118 to 0.124 |
| Tree pipit, across year | wav2vec 2.0 Base | 0.070 to 0.064 | 0.134 to 0.114 |
| Tree pipit, across year | wav2vec 2.0 Large (robust) | 0.073 to 0.064 | 0.121 to 0.114 |
| Tree pipit, across year | XLS-R 300M | 0.058 to 0.026 | 0.124 to 0.108 |
| Tree pipit, across year | wav2vec 2.0 Conformer Large | 0.048 to 0.029 | 0.108 to 0.085 |
| Tree pipit, across year | MMS 300M | 0.061 to 0.038 | 0.121 to 0.114 |
| Tree pipit, across year | HuBERT Base | 0.045 to 0.045 | 0.059 to 0.033 |
| Tree pipit, across year | HuBERT Large | 0.070 to 0.048 | 0.127 to 0.108 |
| Tree pipit, across year | data2vec Base (100 h) | 0.077 to 0.054 | 0.065 to 0.056 |

| Dataset | Embedding | Calls tested | Backgrounds tested |
|---|---|---|---|
| Tree pipit, across year | data2vec Base (960 h) | 0.070 to 0.045 | 0.069 to 0.059 |
| Tree pipit, across year | WavLM Base+ | 0.035 to 0.045 | 0.042 to 0.013 |
| Tree pipit, across year | WavLM Large | 0.067 to 0.038 | 0.124 to 0.098 |
| Tree pipit, across year | UniSpeech-SAT Base+ | 0.032 to 0.045 | 0.056 to 0.026 |
| Tree pipit, across year | XEUS | 0.048 to 0.054 | 0.059 to 0.042 |
| Chiffchaff, across year | BirdNET v2.4 | 0.125 to 0.150 | 0.096 to 0.142 |
| Chiffchaff, across year | Perch 2.0 | 0.115 to 0.105 | 0.132 to 0.117 |
| Chiffchaff, across year | Perch | 0.105 to 0.115 | 0.152 to 0.152 |
| Chiffchaff, across year | SurfPerch | 0.120 to 0.165 | 0.127 to 0.168 |
| Chiffchaff, across year | BirdNET v3 (preview) | 0.065 to 0.065 | 0.091 to 0.107 |
| Chiffchaff, across year | AVEX sl-BEATs | 0.150 to 0.120 | 0.137 to 0.132 |
| Chiffchaff, across year | AVEX EfficientNet-B0 | 0.075 to 0.065 | 0.142 to 0.162 |
| Chiffchaff, across year | AvesEcho | 0.135 to 0.150 | 0.162 to 0.203 |
| Chiffchaff, across year | AudioProtoPNet | 0.085 to 0.095 | 0.157 to 0.152 |
| Chiffchaff, across year | ConvNeXt (BirdSet) | 0.105 to 0.120 | 0.117 to 0.137 |
| Chiffchaff, across year | Bird-MAE | 0.115 to 0.135 | 0.081 to 0.112 |
| Chiffchaff, across year | ProtoCLR | 0.185 to 0.140 | 0.112 to 0.112 |
| Chiffchaff, across year | RCL_FS_BSED | 0.135 to 0.060 | 0.081 to 0.112 |
| Chiffchaff, across year | BirdAVES | 0.150 to 0.105 | 0.183 to 0.178 |
| Chiffchaff, across year | AVES | 0.175 to 0.165 | 0.193 to 0.203 |
| Chiffchaff, across year | NatureBEATs | 0.120 to 0.140 | 0.102 to 0.132 |
| Chiffchaff, across year | BioLingual | 0.090 to 0.125 | 0.117 to 0.117 |
| Chiffchaff, across year | BEATs | 0.155 to 0.170 | 0.096 to 0.117 |
| Chiffchaff, across year | AudioMAE | 0.095 to 0.110 | 0.102 to 0.096 |
| Chiffchaff, across year | VGGish | 0.120 to 0.080 | 0.091 to 0.086 |
| Chiffchaff, across year | ECAPA-TDNN (speaker) | 0.105 to 0.085 | 0.152 to 0.132 |
| Chiffchaff, across year | ResNet (speaker) | 0.140 to 0.175 | 0.066 to 0.102 |
| Chiffchaff, across year | x-vector (speaker) | 0.100 to 0.095 | 0.112 to 0.102 |
| Chiffchaff, across year | wav2vec 2.0 Base | 0.180 to 0.105 | 0.168 to 0.168 |
| Chiffchaff, across year | wav2vec 2.0 Large (robust) | 0.160 to 0.175 | 0.137 to 0.122 |
| Chiffchaff, across year | XLS-R 300M | 0.155 to 0.170 | 0.132 to 0.122 |
| Chiffchaff, across year | wav2vec 2.0 Conformer Large | 0.165 to 0.155 | 0.127 to 0.122 |
| Chiffchaff, across year | MMS 300M | 0.165 to 0.170 | 0.137 to 0.117 |
| Chiffchaff, across year | HuBERT Base | 0.150 to 0.140 | 0.198 to 0.188 |
| Chiffchaff, across year | HuBERT Large | 0.170 to 0.170 | 0.137 to 0.132 |
| Chiffchaff, across year | data2vec Base (100 h) | 0.185 to 0.160 | 0.137 to 0.152 |
| Chiffchaff, across year | data2vec Base (960 h) | 0.185 to 0.155 | 0.112 to 0.147 |
| Chiffchaff, across year | WavLM Base+ | 0.140 to 0.130 | 0.193 to 0.188 |
| Chiffchaff, across year | WavLM Large | 0.150 to 0.170 | 0.076 to 0.132 |
| Chiffchaff, across year | UniSpeech-SAT Base+ | 0.140 to 0.130 | 0.198 to 0.193 |
| Chiffchaff, across year | XEUS | 0.160 to 0.180 | 0.122 to 0.132 |

*Notes:* Backgrounds tested: classifier fitted on backgrounds and tested on backgrounds. Intervals are in the numerical ledger. Duration and loudness controls are omitted because their unit-scaled values do not measure the intended magnitude controls; their kernel-ridge results elsewhere are retained.

**Table S11.** Session compensation on the fruit bat with BirdNET: re-ID accuracy and how well the recording day could still be predicted, for every setting specified in advance.

| Method | Setting | Re-ID accuracy | Day predicted | Day, majority rate |
|---|---|---|---|---|
| directions of between-session variation removed | 0 | 0.381 | 0.258 | 0.166 |
| directions of between-session variation removed | 1 | 0.370 | 0.261 | 0.166 |
| directions of between-session variation removed | 2 | 0.362 | 0.264 | 0.166 |
| directions of between-session variation removed | 4 | 0.369 | 0.261 | 0.166 |
| directions of between-session variation removed | 8 | 0.383 | 0.269 | 0.166 |
| directions of between-session variation removed | 16 | 0.331 | 0.261 | 0.166 |
| directions of between-session variation removed | 32 | 0.306 | 0.250 | 0.166 |
| none | n/a | 0.381 | 0.258 | 0.166 |
| session mean subtracted, whole session | n/a | 0.284 | 0.016 | 0.166 |
| session mean subtracted, running | n/a | 0.279 | 0.131 | 0.166 |
| within-class covariance normalisation | 0.0 | 0.281 | 0.265 | 0.166 |
| within-class covariance normalisation | 0.01 | 0.300 | 0.256 | 0.166 |
| within-class covariance normalisation | 0.03 | 0.312 | 0.252 | 0.166 |
| within-class covariance normalisation | 0.1 | 0.333 | 0.263 | 0.166 |
| within-class covariance normalisation | 0.3 | 0.352 | 0.269 | 0.166 |
| within-class covariance normalisation | 0.6 | 0.374 | 0.273 | 0.166 |
| within-class covariance normalisation | 0.9 | 0.378 | 0.278 | 0.166 |
| within-class covariance normalisation | 1.0 | 0.381 | 0.258 | 0.166 |

| Method | Setting | Classifier | Re-ID accuracy (95% interval) |
|---|---|---|---|
| none | n/a | kernel ridge | 0.501 (0.374 to 0.631) |
| none | n/a | class mean | 0.393 (0.244 to 0.553) |
| none | n/a | nearest clip | 0.548 (0.407 to 0.696) |
| within-class covariance normalisation | 0.0 | kernel ridge | 0.484 (0.360 to 0.609) |
| within-class covariance normalisation | 0.0 | class mean | 0.501 (0.378 to 0.632) |
| within-class covariance normalisation | 0.0 | nearest clip | 0.501 (0.381 to 0.629) |
| within-class covariance normalisation | 0.01 | kernel ridge | 0.516 (0.394 to 0.643) |
| within-class covariance normalisation | 0.01 | class mean | 0.526 (0.411 to 0.650) |
| within-class covariance normalisation | 0.01 | nearest clip | 0.514 (0.390 to 0.638) |
| within-class covariance normalisation | 0.03 | kernel ridge | 0.526 (0.409 to 0.656) |
| within-class covariance normalisation | 0.03 | class mean | 0.528 (0.406 to 0.648) |
| within-class covariance normalisation | 0.03 | nearest clip | 0.533 (0.414 to 0.654) |
| within-class covariance normalisation | 0.1 | kernel ridge | 0.541 (0.405 to 0.680) |
| within-class covariance normalisation | 0.1 | class mean | 0.533 (0.405 to 0.657) |
| within-class covariance normalisation | 0.1 | nearest clip | 0.555 (0.426 to 0.688) |
| within-class covariance normalisation | 0.3 | kernel ridge | 0.533 (0.389 to 0.672) |
| within-class covariance normalisation | 0.3 | class mean | 0.543 (0.410 to 0.672) |
| within-class covariance normalisation | 0.3 | nearest clip | 0.560 (0.426 to 0.685) |
| within-class covariance normalisation | 0.6 | kernel ridge | 0.526 (0.394 to 0.661) |
| within-class covariance normalisation | 0.6 | class mean | 0.521 (0.391 to 0.651) |
| within-class covariance normalisation | 0.6 | nearest clip | 0.565 (0.426 to 0.693) |
| within-class covariance normalisation | 0.9 | kernel ridge | 0.523 (0.379 to 0.658) |
| within-class covariance normalisation | 0.9 | class mean | 0.457 (0.324 to 0.590) |

| Method | Setting | Classifier | Re-ID accuracy (95% interval) |
|---|---|---|---|
| within-class covariance normalisation | 0.9 | nearest clip | 0.572 (0.436 to 0.707) |
| within-class covariance normalisation | 1.0 | kernel ridge | 0.501 (0.377 to 0.631) |
| within-class covariance normalisation | 1.0 | class mean | 0.393 (0.250 to 0.562) |
| within-class covariance normalisation | 1.0 | nearest clip | 0.548 (0.394 to 0.689) |

*Notes:* Upper: fruit bat. Day predicted: accuracy of a classifier fitted on all but one bat and tested on the one left out, for each bat in turn. Lower: within-class covariance normalisation on the little owl with three classifiers. Removing 0 directions and shrinkage 1.0 leave the embedding unchanged and repeat the row without compensation. Spearman correlation between re-ID accuracy and day prediction over the 16 distinct bat settings: 0.616 (p = 0.0111); without the two session-mean rows, 0.477 (p = 0.0843). Subtracting each day's mean over all bats places a bat's clips opposite the other bats' clips of the same day, so a day classifier fitted on the other bats is systematically wrong; the day prediction of 0.016 is a result of that design.

**Table S12.** The 35 neural networks compared with BirdNET v2.4.

| Neural network | Group | Source | Input sample rate (Hz) | Highest frequency analysed (kHz) | Window (s) | Candidate embeddings per dataset | Playback slowing |
|---|---|---|---|---|---|---|---|
| Perch 2.0 | animal or general sound | van Merriënboer et al. (2025) | 32,000 | 16 | 5 | 1 | 1 |
| Perch | animal or general sound | Ghani et al. (2023) | 32,000 | 16 | 5 | 1 | 1 |
| SurfPerch | animal or general sound | Williams et al. (2025) | 32,000 | 16 | 5 | 1 | 1 |
| BirdNET v3 (preview) | animal or general sound | developer preview, as supplied by Bacpipe 1.3.5 (Kather et al. 2026) | 32,000 | 16 | 3 | 1 | 1 |
| AVEX sl-BEATs | animal or general sound | Miron et al. (2026) | 16,000 | 8 | 5 | 13 | 1 |
| AVEX EfficientNet-B0 | animal or general sound | Miron et al. (2026) | 16,000 | 8 | 5 | 1 | 1 |
| AvesEcho | animal or general sound | Ghani et al. (2025) | 32,000 | 16 | 3 | 13 | 1 |
| AudioProtoPNet | animal or general sound | Heinrich et al. (2025) | 32,000 | 16 | 5 | 1 | 1 |
| ConvNeXt (BirdSet) | animal or general sound | Rauch et al. (2025b) | 32,000 | 16 | 5 | 1 | 1 |
| Bird-MAE | animal or general sound | Rauch et al. (2025a) | 32,000 | 16 | 5 | 33 | 1 |
| ProtoCLR | animal or general sound | Moummad et al. (2026) | 16,000 | 8 | 6 | 14 | 1 |
| RCL_FS_BSED | animal or general sound | Moummad et al. (2024) | 22,050 | 11.025 | 0.2 | 1 | 1 |
| BirdAVES | animal or general sound | Hagiwara (2023) | 16,000 | 8 | 1 | 25 | 1 |
| AVES | animal or general sound | Hagiwara (2023) | 16,000 | 8 | 1 | 13 | 1 |
| NatureBEATs | animal or general sound | Robinson et al. (2025) | 16,000 | 8 | 5 | 13 | 1 |
| BioLingual | animal or general sound | Robinson et al. (2024) | 48,000 | 24 | 10 | 13 | 1 |
| BEATs | animal or general sound | Chen et al. (2023) | 16,000 | 8 | 5 | 13 | 1 |
| AudioMAE | animal or general sound | Huang et al. (2022) | 16,000 | 8 | 10 | 13 | 1 |
| VGGish | animal or general sound | Hershey et al. (2017) | 16,000 | 8 | 1 | 1 | 1 |
| ECAPA-TDNN (speaker) | human speech | Desplanques et al. (2020); Ravanelli et al. (2021); speechbrain/spkrec-ecapa-voxceleb | 16,000 | 8; 16 and 24 at slowed playback | whole clip | 3 | 1, 2, 3 |
| ResNet (speaker) | human speech | Villalba et al. (2020); Ravanelli et al. (2021); speechbrain/spkrec-resnet-voxceleb | 16,000 | 8; 16 and 24 at slowed playback | whole clip | 3 | 1, 2, 3 |

| Neural network | Group | Source | Input sample rate (Hz) | Highest frequency analysed (kHz) | Window (s) | Candidate embeddings per dataset | Playback slowing |
|---|---|---|---|---|---|---|---|
| x-vector (speaker) | human speech | Snyder et al. (2018); Ravanelli et al. (2021); speechbrain/spkrec-xvect-voxceleb | 16,000 | 8; 16 and 24 at slowed playback | whole clip | 3 | 1, 2, 3 |
| wav2vec 2.0 Base | human speech | Baevski et al. (2020); facebook/wav2vec2-base | 16,000 | 8; 16 and 24 at slowed playback | whole clip | 39 | 1, 2, 3 |
| wav2vec 2.0 Large (robust) | human speech | Hsu et al. (2021b); facebook/wav2vec2-large-robust | 16,000 | 8; 16 and 24 at slowed playback | whole clip | 75 | 1, 2, 3 |
| XLS-R 300M | human speech | Babu et al. (2022); facebook/wav2vec2-xls-r-300m | 16,000 | 8; 16 and 24 at slowed playback | whole clip | 75 | 1, 2, 3 |
| wav2vec 2.0 Conformer Large | human speech | Gulati et al. (2020); Wang et al. (2020); facebook/wav2vec2-conformer-rope-large | 16,000 | 8; 16 and 24 at slowed playback | whole clip | 75 | 1, 2, 3 |
| MMS 300M | human speech | Pratap et al. (2024); facebook/mms-300m | 16,000 | 8; 16 and 24 at slowed playback | whole clip | 75 | 1, 2, 3 |
| HuBERT Base | human speech | Hsu et al. (2021a); facebook/hubert-base-ls960 | 16,000 | 8; 16 and 24 at slowed playback | whole clip | 39 | 1, 2, 3 |
| HuBERT Large | human speech | Hsu et al. (2021a); facebook/hubert-large-ll60k | 16,000 | 8; 16 and 24 at slowed playback | whole clip | 75 | 1, 2, 3 |
| data2vec Base (100 h) | human speech | Baevski et al. (2022); facebook/data2vec-audio-base-100h | 16,000 | 8; 16 and 24 at slowed playback | whole clip | 39 | 1, 2, 3 |
| data2vec Base (960 h) | human speech | Baevski et al. (2022); facebook/data2vec-audio-base-960h | 16,000 | 8; 16 and 24 at slowed playback | whole clip | 39 | 1, 2, 3 |
| WavLM Base+ | human speech | Chen et al. (2022a); microsoft/wavlm-base-plus | 16,000 | 8; 16 and 24 at slowed playback | whole clip | 39 | 1, 2, 3 |
| WavLM Large | human speech | Chen et al. (2022a); microsoft/wavlm-large | 16,000 | 8; 16 and 24 at slowed playback | whole clip | 75 | 1, 2, 3 |
| UniSpeech-SAT Base+ | human speech | Chen et al. (2022b); microsoft/unispeech-sat-base-plus | 16,000 | 8; 16 and 24 at slowed playback | whole clip | 39 | 1, 2, 3 |
| XEUS | human speech | Chen et al. (2024); espnet/xeus | 16,000 | 8; 16 and 24 at slowed playback | whole clip | 57 | 1, 2, 3 |

*Notes:* Speech networks are identified by their method papers and published checkpoint names. Window: the input length each clip was cut into; speech networks read whole clips, or 60-s windows where a clip exceeded the graphics card’s memory. Candidate embeddings: layers or blocks read, times playback speeds, where computed on that dataset. Playback slowing: 1, the recording’s own speed; 2 and 3, half and one third of it. Highest frequency analysed: half the input sample rate, the most a network can represent; for speech networks, 8 kHz at the recording’s own speed and 16 and 24 kHz at half and one third of it. BirdNET v2.4 reads 48 kHz audio. Sample rates and windows come from a record made by reloading each network. The speaker-verification checkpoint of WavLM Base+ (microsoft/wavlm-base-plus-sv) was also measured; the layers read have the weights of WavLM Base+ and every value was identical, so it is not counted separately. BirdNET v3 (preview) is the ONNX file Bacpipe 1.3.5 supplies (SHA-256 6f58d7ffa4c33bf49c8c67ac27bc5265a940a139cb67254477997bad41efc16d).

**Table S13.** Each neural network and control against BirdNET v2.4 on every dataset, under the two layer rules specified in advance.

*Zebra finch, group of four*: BirdNET v2.4 0.807 (0.669 to 0.963).

| Neural network | Fisher rule: layer, re-ID accuracy | Difference from BirdNET | Holm p | Held-out rule: layer, re-ID accuracy | Difference from BirdNET | Holm p |
|---|---|---|---|---|---|---|
| Perch 2.0 | embedding: 0.830 (0.679 to 0.923) | 0.023 (−0.037 to 0.073) | 1.0000 | embedding: 0.830 (0.679 to 0.923) | 0.023 (−0.037 to 0.073) | 1.0000 |
| Perch | embedding: 0.803 (0.717 to 0.838) | −0.005 (−0.125 to 0.079) | 1.0000 | embedding: 0.803 (0.717 to 0.838) | −0.005 (−0.125 to 0.079) | 1.0000 |
| SurfPerch | embedding: 0.780 (0.682 to 0.860) | −0.028 (−0.092 to 0.057) | 1.0000 | embedding: 0.780 (0.682 to 0.860) | −0.028 (−0.092 to 0.057) | 1.0000 |
| BirdNET v3 (preview) | embedding: 0.794 (0.696 to 0.900) | −0.014 (−0.062 to 0.027) | 1.0000 | embedding: 0.794 (0.696 to 0.900) | −0.014 (−0.062 to 0.027) | 1.0000 |
| AVEX sl-BEATs | block 0: 0.417 (0.058 to 0.675) | −0.390 (−0.904 to −0.035) | 0.8274 | block 5: 0.683 (0.604 to 0.776) | −0.124 (−0.183 to −0.027) | 0.8556 |
| AVEX EfficientNet-B0 | embedding: 0.798 (0.660 to 0.867) | −0.009 (−0.094 to 0.055) | 1.0000 | embedding: 0.798 (0.660 to 0.867) | −0.009 (−0.094 to 0.055) | 1.0000 |
| AvesEcho | block 3: 0.541 (0.236 to 0.921) | −0.266 (−0.421 to −0.042) | 0.2184 | embedding: 0.798 (0.702 to 0.942) | −0.009 (−0.036 to 0.075) | 1.0000 |
| AudioProtoPNet | embedding: 0.835 (0.679 to 0.950) | 0.028 (−0.013 to 0.064) | 1.0000 | embedding: 0.835 (0.679 to 0.950) | 0.028 (−0.013 to 0.064) | 1.0000 |
| ConvNeXt (BirdSet) | embedding: 0.839 (0.698 to 0.950) | 0.032 (−0.010 to 0.070) | 1.0000 | embedding: 0.839 (0.698 to 0.950) | 0.032 (−0.010 to 0.070) | 1.0000 |
| Bird-MAE | block 12: 0.367 (0.123 to 0.638) | −0.440 (−0.588 to −0.264) | < 0.0074 | block 18: 0.404 (0.202 to 0.625) | −0.404 (−0.509 to −0.264) | < 0.0074 |
| ProtoCLR | block 0: 0.372 (0.086 to 0.598) | −0.436 (−0.878 to −0.047) | 0.2184 | block 12: 0.610 (0.541 to 0.688) | −0.197 (−0.375 to −0.019) | 0.2494 |
| RCL_FS_BSED | embedding: 0.661 (0.491 to 0.835) | −0.147 (−0.273 to −0.027) | 0.8880 | embedding: 0.661 (0.491 to 0.835) | −0.147 (−0.273 to −0.027) | 0.9768 |
| BirdAVES | block 23: 0.638 (0.396 to 0.774) | −0.170 (−0.319 to −0.123) | < 0.0074 | block 9: 0.560 (0.270 to 0.801) | −0.248 (−0.360 to −0.125) | < 0.0074 |
| AVES | block 6: 0.445 (0.228 to 0.683) | −0.362 (−0.482 to −0.182) | < 0.0074 | block 5: 0.454 (0.211 to 0.721) | −0.353 (−0.500 to −0.160) | < 0.0074 |
| NatureBEATs | block 6: 0.734 (0.585 to 0.892) | −0.073 (−0.108 to −0.047) | 0.2184 | block 10: 0.784 (0.682 to 0.900) | −0.023 (−0.062 to 0.019) | 1.0000 |
| BioLingual | block 9: 0.417 (0.053 to 0.896) | −0.390 (−0.658 to −0.068) | < 0.0074 | block 4: 0.679 (0.482 to 0.954) | −0.128 (−0.245 to 0.000) | 1.0000 |
| BEATs | block 4: 0.495 (0.059 to 0.859) | −0.312 (−0.892 to 0.116) | 1.0000 | block 10: 0.734 (0.574 to 0.938) | −0.073 (−0.110 to −0.024) | < 0.0074 |
| AudioMAE | block 6: 0.509 (0.263 to 0.851) | −0.298 (−0.447 to −0.113) | < 0.0074 | block 11: 0.683 (0.482 to 0.958) | −0.124 (−0.251 to 0.023) | 1.0000 |
| VGGish | embedding: 0.702 (0.486 to 0.873) | −0.106 (−0.182 to −0.058) | < 0.0074 | embedding: 0.702 (0.486 to 0.873) | −0.106 (−0.182 to −0.058) | < 0.0074 |
| ECAPA-TDNN (speaker) | embedding, 1/3 speed: 0.711 (0.509 to 0.796) | −0.096 (−0.218 to −0.016) | 0.2184 | embedding, 1/3 speed: 0.711 (0.509 to 0.796) | −0.096 (−0.218 to −0.016) | 0.2494 |
| ResNet (speaker) | embedding, 1/2 speed: 0.771 (0.647 to 0.816) | −0.037 (−0.188 to 0.073) | 1.0000 | embedding: 0.794 (0.669 to 0.896) | −0.014 (−0.059 to 0.038) | 1.0000 |
| x-vector (speaker) | embedding, 1/3 speed: 0.789 (0.623 to 0.941) | −0.018 (−0.059 to −0.003) | 0.2184 | embedding, 1/3 speed: 0.789 (0.623 to 0.941) | −0.018 (−0.059 to −0.003) | 0.2494 |
| wav2vec 2.0 Base | layer 1, 1/2 speed: 0.826 (0.660 to 0.879) | 0.018 (−0.118 to 0.120) | 1.0000 | layer 4, 1/3 speed: 0.697 (0.585 to 0.733) | −0.110 (−0.254 to −0.009) | 0.2494 |
| wav2vec 2.0 Large (robust) | layer 21, 1/2 speed: 0.555 (0.272 to 0.865) | −0.252 (−0.439 to −0.017) | 0.2184 | layer 22, 1/2 speed: 0.394 (0.224 to 0.610) | −0.413 (−0.682 to −0.113) | 0.2494 |
| XLS-R 300M | layer 1, 1/3 speed: 0.789 (0.608 to 0.891) | −0.018 (−0.126 to 0.030) | 1.0000 | layer 2, full speed: 0.839 (0.736 to 0.958) | 0.032 (−0.004 to 0.075) | 1.0000 |
| wav2vec 2.0 Conformer Large | layer 1, 1/3 speed: 0.780 (0.601 to 0.875) | −0.028 (−0.129 to 0.040) | 1.0000 | layer 4, full speed: 0.780 (0.689 to 0.827) | −0.028 (−0.141 to 0.059) | 1.0000 |
| MMS 300M | layer 1, 1/3 speed: 0.798 (0.622 to 0.923) | −0.009 (−0.092 to 0.023) | 1.0000 | layer 3, 1/2 speed: 0.784 (0.615 to 0.959) | −0.023 (−0.053 to 0.016) | 1.0000 |
| HuBERT Base | layer 2, full speed: 0.638 (0.471 to 0.760) | −0.170 (−0.475 to 0.026) | 1.0000 | layer 1, full speed: 0.656 (0.506 to 0.766) | −0.151 (−0.425 to 0.028) | 1.0000 |
| HuBERT Large | layer 3, 1/3 speed: 0.716 (0.412 to 0.923) | −0.092 (−0.257 to 0.000) | 1.0000 | layer 11, 1/3 speed: 0.679 (0.351 to 0.923) | −0.128 (−0.299 to 0.010) | 1.0000 |
| data2vec Base (100 h) | layer 1, 1/2 speed: 0.743 (0.622 to 0.892) | −0.064 (−0.073 to −0.038) | 0.2184 | layer 1, full speed: 0.739 (0.628 to 0.808) | −0.069 (−0.153 to 0.000) | 1.0000 |

| Neural network | Fisher rule: layer, re-ID accuracy | Difference from BirdNET | Holm p | Held-out rule: layer, re-ID accuracy | Difference from BirdNET | Holm p |
|---|---|---|---|---|---|---|
| data2vec Base (960 h) | layer 1, 1/2 speed: 0.743 (0.581 to 0.882) | −0.064 (−0.118 to −0.045) | < 0.0074 | layer 1, full speed: 0.725 (0.574 to 0.846) | −0.083 (−0.118 to −0.038) | 0.2494 |
| WavLM Base+ | layer 0, 1/3 speed: 0.817 (0.681 to 0.852) | 0.009 (−0.141 to 0.105) | 1.0000 | layer 5, 1/2 speed: 0.716 (0.534 to 0.864) | −0.092 (−0.151 to −0.068) | < 0.0074 |
| WavLM Large | layer 23, 1/3 speed: 0.743 (0.472 to 0.921) | −0.064 (−0.208 to −0.018) | < 0.0074 | layer 19, full speed: 0.711 (0.585 to 0.829) | −0.096 (−0.135 to −0.057) | 0.2340 |
| UniSpeech-SAT Base+ | layer 0, 1/3 speed: 0.794 (0.630 to 0.837) | −0.014 (−0.165 to 0.083) | 1.0000 | layer 1, full speed: 0.720 (0.565 to 0.838) | −0.087 (−0.367 to 0.098) | 1.0000 |
| XEUS | layer 0, 1/2 speed: 0.817 (0.660 to 0.905) | 0.009 (−0.059 to 0.064) | 1.0000 | layer 11, 1/2 speed: 0.812 (0.642 to 0.955) | 0.005 (−0.050 to 0.025) | 1.0000 |
| Loudness control | summary: 0.381 (0.018 to 0.846) | −0.427 (−0.693 to −0.118) | < 0.0074 | summary: 0.381 (0.018 to 0.846) | −0.427 (−0.693 to −0.118) | < 0.0074 |
| Duration control | summary: 0.468 (0.000 to 0.850) | −0.339 (−0.906 to 0.107) | 1.0000 | summary: 0.468 (0.000 to 0.850) | −0.339 (−0.906 to 0.107) | 1.0000 |

*Zebra finch, one bird per recording*: BirdNET v2.4 0.967 (0.933 to 0.989).

| Neural network | Fisher rule: layer, re-ID accuracy | Difference from BirdNET | Holm p | Held-out rule: layer, re-ID accuracy | Difference from BirdNET | Holm p |
|---|---|---|---|---|---|---|
| Perch 2.0 | embedding: 0.956 (0.911 to 1.000) | −0.011 (−0.044 to 0.022) | 1.0000 | embedding: 0.956 (0.911 to 1.000) | −0.011 (−0.044 to 0.022) | 1.0000 |
| Perch | embedding: 0.878 (0.722 to 0.978) | −0.089 (−0.222 to 0.000) | 0.4256 | embedding: 0.878 (0.722 to 0.978) | −0.089 (−0.222 to 0.000) | 0.4820 |
| SurfPerch | embedding: 0.900 (0.833 to 0.967) | −0.067 (−0.111 to −0.022) | 0.0182 | embedding: 0.900 (0.833 to 0.967) | −0.067 (−0.111 to −0.022) | 0.0252 |
| BirdNET v3 (preview) | embedding: 0.844 (0.744 to 0.933) | −0.122 (−0.200 to −0.044) | 0.0240 | embedding: 0.844 (0.744 to 0.933) | −0.122 (−0.200 to −0.044) | 0.0336 |
| AVEX sl-BEATs | block 6: 0.733 (0.533 to 0.878) | −0.233 (−0.444 to −0.078) | < 0.0074 | block 11: 0.889 (0.811 to 0.967) | −0.078 (−0.156 to −0.022) | 0.4820 |
| AVEX EfficientNet-B0 | embedding: 0.878 (0.767 to 0.978) | −0.089 (−0.189 to 0.000) | 0.4416 | embedding: 0.878 (0.767 to 0.978) | −0.089 (−0.189 to 0.000) | 0.4820 |
| AvesEcho | block 4: 0.756 (0.567 to 0.911) | −0.211 (−0.389 to −0.056) | 0.0306 | embedding: 0.889 (0.756 to 0.978) | −0.078 (−0.189 to 0.000) | 0.4820 |
| AudioProtoPNet | embedding: 0.933 (0.844 to 1.000) | −0.033 (−0.111 to 0.022) | 1.0000 | embedding: 0.933 (0.844 to 1.000) | −0.033 (−0.111 to 0.022) | 1.0000 |
| ConvNeXt (BirdSet) | embedding: 0.978 (0.933 to 1.000) | 0.011 (−0.022 to 0.044) | 1.0000 | embedding: 0.978 (0.933 to 1.000) | 0.011 (−0.022 to 0.044) | 1.0000 |
| Bird-MAE | block 16: 0.567 (0.356 to 0.744) | −0.400 (−0.600 to −0.222) | < 0.0074 | block 24: 0.656 (0.444 to 0.833) | −0.311 (−0.511 to −0.133) | < 0.0074 |
| ProtoCLR | block 11: 0.378 (0.167 to 0.589) | −0.589 (−0.789 to −0.389) | < 0.0074 | block 1: 0.344 (0.144 to 0.578) | −0.622 (−0.811 to −0.411) | < 0.0074 |
| RCL_FS_BSED | embedding: 0.456 (0.189 to 0.722) | −0.511 (−0.756 to −0.267) | < 0.0074 | embedding: 0.456 (0.189 to 0.722) | −0.511 (−0.756 to −0.267) | < 0.0074 |
| BirdAVES | block 23: 0.511 (0.333 to 0.678) | −0.456 (−0.611 to −0.300) | < 0.0074 | block 22: 0.556 (0.367 to 0.744) | −0.411 (−0.578 to −0.244) | < 0.0074 |
| AVES | block 10: 0.556 (0.389 to 0.711) | −0.411 (−0.556 to −0.267) | < 0.0074 | block 10: 0.556 (0.389 to 0.711) | −0.411 (−0.556 to −0.267) | < 0.0074 |
| NatureBEATs | block 2: 0.600 (0.400 to 0.789) | −0.367 (−0.567 to −0.189) | < 0.0074 | block 7: 0.700 (0.578 to 0.811) | −0.267 (−0.389 to −0.167) | < 0.0074 |
| BioLingual | embedding: 0.956 (0.911 to 0.989) | −0.011 (−0.044 to 0.022) | 1.0000 | embedding: 0.956 (0.911 to 0.989) | −0.011 (−0.044 to 0.022) | 1.0000 |
| BEATs | block 4: 0.500 (0.278 to 0.722) | −0.467 (−0.689 to −0.244) | < 0.0074 | block 9: 0.767 (0.544 to 0.933) | −0.200 (−0.400 to −0.056) | 0.0050 |
| AudioMAE | block 10: 0.733 (0.578 to 0.867) | −0.233 (−0.400 to −0.089) | 0.0120 | block 9: 0.667 (0.511 to 0.800) | −0.300 (−0.456 to −0.167) | < 0.0074 |
| VGGish | embedding: 0.533 (0.333 to 0.733) | −0.433 (−0.622 to −0.244) | < 0.0074 | embedding: 0.533 (0.333 to 0.733) | −0.433 (−0.622 to −0.244) | < 0.0074 |
| ECAPA-TDNN (speaker) | embedding, 1/2 speed: 0.844 (0.744 to 0.933) | −0.122 (−0.222 to −0.033) | 0.0168 | embedding, 1/2 speed: 0.844 (0.744 to 0.933) | −0.122 (−0.222 to −0.033) | 0.0228 |
| ResNet (speaker) | embedding: 0.878 (0.756 to 0.967) | −0.089 (−0.200 to 0.000) | 0.4416 | embedding, 1/3 speed: 0.889 (0.822 to 0.944) | −0.078 (−0.133 to −0.033) | 0.0252 |
| x-vector (speaker) | embedding, 1/2 speed: 0.767 (0.611 to 0.911) | −0.200 (−0.333 to −0.078) | < 0.0074 | embedding, 1/2 speed: 0.767 (0.611 to 0.911) | −0.200 (−0.333 to −0.078) | < 0.0074 |
| wav2vec 2.0 Base | layer 1, 1/3 speed: 0.622 (0.389 to 0.844) | −0.344 (−0.544 to −0.144) | 0.0040 | layer 0, 1/3 speed: 0.722 (0.556 to 0.889) | −0.244 (−0.389 to −0.100) | 0.0050 |
| wav2vec 2.0 Large (robust) | layer 24, 1/2 speed: 0.256 (0.056 to 0.500) | −0.711 (−0.900 to −0.478) | < 0.0074 | layer 2, full speed: 0.700 (0.467 to 0.900) | −0.267 (−0.478 to −0.089) | < 0.0074 |

| Neural network | Fisher rule: layer, re-ID accuracy | Difference from BirdNET | Holm p | Held-out rule: layer, re-ID accuracy | Difference from BirdNET | Holm p |
|---|---|---|---|---|---|---|
| XLS-R 300M | layer 0, 1/3 speed: 0.756 (0.533 to 0.911) | −0.211 (−0.411 to −0.078) | < 0.0074 | layer 6, 1/3 speed: 0.767 (0.478 to 0.978) | −0.200 (−0.456 to −0.011) | 0.4820 |
| wav2vec 2.0 Conformer Large | layer 1, full speed: 0.700 (0.467 to 0.911) | −0.267 (−0.489 to −0.078) | 0.0072 | layer 5, full speed: 0.689 (0.456 to 0.889) | −0.278 (−0.478 to −0.100) | 0.0050 |
| MMS 300M | layer 0, full speed: 0.722 (0.489 to 0.911) | −0.244 (−0.456 to −0.067) | 0.0072 | layer 16, 1/3 speed: 0.767 (0.556 to 0.933) | −0.200 (−0.378 to −0.044) | 0.1274 |
| HuBERT Base | layer 0, full speed: 0.744 (0.511 to 0.933) | −0.222 (−0.433 to −0.056) | 0.0240 | layer 0, full speed: 0.744 (0.511 to 0.933) | −0.222 (−0.433 to −0.056) | 0.0320 |
| HuBERT Large | layer 2, full speed: 0.656 (0.411 to 0.867) | −0.311 (−0.533 to −0.122) | < 0.0074 | layer 0, full speed: 0.733 (0.500 to 0.922) | −0.233 (−0.444 to −0.067) | 0.0080 |
| data2vec Base (100 h) | layer 5, full speed: 0.478 (0.267 to 0.689) | −0.489 (−0.667 to −0.300) | < 0.0074 | layer 0, full speed: 0.744 (0.522 to 0.911) | −0.222 (−0.422 to −0.067) | 0.0050 |
| data2vec Base (960 h) | layer 10, 1/3 speed: 0.422 (0.233 to 0.611) | −0.544 (−0.722 to −0.367) | < 0.0074 | layer 0, full speed: 0.744 (0.522 to 0.911) | −0.222 (−0.422 to −0.067) | 0.0050 |
| WavLM Base+ | layer 0, full speed: 0.756 (0.533 to 0.933) | −0.211 (−0.411 to −0.056) | 0.0240 | layer 0, full speed: 0.756 (0.533 to 0.933) | −0.211 (−0.411 to −0.056) | 0.0320 |
| WavLM Large | layer 0, full speed: 0.711 (0.489 to 0.900) | −0.256 (−0.456 to −0.089) | 0.0072 | layer 5, full speed: 0.744 (0.467 to 0.956) | −0.222 (−0.467 to −0.022) | 0.1274 |
| UniSpeech-SAT Base+ | layer 2, 1/3 speed: 0.789 (0.689 to 0.900) | −0.178 (−0.256 to −0.089) | 0.0040 | layer 1, full speed: 0.767 (0.522 to 0.956) | −0.200 (−0.411 to −0.033) | 0.1274 |
| XEUS | layer 0, 1/3 speed: 0.867 (0.744 to 0.978) | −0.100 (−0.200 to 0.000) | 0.4256 | layer 6, full speed: 0.778 (0.511 to 0.978) | −0.189 (−0.422 to −0.011) | 0.4820 |
| Loudness control | summary: 0.233 (0.100 to 0.389) | −0.733 (−0.878 to −0.578) | < 0.0074 | summary: 0.233 (0.100 to 0.389) | −0.733 (−0.878 to −0.578) | < 0.0074 |
| Duration control | summary: 0.189 (0.000 to 0.456) | −0.778 (−0.967 to −0.533) | < 0.0074 | summary: 0.189 (0.000 to 0.456) | −0.778 (−0.967 to −0.533) | < 0.0074 |

*Zebra finch, group of eight*: BirdNET v2.4 0.498 (0.181 to 0.731).

| Neural network | Fisher rule: layer, re-ID accuracy | Difference from BirdNET | Holm p | Held-out rule: layer, re-ID accuracy | Difference from BirdNET | Holm p |
|---|---|---|---|---|---|---|
| Perch 2.0 | embedding: 0.559 (0.374 to 0.719) | 0.061 (−0.034 to 0.209) | 1.0000 | embedding: 0.559 (0.374 to 0.719) | 0.061 (−0.034 to 0.209) | 1.0000 |
| Perch | embedding: 0.528 (0.365 to 0.725) | 0.031 (−0.093 to 0.199) | 1.0000 | embedding: 0.528 (0.365 to 0.725) | 0.031 (−0.093 to 0.199) | 1.0000 |
| SurfPerch | embedding: 0.397 (0.171 to 0.689) | −0.100 (−0.213 to 0.016) | 1.0000 | embedding: 0.397 (0.171 to 0.689) | −0.100 (−0.213 to 0.016) | 1.0000 |
| BirdNET v3 (preview) | embedding: 0.515 (0.347 to 0.692) | 0.017 (−0.099 to 0.179) | 1.0000 | embedding: 0.515 (0.347 to 0.692) | 0.017 (−0.099 to 0.179) | 1.0000 |
| AVEX sl-BEATs | block 0: 0.201 (0.014 to 0.593) | −0.297 (−0.544 to −0.019) | 0.5890 | block 11: 0.437 (0.232 to 0.664) | −0.061 (−0.163 to 0.085) | 1.0000 |
| AVEX EfficientNet-B0 | embedding: 0.485 (0.234 to 0.705) | −0.013 (−0.070 to 0.067) | 1.0000 | embedding: 0.485 (0.234 to 0.705) | −0.013 (−0.070 to 0.067) | 1.0000 |
| AvesEcho | block 7: 0.297 (0.094 to 0.667) | −0.201 (−0.391 to −0.009) | 0.8064 | block 9: 0.301 (0.105 to 0.630) | −0.197 (−0.360 to −0.036) | < 0.0074 |
| AudioProtoPNet | embedding: 0.576 (0.397 to 0.748) | 0.079 (−0.037 to 0.228) | 1.0000 | embedding: 0.576 (0.397 to 0.748) | 0.079 (−0.037 to 0.228) | 1.0000 |
| ConvNeXt (BirdSet) | embedding: 0.502 (0.284 to 0.743) | 0.004 (−0.095 to 0.128) | 1.0000 | embedding: 0.502 (0.284 to 0.743) | 0.004 (−0.095 to 0.128) | 1.0000 |
| Bird-MAE | block 5: 0.188 (0.012 to 0.567) | −0.310 (−0.550 to −0.035) | 0.3740 | block 0: 0.188 (0.025 to 0.551) | −0.310 (−0.552 to −0.024) | 0.5016 |
| ProtoCLR | embedding: 0.118 (0.000 to 0.450) | −0.380 (−0.640 to −0.047) | 0.5056 | block 2: 0.183 (0.030 to 0.515) | −0.314 (−0.534 to −0.031) | 0.8384 |
| RCL_FS_BSED | embedding: 0.210 (0.013 to 0.627) | −0.288 (−0.539 to −0.004) | 1.0000 | embedding: 0.210 (0.013 to 0.627) | −0.288 (−0.539 to −0.004) | 1.0000 |
| BirdAVES | block 23: 0.245 (0.091 to 0.481) | −0.253 (−0.409 to −0.059) | < 0.0074 | block 13: 0.197 (0.092 to 0.381) | −0.301 (−0.494 to −0.043) | 0.8580 |
| AVES | block 11: 0.175 (0.061 to 0.406) | −0.323 (−0.550 to −0.038) | 0.4422 | block 4: 0.157 (0.052 to 0.361) | −0.341 (−0.566 to −0.044) | 0.3710 |
| NatureBEATs | block 0: 0.201 (0.014 to 0.593) | −0.297 (−0.544 to −0.019) | 0.5890 | block 8: 0.397 (0.241 to 0.654) | −0.100 (−0.313 to 0.168) | 1.0000 |
| BioLingual | block 5: 0.192 (0.022 to 0.554) | −0.306 (−0.540 to −0.036) | 0.1610 | block 11: 0.354 (0.130 to 0.680) | −0.144 (−0.314 to 0.107) | 1.0000 |
| BEATs | block 5: 0.197 (0.020 to 0.576) | −0.301 (−0.537 to −0.019) | 0.5890 | block 9: 0.485 (0.250 to 0.690) | −0.013 (−0.149 to 0.104) | 1.0000 |
| AudioMAE | block 7: 0.275 (0.077 to 0.681) | −0.223 (−0.470 to 0.068) | 1.0000 | block 5: 0.223 (0.045 to 0.580) | −0.275 (−0.518 to −0.017) | 0.4692 |

| Neural network | Fisher rule: layer, re-ID accuracy | Difference from BirdNET | Holm p | Held-out rule: layer, re-ID accuracy | Difference from BirdNET | Holm p |
|---|---|---|---|---|---|---|
| VGGish | embedding: 0.445 (0.114 to 0.681) | −0.052 (−0.182 to 0.010) | 1.0000 | embedding: 0.445 (0.114 to 0.681) | −0.052 (−0.182 to 0.010) | 1.0000 |
| ECAPA-TDNN (speaker) | embedding, 1/2 speed: 0.454 (0.323 to 0.620) | −0.044 (−0.184 to 0.157) | 1.0000 | embedding: 0.358 (0.222 to 0.535) | −0.140 (−0.291 to 0.076) | 1.0000 |
| ResNet (speaker) | embedding: 0.467 (0.314 to 0.720) | −0.031 (−0.180 to 0.155) | 1.0000 | embedding: 0.467 (0.314 to 0.720) | −0.031 (−0.180 to 0.155) | 1.0000 |
| x-vector (speaker) | embedding, 1/2 speed: 0.454 (0.193 to 0.644) | −0.044 (−0.227 to 0.083) | 1.0000 | embedding, 1/2 speed: 0.454 (0.193 to 0.644) | −0.044 (−0.227 to 0.083) | 1.0000 |
| wav2vec 2.0 Base | layer 0, 1/2 speed: 0.380 (0.230 to 0.669) | −0.118 (−0.326 to 0.116) | 1.0000 | layer 12, 1/3 speed: 0.262 (0.126 to 0.504) | −0.236 (−0.423 to 0.007) | 1.0000 |
| wav2vec 2.0 Large (robust) | layer 0, 1/2 speed: 0.310 (0.104 to 0.711) | −0.188 (−0.475 to 0.124) | 1.0000 | layer 21, 1/2 speed: 0.240 (0.059 to 0.586) | −0.258 (−0.535 to 0.048) | 1.0000 |
| XLS-R 300M | layer 0, 1/2 speed: 0.367 (0.159 to 0.763) | −0.131 (−0.427 to 0.210) | 1.0000 | layer 21, 1/3 speed: 0.380 (0.220 to 0.671) | −0.118 (−0.330 to 0.110) | 1.0000 |
| wav2vec 2.0 Conformer Large | layer 0, 1/3 speed: 0.406 (0.214 to 0.748) | −0.092 (−0.290 to 0.183) | 1.0000 | layer 0, 1/3 speed: 0.406 (0.214 to 0.748) | −0.092 (−0.290 to 0.183) | 1.0000 |
| MMS 300M | layer 0, 1/2 speed: 0.402 (0.205 to 0.780) | −0.096 (−0.373 to 0.222) | 1.0000 | layer 23, 1/2 speed: 0.223 (0.054 to 0.543) | −0.275 (−0.551 to 0.040) | 1.0000 |
| HuBERT Base | layer 0, 1/3 speed: 0.376 (0.202 to 0.698) | −0.122 (−0.318 to 0.124) | 1.0000 | layer 7, 1/2 speed: 0.271 (0.121 to 0.554) | −0.227 (−0.454 to 0.070) | 1.0000 |
| HuBERT Large | layer 18, 1/3 speed: 0.218 (0.038 to 0.562) | −0.279 (−0.552 to 0.054) | 1.0000 | layer 13, full speed: 0.306 (0.110 to 0.548) | −0.192 (−0.383 to 0.034) | 1.0000 |
| data2vec Base (100 h) | layer 3, full speed: 0.480 (0.215 to 0.658) | −0.017 (−0.245 to 0.164) | 1.0000 | layer 3, 1/2 speed: 0.432 (0.197 to 0.646) | −0.066 (−0.213 to 0.064) | 1.0000 |
| data2vec Base (960 h) | layer 4, 1/2 speed: 0.371 (0.068 to 0.628) | −0.127 (−0.260 to −0.045) | < 0.0074 | layer 12, 1/3 speed: 0.170 (0.034 to 0.474) | −0.328 (−0.549 to −0.040) | 0.8384 |
| WavLM Base+ | layer 0, 1/3 speed: 0.397 (0.237 to 0.683) | −0.100 (−0.285 to 0.117) | 1.0000 | layer 5, full speed: 0.384 (0.237 to 0.591) | −0.114 (−0.333 to 0.196) | 1.0000 |
| WavLM Large | layer 0, 1/3 speed: 0.314 (0.166 to 0.622) | −0.183 (−0.420 to 0.098) | 1.0000 | layer 13, full speed: 0.476 (0.237 to 0.683) | −0.022 (−0.198 to 0.153) | 1.0000 |
| UniSpeech-SAT Base+ | layer 0, 1/3 speed: 0.410 (0.247 to 0.702) | −0.087 (−0.271 to 0.124) | 1.0000 | layer 5, full speed: 0.476 (0.189 to 0.698) | −0.022 (−0.211 to 0.106) | 1.0000 |
| XEUS | layer 0, full speed: 0.445 (0.317 to 0.660) | −0.052 (−0.230 to 0.174) | 1.0000 | layer 18, 1/2 speed: 0.284 (0.091 to 0.549) | −0.214 (−0.341 to −0.065) | < 0.0074 |
| Loudness control | summary: 0.210 (0.014 to 0.608) | −0.288 (−0.565 to 0.079) | 1.0000 | summary: 0.210 (0.014 to 0.608) | −0.288 (−0.565 to 0.079) | 1.0000 |
| Duration control | summary: 0.284 (0.000 to 0.572) | −0.214 (−0.661 to 0.000) | 1.0000 | summary: 0.284 (0.000 to 0.572) | −0.214 (−0.661 to 0.000) | 1.0000 |

*Great tit, across year*: BirdNET v2.4 0.444 (0.319 to 0.569).

| Neural network | Fisher rule: layer, re-ID accuracy | Difference from BirdNET | Holm p | Held-out rule: layer, re-ID accuracy | Difference from BirdNET | Holm p |
|---|---|---|---|---|---|---|
| Perch 2.0 | embedding: 0.450 (0.312 to 0.588) | 0.006 (−0.094 to 0.100) | 0.9264 | embedding: 0.450 (0.312 to 0.588) | 0.006 (−0.094 to 0.100) | 1.0000 |
| Perch | embedding: 0.294 (0.194 to 0.400) | −0.150 (−0.256 to −0.056) | 0.0112 | embedding: 0.294 (0.194 to 0.400) | −0.150 (−0.256 to −0.056) | 0.0116 |
| SurfPerch | embedding: 0.375 (0.275 to 0.475) | −0.069 (−0.131 to −0.012) | 0.0968 | embedding: 0.375 (0.275 to 0.475) | −0.069 (−0.131 to −0.012) | 0.1210 |
| BirdNET v3 (preview) | embedding: 0.156 (0.106 to 0.206) | −0.287 (−0.437 to −0.144) | < 0.0074 | embedding: 0.156 (0.106 to 0.206) | −0.287 (−0.437 to −0.144) | < 0.0074 |
| AVEX sl-BEATs | block 6: 0.306 (0.200 to 0.412) | −0.137 (−0.244 to −0.037) | 0.0768 | block 9: 0.331 (0.219 to 0.450) | −0.112 (−0.206 to −0.019) | 0.1164 |
| AVEX EfficientNet-B0 | embedding: 0.338 (0.225 to 0.456) | −0.106 (−0.231 to 0.000) | 0.1188 | embedding: 0.338 (0.225 to 0.456) | −0.106 (−0.231 to 0.000) | 0.1782 |
| AvesEcho | block 5: 0.119 (0.050 to 0.206) | −0.325 (−0.450 to −0.181) | < 0.0074 | embedding: 0.438 (0.294 to 0.581) | −0.006 (−0.069 to 0.056) | 1.0000 |
| AudioProtoPNet | embedding: 0.294 (0.194 to 0.394) | −0.150 (−0.269 to −0.044) | 0.0616 | embedding: 0.294 (0.194 to 0.394) | −0.150 (−0.269 to −0.044) | 0.0616 |

| Neural network | Fisher rule: layer, re-ID accuracy | Difference from BirdNET | Holm p | Held-out rule: layer, re-ID accuracy | Difference from BirdNET | Holm p |
|---|---|---|---|---|---|---|
| ConvNeXt (BirdSet) | embedding: 0.325 (0.200 to 0.463) | −0.119 (−0.225 to −0.025) | 0.0768 | embedding: 0.325 (0.200 to 0.463) | −0.119 (−0.225 to −0.025) | 0.0884 |
| Bird-MAE | block 12: 0.138 (0.044 to 0.250) | −0.306 (−0.469 to −0.131) | 0.0192 | block 9: 0.125 (0.037 to 0.231) | −0.319 (−0.481 to −0.144) | 0.0116 |
| ProtoCLR | embedding: 0.113 (0.031 to 0.212) | −0.331 (−0.500 to −0.150) | 0.0192 | embedding: 0.113 (0.031 to 0.212) | −0.331 (−0.500 to −0.150) | 0.0192 |
| RCL_FS_BSED | embedding: 0.219 (0.094 to 0.362) | −0.225 (−0.338 to −0.106) | 0.0112 | embedding: 0.219 (0.094 to 0.362) | −0.225 (−0.338 to −0.106) | 0.0116 |
| BirdAVES | block 0: 0.287 (0.169 to 0.419) | −0.156 (−0.269 to −0.050) | 0.0540 | block 0: 0.287 (0.169 to 0.419) | −0.156 (−0.269 to −0.050) | 0.0540 |
| AVES | block 0: 0.225 (0.113 to 0.344) | −0.219 (−0.331 to −0.106) | 0.0060 | block 0: 0.225 (0.113 to 0.344) | −0.219 (−0.331 to −0.106) | 0.0062 |
| NatureBEATs | block 1: 0.200 (0.106 to 0.306) | −0.244 (−0.394 to −0.088) | 0.0294 | block 9: 0.244 (0.106 to 0.394) | −0.200 (−0.319 to −0.081) | 0.0342 |
| BioLingual | block 4: 0.188 (0.094 to 0.294) | −0.256 (−0.400 to −0.119) | 0.0060 | embedding: 0.319 (0.206 to 0.444) | −0.125 (−0.206 to −0.050) | 0.0220 |
| BEATs | block 9: 0.300 (0.200 to 0.400) | −0.144 (−0.250 to −0.044) | 0.0768 | block 8: 0.231 (0.125 to 0.344) | −0.212 (−0.344 to −0.081) | 0.0192 |
| AudioMAE | block 3: 0.100 (0.025 to 0.194) | −0.344 (−0.500 to −0.175) | < 0.0074 | block 11: 0.212 (0.131 to 0.294) | −0.231 (−0.350 to −0.125) | < 0.0074 |
| VGGish | embedding: 0.169 (0.087 to 0.250) | −0.275 (−0.394 to −0.150) | < 0.0074 | embedding: 0.169 (0.087 to 0.250) | −0.275 (−0.394 to −0.150) | < 0.0074 |
| ECAPA-TDNN (speaker) | embedding, 1/3 speed: 0.256 (0.150 to 0.375) | −0.188 (−0.331 to −0.069) | 0.0112 | embedding, 1/3 speed: 0.256 (0.150 to 0.375) | −0.188 (−0.331 to −0.069) | 0.0116 |
| ResNet (speaker) | embedding, 1/3 speed: 0.256 (0.175 to 0.344) | −0.188 (−0.312 to −0.069) | 0.0396 | embedding, 1/3 speed: 0.256 (0.175 to 0.344) | −0.188 (−0.312 to −0.069) | 0.0396 |
| x-vector (speaker) | embedding, 1/3 speed: 0.206 (0.106 to 0.319) | −0.237 (−0.344 to −0.131) | < 0.0074 | embedding, 1/2 speed: 0.194 (0.113 to 0.300) | −0.250 (−0.356 to −0.150) | < 0.0074 |
| wav2vec 2.0 Base | layer 0, full speed: 0.231 (0.113 to 0.369) | −0.212 (−0.344 to −0.081) | 0.0342 | layer 2, full speed: 0.231 (0.119 to 0.356) | −0.212 (−0.325 to −0.100) | 0.0062 |
| wav2vec 2.0 Large (robust) | layer 0, full speed: 0.219 (0.113 to 0.331) | −0.225 (−0.338 to −0.100) | 0.0294 | layer 0, full speed: 0.219 (0.113 to 0.331) | −0.225 (−0.338 to −0.100) | 0.0294 |
| XLS-R 300M | layer 0, 1/3 speed: 0.181 (0.087 to 0.287) | −0.262 (−0.394 to −0.119) | 0.0192 | layer 1, full speed: 0.231 (0.125 to 0.338) | −0.212 (−0.331 to −0.094) | 0.0150 |
| wav2vec 2.0 Conformer Large | layer 1, full speed: 0.237 (0.138 to 0.350) | −0.206 (−0.325 to −0.075) | 0.0448 | layer 1, full speed: 0.237 (0.138 to 0.350) | −0.206 (−0.325 to −0.075) | 0.0448 |
| MMS 300M | layer 0, 1/3 speed: 0.244 (0.144 to 0.350) | −0.200 (−0.319 to −0.062) | 0.0650 | layer 0, full speed: 0.269 (0.169 to 0.381) | −0.175 (−0.294 to −0.050) | 0.0884 |
| HuBERT Base | layer 0, full speed: 0.269 (0.144 to 0.406) | −0.175 (−0.306 to −0.044) | 0.0768 | layer 0, full speed: 0.269 (0.144 to 0.406) | −0.175 (−0.306 to −0.044) | 0.0884 |
| HuBERT Large | layer 0, full speed: 0.269 (0.163 to 0.381) | −0.175 (−0.294 to −0.044) | 0.0768 | layer 0, full speed: 0.269 (0.163 to 0.381) | −0.175 (−0.294 to −0.044) | 0.0884 |
| data2vec Base (100 h) | layer 0, full speed: 0.263 (0.156 to 0.381) | −0.181 (−0.306 to −0.050) | 0.0768 | layer 0, full speed: 0.263 (0.156 to 0.381) | −0.181 (−0.306 to −0.050) | 0.0884 |
| data2vec Base (960 h) | layer 0, full speed: 0.281 (0.175 to 0.400) | −0.162 (−0.300 to −0.019) | 0.0968 | layer 0, full speed: 0.281 (0.175 to 0.400) | −0.162 (−0.300 to −0.019) | 0.1210 |
| WavLM Base+ | layer 0, full speed: 0.250 (0.125 to 0.388) | −0.194 (−0.306 to −0.075) | 0.0396 | layer 0, full speed: 0.250 (0.125 to 0.388) | −0.194 (−0.306 to −0.075) | 0.0396 |
| WavLM Large | layer 0, full speed: 0.287 (0.181 to 0.394) | −0.156 (−0.269 to −0.037) | 0.0768 | layer 2, full speed: 0.250 (0.131 to 0.381) | −0.194 (−0.306 to −0.081) | 0.0294 |
| UniSpeech-SAT Base+ | layer 0, full speed: 0.281 (0.156 to 0.412) | −0.162 (−0.281 to −0.038) | 0.0768 | layer 0, full speed: 0.281 (0.156 to 0.412) | −0.162 (−0.281 to −0.038) | 0.0884 |
| XEUS | layer 0, full speed: 0.231 (0.119 to 0.350) | −0.212 (−0.325 to −0.100) | 0.0112 | layer 2, full speed: 0.275 (0.150 to 0.412) | −0.169 (−0.287 to −0.044) | 0.0884 |
| Loudness control | summary: 0.113 (0.031 to 0.206) | −0.331 (−0.494 to −0.169) | < 0.0074 | summary: 0.113 (0.031 to 0.206) | −0.331 (−0.494 to −0.169) | < 0.0074 |
| Duration control | summary: 0.069 (0.000 to 0.175) | −0.375 (−0.537 to −0.200) | < 0.0074 | summary: 0.069 (0.000 to 0.175) | −0.375 (−0.537 to −0.200) | < 0.0074 |

*Tree pipit, across year*: BirdNET v2.4 0.214 (0.117 to 0.336).

| Neural network | Fisher rule: layer, re-ID accuracy | Difference from BirdNET | Holm p | Held-out rule: layer, re-ID accuracy | Difference from BirdNET | Holm p |
|---|---|---|---|---|---|---|
| Perch 2.0 | embedding: 0.447 (0.294 to 0.625) | 0.233 (0.149 to 0.310) | < 0.0074 | n/a | n/a | n/a |
| Perch | embedding: 0.351 (0.219 to 0.517) | 0.137 (0.064 to 0.207) | 0.0204 | n/a | n/a | n/a |
| SurfPerch | embedding: 0.243 (0.153 to 0.358) | 0.029 (−0.015 to 0.083) | 1.0000 | n/a | n/a | n/a |
| BirdNET v3 (preview) | embedding: 0.160 (0.096 to 0.228) | −0.054 (−0.173 to 0.035) | 1.0000 | n/a | n/a | n/a |
| AVEX sl-BEATs | block 0: 0.096 (0.059 to 0.130) | −0.118 (−0.241 to −0.015) | 0.3144 | n/a | n/a | n/a |
| AVEX EfficientNet-B0 | embedding: 0.211 (0.114 to 0.319) | −0.003 (−0.091 to 0.091) | 1.0000 | n/a | n/a | n/a |
| AvesEcho | block 0: 0.019 (0.000 to 0.039) | −0.195 (−0.325 to −0.091) | < 0.0074 | n/a | n/a | n/a |
| AudioProtoPNet | embedding: 0.201 (0.102 to 0.317) | −0.013 (−0.070 to 0.036) | 1.0000 | n/a | n/a | n/a |
| ConvNeXt (BirdSet) | embedding: 0.288 (0.138 to 0.464) | 0.073 (−0.023 to 0.156) | 1.0000 | n/a | n/a | n/a |
| Bird-MAE | block 0: 0.032 (0.000 to 0.096) | −0.182 (−0.292 to −0.100) | < 0.0074 | n/a | n/a | n/a |
| ProtoCLR | embedding: 0.083 (0.030 to 0.156) | −0.131 (−0.261 to 0.000) | 0.4950 | n/a | n/a | n/a |
| RCL_FS_BSED | embedding: 0.045 (0.017 to 0.071) | −0.169 (−0.283 to −0.080) | < 0.0074 | n/a | n/a | n/a |
| BirdAVES | block 0: 0.083 (0.042 to 0.117) | −0.131 (−0.252 to −0.030) | 0.1230 | n/a | n/a | n/a |
| AVES | block 0: 0.038 (0.016 to 0.060) | −0.176 (−0.296 to −0.084) | < 0.0074 | n/a | n/a | n/a |
| NatureBEATs | block 0: 0.070 (0.037 to 0.099) | −0.144 (−0.272 to −0.038) | 0.0864 | n/a | n/a | n/a |
| BioLingual | block 0: 0.058 (0.000 to 0.133) | −0.157 (−0.278 to −0.028) | 0.2782 | n/a | n/a | n/a |
| BEATs | block 9: 0.086 (0.044 to 0.129) | −0.128 (−0.241 to −0.041) | < 0.0074 | n/a | n/a | n/a |
| AudioMAE | block 0: 0.045 (0.000 to 0.137) | −0.169 (−0.318 to −0.015) | 0.3916 | n/a | n/a | n/a |
| VGGish | embedding: 0.070 (0.034 to 0.109) | −0.144 (−0.249 to −0.062) | < 0.0074 | n/a | n/a | n/a |
| ECAPA-TDNN (speaker) | embedding, 1/3 speed: 0.214 (0.116 to 0.343) | 0.000 (−0.100 to 0.098) | 1.0000 | n/a | n/a | n/a |
| ResNet (speaker) | embedding, 1/3 speed: 0.137 (0.065 to 0.228) | −0.077 (−0.177 to −0.003) | 0.4200 | n/a | n/a | n/a |
| x-vector (speaker) | embedding, 1/3 speed: 0.195 (0.081 to 0.351) | −0.019 (−0.127 to 0.097) | 1.0000 | n/a | n/a | n/a |
| wav2vec 2.0 Base | layer 1, 1/3 speed: 0.064 (0.000 to 0.168) | −0.150 (−0.239 to −0.081) | < 0.0074 | n/a | n/a | n/a |
| wav2vec 2.0 Large (robust) | layer 0, 1/3 speed: 0.032 (0.006 to 0.062) | −0.182 (−0.306 to −0.089) | < 0.0074 | n/a | n/a | n/a |
| XLS-R 300M | layer 0, 1/3 speed: 0.022 (0.000 to 0.054) | −0.192 (−0.322 to −0.096) | < 0.0074 | n/a | n/a | n/a |
| wav2vec 2.0 Conformer Large | layer 0, 1/3 speed: 0.026 (0.000 to 0.057) | −0.188 (−0.318 to −0.092) | < 0.0074 | n/a | n/a | n/a |
| MMS 300M | layer 0, 1/3 speed: 0.019 (0.000 to 0.045) | −0.195 (−0.321 to −0.100) | < 0.0074 | n/a | n/a | n/a |
| HuBERT Base | layer 0, 1/3 speed: 0.013 (0.000 to 0.026) | −0.201 (−0.317 to −0.108) | < 0.0074 | n/a | n/a | n/a |
| HuBERT Large | layer 0, 1/3 speed: 0.032 (0.000 to 0.070) | −0.182 (−0.314 to −0.084) | < 0.0074 | n/a | n/a | n/a |
| data2vec Base (100 h) | layer 0, 1/3 speed: 0.029 (0.004 to 0.059) | −0.185 (−0.310 to −0.092) | < 0.0074 | n/a | n/a | n/a |
| data2vec Base (960 h) | layer 0, 1/3 speed: 0.029 (0.009 to 0.051) | −0.185 (−0.307 to −0.091) | < 0.0074 | n/a | n/a | n/a |
| WavLM Base+ | layer 0, 1/3 speed: 0.000 (0.000 to 0.000) | −0.214 (−0.336 to −0.117) | < 0.0074 | n/a | n/a | n/a |
| WavLM Large | layer 0, 1/3 speed: 0.026 (0.000 to 0.054) | −0.188 (−0.317 to −0.091) | < 0.0074 | n/a | n/a | n/a |

| Neural network | Fisher rule: layer, re-ID accuracy | Difference from BirdNET | Holm p | Held-out rule: layer, re-ID accuracy | Difference from BirdNET | Holm p |
|---|---|---|---|---|---|---|
| UniSpeech-SAT Base+ | layer 0, 1/3 speed: 0.016 (0.003 to 0.030) | −0.198 (−0.314 to −0.105) | < 0.0074 | n/a | n/a | n/a |
| XEUS | layer 0, 1/3 speed: 0.010 (0.000 to 0.019) | −0.204 (−0.323 to −0.109) | < 0.0074 | n/a | n/a | n/a |
| Loudness control | summary: 0.073 (0.018 to 0.138) | −0.141 (−0.261 to −0.033) | 0.1288 | n/a | n/a | n/a |
| Duration control | summary: 0.083 (0.000 to 0.234) | −0.131 (−0.307 to 0.079) | 1.0000 | n/a | n/a | n/a |

*Chiffchaff, across year*: BirdNET v2.4 0.160 (0.035 to 0.330).

| Neural network | Fisher rule: layer, re-ID accuracy | Difference from BirdNET | Holm p | Held-out rule: layer, re-ID accuracy | Difference from BirdNET | Holm p |
|---|---|---|---|---|---|---|
| Perch 2.0 | embedding: 0.160 (0.034 to 0.289) | 0.000 (−0.091 to 0.087) | 1.0000 | n/a | n/a | n/a |
| Perch | embedding: 0.150 (0.044 to 0.277) | −0.010 (−0.078 to 0.068) | 1.0000 | n/a | n/a | n/a |
| SurfPerch | embedding: 0.255 (0.103 to 0.421) | 0.095 (0.015 to 0.192) | 0.6808 | n/a | n/a | n/a |
| BirdNET v3 (preview) | embedding: 0.085 (0.033 to 0.151) | −0.075 (−0.245 to 0.077) | 1.0000 | n/a | n/a | n/a |
| AVEX sl-BEATs | block 9: 0.210 (0.074 to 0.361) | 0.050 (−0.026 to 0.119) | 1.0000 | n/a | n/a | n/a |
| AVEX EfficientNet-B0 | embedding: 0.120 (0.036 to 0.234) | −0.040 (−0.116 to 0.033) | 1.0000 | n/a | n/a | n/a |
| AvesEcho | block 6: 0.180 (0.012 to 0.382) | 0.020 (−0.058 to 0.097) | 1.0000 | n/a | n/a | n/a |
| AudioProtoPNet | embedding: 0.115 (0.047 to 0.197) | −0.045 (−0.155 to 0.054) | 1.0000 | n/a | n/a | n/a |
| ConvNeXt (BirdSet) | embedding: 0.135 (0.034 to 0.262) | −0.025 (−0.098 to 0.051) | 1.0000 | n/a | n/a | n/a |
| Bird-MAE | block 11: 0.080 (0.014 to 0.173) | −0.080 (−0.177 to 0.019) | 1.0000 | n/a | n/a | n/a |
| ProtoCLR | embedding: 0.195 (0.041 to 0.379) | 0.035 (−0.141 to 0.222) | 1.0000 | n/a | n/a | n/a |
| RCL_FS_BSED | embedding: 0.150 (0.016 to 0.312) | −0.010 (−0.083 to 0.064) | 1.0000 | n/a | n/a | n/a |
| BirdAVES | block 0: 0.190 (0.022 to 0.400) | 0.030 (−0.056 to 0.111) | 1.0000 | n/a | n/a | n/a |
| AVES | block 0: 0.205 (0.000 to 0.414) | 0.045 (−0.056 to 0.141) | 1.0000 | n/a | n/a | n/a |
| NatureBEATs | block 9: 0.160 (0.033 to 0.302) | 0.000 (−0.065 to 0.065) | 1.0000 | n/a | n/a | n/a |
| BioLingual | embedding: 0.160 (0.031 to 0.289) | 0.000 (−0.086 to 0.078) | 1.0000 | n/a | n/a | n/a |
| BEATs | block 9: 0.155 (0.043 to 0.287) | −0.005 (−0.070 to 0.053) | 1.0000 | n/a | n/a | n/a |
| AudioMAE | embedding: 0.150 (0.046 to 0.279) | −0.010 (−0.071 to 0.045) | 1.0000 | n/a | n/a | n/a |
| VGGish | embedding: 0.095 (0.000 to 0.275) | −0.065 (−0.126 to −0.005) | 1.0000 | n/a | n/a | n/a |
| ECAPA-TDNN (speaker) | embedding, 1/3 speed: 0.150 (0.054 to 0.275) | −0.010 (−0.090 to 0.075) | 1.0000 | n/a | n/a | n/a |
| ResNet (speaker) | embedding, 1/3 speed: 0.145 (0.046 to 0.289) | −0.015 (−0.166 to 0.140) | 1.0000 | n/a | n/a | n/a |
| x-vector (speaker) | embedding, 1/3 speed: 0.140 (0.033 to 0.313) | −0.020 (−0.084 to 0.040) | 1.0000 | n/a | n/a | n/a |
| wav2vec 2.0 Base | layer 1, full speed: 0.165 (0.000 to 0.380) | 0.005 (−0.077 to 0.093) | 1.0000 | n/a | n/a | n/a |
| wav2vec 2.0 Large (robust) | layer 0, full speed: 0.210 (0.017 to 0.426) | 0.050 (−0.045 to 0.168) | 1.0000 | n/a | n/a | n/a |
| XLS-R 300M | layer 0, full speed: 0.235 (0.057 to 0.439) | 0.075 (−0.022 to 0.176) | 1.0000 | n/a | n/a | n/a |
| wav2vec 2.0 Conformer Large | layer 0, full speed: 0.205 (0.000 to 0.389) | 0.045 (−0.067 to 0.161) | 1.0000 | n/a | n/a | n/a |
| MMS 300M | layer 0, full speed: 0.220 (0.034 to 0.433) | 0.060 (−0.027 to 0.156) | 1.0000 | n/a | n/a | n/a |

| Neural network | Fisher rule: layer, re-ID accuracy | Difference from BirdNET | Holm p | Held-out rule: layer, re-ID accuracy | Difference from BirdNET | Holm p |
|---|---|---|---|---|---|---|
| HuBERT Base | layer 0, full speed: 0.125 (0.000 to 0.344) | −0.035 (−0.104 to 0.036) | 1.0000 | n/a | n/a | n/a |
| HuBERT Large | layer 0, full speed: 0.215 (0.000 to 0.439) | 0.055 (−0.052 to 0.176) | 1.0000 | n/a | n/a | n/a |
| data2vec Base (100 h) | layer 0, full speed: 0.180 (0.000 to 0.392) | 0.020 (−0.067 to 0.123) | 1.0000 | n/a | n/a | n/a |
| data2vec Base (960 h) | layer 0, full speed: 0.180 (0.000 to 0.386) | 0.020 (−0.065 to 0.112) | 1.0000 | n/a | n/a | n/a |
| WavLM Base+ | layer 0, full speed: 0.125 (0.000 to 0.344) | −0.035 (−0.104 to 0.036) | 1.0000 | n/a | n/a | n/a |
| WavLM Large | layer 0, 1/2 speed: 0.200 (0.029 to 0.384) | 0.040 (−0.043 to 0.129) | 1.0000 | n/a | n/a | n/a |
| UniSpeech-SAT Base+ | layer 0, full speed: 0.130 (0.000 to 0.346) | −0.030 (−0.096 to 0.038) | 1.0000 | n/a | n/a | n/a |
| XEUS | layer 0, 1/2 speed: 0.205 (0.017 to 0.407) | 0.045 (−0.059 to 0.173) | 1.0000 | n/a | n/a | n/a |
| Loudness control | summary: 0.025 (0.000 to 0.058) | −0.135 (−0.308 to 0.000) | 1.0000 | n/a | n/a | n/a |
| Duration control | summary: 0.105 (0.000 to 0.298) | −0.055 (−0.271 to 0.156) | 1.0000 | n/a | n/a | n/a |

*Tree pipit, within year*: BirdNET v2.4 0.376 (0.195 to 0.564).

| Neural network | Fisher rule: layer, re-ID accuracy | Difference from BirdNET | Holm p | Held-out rule: layer, re-ID accuracy | Difference from BirdNET | Holm p |
|---|---|---|---|---|---|---|
| Perch 2.0 | embedding: 0.548 (0.361 to 0.738) | 0.172 (0.103 to 0.251) | < 0.0074 | n/a | n/a | n/a |
| Perch | embedding: 0.426 (0.230 to 0.628) | 0.050 (−0.054 to 0.171) | 1.0000 | n/a | n/a | n/a |
| SurfPerch | embedding: 0.455 (0.265 to 0.634) | 0.079 (−0.010 to 0.171) | 1.0000 | n/a | n/a | n/a |
| BirdNET v3 (preview) | embedding: 0.257 (0.128 to 0.417) | −0.119 (−0.304 to 0.054) | 1.0000 | n/a | n/a | n/a |
| AVEX sl-BEATs | block 0: 0.162 (0.056 to 0.286) | −0.215 (−0.377 to −0.048) | 0.3696 | n/a | n/a | n/a |
| AVEX EfficientNet-B0 | embedding: 0.281 (0.111 to 0.475) | −0.096 (−0.244 to 0.051) | 1.0000 | n/a | n/a | n/a |
| AvesEcho | block 0: 0.135 (0.015 to 0.329) | −0.241 (−0.469 to −0.025) | 0.8004 | n/a | n/a | n/a |
| AudioProtoPNet | embedding: 0.442 (0.239 to 0.657) | 0.066 (−0.087 to 0.237) | 1.0000 | n/a | n/a | n/a |
| ConvNeXt (BirdSet) | embedding: 0.502 (0.298 to 0.704) | 0.125 (0.012 to 0.265) | 0.8456 | n/a | n/a | n/a |
| Bird-MAE | block 0: 0.069 (0.020 to 0.134) | −0.307 (−0.511 to −0.108) | 0.0748 | n/a | n/a | n/a |
| ProtoCLR | embedding: 0.152 (0.025 to 0.308) | −0.224 (−0.448 to −0.012) | 1.0000 | n/a | n/a | n/a |
| RCL_FS_BSED | embedding: 0.172 (0.054 to 0.337) | −0.205 (−0.441 to 0.051) | 1.0000 | n/a | n/a | n/a |
| BirdAVES | block 0: 0.300 (0.105 to 0.515) | −0.076 (−0.269 to 0.157) | 1.0000 | n/a | n/a | n/a |
| AVES | block 0: 0.228 (0.065 to 0.429) | −0.149 (−0.343 to 0.100) | 1.0000 | n/a | n/a | n/a |
| NatureBEATs | block 0: 0.182 (0.085 to 0.281) | −0.195 (−0.350 to −0.028) | 0.6120 | n/a | n/a | n/a |
| BioLingual | block 0: 0.112 (0.014 to 0.228) | −0.264 (−0.427 to −0.116) | < 0.0074 | n/a | n/a | n/a |
| BEATs | block 9: 0.211 (0.096 to 0.329) | −0.165 (−0.308 to −0.033) | 0.4340 | n/a | n/a | n/a |
| AudioMAE | block 0: 0.125 (0.023 to 0.241) | −0.251 (−0.412 to −0.099) | < 0.0074 | n/a | n/a | n/a |
| VGGish | embedding: 0.116 (0.028 to 0.216) | −0.261 (−0.489 to −0.052) | 0.3776 | n/a | n/a | n/a |
| ECAPA-TDNN (speaker) | embedding, 1/3 speed: 0.356 (0.167 to 0.567) | −0.020 (−0.189 to 0.142) | 1.0000 | n/a | n/a | n/a |
| ResNet (speaker) | embedding, 1/3 speed: 0.264 (0.130 to 0.424) | −0.112 (−0.259 to 0.036) | 1.0000 | n/a | n/a | n/a |

| Neural network | Fisher rule: layer, re-ID accuracy | Difference from BirdNET | Holm p | Held-out rule: layer, re-ID accuracy | Difference from BirdNET | Holm p |
|---|---|---|---|---|---|---|
| x-vector (speaker) | embedding, 1/3 speed: 0.304 (0.117 to 0.517) | −0.073 (−0.243 to 0.106) | 1.0000 | n/a | n/a | n/a |
| wav2vec 2.0 Base | layer 1, 1/3 speed: 0.231 (0.061 to 0.426) | −0.145 (−0.326 to 0.084) | 1.0000 | n/a | n/a | n/a |
| wav2vec 2.0 Large (robust) | layer 0, 1/3 speed: 0.168 (0.006 to 0.400) | −0.208 (−0.474 to 0.090) | 1.0000 | n/a | n/a | n/a |
| XLS-R 300M | layer 0, 1/3 speed: 0.135 (0.015 to 0.329) | −0.241 (−0.477 to 0.043) | 1.0000 | n/a | n/a | n/a |
| wav2vec 2.0 Conformer Large | layer 0, 1/3 speed: 0.149 (0.021 to 0.346) | −0.228 (−0.466 to 0.058) | 1.0000 | n/a | n/a | n/a |
| MMS 300M | layer 0, 1/3 speed: 0.125 (0.016 to 0.316) | −0.251 (−0.486 to 0.036) | 1.0000 | n/a | n/a | n/a |
| HuBERT Base | layer 0, 1/3 speed: 0.191 (0.036 to 0.393) | −0.185 (−0.414 to 0.081) | 1.0000 | n/a | n/a | n/a |
| HuBERT Large | layer 0, 1/3 speed: 0.188 (0.016 to 0.429) | −0.188 (−0.457 to 0.099) | 1.0000 | n/a | n/a | n/a |
| data2vec Base (100 h) | layer 0, 1/3 speed: 0.182 (0.040 to 0.380) | −0.195 (−0.417 to 0.077) | 1.0000 | n/a | n/a | n/a |
| data2vec Base (960 h) | layer 0, 1/3 speed: 0.172 (0.041 to 0.360) | −0.205 (−0.426 to 0.062) | 1.0000 | n/a | n/a | n/a |
| WavLM Base+ | layer 0, 1/3 speed: 0.158 (0.025 to 0.355) | −0.218 (−0.445 to 0.059) | 1.0000 | n/a | n/a | n/a |
| WavLM Large | layer 0, 1/3 speed: 0.165 (0.009 to 0.386) | −0.211 (−0.472 to 0.070) | 1.0000 | n/a | n/a | n/a |
| UniSpeech-SAT Base+ | layer 0, 1/3 speed: 0.145 (0.018 to 0.329) | −0.231 (−0.452 to 0.033) | 1.0000 | n/a | n/a | n/a |
| XEUS | layer 0, 1/3 speed: 0.139 (0.011 to 0.333) | −0.238 (−0.481 to 0.047) | 1.0000 | n/a | n/a | n/a |
| Loudness control | summary: 0.228 (0.032 to 0.461) | −0.149 (−0.407 to 0.140) | 1.0000 | n/a | n/a | n/a |
| Duration control | summary: 0.145 (0.000 to 0.334) | −0.231 (−0.490 to 0.023) | 1.0000 | n/a | n/a | n/a |

*Little owl, across year*: BirdNET v2.4 0.501 (0.372 to 0.631).

| Neural network | Fisher rule: layer, re-ID accuracy | Difference from BirdNET | Holm p | Held-out rule: layer, re-ID accuracy | Difference from BirdNET | Holm p |
|---|---|---|---|---|---|---|
| Perch 2.0 | embedding: 0.651 (0.494 to 0.786) | 0.150 (0.044 to 0.244) | 0.1932 | n/a | n/a | n/a |
| Perch | embedding: 0.646 (0.501 to 0.775) | 0.145 (0.057 to 0.229) | 0.0432 | n/a | n/a | n/a |
| SurfPerch | embedding: 0.442 (0.291 to 0.593) | −0.059 (−0.218 to 0.097) | 1.0000 | n/a | n/a | n/a |
| BirdNET v3 (preview) | embedding: 0.373 (0.237 to 0.523) | −0.128 (−0.258 to 0.003) | 0.4944 | n/a | n/a | n/a |
| AVEX sl-BEATs | block 2: 0.194 (0.077 to 0.331) | −0.307 (−0.446 to −0.168) | < 0.0074 | n/a | n/a | n/a |
| AVEX EfficientNet-B0 | embedding: 0.373 (0.237 to 0.522) | −0.128 (−0.253 to −0.003) | 0.4944 | n/a | n/a | n/a |
| AvesEcho | block 0: 0.138 (0.029 to 0.266) | −0.364 (−0.532 to −0.181) | 0.0168 | n/a | n/a | n/a |
| AudioProtoPNet | embedding: 0.631 (0.472 to 0.782) | 0.130 (0.033 to 0.246) | 0.0920 | n/a | n/a | n/a |
| ConvNeXt (BirdSet) | embedding: 0.624 (0.479 to 0.754) | 0.123 (0.006 to 0.229) | 0.4944 | n/a | n/a | n/a |
| Bird-MAE | embedding: 0.251 (0.103 to 0.412) | −0.251 (−0.453 to −0.036) | 0.3424 | n/a | n/a | n/a |
| ProtoCLR | embedding: 0.172 (0.067 to 0.305) | −0.329 (−0.467 to −0.182) | < 0.0074 | n/a | n/a | n/a |
| RCL_FS_BSED | embedding: 0.189 (0.071 to 0.331) | −0.312 (−0.495 to −0.111) | 0.0864 | n/a | n/a | n/a |

| Neural network | Fisher rule: layer, re-ID accuracy | Difference from BirdNET | Holm p | Held-out rule: layer, re-ID accuracy | Difference from BirdNET | Holm p |
|---|---|---|---|---|---|---|
| BirdAVES | block 23: 0.285 (0.155 to 0.427) | −0.216 (−0.346 to −0.089) | 0.0120 | n/a | n/a | n/a |
| AVES | block 0: 0.285 (0.160 to 0.406) | −0.216 (−0.373 to −0.071) | 0.0600 | n/a | n/a | n/a |
| NatureBEATs | block 2: 0.241 (0.131 to 0.357) | −0.260 (−0.415 to −0.098) | 0.0468 | n/a | n/a | n/a |
| BioLingual | block 1: 0.133 (0.021 to 0.256) | −0.369 (−0.547 to −0.172) | 0.0120 | n/a | n/a | n/a |
| BEATs | block 2: 0.216 (0.087 to 0.369) | −0.285 (−0.447 to −0.091) | 0.1364 | n/a | n/a | n/a |
| AudioMAE | block 3: 0.170 (0.064 to 0.292) | −0.332 (−0.478 to −0.176) | < 0.0074 | n/a | n/a | n/a |
| VGGish | embedding: 0.231 (0.098 to 0.379) | −0.270 (−0.398 to −0.157) | < 0.0074 | n/a | n/a | n/a |
| ECAPA-TDNN (speaker) | embedding, 1/3 speed: 0.437 (0.279 to 0.595) | −0.064 (−0.255 to 0.123) | 1.0000 | n/a | n/a | n/a |
| ResNet (speaker) | embedding, 1/3 speed: 0.437 (0.280 to 0.584) | −0.064 (−0.251 to 0.115) | 1.0000 | n/a | n/a | n/a |
| x-vector (speaker) | embedding, 1/3 speed: 0.393 (0.252 to 0.532) | −0.108 (−0.299 to 0.085) | 1.0000 | n/a | n/a | n/a |
| wav2vec 2.0 Base | layer 0, full speed: 0.263 (0.104 to 0.442) | −0.238 (−0.412 to −0.050) | 0.2840 | n/a | n/a | n/a |
| wav2vec 2.0 Large (robust) | layer 0, full speed: 0.270 (0.132 to 0.419) | −0.231 (−0.400 to −0.049) | 0.2840 | n/a | n/a | n/a |
| XLS-R 300M | layer 0, full speed: 0.292 (0.136 to 0.457) | −0.209 (−0.400 to −0.002) | 0.4944 | n/a | n/a | n/a |
| wav2vec 2.0 Conformer Large | layer 1, full speed: 0.349 (0.201 to 0.507) | −0.152 (−0.290 to 0.009) | 0.4944 | n/a | n/a | n/a |
| MMS 300M | layer 0, full speed: 0.290 (0.137 to 0.461) | −0.211 (−0.395 to −0.012) | 0.4944 | n/a | n/a | n/a |
| HuBERT Base | layer 0, full speed: 0.290 (0.155 to 0.439) | −0.211 (−0.356 to −0.045) | 0.2840 | n/a | n/a | n/a |
| HuBERT Large | layer 1, full speed: 0.187 (0.089 to 0.305) | −0.314 (−0.460 to −0.178) | < 0.0074 | n/a | n/a | n/a |
| data2vec Base (100 h) | layer 0, full speed: 0.361 (0.217 to 0.518) | −0.140 (−0.281 to 0.018) | 0.4944 | n/a | n/a | n/a |
| data2vec Base (960 h) | layer 0, full speed: 0.378 (0.222 to 0.549) | −0.123 (−0.270 to 0.040) | 0.6800 | n/a | n/a | n/a |
| WavLM Base+ | layer 0, full speed: 0.305 (0.164 to 0.455) | −0.197 (−0.332 to −0.034) | 0.3424 | n/a | n/a | n/a |
| WavLM Large | layer 0, full speed: 0.268 (0.129 to 0.419) | −0.233 (−0.413 to −0.030) | 0.4116 | n/a | n/a | n/a |
| UniSpeech-SAT Base+ | layer 0, full speed: 0.270 (0.131 to 0.427) | −0.231 (−0.390 to −0.051) | 0.2840 | n/a | n/a | n/a |
| XEUS | layer 0, full speed: 0.297 (0.156 to 0.458) | −0.204 (−0.355 to −0.025) | 0.4116 | n/a | n/a | n/a |
| Loudness control | summary: 0.069 (0.014 to 0.134) | −0.432 (−0.570 to −0.305) | < 0.0074 | n/a | n/a | n/a |
| Duration control | summary: 0.115 (0.000 to 0.293) | −0.386 (−0.526 to −0.243) | < 0.0074 | n/a | n/a | n/a |

*Little penguin, across night*: BirdNET v2.4 0.693 (0.607 to 0.773).

| Neural network | Fisher rule: layer, re-ID accuracy | Difference from BirdNET | Holm p | Held-out rule: layer, re-ID accuracy | Difference from BirdNET | Holm p |
|---|---|---|---|---|---|---|
| Perch 2.0 | embedding: 0.705 (0.599 to 0.791) | 0.011 (−0.051 to 0.069) | 1.0000 | n/a | n/a | n/a |

| Neural network | Fisher rule: layer, re-ID accuracy | Difference from BirdNET | Holm p | Held-out rule: layer, re-ID accuracy | Difference from BirdNET | Holm p |
|---|---|---|---|---|---|---|
| Perch | embedding: 0.635 (0.528 to 0.727) | −0.058 (−0.134 to 0.013) | 1.0000 | n/a | n/a | n/a |
| SurfPerch | embedding: 0.646 (0.551 to 0.728) | −0.047 (−0.107 to 0.011) | 1.0000 | n/a | n/a | n/a |
| BirdNET v3 (preview) | embedding: 0.501 (0.371 to 0.611) | −0.192 (−0.268 to −0.124) | < 0.0074 | n/a | n/a | n/a |
| AVEX sl-BEATs | block 3: 0.518 (0.396 to 0.621) | −0.176 (−0.255 to −0.102) | < 0.0074 | n/a | n/a | n/a |
| AVEX EfficientNet-B0 | embedding: 0.569 (0.444 to 0.681) | −0.124 (−0.199 to −0.051) | 0.0040 | n/a | n/a | n/a |
| AvesEcho | block 1: 0.407 (0.271 to 0.520) | −0.287 (−0.390 to −0.188) | < 0.0074 | n/a | n/a | n/a |
| AudioProtoPNet | embedding: 0.674 (0.569 to 0.760) | −0.019 (−0.076 to 0.036) | 1.0000 | n/a | n/a | n/a |
| ConvNeXt (BirdSet) | embedding: 0.658 (0.538 to 0.755) | −0.035 (−0.130 to 0.052) | 1.0000 | n/a | n/a | n/a |
| Bird-MAE | block 18: 0.385 (0.222 to 0.523) | −0.308 (−0.434 to −0.196) | < 0.0074 | n/a | n/a | n/a |
| ProtoCLR | embedding: 0.447 (0.310 to 0.560) | −0.246 (−0.366 to −0.132) | < 0.0074 | n/a | n/a | n/a |
| RCL_FS_BSED | embedding: 0.622 (0.532 to 0.711) | −0.071 (−0.117 to −0.022) | 0.0936 | n/a | n/a | n/a |
| BirdAVES | block 0: 0.726 (0.631 to 0.818) | 0.033 (−0.023 to 0.097) | 1.0000 | n/a | n/a | n/a |
| AVES | block 0: 0.761 (0.673 to 0.845) | 0.068 (−0.002 to 0.149) | 0.8736 | n/a | n/a | n/a |
| NatureBEATs | block 1: 0.490 (0.373 to 0.593) | −0.203 (−0.290 to −0.120) | < 0.0074 | n/a | n/a | n/a |
| BioLingual | block 2: 0.343 (0.184 to 0.489) | −0.350 (−0.490 to −0.225) | < 0.0074 | n/a | n/a | n/a |
| BEATs | block 4: 0.564 (0.450 to 0.661) | −0.129 (−0.195 to −0.070) | < 0.0074 | n/a | n/a | n/a |
| AudioMAE | block 3: 0.307 (0.168 to 0.431) | −0.386 (−0.512 to −0.266) | < 0.0074 | n/a | n/a | n/a |
| VGGish | embedding: 0.463 (0.341 to 0.576) | −0.230 (−0.300 to −0.166) | < 0.0074 | n/a | n/a | n/a |
| ECAPA-TDNN (speaker) | embedding, 1/3 speed: 0.604 (0.461 to 0.723) | −0.090 (−0.183 to −0.017) | 0.2006 | n/a | n/a | n/a |
| ResNet (speaker) | embedding, 1/3 speed: 0.558 (0.419 to 0.681) | −0.135 (−0.226 to −0.060) | < 0.0074 | n/a | n/a | n/a |
| x-vector (speaker) | embedding, 1/3 speed: 0.518 (0.362 to 0.645) | −0.176 (−0.290 to −0.084) | 0.0040 | n/a | n/a | n/a |
| wav2vec 2.0 Base | layer 0, 1/3 speed: 0.740 (0.649 to 0.823) | 0.047 (−0.003 to 0.094) | 0.8736 | n/a | n/a | n/a |
| wav2vec 2.0 Large (robust) | layer 0, 1/3 speed: 0.710 (0.591 to 0.819) | 0.016 (−0.046 to 0.077) | 1.0000 | n/a | n/a | n/a |
| XLS-R 300M | layer 0, 1/3 speed: 0.722 (0.617 to 0.817) | 0.029 (−0.039 to 0.102) | 1.0000 | n/a | n/a | n/a |
| wav2vec 2.0 Conformer Large | layer 0, 1/3 speed: 0.730 (0.625 to 0.825) | 0.037 (−0.033 to 0.113) | 1.0000 | n/a | n/a | n/a |
| MMS 300M | layer 0, 1/3 speed: 0.737 (0.643 to 0.826) | 0.044 (−0.013 to 0.107) | 1.0000 | n/a | n/a | n/a |
| HuBERT Base | layer 0, 1/3 speed: 0.760 (0.677 to 0.840) | 0.067 (0.004 to 0.138) | 0.5880 | n/a | n/a | n/a |
| HuBERT Large | layer 0, 1/3 speed: 0.677 (0.550 to 0.792) | −0.016 (−0.090 to 0.054) | 1.0000 | n/a | n/a | n/a |
| data2vec Base (100 h) | layer 0, 1/3 speed: 0.794 (0.720 to 0.869) | 0.101 (0.062 to 0.145) | < 0.0074 | n/a | n/a | n/a |
| data2vec Base (960 h) | layer 0, 1/3 speed: 0.797 (0.716 to 0.875) | 0.104 (0.055 to 0.161) | < 0.0074 | n/a | n/a | n/a |
| WavLM Base+ | layer 0, 1/3 speed: 0.797 (0.723 to 0.870) | 0.104 (0.055 to 0.160) | < 0.0074 | n/a | n/a | n/a |
| WavLM Large | layer 0, 1/3 speed: 0.686 (0.560 to 0.802) | −0.008 (−0.094 to 0.077) | 1.0000 | n/a | n/a | n/a |

| Neural network | Fisher rule: layer, re-ID accuracy | Difference from BirdNET | Holm p | Held-out rule: layer, re-ID accuracy | Difference from BirdNET | Holm p |
| --- | --- | --- | --- | --- | --- | --- |
| UniSpeech-SAT Base+ | layer 0, 1/3 speed: 0.764 (0.670 to 0.853) | 0.071 (0.008 to 0.136) | 0.4288 | n/a | n/a | n/a |
| XEUS | layer 0, 1/3 speed: 0.798 (0.719 to 0.875) | 0.105 (0.051 to 0.173) | < 0.0074 | n/a | n/a | n/a |
| Loudness control | summary: 0.159 (0.048 to 0.267) | −0.534 (−0.659 to −0.411) | < 0.0074 | n/a | n/a | n/a |
| Duration control | summary: 0.167 (0.000 to 0.394) | −0.527 (−0.738 to −0.287) | < 0.0074 | n/a | n/a | n/a |

*Red-tailed black cockatoo, random split*: BirdNET v2.4 0.961 (0.930 to 0.980).

| Neural network | Fisher rule: layer, re-ID accuracy | Difference from BirdNET | Holm p | Held-out rule: layer, re-ID accuracy | Difference from BirdNET | Holm p |
| --- | --- | --- | --- | --- | --- | --- |
| Perch 2.0 | embedding: 0.961 (0.933 to 0.979) | 0.000 (−0.015 to 0.023) | 1.0000 | n/a | n/a | n/a |
| Perch | embedding: 0.947 (0.920 to 0.973) | −0.014 (−0.043 to 0.029) | 1.0000 | n/a | n/a | n/a |
| SurfPerch | embedding: 0.958 (0.920 to 0.981) | −0.003 (−0.019 to 0.017) | 1.0000 | n/a | n/a | n/a |
| BirdNET v3 (preview) | embedding: 0.854 (0.791 to 0.892) | −0.106 (−0.158 to −0.070) | < 0.0074 | n/a | n/a | n/a |
| AVEX sl-BEATs | block 0: 0.577 (0.279 to 0.742) | −0.384 (−0.672 to −0.223) | < 0.0074 | n/a | n/a | n/a |
| AVEX EfficientNet-B0 | embedding: 0.891 (0.839 to 0.925) | −0.070 (−0.107 to −0.034) | 0.0080 | n/a | n/a | n/a |
| AvesEcho | block 6: 0.798 (0.570 to 0.909) | −0.162 (−0.371 to −0.062) | < 0.0074 | n/a | n/a | n/a |
| AudioProtoPNet | embedding: 0.941 (0.910 to 0.967) | −0.020 (−0.050 to 0.023) | 1.0000 | n/a | n/a | n/a |
| ConvNeXt (BirdSet) | embedding: 0.969 (0.936 to 0.991) | 0.008 (−0.019 to 0.045) | 1.0000 | n/a | n/a | n/a |
| Bird-MAE | block 15: 0.504 (0.133 to 0.731) | −0.457 (−0.810 to −0.236) | < 0.0074 | n/a | n/a | n/a |
| ProtoCLR | embedding: 0.515 (0.219 to 0.713) | −0.445 (−0.726 to −0.259) | < 0.0074 | n/a | n/a | n/a |
| RCL_FS_BSED | embedding: 0.709 (0.436 to 0.841) | −0.252 (−0.510 to −0.122) | < 0.0074 | n/a | n/a | n/a |
| BirdAVES | block 0: 0.910 (0.838 to 0.956) | −0.050 (−0.106 to −0.014) | 0.0408 | n/a | n/a | n/a |
| AVES | block 0: 0.964 (0.917 to 0.986) | 0.003 (−0.025 to 0.021) | 1.0000 | n/a | n/a | n/a |
| NatureBEATs | block 9: 0.947 (0.885 to 0.978) | −0.014 (−0.056 to 0.009) | 1.0000 | n/a | n/a | n/a |
| BioLingual | block 1: 0.468 (0.009 to 0.729) | −0.493 (−0.926 to −0.241) | < 0.0074 | n/a | n/a | n/a |
| BEATs | block 4: 0.678 (0.413 to 0.829) | −0.283 (−0.535 to −0.142) | < 0.0074 | n/a | n/a | n/a |
| AudioMAE | block 3: 0.476 (0.024 to 0.733) | −0.485 (−0.913 to −0.236) | < 0.0074 | n/a | n/a | n/a |
| VGGish | embedding: 0.812 (0.680 to 0.902) | −0.148 (−0.258 to −0.071) | < 0.0074 | n/a | n/a | n/a |
| ECAPA-TDNN (speaker) | embedding, 1/3 speed: 0.810 (0.705 to 0.868) | −0.151 (−0.241 to −0.095) | < 0.0074 | n/a | n/a | n/a |
| ResNet (speaker) | embedding, 1/3 speed: 0.810 (0.709 to 0.864) | −0.151 (−0.231 to −0.102) | < 0.0074 | n/a | n/a | n/a |
| x-vector (speaker) | embedding, 1/3 speed: 0.728 (0.548 to 0.829) | −0.232 (−0.393 to −0.135) | < 0.0074 | n/a | n/a | n/a |
| wav2vec 2.0 Base | layer 1, full speed: 0.871 (0.787 to 0.915) | −0.090 (−0.154 to −0.053) | < 0.0074 | n/a | n/a | n/a |
| wav2vec 2.0 Large (robust) | layer 20, 1/2 speed: 0.922 (0.822 to 0.969) | −0.039 (−0.122 to 0.004) | 1.0000 | n/a | n/a | n/a |
| XLS-R 300M | layer 0, 1/3 speed: 0.910 (0.825 to 0.950) | −0.050 (−0.121 to −0.014) | 0.1066 | n/a | n/a | n/a |
| wav2vec 2.0 Conformer Large | layer 0, 1/3 speed: 0.894 (0.801 to 0.938) | −0.067 (−0.146 to −0.021) | 0.0420 | n/a | n/a | n/a |
| MMS 300M | layer 0, 1/3 speed: 0.908 (0.814 to 0.951) | −0.053 (−0.131 to −0.016) | 0.0504 | n/a | n/a | n/a |

| Neural network | Fisher rule: layer, re-ID accuracy | Difference from BirdNET | Holm p | Held-out rule: layer, re-ID accuracy | Difference from BirdNET | Holm p |
|---|---|---|---|---|---|---|
| HuBERT Base | layer 0, 1/3 speed: 0.916 (0.843 to 0.950) | −0.045 (−0.104 to −0.014) | 0.0408 | n/a | n/a | n/a |
| HuBERT Large | layer 0, 1/3 speed: 0.846 (0.717 to 0.917) | −0.115 (−0.231 to −0.046) | < 0.0074 | n/a | n/a | n/a |
| data2vec Base (100 h) | layer 0, 1/3 speed: 0.936 (0.877 to 0.967) | −0.025 (−0.072 to 0.008) | 1.0000 | n/a | n/a | n/a |
| data2vec Base (960 h) | layer 0, 1/3 speed: 0.938 (0.878 to 0.968) | −0.022 (−0.073 to 0.012) | 1.0000 | n/a | n/a | n/a |
| WavLM Base+ | layer 0, 1/3 speed: 0.924 (0.856 to 0.954) | −0.036 (−0.092 to −0.007) | 0.1536 | n/a | n/a | n/a |
| WavLM Large | layer 0, 1/3 speed: 0.846 (0.730 to 0.906) | −0.115 (−0.219 to −0.054) | 0.0080 | n/a | n/a | n/a |
| UniSpeech-SAT Base+ | layer 0, 1/3 speed: 0.916 (0.843 to 0.947) | −0.045 (−0.103 to −0.016) | 0.0180 | n/a | n/a | n/a |
| XEUS | layer 0, 1/3 speed: 0.899 (0.822 to 0.943) | −0.062 (−0.133 to −0.010) | 0.1826 | n/a | n/a | n/a |
| Loudness control | summary: 0.406 (0.153 to 0.587) | −0.555 (−0.793 to −0.375) | < 0.0074 | n/a | n/a | n/a |
| Duration control | summary: 0.336 (0.000 to 0.666) | −0.625 (−0.958 to −0.304) | < 0.0074 | n/a | n/a | n/a |

*Egyptian fruit bat, across year*: BirdNET v2.4 0.381 (0.189 to 0.561).

| Neural network | Fisher rule: layer, re-ID accuracy | Difference from BirdNET | Holm p | Held-out rule: layer, re-ID accuracy | Difference from BirdNET | Holm p |
|---|---|---|---|---|---|---|
| Perch 2.0 | embedding: 0.350 (0.161 to 0.532) | −0.031 (−0.115 to 0.104) | 1.0000 | embedding: 0.350 (0.161 to 0.532) | −0.031 (−0.115 to 0.104) | 1.0000 |
| Perch | embedding: 0.268 (0.131 to 0.537) | −0.113 (−0.353 to 0.180) | 1.0000 | embedding: 0.268 (0.131 to 0.537) | −0.113 (−0.353 to 0.180) | 1.0000 |
| SurfPerch | embedding: 0.353 (0.202 to 0.496) | −0.028 (−0.136 to 0.104) | 1.0000 | embedding: 0.353 (0.202 to 0.496) | −0.028 (−0.136 to 0.104) | 1.0000 |
| BirdNET v3 (preview) | embedding: 0.293 (0.194 to 0.475) | −0.088 (−0.272 to 0.103) | 1.0000 | embedding: 0.293 (0.194 to 0.475) | −0.088 (−0.272 to 0.103) | 1.0000 |
| AVEX sl-BEATs | block 0: 0.114 (0.037 to 0.238) | −0.267 (−0.495 to −0.054) | 0.0072 | block 10: 0.378 (0.207 to 0.512) | −0.003 (−0.082 to 0.085) | 1.0000 |
| AVEX EfficientNet-B0 | embedding: 0.307 (0.258 to 0.359) | −0.074 (−0.237 to 0.088) | 1.0000 | embedding: 0.307 (0.258 to 0.359) | −0.074 (−0.237 to 0.088) | 1.0000 |
| AvesEcho | block 5: 0.151 (0.028 to 0.381) | −0.231 (−0.472 to 0.028) | 1.0000 | embedding: 0.223 (0.160 to 0.292) | −0.158 (−0.299 to −0.027) | 0.0518 |
| AudioProtoPNet | embedding: 0.305 (0.131 to 0.512) | −0.077 (−0.189 to 0.100) | 1.0000 | embedding: 0.305 (0.131 to 0.512) | −0.077 (−0.189 to 0.100) | 1.0000 |
| ConvNeXt (BirdSet) | embedding: 0.333 (0.127 to 0.530) | −0.048 (−0.121 to 0.061) | 1.0000 | embedding: 0.333 (0.127 to 0.530) | −0.048 (−0.121 to 0.061) | 1.0000 |
| Bird-MAE | block 8: 0.114 (0.046 to 0.200) | −0.267 (−0.490 to −0.041) | 0.1768 | block 25: 0.123 (0.048 to 0.220) | −0.259 (−0.479 to −0.031) | 0.4968 |
| ProtoCLR | block 10: 0.184 (0.019 to 0.384) | −0.197 (−0.518 to 0.172) | 1.0000 | embedding: 0.133 (0.052 to 0.233) | −0.249 (−0.494 to −0.001) | 1.0000 |
| RCL_FS_BSED | embedding: 0.191 (0.077 to 0.327) | −0.191 (−0.467 to 0.056) | 1.0000 | embedding: 0.191 (0.077 to 0.327) | −0.191 (−0.467 to 0.056) | 1.0000 |
| BirdAVES | block 0: 0.246 (0.138 to 0.329) | −0.136 (−0.336 to 0.100) | 1.0000 | block 21: 0.348 (0.123 to 0.569) | −0.033 (−0.163 to 0.041) | 1.0000 |
| AVES | block 0: 0.283 (0.024 to 0.473) | −0.098 (−0.311 to 0.057) | 1.0000 | block 2: 0.280 (0.041 to 0.473) | −0.101 (−0.294 to 0.030) | 1.0000 |
| NatureBEATs | block 3: 0.131 (0.051 to 0.258) | −0.250 (−0.481 to −0.019) | 1.0000 | block 9: 0.350 (0.114 to 0.584) | −0.031 (−0.111 to 0.023) | 1.0000 |
| BioLingual | block 2: 0.119 (0.034 to 0.231) | −0.263 (−0.513 to −0.010) | 1.0000 | embedding: 0.209 (0.122 to 0.319) | −0.172 (−0.417 to 0.054) | 1.0000 |
| BEATs | block 7: 0.223 (0.156 to 0.298) | −0.158 (−0.382 to 0.062) | 1.0000 | block 9: 0.259 (0.180 to 0.395) | −0.123 (−0.352 to 0.050) | 1.0000 |
| AudioMAE | block 1: 0.190 (0.005 to 0.415) | −0.192 (−0.534 to 0.204) | 1.0000 | block 10: 0.184 (0.104 to 0.328) | −0.197 (−0.423 to 0.037) | 1.0000 |
| VGGish | embedding: 0.166 (0.081 to 0.216) | −0.216 (−0.370 to −0.033) | 0.4950 | embedding: 0.166 (0.081 to 0.216) | −0.216 (−0.370 to −0.033) | 0.5250 |
| ECAPA-TDNN (speaker) | embedding, 1/2 speed: 0.297 (0.187 to 0.411) | −0.084 (−0.199 to 0.048) | 1.0000 | embedding, 1/3 speed: 0.272 (0.147 to 0.406) | −0.110 (−0.232 to 0.034) | 1.0000 |
| ResNet (speaker) | embedding: 0.319 (0.204 to 0.412) | −0.062 (−0.244 to 0.144) | 1.0000 | embedding, 1/3 speed: 0.265 (0.182 to 0.375) | −0.116 (−0.287 to 0.053) | 1.0000 |

| Neural network | Fisher rule: layer, re-ID accuracy | Difference from BirdNET | Holm p | Held-out rule: layer, re-ID accuracy | Difference from BirdNET | Holm p |
|---|---|---|---|---|---|---|
| x-vector (speaker) | embedding, 1/3 speed: 0.305 (0.144 to 0.422) | −0.077 (−0.186 to 0.057) | 1.0000 | embedding, 1/3 speed: 0.305 (0.144 to 0.422) | −0.077 (−0.186 to 0.057) | 1.0000 |
| wav2vec 2.0 Base | layer 0, full speed: 0.233 (0.063 to 0.364) | −0.149 (−0.384 to 0.125) | 1.0000 | layer 3, 1/3 speed: 0.365 (0.139 to 0.561) | −0.016 (−0.120 to 0.053) | 1.0000 |
| wav2vec 2.0 Large (robust) | layer 0, full speed: 0.231 (0.045 to 0.349) | −0.151 (−0.320 to 0.104) | 1.0000 | layer 3, 1/2 speed: 0.394 (0.159 to 0.597) | 0.013 (−0.103 to 0.133) | 1.0000 |
| XLS-R 300M | layer 1, full speed: 0.258 (0.146 to 0.349) | −0.124 (−0.234 to −0.007) | 1.0000 | layer 5, 1/3 speed: 0.265 (0.134 to 0.456) | −0.116 (−0.361 to 0.140) | 1.0000 |
| wav2vec 2.0 Conformer Large | layer 2, full speed: 0.283 (0.091 to 0.429) | −0.098 (−0.288 to 0.171) | 1.0000 | layer 5, 1/3 speed: 0.310 (0.153 to 0.486) | −0.071 (−0.233 to 0.125) | 1.0000 |
| MMS 300M | layer 0, full speed: 0.218 (0.060 to 0.320) | −0.164 (−0.269 to −0.001) | 1.0000 | layer 4, 1/3 speed: 0.367 (0.166 to 0.530) | −0.014 (−0.147 to 0.155) | 1.0000 |
| HuBERT Base | layer 0, full speed: 0.249 (0.026 to 0.385) | −0.133 (−0.337 to 0.116) | 1.0000 | layer 2, 1/3 speed: 0.345 (0.103 to 0.578) | −0.037 (−0.216 to 0.068) | 1.0000 |
| HuBERT Large | layer 0, full speed: 0.216 (0.049 to 0.358) | −0.166 (−0.242 to −0.060) | 0.0700 | layer 5, 1/3 speed: 0.319 (0.159 to 0.475) | −0.062 (−0.217 to 0.118) | 1.0000 |
| data2vec Base (100 h) | layer 0, full speed: 0.255 (0.027 to 0.413) | −0.126 (−0.335 to 0.169) | 1.0000 | layer 1, 1/3 speed: 0.337 (0.196 to 0.483) | −0.044 (−0.215 to 0.154) | 1.0000 |
| data2vec Base (960 h) | layer 0, full speed: 0.258 (0.034 to 0.419) | −0.124 (−0.327 to 0.170) | 1.0000 | layer 1, 1/3 speed: 0.341 (0.243 to 0.412) | −0.041 (−0.189 to 0.118) | 1.0000 |
| WavLM Base+ | layer 0, full speed: 0.250 (0.040 to 0.389) | −0.131 (−0.328 to 0.128) | 1.0000 | layer 1, 1/3 speed: 0.369 (0.136 to 0.586) | −0.013 (−0.187 to 0.088) | 1.0000 |
| WavLM Large | layer 0, full speed: 0.238 (0.018 to 0.458) | −0.143 (−0.214 to −0.093) | < 0.0074 | layer 6, 1/3 speed: 0.414 (0.157 to 0.636) | 0.032 (−0.086 to 0.096) | 1.0000 |
| UniSpeech-SAT Base+ | layer 0, full speed: 0.235 (0.023 to 0.420) | −0.147 (−0.377 to 0.180) | 1.0000 | layer 2, 1/3 speed: 0.334 (0.132 to 0.518) | −0.047 (−0.153 to 0.019) | 1.0000 |
| XEUS | layer 0, 1/2 speed: 0.306 (0.047 to 0.547) | −0.075 (−0.262 to 0.034) | 1.0000 | layer 4, 1/3 speed: 0.364 (0.075 to 0.626) | −0.017 (−0.213 to 0.103) | 1.0000 |
| Loudness control | summary: 0.248 (0.013 to 0.620) | −0.134 (−0.531 to 0.395) | 1.0000 | summary: 0.248 (0.013 to 0.620) | −0.134 (−0.531 to 0.395) | 1.0000 |
| Duration control | summary: 0.208 (0.000 to 0.462) | −0.173 (−0.537 to 0.249) | 1.0000 | summary: 0.208 (0.000 to 0.462) | −0.173 (−0.537 to 0.249) | 1.0000 |

*Chiffchaff, within year*: BirdNET v2.4 0.765 (0.680 to 0.841).

| Neural network | Fisher rule: layer, re-ID accuracy | Difference from BirdNET | Holm p | Held-out rule: layer, re-ID accuracy | Difference from BirdNET | Holm p |
|---|---|---|---|---|---|---|
| Perch 2.0 | embedding: 0.831 (0.756 to 0.884) | 0.066 (−0.008 to 0.122) | 0.5782 | n/a | n/a | n/a |
| Perch | embedding: 0.778 (0.656 to 0.858) | 0.013 (−0.105 to 0.094) | 1.0000 | n/a | n/a | n/a |
| SurfPerch | embedding: 0.729 (0.632 to 0.808) | −0.036 (−0.115 to 0.013) | 0.8892 | n/a | n/a | n/a |
| BirdNET v3 (preview) | embedding: 0.624 (0.370 to 0.770) | −0.141 (−0.381 to −0.003) | 0.3504 | n/a | n/a | n/a |
| AVEX sl-BEATs | block 9: 0.737 (0.585 to 0.817) | −0.027 (−0.127 to 0.023) | 1.0000 | n/a | n/a | n/a |
| AVEX EfficientNet-B0 | embedding: 0.691 (0.577 to 0.760) | −0.074 (−0.173 to −0.021) | 0.0400 | n/a | n/a | n/a |
| AvesEcho | embedding: 0.691 (0.518 to 0.772) | −0.073 (−0.217 to −0.008) | 0.2160 | n/a | n/a | n/a |
| AudioProtoPNet | embedding: 0.806 (0.646 to 0.890) | 0.042 (−0.096 to 0.114) | 1.0000 | n/a | n/a | n/a |
| ConvNeXt (BirdSet) | embedding: 0.805 (0.687 to 0.866) | 0.041 (−0.066 to 0.089) | 1.0000 | n/a | n/a | n/a |
| Bird-MAE | block 12: 0.319 (0.114 to 0.447) | −0.446 (−0.633 to −0.339) | < 0.0074 | n/a | n/a | n/a |
| ProtoCLR | embedding: 0.325 (0.211 to 0.477) | −0.439 (−0.548 to −0.305) | < 0.0074 | n/a | n/a | n/a |
| RCL_FS_BSED | embedding: 0.256 (0.121 to 0.494) | −0.508 (−0.643 to −0.289) | < 0.0074 | n/a | n/a | n/a |

| Neural network | Fisher rule: layer, re-ID accuracy | Difference from BirdNET | Holm p | Held-out rule: layer, re-ID accuracy | Difference from BirdNET | Holm p |
|---|---|---|---|---|---|---|
| BirdAVES | block 0: 0.420 (0.331 to 0.588) | −0.345 (−0.430 to −0.205) | < 0.0074 | n/a | n/a | n/a |
| AVES | block 0: 0.397 (0.309 to 0.554) | −0.368 (−0.438 to −0.251) | < 0.0074 | n/a | n/a | n/a |
| NatureBEATs | block 9: 0.741 (0.577 to 0.816) | −0.024 (−0.156 to 0.035) | 1.0000 | n/a | n/a | n/a |
| BioLingual | block 10: 0.637 (0.449 to 0.743) | −0.127 (−0.281 to −0.052) | < 0.0074 | n/a | n/a | n/a |
| BEATs | block 9: 0.558 (0.340 to 0.673) | −0.207 (−0.401 to −0.118) | < 0.0074 | n/a | n/a | n/a |
| AudioMAE | block 2: 0.184 (0.122 to 0.279) | −0.581 (−0.658 to −0.468) | < 0.0074 | n/a | n/a | n/a |
| VGGish | embedding: 0.458 (0.256 to 0.561) | −0.307 (−0.501 to −0.223) | < 0.0074 | n/a | n/a | n/a |
| ECAPA-TDNN (speaker) | embedding, 1/3 speed: 0.416 (0.303 to 0.545) | −0.348 (−0.438 to −0.258) | < 0.0074 | n/a | n/a | n/a |
| ResNet (speaker) | embedding, 1/3 speed: 0.523 (0.327 to 0.633) | −0.242 (−0.411 to −0.153) | < 0.0074 | n/a | n/a | n/a |
| x-vector (speaker) | embedding, 1/2 speed: 0.517 (0.326 to 0.634) | −0.248 (−0.404 to −0.170) | < 0.0074 | n/a | n/a | n/a |
| wav2vec 2.0 Base | layer 0, full speed: 0.331 (0.234 to 0.522) | −0.434 (−0.519 to −0.274) | < 0.0074 | n/a | n/a | n/a |
| wav2vec 2.0 Large (robust) | layer 0, 1/2 speed: 0.305 (0.208 to 0.444) | −0.460 (−0.555 to −0.343) | < 0.0074 | n/a | n/a | n/a |
| XLS-R 300M | layer 0, 1/2 speed: 0.321 (0.213 to 0.440) | −0.444 (−0.548 to −0.345) | < 0.0074 | n/a | n/a | n/a |
| wav2vec 2.0 Conformer Large | layer 2, full speed: 0.414 (0.304 to 0.575) | −0.351 (−0.425 to −0.236) | < 0.0074 | n/a | n/a | n/a |
| MMS 300M | layer 0, 1/2 speed: 0.324 (0.224 to 0.473) | −0.441 (−0.531 to −0.319) | < 0.0074 | n/a | n/a | n/a |
| HuBERT Base | layer 0, full speed: 0.314 (0.218 to 0.496) | −0.451 (−0.536 to −0.298) | < 0.0074 | n/a | n/a | n/a |
| HuBERT Large | layer 0, 1/2 speed: 0.316 (0.215 to 0.442) | −0.449 (−0.540 to −0.351) | < 0.0074 | n/a | n/a | n/a |
| data2vec Base (100 h) | layer 1, 1/3 speed: 0.382 (0.279 to 0.503) | −0.383 (−0.489 to −0.276) | < 0.0074 | n/a | n/a | n/a |
| data2vec Base (960 h) | layer 0, 1/3 speed: 0.378 (0.268 to 0.491) | −0.386 (−0.487 to −0.298) | < 0.0074 | n/a | n/a | n/a |
| WavLM Base+ | layer 0, full speed: 0.327 (0.233 to 0.512) | −0.438 (−0.519 to −0.283) | < 0.0074 | n/a | n/a | n/a |
| WavLM Large | layer 0, 1/2 speed: 0.347 (0.225 to 0.470) | −0.417 (−0.524 to −0.325) | < 0.0074 | n/a | n/a | n/a |
| UniSpeech-SAT Base+ | layer 0, full speed: 0.317 (0.222 to 0.497) | −0.447 (−0.534 to −0.296) | < 0.0074 | n/a | n/a | n/a |
| XEUS | layer 0, 1/2 speed: 0.341 (0.236 to 0.465) | −0.424 (−0.518 to −0.324) | < 0.0074 | n/a | n/a | n/a |
| Loudness control | summary: 0.094 (0.010 to 0.267) | −0.671 (−0.747 to −0.522) | < 0.0074 | n/a | n/a | n/a |
| Duration control | summary: 0.040 (0.000 to 0.147) | −0.725 (−0.794 to −0.607) | < 0.0074 | n/a | n/a | n/a |

*Rook, across year*: BirdNET v2.4 0.375 (0.261 to 0.506).

| Neural network | Fisher rule: layer, re-ID accuracy | Difference from BirdNET | Holm p | Held-out rule: layer, re-ID accuracy | Difference from BirdNET | Holm p |
|---|---|---|---|---|---|---|
| Perch 2.0 | embedding: 0.418 (0.306 to 0.535) | 0.043 (0.017 to 0.065) | 0.0384 | embedding: 0.418 (0.306 to 0.535) | 0.043 (0.017 to 0.065) | 0.0336 |

| Neural network | Fisher rule: layer, re-ID accuracy | Difference from BirdNET | Holm p | Held-out rule: layer, re-ID accuracy | Difference from BirdNET | Holm p |
|---|---|---|---|---|---|---|
| Perch | embedding: 0.393 (0.282 to 0.516) | 0.018 (−0.027 to 0.062) | 0.9816 | embedding: 0.393 (0.282 to 0.516) | 0.018 (−0.027 to 0.062) | 1.0000 |
| SurfPerch | embedding: 0.306 (0.200 to 0.425) | −0.069 (−0.106 to −0.038) | 0.0050 | embedding: 0.306 (0.200 to 0.425) | −0.069 (−0.106 to −0.038) | 0.0046 |
| BirdNET v3 (preview) | embedding: 0.328 (0.212 to 0.439) | −0.048 (−0.098 to −0.008) | 0.1632 | embedding: 0.328 (0.212 to 0.439) | −0.048 (−0.098 to −0.008) | 0.1632 |
| AVEX sl-BEATs | block 5: 0.284 (0.153 to 0.442) | −0.092 (−0.177 to −0.020) | 0.1152 | block 10: 0.342 (0.202 to 0.457) | −0.033 (−0.102 to 0.019) | 1.0000 |
| AVEX EfficientNet-B0 | embedding: 0.345 (0.247 to 0.442) | −0.031 (−0.073 to −0.001) | 0.2580 | embedding: 0.345 (0.247 to 0.442) | −0.031 (−0.073 to −0.001) | 0.3010 |
| AvesEcho | embedding: 0.343 (0.223 to 0.437) | −0.033 (−0.125 to 0.024) | 0.9816 | embedding: 0.343 (0.223 to 0.437) | −0.033 (−0.125 to 0.024) | 1.0000 |
| AudioProtoPNet | embedding: 0.386 (0.269 to 0.506) | 0.011 (−0.008 to 0.024) | 0.9248 | embedding: 0.386 (0.269 to 0.506) | 0.011 (−0.008 to 0.024) | 1.0000 |
| ConvNeXt (BirdSet) | embedding: 0.409 (0.260 to 0.536) | 0.034 (−0.044 to 0.083) | 0.9816 | embedding: 0.409 (0.260 to 0.536) | 0.034 (−0.044 to 0.083) | 1.0000 |
| Bird-MAE | block 1: 0.125 (0.055 to 0.225) | −0.250 (−0.341 to −0.145) | < 0.0074 | block 0: 0.095 (0.038 to 0.200) | −0.281 (−0.375 to −0.166) | < 0.0074 |
| ProtoCLR | block 1: 0.101 (0.039 to 0.211) | −0.274 (−0.352 to −0.164) | < 0.0074 | block 6: 0.079 (0.015 to 0.213) | −0.296 (−0.383 to −0.167) | < 0.0074 |
| RCL_FS_BSED | embedding: 0.105 (0.032 to 0.215) | −0.270 (−0.350 to −0.178) | < 0.0074 | embedding: 0.105 (0.032 to 0.215) | −0.270 (−0.350 to −0.178) | < 0.0074 |
| BirdAVES | block 11: 0.151 (0.032 to 0.366) | −0.225 (−0.322 to −0.095) | 0.0050 | block 23: 0.174 (0.053 to 0.384) | −0.202 (−0.305 to −0.073) | 0.0144 |
| AVES | block 10: 0.179 (0.055 to 0.375) | −0.197 (−0.283 to −0.091) | 0.0050 | block 9: 0.173 (0.047 to 0.374) | −0.202 (−0.293 to −0.093) | 0.0046 |
| NatureBEATs | block 3: 0.166 (0.044 to 0.394) | −0.209 (−0.320 to −0.064) | 0.0660 | block 7: 0.277 (0.109 to 0.455) | −0.099 (−0.206 to −0.014) | 0.1458 |
| BioLingual | block 11: 0.281 (0.084 to 0.479) | −0.095 (−0.233 to 0.018) | 0.5330 | embedding: 0.303 (0.186 to 0.404) | −0.073 (−0.139 to −0.025) | 0.0170 |
| BEATs | block 4: 0.207 (0.086 to 0.410) | −0.168 (−0.272 to −0.044) | 0.0728 | block 10: 0.341 (0.208 to 0.456) | −0.034 (−0.092 to 0.005) | 0.5436 |
| AudioMAE | block 7: 0.185 (0.038 to 0.418) | −0.191 (−0.310 to −0.038) | 0.1520 | block 11: 0.234 (0.115 to 0.410) | −0.141 (−0.226 to −0.048) | 0.0270 |
| VGGish | embedding: 0.180 (0.086 to 0.329) | −0.195 (−0.266 to −0.117) | < 0.0074 | embedding: 0.180 (0.086 to 0.329) | −0.195 (−0.266 to −0.117) | < 0.0074 |
| ECAPA-TDNN (speaker) | embedding, 1/3 speed: 0.271 (0.180 to 0.365) | −0.105 (−0.200 to −0.023) | 0.1520 | embedding, 1/2 speed: 0.281 (0.169 to 0.400) | −0.094 (−0.160 to −0.035) | 0.0552 |
| ResNet (speaker) | embedding, 1/2 speed: 0.297 (0.152 to 0.451) | −0.078 (−0.165 to −0.016) | 0.1092 | embedding: 0.239 (0.125 to 0.435) | −0.137 (−0.221 to −0.035) | 0.0552 |
| x-vector (speaker) | embedding, 1/2 speed: 0.296 (0.161 to 0.447) | −0.079 (−0.174 to −0.004) | 0.2534 | embedding: 0.262 (0.136 to 0.417) | −0.114 (−0.163 to −0.064) | < 0.0074 |
| wav2vec 2.0 Base | layer 0, full speed: 0.153 (0.056 to 0.344) | −0.222 (−0.327 to −0.091) | < 0.0074 | layer 3, full speed: 0.202 (0.096 to 0.369) | −0.174 (−0.258 to −0.082) | < 0.0074 |
| wav2vec 2.0 Large (robust) | layer 24, full speed: 0.125 (0.012 to 0.267) | −0.250 (−0.409 to −0.089) | 0.0136 | layer 23, full speed: 0.125 (0.012 to 0.264) | −0.251 (−0.408 to −0.091) | 0.0120 |
| XLS-R 300M | layer 22, full speed: 0.187 (0.067 to 0.384) | −0.188 (−0.278 to −0.083) | 0.0080 | layer 14, full speed: 0.226 (0.088 to 0.370) | −0.150 (−0.224 to −0.101) | < 0.0074 |
| wav2vec 2.0 Conformer Large | layer 11, 1/2 speed: 0.256 (0.138 to 0.395) | −0.120 (−0.227 to −0.024) | 0.1152 | layer 8, full speed: 0.281 (0.134 to 0.418) | −0.094 (−0.164 to −0.049) | < 0.0074 |
| MMS 300M | layer 24, 1/2 speed: 0.163 (0.048 to 0.344) | −0.212 (−0.316 to −0.112) | < 0.0074 | layer 8, full speed: 0.224 (0.090 to 0.397) | −0.151 (−0.234 to −0.075) | < 0.0074 |
| HuBERT Base | layer 12, 1/2 speed: 0.235 (0.089 to 0.400) | −0.140 (−0.226 to −0.071) | < 0.0074 | layer 8, full speed: 0.250 (0.112 to 0.420) | −0.125 (−0.212 to −0.051) | 0.0224 |
| HuBERT Large | layer 1, 1/3 speed: 0.197 (0.058 to 0.319) | −0.178 (−0.295 to −0.113) | < 0.0074 | layer 17, full speed: 0.170 (0.070 to 0.317) | −0.206 (−0.299 to −0.122) | < 0.0074 |
| data2vec Base (100 h) | layer 11, full speed: 0.169 (0.060 to 0.319) | −0.207 (−0.293 to −0.137) | < 0.0074 | layer 5, full speed: 0.213 (0.078 to 0.377) | −0.162 (−0.240 to −0.094) | < 0.0074 |
| data2vec Base (960 h) | layer 2, full speed: 0.236 (0.095 to 0.392) | −0.139 (−0.223 to −0.077) | < 0.0074 | layer 9, full speed: 0.163 (0.050 to 0.349) | −0.213 (−0.306 to −0.114) | < 0.0074 |
| WavLM Base+ | layer 12, 1/2 speed: 0.105 (0.022 to 0.287) | −0.270 (−0.367 to −0.151) | < 0.0074 | layer 0, full speed: 0.162 (0.061 to 0.354) | −0.213 (−0.310 to −0.098) | < 0.0074 |
| WavLM Large | layer 1, 1/3 speed: 0.166 (0.049 to 0.365) | −0.210 (−0.312 to −0.094) | 0.0050 | layer 3, 1/2 speed: 0.210 (0.091 to 0.371) | −0.165 (−0.289 to −0.057) | 0.0120 |

| Neural network | Fisher rule: layer, re-ID accuracy | Difference from BirdNET | Holm p | Held-out rule: layer, re-ID accuracy | Difference from BirdNET | Holm p |
|---|---|---|---|---|---|---|
| UniSpeech-SAT Base+ | layer 1, full speed: 0.181 (0.059 to 0.375) | −0.195 (−0.293 to −0.084) | 0.0080 | layer 9, 1/2 speed: 0.205 (0.076 to 0.392) | −0.170 (−0.287 to −0.055) | 0.0336 |
| XEUS | layer 1, 1/2 speed: 0.247 (0.111 to 0.408) | −0.128 (−0.236 to −0.043) | 0.0108 | layer 18, 1/2 speed: 0.254 (0.077 to 0.430) | −0.122 (−0.260 to −0.021) | 0.0880 |
| Loudness control | summary: 0.126 (0.019 to 0.342) | −0.249 (−0.363 to −0.108) | < 0.0074 | summary: 0.126 (0.019 to 0.342) | −0.249 (−0.363 to −0.108) | < 0.0074 |
| Duration control | summary: 0.030 (0.000 to 0.113) | −0.345 (−0.467 to −0.210) | 0.0050 | summary: 0.030 (0.000 to 0.113) | −0.345 (−0.467 to −0.210) | 0.0046 |

*Notes:* Re-ID accuracy with a 95% interval over individuals; difference from BirdNET with a paired interval over the same 10,000 draws of individuals; two-sided bootstrap p = min(1, 2 × min(proportion of differences ≤ 0, proportion ≥ 0)), with zero differences counted on both sides, then adjusted by Holm's method over the 37 comparisons on that dataset and rule. When no draw crossed zero the empirical estimate is zero; < 0.0074 denotes the display resolution 37 × 2 / 10,000, not a confidence bound on the underlying p value. BirdNET's interval comes from the same paired draws, so its bounds can differ slightly from those in Table 2. n/a: the held-out rule does not apply, because the dataset has fewer than two enrolment sessions per individual, and on such datasets it would choose what the Fisher rule chose.

**Table S14.** Consistency of the ranking of embeddings across datasets under the three layer rules, and each entry's average rank.

| Layer rule | Entries ranked | Friedman χ² | p | Kendall's W | Nemenyi critical difference |
|---|---|---|---|---|---|
| Fisher, 13 datasets | 38 | 224.7 | < 0.0001 | 0.47 | 16.8 |
| Held-out, 6 datasets | 38 | 138.3 | < 0.0001 | 0.62 | 24.8 |
| Own final output, 13 datasets | 38 | 307.7 | < 0.0001 | 0.64 | 16.8 |

| Embedding | Average rank, Fisher rule | Average rank, held-out rule | Average rank, own final output |
|---|---|---|---|
| Perch 2.0 | 4.12 | 3.67 | 3.38 |
| ConvNeXt (BirdSet) | 6.00 | 5.58 | 5.77 |
| BirdNET v2.4 | 6.27 | 4.00 | 5.62 |
| SurfPerch | 7.69 | 10.50 | 6.96 |
| AudioProtoPNet | 8.08 | 7.33 | 7.81 |
| Perch | 8.35 | 9.83 | 7.92 |
| AVEX EfficientNet-B0 | 11.92 | 9.58 | 12.12 |
| ResNet (speaker) | 14.50 | 15.92 | 18.81 |
| ECAPA-TDNN (speaker) | 15.19 | 18.92 | 19.73 |
| XEUS | 16.04 | 13.92 | 22.46 |
| x-vector (speaker) | 16.08 | 18.42 | 18.62 |
| BirdNET v3 (preview) | 16.19 | 15.67 | 15.81 |
| data2vec Base (960 h) | 17.42 | 22.67 | 30.54 |
| data2vec Base (100 h) | 17.54 | 18.42 | 30.38 |
| wav2vec 2.0 Conformer Large | 17.73 | 19.58 | 19.31 |
| BirdAVES | 19.00 | 25.08 | 14.08 |
| wav2vec 2.0 Base | 19.42 | 23.00 | 27.04 |
| AVES | 19.58 | 31.08 | 19.46 |
| XLS-R 300M | 19.85 | 18.83 | 26.77 |
| WavLM Base+ | 20.08 | 19.75 | 24.04 |
| UniSpeech-SAT Base+ | 20.19 | 16.67 | 21.00 |
| MMS 300M | 20.46 | 17.50 | 25.00 |
| HuBERT Base | 20.88 | 21.25 | 21.00 |
| WavLM Large | 21.19 | 16.83 | 18.00 |
| AVEX sl-BEATs | 21.62 | 11.08 | 9.15 |
| HuBERT Large | 22.31 | 23.42 | 22.73 |
| NatureBEATs | 22.73 | 17.83 | 9.96 |
| BEATs | 22.77 | 17.75 | 18.77 |
| wav2vec 2.0 Large (robust) | 23.69 | 26.50 | 33.31 |
| AvesEcho | 25.38 | 13.58 | 9.81 |
| VGGish | 26.38 | 27.83 | 25.88 |
| BioLingual | 26.58 | 17.67 | 12.58 |
| RCL_FS_BSED | 28.04 | 32.67 | 28.58 |
| AudioMAE | 29.85 | 28.83 | 19.46 |
| ProtoCLR | 30.19 | 35.75 | 30.54 |
| Loudness control | 31.58 | 34.50 | 33.88 |
| Duration control | 33.04 | 34.42 | 34.69 |
| Bird-MAE | 33.08 | 35.17 | 30.04 |

*Notes:* Rank 1 is the highest re-ID accuracy on a dataset. Held-out rule: the six datasets where it applies. Own final output: specified after the review, so exploratory. Comparisons with BirdNET after Holm's correction: Fisher rule 9 of 481 above and 183 below; held-out rule 1 of 222 above and 79 below; own final output 5 of 481 above and 200 below. On the six datasets, the held-out rule changed the chosen layer in 130 of 222 comparisons, raising re-ID accuracy in 82 and lowering it in 45.

**Table S15.** Re-ID with strangers present for all 36 neural networks: medians over 16 allocations, with ranges.

| Dataset | Embedding | Names correctly | Separation | Accepted and named, calibrated | Strangers accepted, calibrated | Balanced accuracy, known | Balanced accuracy, strangers | Geometric mean |
|---|---|---|---|---|---|---|---|---|
| Great tit, across year | BirdNET v2.4 | 0.600 (0.375 to 0.700) | 0.617 (0.466 to 0.733) | 0.175 (0.050 to 0.350) | 0.087 (0.000 to 0.350) | 0.175 (0.050 to 0.350) | 0.913 (0.650 to 1.000) | 0.394 (0.221 to 0.494) |
| Great tit, across year | Perch 2.0 | 0.625 (0.400 to 0.700) | 0.616 (0.497 to 0.667) | 0.225 (0.025 to 0.350) | 0.087 (0.000 to 0.275) | 0.225 (0.025 to 0.350) | 0.913 (0.725 to 1.000) | 0.419 (0.158 to 0.556) |
| Great tit, across year | Perch | 0.425 (0.275 to 0.625) | 0.534 (0.324 to 0.676) | 0.062 (0.000 to 0.225) | 0.087 (0.025 to 0.300) | 0.062 (0.000 to 0.225) | 0.913 (0.700 to 0.975) | 0.239 (0.000 to 0.400) |
| Great tit, across year | SurfPerch | 0.562 (0.350 to 0.750) | 0.584 (0.443 to 0.675) | 0.150 (0.025 to 0.250) | 0.087 (0.000 to 0.350) | 0.150 (0.025 to 0.250) | 0.912 (0.650 to 1.000) | 0.359 (0.156 to 0.468) |
| Great tit, across year | BirdNET v3 (preview) | 0.325 (0.200 to 0.500) | 0.540 (0.349 to 0.610) | 0.038 (0.000 to 0.125) | 0.087 (0.000 to 0.350) | 0.038 (0.000 to 0.125) | 0.913 (0.650 to 1.000) | 0.176 (0.000 to 0.285) |
| Great tit, across year | AVEX sl-BEATs | 0.412 (0.200 to 0.625) | 0.530 (0.440 to 0.666) | 0.062 (0.000 to 0.225) | 0.100 (0.000 to 0.375) | 0.062 (0.000 to 0.225) | 0.900 (0.625 to 1.000) | 0.238 (0.000 to 0.404) |
| Great tit, across year | AVEX EfficientNet-B0 | 0.500 (0.375 to 0.600) | 0.603 (0.437 to 0.728) | 0.138 (0.025 to 0.200) | 0.087 (0.000 to 0.250) | 0.138 (0.025 to 0.200) | 0.913 (0.750 to 1.000) | 0.349 (0.141 to 0.402) |
| Great tit, across year | AvesEcho | 0.388 (0.225 to 0.550) | 0.559 (0.351 to 0.700) | 0.062 (0.000 to 0.325) | 0.100 (0.000 to 0.650) | 0.062 (0.000 to 0.325) | 0.900 (0.350 to 1.000) | 0.223 (0.000 to 0.406) |
| Great tit, across year | AudioProtoPNet | 0.463 (0.350 to 0.600) | 0.535 (0.454 to 0.614) | 0.075 (0.025 to 0.175) | 0.087 (0.025 to 0.350) | 0.075 (0.025 to 0.175) | 0.913 (0.650 to 0.975) | 0.267 (0.135 to 0.391) |
| Great tit, across year | ConvNeXt (BirdSet) | 0.450 (0.275 to 0.650) | 0.523 (0.408 to 0.654) | 0.062 (0.000 to 0.150) | 0.087 (0.000 to 0.375) | 0.062 (0.000 to 0.150) | 0.913 (0.625 to 1.000) | 0.240 (0.000 to 0.387) |
| Great tit, across year | Bird-MAE | 0.375 (0.200 to 0.575) | 0.554 (0.342 to 0.667) | 0.100 (0.000 to 0.200) | 0.075 (0.000 to 0.325) | 0.100 (0.000 to 0.200) | 0.925 (0.675 to 1.000) | 0.293 (0.000 to 0.424) |
| Great tit, across year | ProtoCLR | 0.438 (0.275 to 0.625) | 0.526 (0.367 to 0.649) | 0.050 (0.000 to 0.225) | 0.087 (0.000 to 0.350) | 0.050 (0.000 to 0.225) | 0.913 (0.650 to 1.000) | 0.211 (0.000 to 0.382) |
| Great tit, across year | RCL_FS_BSED | 0.450 (0.200 to 0.825) | 0.615 (0.410 to 0.862) | 0.087 (0.000 to 0.375) | 0.087 (0.000 to 0.650) | 0.088 (0.000 to 0.375) | 0.913 (0.350 to 1.000) | 0.266 (0.000 to 0.466) |
| Great tit, across year | BirdAVES | 0.487 (0.350 to 0.625) | 0.600 (0.506 to 0.801) | 0.225 (0.000 to 0.375) | 0.087 (0.000 to 0.425) | 0.225 (0.000 to 0.375) | 0.913 (0.575 to 1.000) | 0.439 (0.000 to 0.556) |
| Great tit, across year | AVES | 0.500 (0.250 to 0.650) | 0.597 (0.445 to 0.767) | 0.175 (0.050 to 0.300) | 0.087 (0.000 to 0.300) | 0.175 (0.050 to 0.300) | 0.913 (0.700 to 1.000) | 0.380 (0.224 to 0.512) |
| Great tit, across year | NatureBEATs | 0.375 (0.150 to 0.550) | 0.524 (0.389 to 0.664) | 0.050 (0.000 to 0.225) | 0.075 (0.000 to 0.375) | 0.050 (0.000 to 0.225) | 0.925 (0.625 to 1.000) | 0.217 (0.000 to 0.375) |
| Great tit, across year | BioLingual | 0.450 (0.300 to 0.625) | 0.542 (0.431 to 0.649) | 0.050 (0.000 to 0.200) | 0.087 (0.000 to 0.275) | 0.050 (0.000 to 0.200) | 0.912 (0.725 to 1.000) | 0.221 (0.000 to 0.394) |
| Great tit, across year | BEATs | 0.438 (0.250 to 0.625) | 0.593 (0.341 to 0.749) | 0.075 (0.000 to 0.450) | 0.087 (0.000 to 0.400) | 0.075 (0.000 to 0.450) | 0.913 (0.600 to 1.000) | 0.254 (0.000 to 0.636) |
| Great tit, across year | AudioMAE | 0.375 (0.200 to 0.550) | 0.536 (0.296 to 0.664) | 0.075 (0.000 to 0.175) | 0.087 (0.000 to 0.325) | 0.075 (0.000 to 0.175) | 0.913 (0.675 to 1.000) | 0.263 (0.000 to 0.391) |
| Great tit, across year | VGGish | 0.362 (0.200 to 0.750) | 0.542 (0.370 to 0.793) | 0.075 (0.025 to 0.200) | 0.087 (0.000 to 0.250) | 0.075 (0.025 to 0.200) | 0.913 (0.750 to 1.000) | 0.260 (0.139 to 0.424) |
| Great tit, across year | ECAPA-TDNN (speaker) | 0.438 (0.175 to 0.575) | 0.528 (0.415 to 0.680) | 0.100 (0.000 to 0.225) | 0.087 (0.025 to 0.200) | 0.100 (0.000 to 0.225) | 0.913 (0.800 to 0.975) | 0.292 (0.000 to 0.431) |
| Great tit, across year | ResNet (speaker) | 0.450 (0.200 to 0.625) | 0.537 (0.412 to 0.671) | 0.025 (0.000 to 0.350) | 0.087 (0.000 to 0.400) | 0.025 (0.000 to 0.350) | 0.913 (0.600 to 1.000) | 0.155 (0.000 to 0.458) |

| Dataset | Embedding | Names correctly | Separation | Accepted and named, calibrated | Strangers accepted, calibrated | Balanced accuracy, known | Balanced accuracy, strangers | Geometric mean |
|---|---|---|---|---|---|---|---|---|
| Great tit, across year | x-vector (speaker) | 0.388 (0.175 to 0.550) | 0.562 (0.394 to 0.731) | 0.050 (0.000 to 0.125) | 0.087 (0.000 to 0.300) | 0.050 (0.000 to 0.125) | 0.913 (0.700 to 1.000) | 0.213 (0.000 to 0.304) |
| Great tit, across year | wav2vec 2.0 Base | 0.425 (0.225 to 0.575) | 0.603 (0.379 to 0.861) | 0.100 (0.000 to 0.250) | 0.075 (0.000 to 0.525) | 0.100 (0.000 to 0.250) | 0.925 (0.475 to 1.000) | 0.304 (0.000 to 0.454) |
| Great tit, across year | wav2vec 2.0 Large (robust) | 0.400 (0.200 to 0.700) | 0.625 (0.409 to 0.828) | 0.075 (0.000 to 0.225) | 0.087 (0.000 to 0.250) | 0.075 (0.000 to 0.225) | 0.913 (0.750 to 1.000) | 0.260 (0.000 to 0.462) |
| Great tit, across year | XLS-R 300M | 0.388 (0.225 to 0.675) | 0.578 (0.401 to 0.863) | 0.100 (0.000 to 0.225) | 0.087 (0.000 to 0.225) | 0.100 (0.000 to 0.225) | 0.913 (0.775 to 1.000) | 0.300 (0.000 to 0.450) |
| Great tit, across year | wav2vec 2.0 Conformer Large | 0.450 (0.250 to 0.775) | 0.660 (0.482 to 0.873) | 0.125 (0.000 to 0.300) | 0.087 (0.000 to 0.475) | 0.125 (0.000 to 0.300) | 0.913 (0.525 to 1.000) | 0.344 (0.000 to 0.518) |
| Great tit, across year | MMS 300M | 0.388 (0.275 to 0.750) | 0.606 (0.448 to 0.865) | 0.125 (0.000 to 0.200) | 0.100 (0.000 to 0.200) | 0.125 (0.000 to 0.200) | 0.900 (0.800 to 1.000) | 0.321 (0.000 to 0.430) |
| Great tit, across year | HuBERT Base | 0.450 (0.250 to 0.650) | 0.658 (0.356 to 0.908) | 0.100 (0.000 to 0.350) | 0.087 (0.000 to 0.550) | 0.100 (0.000 to 0.350) | 0.913 (0.450 to 1.000) | 0.277 (0.000 to 0.569) |
| Great tit, across year | HuBERT Large | 0.500 (0.325 to 0.825) | 0.664 (0.422 to 0.860) | 0.125 (0.050 to 0.275) | 0.087 (0.000 to 0.425) | 0.125 (0.050 to 0.275) | 0.913 (0.575 to 1.000) | 0.342 (0.221 to 0.504) |
| Great tit, across year | data2vec Base (100 h) | 0.463 (0.275 to 0.850) | 0.657 (0.437 to 0.770) | 0.113 (0.000 to 0.350) | 0.100 (0.000 to 0.500) | 0.113 (0.000 to 0.350) | 0.900 (0.500 to 1.000) | 0.311 (0.000 to 0.495) |
| Great tit, across year | data2vec Base (960 h) | 0.475 (0.300 to 0.825) | 0.637 (0.436 to 0.772) | 0.113 (0.000 to 0.400) | 0.087 (0.000 to 0.500) | 0.113 (0.000 to 0.400) | 0.913 (0.500 to 1.000) | 0.316 (0.000 to 0.520) |
| Great tit, across year | WavLM Base+ | 0.450 (0.250 to 0.600) | 0.650 (0.372 to 0.879) | 0.138 (0.000 to 0.350) | 0.087 (0.000 to 0.575) | 0.138 (0.000 to 0.350) | 0.913 (0.425 to 1.000) | 0.314 (0.000 to 0.561) |
| Great tit, across year | WavLM Large | 0.475 (0.250 to 0.775) | 0.638 (0.458 to 0.885) | 0.125 (0.025 to 0.375) | 0.087 (0.000 to 0.375) | 0.125 (0.025 to 0.375) | 0.913 (0.625 to 1.000) | 0.345 (0.152 to 0.556) |
| Great tit, across year | UniSpeech-SAT Base+ | 0.425 (0.275 to 0.575) | 0.659 (0.359 to 0.888) | 0.138 (0.000 to 0.400) | 0.100 (0.000 to 0.550) | 0.138 (0.000 to 0.400) | 0.900 (0.450 to 1.000) | 0.322 (0.000 to 0.616) |
| Great tit, across year | XEUS | 0.425 (0.200 to 0.700) | 0.638 (0.472 to 0.843) | 0.100 (0.000 to 0.350) | 0.087 (0.000 to 0.450) | 0.100 (0.000 to 0.350) | 0.912 (0.550 to 1.000) | 0.310 (0.000 to 0.545) |
| Great tit, 50 birds, across year | BirdNET v2.4 | 0.348 (0.275 to 0.450) | 0.545 (0.414 to 0.740) | 0.092 (0.015 to 0.177) | 0.092 (0.025 to 0.275) | 0.092 (0.015 to 0.177) | 0.908 (0.725 to 0.975) | 0.288 (0.116 to 0.395) |
| Great tit, 50 birds, across year | Perch 2.0 | 0.436 (0.223 to 0.508) | 0.549 (0.430 to 0.718) | 0.112 (0.015 to 0.250) | 0.103 (0.025 to 0.277) | 0.112 (0.015 to 0.250) | 0.897 (0.723 to 0.975) | 0.317 (0.121 to 0.462) |
| Great tit, 50 birds, across year | Perch | 0.204 (0.131 to 0.285) | 0.525 (0.391 to 0.616) | 0.023 (0.008 to 0.046) | 0.092 (0.042 to 0.208) | 0.023 (0.008 to 0.046) | 0.908 (0.792 to 0.958) | 0.147 (0.084 to 0.207) |
| Great tit, 50 birds, across year | SurfPerch | 0.308 (0.177 to 0.450) | 0.552 (0.425 to 0.641) | 0.080 (0.023 to 0.192) | 0.092 (0.017 to 0.262) | 0.080 (0.023 to 0.192) | 0.908 (0.738 to 0.983) | 0.272 (0.140 to 0.392) |
| Great tit, 50 birds, across year | BirdNET v3 (preview) | 0.144 (0.092 to 0.223) | 0.509 (0.418 to 0.596) | 0.004 (0.000 to 0.031) | 0.096 (0.015 to 0.217) | 0.004 (0.000 to 0.031) | 0.904 (0.783 to 0.985) | 0.042 (0.000 to 0.160) |
| Great tit, 50 birds, across year | AVEX sl-BEATs | 0.192 (0.100 to 0.333) | 0.506 (0.387 to 0.629) | 0.016 (0.000 to 0.054) | 0.088 (0.000 to 0.315) | 0.016 (0.000 to 0.054) | 0.912 (0.685 to 1.000) | 0.120 (0.000 to 0.208) |
| Great tit, 50 birds, across year | AVEX EfficientNet-B0 | 0.332 (0.177 to 0.483) | 0.555 (0.438 to 0.638) | 0.068 (0.025 to 0.183) | 0.096 (0.050 to 0.169) | 0.068 (0.025 to 0.183) | 0.904 (0.831 to 0.950) | 0.250 (0.154 to 0.393) |
| Great tit, 50 birds, across year | AvesEcho | 0.168 (0.100 to 0.275) | 0.510 (0.440 to 0.637) | 0.023 (0.008 to 0.092) | 0.104 (0.000 to 0.154) | 0.023 (0.008 to 0.092) | 0.896 (0.846 to 1.000) | 0.143 (0.081 to 0.280) |
| Great tit, 50 birds, across year | AudioProtoPNet | 0.272 (0.200 to 0.358) | 0.504 (0.362 to 0.665) | 0.033 (0.008 to 0.075) | 0.096 (0.017 to 0.192) | 0.033 (0.008 to 0.075) | 0.904 (0.808 to 0.983) | 0.176 (0.084 to 0.258) |

| Dataset | Embedding | Names correctly | Separation | Accepted and named, calibrated | Strangers accepted, calibrated | Balanced accuracy, known | Balanced accuracy, strangers | Geometric mean |
|---|---|---|---|---|---|---|---|---|
| Great tit, 50 birds, across year | ConvNeXt (BirdSet) | 0.260 (0.200 to 0.325) | 0.498 (0.410 to 0.612) | 0.028 (0.008 to 0.067) | 0.096 (0.067 to 0.177) | 0.028 (0.008 to 0.067) | 0.904 (0.823 to 0.933) | 0.155 (0.084 to 0.242) |
| Great tit, 50 birds, across year | Bird-MAE | 0.154 (0.092 to 0.258) | 0.518 (0.411 to 0.595) | 0.015 (0.000 to 0.046) | 0.096 (0.008 to 0.231) | 0.015 (0.000 to 0.046) | 0.904 (0.769 to 0.992) | 0.114 (0.000 to 0.189) |
| Great tit, 50 birds, across year | ProtoCLR | 0.162 (0.069 to 0.238) | 0.507 (0.370 to 0.619) | 0.012 (0.000 to 0.046) | 0.096 (0.033 to 0.338) | 0.012 (0.000 to 0.046) | 0.904 (0.662 to 0.967) | 0.100 (0.000 to 0.194) |
| Great tit, 50 birds, across year | RCL_FS_BSED | 0.254 (0.158 to 0.392) | 0.520 (0.449 to 0.670) | 0.044 (0.008 to 0.125) | 0.096 (0.025 to 0.262) | 0.044 (0.008 to 0.125) | 0.904 (0.738 to 0.975) | 0.194 (0.085 to 0.329) |
| Great tit, 50 birds, across year | BirdAVES | 0.240 (0.162 to 0.325) | 0.559 (0.467 to 0.662) | 0.076 (0.008 to 0.125) | 0.096 (0.042 to 0.215) | 0.076 (0.008 to 0.125) | 0.904 (0.785 to 0.958) | 0.263 (0.089 to 0.321) |
| Great tit, 50 birds, across year | AVES | 0.212 (0.138 to 0.250) | 0.531 (0.419 to 0.676) | 0.050 (0.015 to 0.085) | 0.096 (0.025 to 0.215) | 0.050 (0.015 to 0.085) | 0.904 (0.785 to 0.975) | 0.214 (0.120 to 0.267) |
| Great tit, 50 birds, across year | NatureBEATs | 0.196 (0.123 to 0.292) | 0.504 (0.379 to 0.632) | 0.015 (0.008 to 0.023) | 0.096 (0.000 to 0.300) | 0.015 (0.008 to 0.023) | 0.904 (0.700 to 1.000) | 0.110 (0.080 to 0.135) |
| Great tit, 50 birds, across year | BioLingual | 0.128 (0.069 to 0.208) | 0.471 (0.418 to 0.603) | 0.008 (0.000 to 0.038) | 0.096 (0.000 to 0.385) | 0.008 (0.000 to 0.038) | 0.904 (0.615 to 1.000) | 0.086 (0.000 to 0.176) |
| Great tit, 50 birds, across year | BEATs | 0.216 (0.131 to 0.325) | 0.527 (0.456 to 0.690) | 0.040 (0.000 to 0.108) | 0.097 (0.017 to 0.331) | 0.040 (0.000 to 0.108) | 0.903 (0.669 to 0.983) | 0.189 (0.000 to 0.303) |
| Great tit, 50 birds, across year | AudioMAE | 0.124 (0.100 to 0.233) | 0.519 (0.394 to 0.594) | 0.016 (0.000 to 0.038) | 0.096 (0.017 to 0.215) | 0.016 (0.000 to 0.038) | 0.904 (0.785 to 0.983) | 0.122 (0.000 to 0.177) |
| Great tit, 50 birds, across year | VGGish | 0.168 (0.115 to 0.342) | 0.533 (0.434 to 0.662) | 0.032 (0.000 to 0.069) | 0.090 (0.031 to 0.238) | 0.032 (0.000 to 0.069) | 0.910 (0.762 to 0.969) | 0.162 (0.000 to 0.237) |
| Great tit, 50 birds, across year | ECAPA-TDNN (speaker) | 0.242 (0.123 to 0.323) | 0.513 (0.388 to 0.631) | 0.062 (0.000 to 0.092) | 0.088 (0.017 to 0.223) | 0.062 (0.000 to 0.092) | 0.912 (0.777 to 0.983) | 0.236 (0.000 to 0.283) |
| Great tit, 50 birds, across year | ResNet (speaker) | 0.232 (0.167 to 0.358) | 0.510 (0.422 to 0.657) | 0.015 (0.000 to 0.067) | 0.096 (0.000 to 0.223) | 0.015 (0.000 to 0.067) | 0.904 (0.777 to 1.000) | 0.121 (0.000 to 0.236) |
| Great tit, 50 birds, across year | x-vector (speaker) | 0.232 (0.146 to 0.375) | 0.525 (0.402 to 0.716) | 0.052 (0.008 to 0.138) | 0.092 (0.031 to 0.225) | 0.052 (0.008 to 0.138) | 0.908 (0.775 to 0.969) | 0.222 (0.082 to 0.340) |
| Great tit, 50 birds, across year | wav2vec 2.0 Base | 0.212 (0.175 to 0.342) | 0.531 (0.424 to 0.621) | 0.031 (0.008 to 0.083) | 0.099 (0.017 to 0.292) | 0.031 (0.008 to 0.083) | 0.901 (0.708 to 0.983) | 0.160 (0.086 to 0.272) |
| Great tit, 50 birds, across year | wav2vec 2.0 Large (robust) | 0.224 (0.146 to 0.275) | 0.555 (0.390 to 0.685) | 0.024 (0.000 to 0.083) | 0.093 (0.025 to 0.223) | 0.024 (0.000 to 0.083) | 0.907 (0.777 to 0.975) | 0.143 (0.000 to 0.262) |
| Great tit, 50 birds, across year | XLS-R 300M | 0.208 (0.167 to 0.317) | 0.542 (0.416 to 0.694) | 0.031 (0.008 to 0.077) | 0.097 (0.015 to 0.267) | 0.031 (0.008 to 0.077) | 0.903 (0.733 to 0.985) | 0.165 (0.087 to 0.267) |
| Great tit, 50 birds, across year | wav2vec 2.0 Conformer Large | 0.220 (0.158 to 0.338) | 0.553 (0.425 to 0.692) | 0.035 (0.008 to 0.123) | 0.092 (0.017 to 0.200) | 0.035 (0.008 to 0.123) | 0.908 (0.800 to 0.983) | 0.174 (0.089 to 0.333) |
| Great tit, 50 birds, across year | MMS 300M | 0.225 (0.169 to 0.315) | 0.556 (0.408 to 0.689) | 0.044 (0.015 to 0.131) | 0.096 (0.008 to 0.283) | 0.044 (0.015 to 0.131) | 0.904 (0.717 to 0.992) | 0.196 (0.111 to 0.307) |
| Great tit, 50 birds, across year | HuBERT Base | 0.232 (0.125 to 0.292) | 0.535 (0.417 to 0.672) | 0.028 (0.000 to 0.083) | 0.092 (0.008 to 0.285) | 0.028 (0.000 to 0.083) | 0.908 (0.715 to 0.992) | 0.158 (0.000 to 0.254) |
| Great tit, 50 birds, across year | HuBERT Large | 0.227 (0.133 to 0.300) | 0.560 (0.425 to 0.689) | 0.044 (0.000 to 0.115) | 0.096 (0.017 to 0.262) | 0.044 (0.000 to 0.115) | 0.904 (0.738 to 0.983) | 0.195 (0.000 to 0.299) |
| Great tit, 50 birds, across year | data2vec Base (100 h) | 0.250 (0.183 to 0.300) | 0.564 (0.428 to 0.662) | 0.058 (0.023 to 0.115) | 0.096 (0.008 to 0.300) | 0.058 (0.023 to 0.115) | 0.904 (0.700 to 0.992) | 0.225 (0.143 to 0.284) |
| Great tit, 50 birds, across year | data2vec Base (960 h) | 0.258 (0.175 to 0.308) | 0.572 (0.429 to 0.671) | 0.062 (0.008 to 0.115) | 0.092 (0.008 to 0.308) | 0.062 (0.008 to 0.115) | 0.908 (0.692 to 0.992) | 0.230 (0.085 to 0.300) |

| **Dataset** | **Embedding** | **Names correctly** | **Separation** | **Accepted and named, calibrated** | **Strangers accepted, calibrated** | **Balanced accuracy, known** | **Balanced accuracy, strangers** | **Geometric mean** |
|---|---|---|---|---|---|---|---|---|
| Great tit, 50 birds, across year | WavLM Base+ | 0.208 (0.133 to 0.315) | 0.536 (0.383 to 0.668) | 0.036 (0.000 to 0.083) | 0.099 (0.000 to 0.246) | 0.036 (0.000 to 0.083) | 0.901 (0.754 to 1.000) | 0.179 (0.000 to 0.272) |
| Great tit, 50 birds, across year | WavLM Large | 0.216 (0.150 to 0.300) | 0.545 (0.386 to 0.691) | 0.036 (0.000 to 0.077) | 0.096 (0.017 to 0.215) | 0.036 (0.000 to 0.077) | 0.904 (0.785 to 0.983) | 0.183 (0.000 to 0.246) |
| Great tit, 50 birds, across year | UniSpeech-SAT Base+ | 0.204 (0.108 to 0.285) | 0.531 (0.412 to 0.668) | 0.025 (0.000 to 0.067) | 0.099 (0.008 to 0.262) | 0.025 (0.000 to 0.067) | 0.901 (0.738 to 0.992) | 0.146 (0.000 to 0.243) |
| Great tit, 50 birds, across year | XEUS | 0.192 (0.117 to 0.331) | 0.543 (0.431 to 0.719) | 0.040 (0.000 to 0.083) | 0.100 (0.025 to 0.254) | 0.040 (0.000 to 0.083) | 0.900 (0.746 to 0.975) | 0.183 (0.000 to 0.257) |
| Little owl, across year | BirdNET v2.4 | 0.659 (0.480 to 0.892) | 0.691 (0.436 to 0.881) | 0.200 (0.000 to 0.520) | 0.084 (0.000 to 0.697) | 0.213 (0.000 to 0.538) | 0.894 (0.294 to 1.000) | 0.388 (0.000 to 0.525) |
| Little owl, across year | Perch 2.0 | 0.751 (0.500 to 0.957) | 0.743 (0.468 to 0.880) | 0.374 (0.000 to 0.500) | 0.100 (0.000 to 0.587) | 0.367 (0.000 to 0.533) | 0.918 (0.422 to 1.000) | 0.481 (0.000 to 0.614) |
| Little owl, across year | Perch | 0.742 (0.500 to 0.968) | 0.759 (0.532 to 0.890) | 0.322 (0.000 to 0.649) | 0.091 (0.000 to 0.688) | 0.340 (0.000 to 0.615) | 0.910 (0.317 to 1.000) | 0.497 (0.000 to 0.607) |
| Little owl, across year | SurfPerch | 0.598 (0.337 to 0.902) | 0.619 (0.501 to 0.837) | 0.197 (0.000 to 0.431) | 0.080 (0.000 to 0.447) | 0.201 (0.000 to 0.502) | 0.923 (0.555 to 1.000) | 0.412 (0.000 to 0.553) |
| Little owl, across year | BirdNET v3 (preview) | 0.423 (0.275 to 0.771) | 0.593 (0.279 to 0.827) | 0.125 (0.000 to 0.276) | 0.085 (0.000 to 0.660) | 0.156 (0.000 to 0.324) | 0.921 (0.319 to 1.000) | 0.320 (0.000 to 0.489) |
| Little owl, across year | AVEX sl-BEATs | 0.519 (0.354 to 0.800) | 0.584 (0.358 to 0.817) | 0.107 (0.000 to 0.360) | 0.086 (0.000 to 0.670) | 0.098 (0.000 to 0.388) | 0.917 (0.317 to 1.000) | 0.264 (0.000 to 0.441) |
| Little owl, across year | AVEX EfficientNet-B0 | 0.614 (0.429 to 0.849) | 0.665 (0.408 to 0.816) | 0.194 (0.000 to 0.459) | 0.091 (0.000 to 0.523) | 0.219 (0.000 to 0.436) | 0.930 (0.480 to 1.000) | 0.372 (0.000 to 0.581) |
| Little owl, across year | AvesEcho | 0.449 (0.255 to 0.567) | 0.514 (0.381 to 0.817) | 0.071 (0.000 to 0.174) | 0.096 (0.022 to 0.367) | 0.072 (0.000 to 0.192) | 0.898 (0.621 to 0.977) | 0.245 (0.000 to 0.417) |
| Little owl, across year | AudioProtoPNet | 0.618 (0.469 to 0.926) | 0.698 (0.376 to 0.904) | 0.328 (0.000 to 0.713) | 0.090 (0.000 to 0.578) | 0.321 (0.000 to 0.633) | 0.914 (0.451 to 1.000) | 0.463 (0.000 to 0.626) |
| Little owl, across year | ConvNeXt (BirdSet) | 0.777 (0.520 to 0.925) | 0.741 (0.412 to 0.925) | 0.284 (0.000 to 0.585) | 0.100 (0.000 to 0.664) | 0.275 (0.000 to 0.568) | 0.885 (0.293 to 1.000) | 0.418 (0.000 to 0.601) |
| Little owl, across year | Bird-MAE | 0.631 (0.245 to 0.850) | 0.692 (0.447 to 0.816) | 0.166 (0.000 to 0.542) | 0.086 (0.000 to 0.679) | 0.152 (0.000 to 0.557) | 0.917 (0.304 to 1.000) | 0.320 (0.000 to 0.596) |
| Little owl, across year | ProtoCLR | 0.588 (0.357 to 0.950) | 0.578 (0.419 to 0.810) | 0.106 (0.010 to 0.330) | 0.107 (0.000 to 0.431) | 0.100 (0.010 to 0.327) | 0.897 (0.538 to 1.000) | 0.299 (0.100 to 0.457) |
| Little owl, across year | RCL_FS_BSED | 0.404 (0.229 to 0.657) | 0.545 (0.401 to 0.814) | 0.052 (0.000 to 0.153) | 0.104 (0.000 to 0.367) | 0.056 (0.000 to 0.149) | 0.887 (0.638 to 1.000) | 0.231 (0.000 to 0.345) |
| Little owl, across year | BirdAVES | 0.474 (0.284 to 0.608) | 0.580 (0.215 to 0.949) | 0.096 (0.000 to 0.341) | 0.110 (0.000 to 0.670) | 0.117 (0.000 to 0.362) | 0.903 (0.324 to 1.000) | 0.281 (0.000 to 0.510) |
| Little owl, across year | AVES | 0.525 (0.202 to 0.703) | 0.595 (0.267 to 0.813) | 0.099 (0.000 to 0.383) | 0.088 (0.000 to 0.553) | 0.104 (0.000 to 0.398) | 0.904 (0.453 to 1.000) | 0.274 (0.000 to 0.492) |
| Little owl, across year | NatureBEATs | 0.544 (0.357 to 0.706) | 0.611 (0.396 to 0.864) | 0.071 (0.000 to 0.470) | 0.086 (0.000 to 0.714) | 0.071 (0.000 to 0.523) | 0.907 (0.316 to 1.000) | 0.263 (0.000 to 0.472) |
| Little owl, across year | BioLingual | 0.534 (0.286 to 0.840) | 0.530 (0.392 to 0.757) | 0.071 (0.000 to 0.360) | 0.091 (0.000 to 0.358) | 0.061 (0.000 to 0.392) | 0.913 (0.631 to 1.000) | 0.226 (0.000 to 0.524) |
| Little owl, across year | BEATs | 0.481 (0.380 to 0.810) | 0.597 (0.345 to 0.850) | 0.097 (0.000 to 0.490) | 0.096 (0.000 to 0.765) | 0.088 (0.000 to 0.553) | 0.914 (0.240 to 1.000) | 0.249 (0.000 to 0.505) |
| Little owl, across year | AudioMAE | 0.579 (0.276 to 0.775) | 0.610 (0.362 to 0.795) | 0.095 (0.000 to 0.250) | 0.086 (0.000 to 0.495) | 0.085 (0.000 to 0.291) | 0.921 (0.494 to 1.000) | 0.279 (0.000 to 0.427) |

| Dataset | Embedding | Names correctly | Separation | Accepted and named, calibrated | Strangers accepted, calibrated | Balanced accuracy, known | Balanced accuracy, strangers | Geometric mean |
|---|---|---|---|---|---|---|---|---|
| Little owl, across year | VGGish | 0.375 (0.153 to 0.784) | 0.554 (0.242 to 0.771) | 0.070 (0.000 to 0.160) | 0.083 (0.000 to 0.442) | 0.071 (0.000 to 0.171) | 0.932 (0.515 to 1.000) | 0.233 (0.000 to 0.361) |
| Little owl, across year | ECAPA-TDNN (speaker) | 0.559 (0.286 to 0.882) | 0.660 (0.393 to 0.830) | 0.210 (0.024 to 0.392) | 0.091 (0.000 to 0.385) | 0.184 (0.022 to 0.434) | 0.904 (0.579 to 1.000) | 0.389 (0.143 to 0.592) |
| Little owl, across year | ResNet (speaker) | 0.599 (0.424 to 0.951) | 0.646 (0.488 to 0.789) | 0.126 (0.010 to 0.417) | 0.093 (0.000 to 0.495) | 0.124 (0.009 to 0.441) | 0.903 (0.477 to 1.000) | 0.347 (0.096 to 0.504) |
| Little owl, across year | x-vector (speaker) | 0.569 (0.353 to 0.784) | 0.575 (0.383 to 0.814) | 0.109 (0.010 to 0.385) | 0.105 (0.000 to 0.565) | 0.100 (0.008 to 0.376) | 0.900 (0.429 to 1.000) | 0.304 (0.087 to 0.484) |
| Little owl, across year | wav2vec 2.0 Base | 0.450 (0.240 to 0.863) | 0.560 (0.158 to 0.833) | 0.029 (0.000 to 0.324) | 0.090 (0.000 to 0.638) | 0.025 (0.000 to 0.316) | 0.906 (0.304 to 1.000) | 0.151 (0.000 to 0.481) |
| Little owl, across year | wav2vec 2.0 Large (robust) | 0.515 (0.255 to 0.843) | 0.578 (0.325 to 0.800) | 0.020 (0.000 to 0.233) | 0.102 (0.000 to 0.585) | 0.021 (0.000 to 0.235) | 0.879 (0.413 to 1.000) | 0.134 (0.000 to 0.378) |
| Little owl, across year | XLS-R 300M | 0.525 (0.276 to 0.814) | 0.555 (0.307 to 0.769) | 0.038 (0.000 to 0.402) | 0.129 (0.000 to 0.691) | 0.037 (0.000 to 0.389) | 0.845 (0.286 to 1.000) | 0.155 (0.000 to 0.373) |
| Little owl, across year | wav2vec 2.0 Conformer Large | 0.523 (0.459 to 0.896) | 0.607 (0.386 to 0.836) | 0.061 (0.000 to 0.245) | 0.086 (0.000 to 0.394) | 0.059 (0.000 to 0.238) | 0.898 (0.542 to 1.000) | 0.226 (0.000 to 0.413) |
| Little owl, across year | MMS 300M | 0.548 (0.235 to 0.863) | 0.563 (0.332 to 0.810) | 0.038 (0.000 to 0.324) | 0.126 (0.000 to 0.617) | 0.038 (0.000 to 0.295) | 0.853 (0.344 to 1.000) | 0.163 (0.000 to 0.386) |
| Little owl, across year | HuBERT Base | 0.527 (0.237 to 0.900) | 0.593 (0.348 to 0.835) | 0.061 (0.000 to 0.239) | 0.096 (0.000 to 0.426) | 0.072 (0.000 to 0.240) | 0.882 (0.464 to 1.000) | 0.238 (0.000 to 0.448) |
| Little owl, across year | HuBERT Large | 0.633 (0.220 to 0.858) | 0.569 (0.418 to 0.811) | 0.010 (0.000 to 0.533) | 0.086 (0.000 to 0.643) | 0.013 (0.000 to 0.537) | 0.917 (0.354 to 1.000) | 0.107 (0.000 to 0.500) |
| Little owl, across year | data2vec Base (100 h) | 0.562 (0.431 to 0.906) | 0.611 (0.362 to 0.810) | 0.068 (0.000 to 0.458) | 0.090 (0.000 to 0.659) | 0.087 (0.000 to 0.452) | 0.910 (0.348 to 1.000) | 0.256 (0.000 to 0.479) |
| Little owl, across year | data2vec Base (960 h) | 0.589 (0.479 to 0.917) | 0.614 (0.385 to 0.803) | 0.128 (0.000 to 0.450) | 0.098 (0.000 to 0.565) | 0.130 (0.000 to 0.451) | 0.907 (0.438 to 1.000) | 0.343 (0.000 to 0.451) |
| Little owl, across year | WavLM Base+ | 0.537 (0.247 to 0.730) | 0.587 (0.308 to 0.786) | 0.048 (0.000 to 0.206) | 0.106 (0.000 to 0.532) | 0.049 (0.000 to 0.226) | 0.862 (0.398 to 1.000) | 0.200 (0.000 to 0.402) |
| Little owl, across year | WavLM Large | 0.514 (0.347 to 0.825) | 0.558 (0.289 to 0.794) | 0.047 (0.000 to 0.363) | 0.086 (0.000 to 0.489) | 0.047 (0.000 to 0.332) | 0.903 (0.470 to 1.000) | 0.185 (0.000 to 0.405) |
| Little owl, across year | UniSpeech-SAT Base+ | 0.531 (0.269 to 0.930) | 0.576 (0.409 to 0.794) | 0.061 (0.000 to 0.193) | 0.091 (0.000 to 0.477) | 0.062 (0.000 to 0.246) | 0.891 (0.499 to 1.000) | 0.241 (0.000 to 0.406) |
| Little owl, across year | XEUS | 0.565 (0.172 to 0.833) | 0.579 (0.290 to 0.804) | 0.045 (0.000 to 0.353) | 0.096 (0.000 to 0.755) | 0.045 (0.000 to 0.333) | 0.913 (0.221 to 1.000) | 0.200 (0.000 to 0.428) |
| Rook, across year | BirdNET v2.4 | 0.611 (0.429 to 0.879) | 0.523 (0.408 to 0.679) | 0.080 (0.000 to 0.234) | 0.100 (0.000 to 0.503) | 0.064 (0.000 to 0.224) | 0.867 (0.413 to 1.000) | 0.216 (0.000 to 0.421) |
| Rook, across year | Perch 2.0 | 0.575 (0.401 to 0.842) | 0.610 (0.417 to 0.904) | 0.101 (0.000 to 0.291) | 0.100 (0.000 to 0.403) | 0.094 (0.000 to 0.373) | 0.876 (0.475 to 1.000) | 0.293 (0.000 to 0.550) |
| Rook, across year | Perch | 0.523 (0.367 to 0.765) | 0.570 (0.373 to 0.702) | 0.036 (0.000 to 0.165) | 0.102 (0.000 to 0.494) | 0.023 (0.000 to 0.154) | 0.880 (0.471 to 1.000) | 0.130 (0.000 to 0.361) |
| Rook, across year | SurfPerch | 0.500 (0.310 to 0.821) | 0.557 (0.356 to 0.707) | 0.049 (0.004 to 0.125) | 0.098 (0.003 to 0.437) | 0.054 (0.010 to 0.142) | 0.897 (0.504 to 0.997) | 0.216 (0.101 to 0.329) |
| Rook, across year | BirdNET v3 (preview) | 0.504 (0.281 to 0.791) | 0.578 (0.365 to 0.861) | 0.071 (0.000 to 0.282) | 0.100 (0.000 to 0.407) | 0.063 (0.000 to 0.344) | 0.877 (0.459 to 1.000) | 0.239 (0.000 to 0.494) |
| Rook, across year | AVEX sl-BEATs | 0.552 (0.361 to 0.835) | 0.570 (0.332 to 0.737) | 0.119 (0.001 to 0.451) | 0.095 (0.004 to 0.270) | 0.140 (0.000 to 0.395) | 0.891 (0.695 to 0.996) | 0.353 (0.022 to 0.583) |

| Dataset | Embedding | Names correctly | Separation | Accepted and named, calibrated | Strangers accepted, calibrated | Balanced accuracy, known | Balanced accuracy, strangers | Geometric mean |
|---|---|---|---|---|---|---|---|---|
| Rook, across year | AVEX EfficientNet-B0 | 0.540 (0.407 to 0.828) | 0.582 (0.397 to 0.890) | 0.064 (0.001 to 0.211) | 0.100 (0.000 to 0.383) | 0.043 (0.000 to 0.261) | 0.881 (0.521 to 1.000) | 0.188 (0.022 to 0.414) |
| Rook, across year | AvesEcho | 0.571 (0.326 to 0.816) | 0.560 (0.355 to 0.755) | 0.060 (0.000 to 0.228) | 0.093 (0.000 to 0.679) | 0.045 (0.000 to 0.156) | 0.888 (0.320 to 1.000) | 0.192 (0.000 to 0.329) |
| Rook, across year | AudioProtoPNet | 0.574 (0.457 to 0.805) | 0.570 (0.382 to 0.849) | 0.064 (0.000 to 0.222) | 0.099 (0.000 to 0.508) | 0.048 (0.000 to 0.213) | 0.864 (0.448 to 1.000) | 0.183 (0.000 to 0.420) |
| Rook, across year | ConvNeXt (BirdSet) | 0.591 (0.444 to 0.889) | 0.602 (0.412 to 0.879) | 0.077 (0.000 to 0.279) | 0.099 (0.000 to 0.352) | 0.053 (0.000 to 0.341) | 0.883 (0.532 to 1.000) | 0.189 (0.000 to 0.487) |
| Rook, across year | Bird-MAE | 0.469 (0.293 to 0.771) | 0.582 (0.382 to 0.682) | 0.073 (0.003 to 0.382) | 0.100 (0.003 to 0.276) | 0.113 (0.002 to 0.391) | 0.921 (0.709 to 0.999) | 0.322 (0.044 to 0.532) |
| Rook, across year | ProtoCLR | 0.471 (0.256 to 0.881) | 0.575 (0.348 to 0.739) | 0.078 (0.000 to 0.440) | 0.099 (0.000 to 0.255) | 0.112 (0.000 to 0.410) | 0.912 (0.723 to 1.000) | 0.315 (0.000 to 0.544) |
| Rook, across year | RCL_FS_BSED | 0.394 (0.225 to 0.859) | 0.504 (0.341 to 0.727) | 0.061 (0.001 to 0.522) | 0.097 (0.001 to 0.280) | 0.103 (0.000 to 0.573) | 0.929 (0.695 to 1.000) | 0.294 (0.021 to 0.631) |
| Rook, across year | BirdAVES | 0.502 (0.156 to 0.764) | 0.525 (0.383 to 0.786) | 0.069 (0.018 to 0.428) | 0.099 (0.016 to 0.293) | 0.074 (0.012 to 0.411) | 0.914 (0.730 to 0.992) | 0.258 (0.107 to 0.623) |
| Rook, across year | AVES | 0.494 (0.172 to 0.764) | 0.534 (0.381 to 0.804) | 0.105 (0.023 to 0.331) | 0.097 (0.017 to 0.459) | 0.074 (0.014 to 0.365) | 0.906 (0.572 to 0.983) | 0.239 (0.117 to 0.598) |
| Rook, across year | NatureBEATs | 0.457 (0.299 to 0.867) | 0.619 (0.312 to 0.778) | 0.110 (0.016 to 0.491) | 0.098 (0.007 to 0.382) | 0.136 (0.012 to 0.416) | 0.905 (0.609 to 0.996) | 0.344 (0.105 to 0.607) |
| Rook, across year | BioLingual | 0.585 (0.353 to 0.791) | 0.550 (0.400 to 0.727) | 0.060 (0.003 to 0.319) | 0.105 (0.000 to 0.580) | 0.048 (0.002 to 0.282) | 0.883 (0.347 to 1.000) | 0.209 (0.044 to 0.372) |
| Rook, across year | BEATs | 0.486 (0.217 to 0.762) | 0.592 (0.278 to 0.762) | 0.091 (0.015 to 0.178) | 0.096 (0.013 to 0.458) | 0.069 (0.021 to 0.161) | 0.894 (0.648 to 0.993) | 0.244 (0.141 to 0.397) |
| Rook, across year | AudioMAE | 0.442 (0.192 to 0.768) | 0.602 (0.288 to 0.698) | 0.090 (0.020 to 0.223) | 0.105 (0.024 to 0.239) | 0.082 (0.025 to 0.230) | 0.896 (0.773 to 0.979) | 0.268 (0.151 to 0.470) |
| Rook, across year | VGGish | 0.407 (0.279 to 0.742) | 0.547 (0.353 to 0.634) | 0.026 (0.007 to 0.084) | 0.099 (0.011 to 0.259) | 0.038 (0.007 to 0.097) | 0.918 (0.785 to 0.991) | 0.190 (0.083 to 0.282) |
| Rook, across year | ECAPA-TDNN (speaker) | 0.472 (0.227 to 0.794) | 0.518 (0.311 to 0.679) | 0.051 (0.000 to 0.129) | 0.099 (0.001 to 0.513) | 0.045 (0.000 to 0.093) | 0.904 (0.583 to 0.999) | 0.199 (0.000 to 0.284) |
| Rook, across year | ResNet (speaker) | 0.496 (0.273 to 0.901) | 0.537 (0.363 to 0.757) | 0.081 (0.020 to 0.199) | 0.097 (0.012 to 0.279) | 0.076 (0.017 to 0.203) | 0.918 (0.704 to 0.986) | 0.264 (0.129 to 0.444) |
| Rook, across year | x-vector (speaker) | 0.582 (0.147 to 0.847) | 0.499 (0.371 to 0.735) | 0.065 (0.002 to 0.329) | 0.100 (0.007 to 0.338) | 0.069 (0.001 to 0.297) | 0.891 (0.639 to 0.989) | 0.248 (0.030 to 0.528) |
| Rook, across year | wav2vec 2.0 Base | 0.385 (0.281 to 0.780) | 0.547 (0.329 to 0.695) | 0.040 (0.000 to 0.136) | 0.105 (0.005 to 0.303) | 0.055 (0.000 to 0.142) | 0.904 (0.750 to 0.998) | 0.218 (0.000 to 0.327) |
| Rook, across year | wav2vec 2.0 Large (robust) | 0.380 (0.205 to 0.776) | 0.531 (0.400 to 0.584) | 0.032 (0.012 to 0.104) | 0.098 (0.031 to 0.223) | 0.030 (0.015 to 0.089) | 0.914 (0.821 to 0.970) | 0.165 (0.118 to 0.285) |
| Rook, across year | XLS-R 300M | 0.432 (0.222 to 0.751) | 0.526 (0.315 to 0.665) | 0.039 (0.018 to 0.088) | 0.098 (0.032 to 0.197) | 0.050 (0.015 to 0.098) | 0.913 (0.848 to 0.975) | 0.212 (0.122 to 0.308) |
| Rook, across year | wav2vec 2.0 Conformer Large | 0.429 (0.261 to 0.716) | 0.547 (0.363 to 0.665) | 0.045 (0.003 to 0.164) | 0.100 (0.003 to 0.343) | 0.047 (0.007 to 0.161) | 0.904 (0.639 to 0.998) | 0.203 (0.080 to 0.343) |
| Rook, across year | MMS 300M | 0.441 (0.228 to 0.772) | 0.541 (0.424 to 0.633) | 0.043 (0.007 to 0.133) | 0.103 (0.020 to 0.209) | 0.040 (0.008 to 0.102) | 0.906 (0.818 to 0.967) | 0.189 (0.088 to 0.302) |
| Rook, across year | HuBERT Base | 0.371 (0.212 to 0.703) | 0.552 (0.374 to 0.729) | 0.088 (0.002 to 0.182) | 0.099 (0.028 to 0.309) | 0.058 (0.002 to 0.166) | 0.913 (0.737 to 0.985) | 0.228 (0.040 to 0.383) |

| Dataset | Embedding | Names correctly | Separation | Accepted and named, calibrated | Strangers accepted, calibrated | Balanced accuracy, known | Balanced accuracy, strangers | Geometric mean |
|---|---|---|---|---|---|---|---|---|
| Rook, across year | HuBERT Large | 0.459 (0.221 to 0.796) | 0.524 (0.336 to 0.682) | 0.043 (0.000 to 0.280) | 0.096 (0.000 to 0.504) | 0.034 (0.000 to 0.181) | 0.897 (0.610 to 1.000) | 0.177 (0.000 to 0.332) |
| Rook, across year | data2vec Base (100 h) | 0.354 (0.227 to 0.714) | 0.532 (0.408 to 0.653) | 0.053 (0.000 to 0.155) | 0.097 (0.022 to 0.271) | 0.052 (0.000 to 0.100) | 0.896 (0.739 to 0.981) | 0.210 (0.000 to 0.313) |
| Rook, across year | data2vec Base (960 h) | 0.400 (0.236 to 0.739) | 0.537 (0.349 to 0.668) | 0.033 (0.011 to 0.100) | 0.098 (0.001 to 0.333) | 0.030 (0.014 to 0.117) | 0.899 (0.651 to 0.998) | 0.161 (0.113 to 0.308) |
| Rook, across year | WavLM Base+ | 0.405 (0.162 to 0.784) | 0.495 (0.359 to 0.696) | 0.033 (0.002 to 0.348) | 0.105 (0.005 to 0.509) | 0.044 (0.001 to 0.268) | 0.917 (0.592 to 0.993) | 0.193 (0.037 to 0.459) |
| Rook, across year | WavLM Large | 0.464 (0.208 to 0.793) | 0.531 (0.329 to 0.688) | 0.042 (0.003 to 0.087) | 0.104 (0.016 to 0.222) | 0.043 (0.003 to 0.066) | 0.905 (0.796 to 0.982) | 0.194 (0.055 to 0.237) |
| Rook, across year | UniSpeech-SAT Base+ | 0.486 (0.321 to 0.797) | 0.540 (0.397 to 0.634) | 0.081 (0.012 to 0.246) | 0.099 (0.046 to 0.176) | 0.059 (0.015 to 0.315) | 0.916 (0.836 to 0.972) | 0.234 (0.119 to 0.513) |
| Rook, across year | XEUS | 0.432 (0.279 to 0.736) | 0.564 (0.344 to 0.669) | 0.064 (0.015 to 0.145) | 0.103 (0.000 to 0.475) | 0.058 (0.009 to 0.129) | 0.902 (0.632 to 1.000) | 0.226 (0.092 to 0.343) |
| Red-tailed black cockatoo, random split | BirdNET v2.4 | 0.980 (0.899 to 1.000) | 0.850 (0.632 to 0.997) | 0.741 (0.029 to 0.921) | 0.082 (0.000 to 0.584) | 0.627 (0.013 to 0.940) | 0.939 (0.530 to 1.000) | 0.718 (0.115 to 0.939) |
| Red-tailed black cockatoo, random split | Perch 2.0 | 0.968 (0.879 to 1.000) | 0.908 (0.642 to 1.000) | 0.814 (0.029 to 0.941) | 0.085 (0.000 to 0.632) | 0.757 (0.013 to 0.961) | 0.939 (0.497 to 1.000) | 0.750 (0.115 to 0.921) |
| Red-tailed black cockatoo, random split | Perch | 0.975 (0.874 to 1.000) | 0.878 (0.718 to 0.998) | 0.736 (0.318 to 0.891) | 0.100 (0.000 to 0.579) | 0.671 (0.288 to 0.920) | 0.939 (0.675 to 1.000) | 0.769 (0.494 to 0.899) |
| Red-tailed black cockatoo, random split | SurfPerch | 0.975 (0.906 to 1.000) | 0.899 (0.681 to 0.999) | 0.644 (0.118 to 1.000) | 0.089 (0.000 to 0.934) | 0.626 (0.089 to 1.000) | 0.945 (0.221 to 1.000) | 0.697 (0.292 to 0.868) |
| Red-tailed black cockatoo, random split | BirdNET v3 (preview) | 0.921 (0.752 to 1.000) | 0.828 (0.616 to 0.983) | 0.602 (0.088 to 0.880) | 0.080 (0.000 to 0.624) | 0.557 (0.099 to 0.810) | 0.943 (0.431 to 1.000) | 0.637 (0.314 to 0.851) |
| Red-tailed black cockatoo, random split | AVEX sl-BEATs | 0.820 (0.582 to 0.896) | 0.675 (0.452 to 0.960) | 0.208 (0.058 to 0.710) | 0.091 (0.000 to 0.441) | 0.192 (0.051 to 0.531) | 0.889 (0.692 to 1.000) | 0.429 (0.207 to 0.667) |
| Red-tailed black cockatoo, random split | AVEX EfficientNet-B0 | 0.968 (0.854 to 0.990) | 0.849 (0.602 to 0.983) | 0.667 (0.029 to 0.812) | 0.090 (0.000 to 0.548) | 0.620 (0.013 to 0.802) | 0.932 (0.676 to 1.000) | 0.705 (0.115 to 0.854) |
| Red-tailed black cockatoo, random split | AvesEcho | 0.964 (0.915 to 1.000) | 0.928 (0.667 to 0.988) | 0.862 (0.000 to 0.981) | 0.107 (0.000 to 0.632) | 0.826 (0.000 to 0.994) | 0.906 (0.650 to 1.000) | 0.836 (0.000 to 0.936) |
| Red-tailed black cockatoo, random split | AudioProtoPNet | 0.968 (0.792 to 1.000) | 0.846 (0.555 to 0.992) | 0.670 (0.059 to 0.901) | 0.090 (0.000 to 0.446) | 0.600 (0.049 to 0.911) | 0.949 (0.640 to 1.000) | 0.706 (0.219 to 0.930) |
| Red-tailed black cockatoo, random split | ConvNeXt (BirdSet) | 0.976 (0.819 to 1.000) | 0.912 (0.596 to 1.000) | 0.783 (0.059 to 0.970) | 0.098 (0.000 to 0.678) | 0.699 (0.049 to 0.974) | 0.928 (0.534 to 1.000) | 0.726 (0.220 to 0.960) |
| Red-tailed black cockatoo, random split | Bird-MAE | 0.941 (0.836 to 1.000) | 0.855 (0.670 to 0.996) | 0.673 (0.136 to 0.940) | 0.104 (0.000 to 0.702) | 0.705 (0.111 to 0.905) | 0.883 (0.377 to 1.000) | 0.701 (0.333 to 0.880) |
| Red-tailed black cockatoo, random split | ProtoCLR | 0.772 (0.617 to 0.907) | 0.715 (0.576 to 0.850) | 0.277 (0.059 to 0.443) | 0.090 (0.000 to 0.439) | 0.234 (0.049 to 0.463) | 0.893 (0.655 to 1.000) | 0.455 (0.220 to 0.636) |
| Red-tailed black cockatoo, random split | RCL_FS_BSED | 0.889 (0.618 to 0.985) | 0.668 (0.522 to 0.981) | 0.352 (0.000 to 0.752) | 0.094 (0.000 to 0.733) | 0.500 (0.000 to 0.839) | 0.760 (0.265 to 1.000) | 0.515 (0.000 to 0.717) |
| Red-tailed black cockatoo, random split | BirdAVES | 0.986 (0.881 to 1.000) | 0.923 (0.733 to 0.998) | 0.815 (0.136 to 0.980) | 0.092 (0.000 to 0.947) | 0.784 (0.138 to 0.986) | 0.911 (0.106 to 1.000) | 0.760 (0.320 to 0.898) |
| Red-tailed black cockatoo, random split | AVES | 0.981 (0.901 to 1.000) | 0.924 (0.778 to 1.000) | 0.848 (0.106 to 0.990) | 0.111 (0.000 to 0.719) | 0.825 (0.069 to 0.991) | 0.913 (0.482 to 1.000) | 0.777 (0.262 to 0.913) |
| Red-tailed black cockatoo, random split | NatureBEATs | 0.944 (0.860 to 1.000) | 0.903 (0.712 to 0.993) | 0.706 (0.212 to 0.947) | 0.094 (0.000 to 0.667) | 0.664 (0.148 to 0.965) | 0.935 (0.510 to 1.000) | 0.765 (0.385 to 0.885) |

| Dataset | Embedding | Names correctly | Separation | Accepted and named, calibrated | Strangers accepted, calibrated | Balanced accuracy, known | Balanced accuracy, strangers | Geometric mean |
|---|---|---|---|---|---|---|---|---|
| Red-tailed black cockatoo, random split | BioLingual | 0.779 (0.585 to 0.955) | 0.732 (0.418 to 0.974) | 0.313 (0.088 to 0.653) | 0.089 (0.000 to 0.801) | 0.332 (0.088 to 0.615) | 0.896 (0.246 to 1.000) | 0.532 (0.285 to 0.660) |
| Red-tailed black cockatoo, random split | BEATs | 0.868 (0.727 to 0.939) | 0.835 (0.568 to 0.961) | 0.501 (0.030 to 0.800) | 0.104 (0.000 to 0.723) | 0.537 (0.017 to 0.828) | 0.862 (0.290 to 1.000) | 0.600 (0.131 to 0.779) |
| Red-tailed black cockatoo, random split | AudioMAE | 0.867 (0.655 to 0.916) | 0.771 (0.476 to 0.954) | 0.429 (0.076 to 0.782) | 0.095 (0.000 to 0.485) | 0.372 (0.073 to 0.735) | 0.853 (0.519 to 1.000) | 0.539 (0.270 to 0.662) |
| Red-tailed black cockatoo, random split | VGGish | 0.849 (0.680 to 0.944) | 0.731 (0.593 to 0.939) | 0.403 (0.045 to 0.732) | 0.090 (0.000 to 0.456) | 0.489 (0.107 to 0.717) | 0.940 (0.557 to 1.000) | 0.654 (0.327 to 0.736) |
| Red-tailed black cockatoo, random split | ECAPA-TDNN (speaker) | 0.866 (0.636 to 0.970) | 0.634 (0.353 to 0.854) | 0.224 (0.053 to 0.358) | 0.090 (0.000 to 0.316) | 0.172 (0.018 to 0.367) | 0.946 (0.829 to 1.000) | 0.400 (0.124 to 0.591) |
| Red-tailed black cockatoo, random split | ResNet (speaker) | 0.773 (0.520 to 0.970) | 0.584 (0.445 to 0.784) | 0.144 (0.000 to 0.430) | 0.105 (0.000 to 0.386) | 0.124 (0.000 to 0.333) | 0.927 (0.713 to 1.000) | 0.343 (0.000 to 0.498) |
| Red-tailed black cockatoo, random split | x-vector (speaker) | 0.933 (0.758 to 0.983) | 0.752 (0.543 to 0.918) | 0.373 (0.099 to 0.649) | 0.079 (0.000 to 0.386) | 0.398 (0.062 to 0.802) | 0.944 (0.678 to 1.000) | 0.624 (0.249 to 0.863) |
| Red-tailed black cockatoo, random split | wav2vec 2.0 Base | 0.937 (0.868 to 0.990) | 0.867 (0.618 to 0.987) | 0.648 (0.015 to 0.888) | 0.113 (0.000 to 0.635) | 0.633 (0.009 to 0.815) | 0.909 (0.494 to 1.000) | 0.687 (0.093 to 0.874) |
| Red-tailed black cockatoo, random split | wav2vec 2.0 Large (robust) | 0.930 (0.800 to 0.972) | 0.839 (0.634 to 0.965) | 0.727 (0.121 to 0.860) | 0.085 (0.000 to 0.526) | 0.623 (0.072 to 0.849) | 0.935 (0.744 to 1.000) | 0.750 (0.268 to 0.883) |
| Red-tailed black cockatoo, random split | XLS-R 300M | 0.956 (0.836 to 0.985) | 0.877 (0.686 to 0.983) | 0.667 (0.182 to 0.879) | 0.085 (0.000 to 0.614) | 0.657 (0.131 to 0.836) | 0.930 (0.653 to 1.000) | 0.761 (0.360 to 0.875) |
| Red-tailed black cockatoo, random split | wav2vec 2.0 Conformer Large | 0.962 (0.800 to 0.985) | 0.874 (0.644 to 0.990) | 0.695 (0.136 to 0.940) | 0.091 (0.000 to 0.609) | 0.670 (0.100 to 0.905) | 0.954 (0.627 to 1.000) | 0.760 (0.317 to 0.832) |
| Red-tailed black cockatoo, random split | MMS 300M | 0.966 (0.800 to 0.985) | 0.882 (0.685 to 0.990) | 0.642 (0.182 to 0.880) | 0.086 (0.000 to 0.614) | 0.690 (0.131 to 0.839) | 0.915 (0.683 to 1.000) | 0.774 (0.358 to 0.855) |
| Red-tailed black cockatoo, random split | HuBERT Base | 0.970 (0.891 to 1.000) | 0.854 (0.555 to 0.992) | 0.701 (0.076 to 0.912) | 0.106 (0.000 to 0.743) | 0.681 (0.043 to 0.893) | 0.958 (0.268 to 1.000) | 0.743 (0.208 to 0.888) |
| Red-tailed black cockatoo, random split | HuBERT Large | 0.956 (0.818 to 0.985) | 0.854 (0.648 to 0.984) | 0.648 (0.106 to 0.877) | 0.111 (0.000 to 0.609) | 0.665 (0.083 to 0.864) | 0.905 (0.539 to 1.000) | 0.754 (0.288 to 0.845) |
| Red-tailed black cockatoo, random split | data2vec Base (100 h) | 0.971 (0.891 to 1.000) | 0.866 (0.577 to 0.994) | 0.759 (0.136 to 0.912) | 0.096 (0.000 to 0.456) | 0.724 (0.100 to 0.893) | 0.962 (0.552 to 1.000) | 0.815 (0.317 to 0.878) |
| Red-tailed black cockatoo, random split | data2vec Base (960 h) | 0.961 (0.862 to 1.000) | 0.855 (0.578 to 0.991) | 0.750 (0.136 to 0.912) | 0.089 (0.000 to 0.525) | 0.719 (0.100 to 0.887) | 0.962 (0.373 to 1.000) | 0.807 (0.317 to 0.887) |
| Red-tailed black cockatoo, random split | WavLM Base+ | 0.968 (0.883 to 1.000) | 0.878 (0.595 to 0.995) | 0.744 (0.045 to 0.930) | 0.092 (0.000 to 0.559) | 0.716 (0.026 to 0.923) | 0.940 (0.445 to 1.000) | 0.796 (0.161 to 0.906) |
| Red-tailed black cockatoo, random split | WavLM Large | 0.961 (0.818 to 1.000) | 0.852 (0.654 to 0.988) | 0.671 (0.121 to 0.900) | 0.124 (0.000 to 0.604) | 0.686 (0.092 to 0.893) | 0.929 (0.511 to 1.000) | 0.774 (0.302 to 0.849) |
| Red-tailed black cockatoo, random split | UniSpeech-SAT Base+ | 0.971 (0.926 to 1.000) | 0.860 (0.589 to 0.994) | 0.731 (0.045 to 0.925) | 0.083 (0.000 to 0.683) | 0.710 (0.026 to 0.901) | 0.957 (0.319 to 1.000) | 0.768 (0.161 to 0.899) |
| Red-tailed black cockatoo, random split | XEUS | 0.969 (0.873 to 1.000) | 0.880 (0.663 to 0.997) | 0.750 (0.121 to 0.947) | 0.106 (0.000 to 0.644) | 0.747 (0.069 to 0.965) | 0.945 (0.391 to 1.000) | 0.843 (0.263 to 0.884) |

*Notes: Calibrated: threshold set on other strangers to admit one in ten; strangers accepted: the proportion of test strangers it actually admitted. Balanced accuracy on known individuals: mean over individuals of the proportion of their calls accepted and correctly named; on strangers: proportion rejected. Geometric mean of the two, as in the AnimalCLEF 2025 benchmark (Adam et al. 2025). Duration and loudness controls are excluded because cosine normalisation removes their magnitude.*

**Table S16.** BirdNET's internal layers, a learned combination of them, and learned pooling over time.

| Dataset | Layer chosen by the Fisher rule | Its re-ID accuracy | Its adjusted p | Final embedding | Best layer chosen with the test labels |
|---|---|---|---|---|---|
| Zebra finch, one bird per recording | stage 4, mean | 0.933 | 0.0012 | 0.967 | embedding, mean (0.967) |
| Great tit, across year | stage 3, mean | 0.338 | 0.0012 | 0.444 | embedding, mean (0.444) |
| Little owl, across year | stage 2, mean | 0.157 | 0.1932 | 0.501 | post-convolution, mean and spread (0.516) |
| Chiffchaff, within year | stage 3, mean | 0.704 | 0.0012 | 0.765 | post-convolution, mean and spread (0.769) |
| Chiffchaff, across year | stage 3, mean | 0.170 | 1.0000 | 0.160 | stage 2, mean (0.195) |
| Tree pipit, within year | stage 1, mean | 0.139 | 1.0000 | 0.376 | embedding, mean (0.376) |
| Tree pipit, across year | stage 1, mean | 0.070 | 1.0000 | 0.214 | embedding, mean (0.214) |
| Zebra finch, group of four | stage 2, mean and spread | 0.729 | 0.0024 | 0.807 | embedding, mean (0.807) |
| Zebra finch, group of eight | stage 4, mean | 0.410 | 0.0624 | 0.498 | post-convolution, mean and spread (0.515) |

| Dataset | Embedding | Kernel ridge | Class mean | Nearest clip |
|---|---|---|---|---|
| Zebra finch, one bird per recording | final layer alone | 0.967 | 0.933 | 0.956 |
| Zebra finch, one bird per recording | final layer projected | 0.978 | 0.989 | 0.989 |
| Zebra finch, one bird per recording | layers concatenated | 0.967 | 0.933 | 0.967 |
| Zebra finch, one bird per recording | learned combination | 0.978 | 0.967 | 0.967 |
| Great tit, across year | final layer alone | 0.444 | 0.375 | 0.425 |
| Great tit, across year | final layer projected | 0.469 | 0.487 | 0.487 |
| Great tit, across year | layers concatenated | 0.381 | 0.325 | 0.362 |
| Great tit, across year | learned combination | 0.481 | 0.481 | 0.475 |
| Little owl, across year | final layer alone | 0.501 | 0.393 | 0.548 |
| Little owl, across year | final layer projected | 0.472 | 0.494 | 0.494 |
| Little owl, across year | layers concatenated | 0.467 | 0.428 | 0.558 |
| Little owl, across year | learned combination | 0.553 | 0.577 | 0.577 |
| Chiffchaff, within year | final layer alone | 0.765 | 0.549 | 0.643 |
| Chiffchaff, within year | final layer projected | 0.767 | 0.763 | 0.737 |
| Chiffchaff, within year | layers concatenated | 0.763 | 0.476 | 0.631 |
| Chiffchaff, within year | learned combination | 0.784 | 0.803 | 0.778 |
| Chiffchaff, across year | final layer alone | 0.160 | 0.125 | 0.115 |
| Chiffchaff, across year | final layer projected | 0.200 | 0.195 | 0.195 |
| Chiffchaff, across year | layers concatenated | 0.165 | 0.145 | 0.135 |
| Chiffchaff, across year | learned combination | 0.205 | 0.205 | 0.205 |
| Tree pipit, within year | final layer alone | 0.376 | 0.327 | 0.363 |
| Tree pipit, within year | final layer projected | 0.300 | 0.333 | 0.333 |
| Tree pipit, within year | layers concatenated | 0.373 | 0.261 | 0.323 |
| Tree pipit, within year | learned combination | 0.314 | 0.356 | 0.360 |
| Tree pipit, across year | final layer alone | 0.214 | 0.173 | 0.176 |
| Tree pipit, across year | final layer projected | 0.208 | 0.198 | 0.198 |
| Tree pipit, across year | layers concatenated | 0.192 | 0.144 | 0.179 |
| Tree pipit, across year | learned combination | 0.204 | 0.192 | 0.195 |

| Dataset | Layer | Individuals scored | Layer averaged over time | Untrained sequence average | Learned sequence pooling |
|---|---|---|---|---|---|
| Zebra finch, one bird per recording | post-activation, sequence | 3 | 0.967 | 0.900 | 0.900 |
| Zebra finch, one bird per recording | stage 1, sequence | 3 | 0.867 | 0.700 | 0.833 |
| Zebra finch, one bird per recording | stage 2, sequence | 3 | 0.967 | 0.733 | 0.633 |
| Zebra finch, one bird per recording | stage 3, sequence | 3 | 0.900 | 0.933 | 0.800 |
| Zebra finch, one bird per recording | stage 4, sequence | 3 | 0.967 | 0.900 | 0.900 |
| Great tit, across year | post-activation, sequence | 7 | 0.429 | 0.243 | 0.314 |
| Great tit, across year | stage 1, sequence | 7 | 0.200 | 0.229 | 0.214 |
| Great tit, across year | stage 2, sequence | 7 | 0.200 | 0.229 | 0.214 |
| Great tit, across year | stage 3, sequence | 7 | 0.343 | 0.343 | 0.386 |
| Great tit, across year | stage 4, sequence | 7 | 0.500 | 0.243 | 0.314 |
| Little owl, across year | post-activation, sequence | 7 | 0.512 | 0.351 | 0.399 |
| Little owl, across year | stage 1, sequence | 7 | 0.262 | 0.286 | 0.375 |
| Little owl, across year | stage 2, sequence | 7 | 0.440 | 0.363 | 0.196 |
| Little owl, across year | stage 3, sequence | 7 | 0.458 | 0.363 | 0.250 |
| Little owl, across year | stage 4, sequence | 7 | 0.506 | 0.375 | 0.363 |
| Chiffchaff, within year | post-activation, sequence | 6 | 0.491 | 0.403 | 0.441 |
| Chiffchaff, within year | stage 1, sequence | 6 | 0.153 | 0.213 | 0.353 |
| Chiffchaff, within year | stage 2, sequence | 6 | 0.214 | 0.345 | 0.592 |
| Chiffchaff, within year | stage 3, sequence | 6 | 0.468 | 0.592 | 0.568 |
| Chiffchaff, within year | stage 4, sequence | 6 | 0.474 | 0.427 | 0.518 |
| Egyptian fruit bat, across year | post-activation, sequence | 3 | 0.368 | 0.361 | 0.265 |
| Egyptian fruit bat, across year | stage 1, sequence | 3 | 0.345 | 0.321 | 0.292 |
| Egyptian fruit bat, across year | stage 2, sequence | 3 | 0.311 | 0.302 | 0.247 |
| Egyptian fruit bat, across year | stage 3, sequence | 3 | 0.255 | 0.303 | 0.255 |
| Egyptian fruit bat, across year | stage 4, sequence | 3 | 0.373 | 0.332 | 0.302 |

*Notes:* Upper: the 14 candidates of Appendix S1.13, with the permutation p (9,999 shuffles) multiplied by the number of distinct candidates (12, since two of the 14 repeat others) and capped at one; the best layer chosen with the test labels cannot be chosen in practice. Middle: learned combination against the final embedding alone, the same projection fitted to the final embedding, and the seven layers joined; no interval was computed on the differences. Lower: learned pooling scored on individuals it never saw (3 to 7 per dataset), so its values are not comparable in scale with Table 2.

**Table S17.** The four pairings on North Atlantic right whale upcalls and tag noise from 11 whales (data and frequency shift of Tolkova et al. 2026), with BirdNET.

| Frequency shift (Hz) | Fisher ratio on enrolment clips | C→C (accuracy / p) | B→B (accuracy / p) | C→B (accuracy / p) | B→C (accuracy / p) |
|---|---|---|---|---|---|
| 0 | 0.660 | 0.833 / 0.0001 | 0.625 / 0.0001 | 0.250 / 0.0451 | 0.292 / 0.0105 |
| 1000 | 0.516 | 0.806 / 0.0001 | 0.750 / 0.0001 | 0.333 / 0.0094 | 0.306 / 0.0046 |
| 2000 | 0.559 | 0.694 / 0.0001 | 0.611 / 0.0001 | 0.181 / 0.2611 | 0.292 / 0.0048 |
| 3000 | 0.605 | 0.681 / 0.0001 | 0.625 / 0.0001 | 0.319 / 0.0175 | 0.417 / 0.0001 |
| 4000 | 0.585 | 0.764 / 0.0001 | 0.639 / 0.0001 | 0.389 / 0.0037 | 0.458 / 0.0001 |
| 5000 | 0.534 | 0.639 / 0.0001 | 0.611 / 0.0001 | 0.264 / 0.0531 | 0.306 / 0.0033 |
| 6000 | 0.715 | 0.708 / 0.0001 | 0.597 / 0.0001 | 0.264 / 0.0445 | 0.333 / 0.0023 |
| 7000 | 0.547 | 0.681 / 0.0001 | 0.611 / 0.0001 | 0.333 / 0.0074 | 0.347 / 0.0004 |
| 8000 | 0.691 | 0.708 / 0.0001 | 0.583 / 0.0001 | 0.222 / 0.1094 | 0.361 / 0.0003 |
| 9000 | 0.639 | 0.708 / 0.0001 | 0.569 / 0.0001 | 0.264 / 0.0372 | 0.264 / 0.0347 |
| 10000 | 0.726 | 0.583 / 0.0001 | 0.556 / 0.0001 | 0.250 / 0.0551 | 0.208 / 0.1054 |

*Notes:* Each whale carried one tag deployment, so enrolment and test clips come from one recording. The shift moves the 50 to 500 Hz upcall band upward before BirdNET reads it, identically for calls and noise. The shift for the result was specified in advance as the one with the highest Fisher ratio on the enrolment calls (10,000 Hz). p from 9,999 label shuffles. After Holm's correction over the 11 shifts, B→C exceeded chance at 9 shifts and C→B at 1, and neither at 10,000 Hz. Data: github.com/avokloti/narw-acoustic-identification, commit 3cad65d, which declares no licence; no clip is redistributed. No identity claim is made from this dataset.

**Table S18.** Beecher's information statistic (bits) for each neural network at its layer chosen by the Fisher rule.

| **Neural network** | **Zebra finch, group of four** | **Zebra finch, one bird per recording** | **Zebra finch, group of eight** | **Great tit, across year** | **Tree pipit, across year** | **Chiffchaff, across year** | **Tree pipit, within year** | **Little owl, across year** | **Little penguin, across night** | **Red-tailed black cockatoo, random split** | **Egyptian fruit bat, across year** | **Chiffchaff, within year** | **Rook, across year** | **Great tit, 50 birds, across year** |
|---|---|---|---|---|---|---|---|---|---|---|---|---|---|---|
| BirdNET v2.4 | 1.12 | 7.92 | 2.10 | 2.80 | 2.08 | 3.45 | 2.34 | 7.58 | 4.01 | 5.82 | 0.81 | 4.37 | 1.19 | 8.19 |
| Perch 2.0 | 1.37 | 7.33 | 2.16 | 2.67 | 1.71 | 2.20 | 2.15 | 7.91 | 3.94 | 5.86 | 0.50 | 3.18 | 1.28 | 7.69 |
| Perch | 1.12 | 7.12 | 2.11 | 2.15 | 1.72 | 1.71 | 1.84 | 8.32 | 3.62 | 5.79 | 0.26 | 2.38 | 1.04 | 5.09 |
| SurfPerch | 1.02 | 4.68 | 1.56 | 2.16 | 1.27 | 1.67 | 1.46 | 4.97 | 3.08 | 6.17 | 0.20 | 2.24 | 0.71 | 6.17 |
| BirdNET v3 (preview) | 1.24 | 6.50 | 1.52 | 1.05 | 1.16 | 1.17 | 1.29 | 4.98 | 2.25 | 3.91 | 0.33 | 1.96 | 0.81 | 2.91 |
| AVEX sl-BEATs | 0.27 | 5.18 | 0.88 | 2.49 | 1.00 | 1.89 | 1.35 | 3.74 | 2.42 | 2.83 | 0.59 | 3.16 | 0.73 | 7.00 |
| AVEX EfficientNet-B0 | 0.99 | 5.04 | 1.62 | 1.98 | 1.11 | 1.52 | 1.42 | 5.41 | 2.65 | 4.19 | 0.55 | 2.03 | 0.89 | 5.45 |
| AvesEcho | 0.99 | 6.01 | 1.88 | 2.83 | 1.27 | 3.14 | 1.84 | 4.07 | 2.51 | 7.59 | 0.74 | 4.30 | 0.96 | 7.95 |
| AudioProtoPNet | 0.97 | 7.58 | 2.07 | 2.24 | 1.68 | 1.97 | 2.17 | 7.56 | 3.55 | 5.34 | 0.68 | 2.86 | 1.00 | 5.12 |
| ConvNeXt (BirdSet) | 1.20 | 7.48 | 2.36 | 1.78 | 2.06 | 2.25 | 2.51 | 7.92 | 4.11 | 6.07 | 0.64 | 2.85 | 1.14 | 4.68 |
| Bird-MAE | 0.94 | 4.40 | 1.33 | 2.51 | 1.02 | 1.82 | 1.60 | 5.61 | 2.96 | 5.51 | 0.58 | 3.29 | 0.50 | 7.37 |
| ProtoCLR | 0.31 | 3.20 | 1.15 | 2.53 | 0.94 | 2.05 | 1.10 | 4.87 | 2.55 | 3.48 | 0.62 | 2.33 | 0.41 | 6.16 |
| RCL_FS_BSED | 0.82 | 3.50 | 1.21 | 2.98 | 1.73 | 2.63 | 2.14 | 3.91 | 3.84 | 4.57 | 0.93 | 3.01 | 0.47 | 8.07 |
| BirdAVES | 0.63 | 2.58 | 1.38 | 2.77 | 1.64 | 2.36 | 1.82 | 4.37 | 4.04 | 6.85 | 0.89 | 2.25 | 0.55 | 6.63 |
| AVES | 0.33 | 2.81 | 1.25 | 2.72 | 1.81 | 2.69 | 2.17 | 4.51 | 4.80 | 6.83 | 0.76 | 2.61 | 0.61 | 7.08 |
| NatureBEATs | 0.89 | 3.08 | 0.94 | 1.51 | 1.08 | 2.17 | 1.34 | 3.98 | 1.94 | 6.28 | 0.54 | 3.37 | 0.64 | 6.64 |
| BioLingual | 0.90 | 6.03 | 1.62 | 2.28 | 1.16 | 1.67 | 1.37 | 3.90 | 2.40 | 3.99 | 0.40 | 3.04 | 0.98 | 3.84 |
| BEATs | 0.65 | 3.35 | 1.35 | 2.43 | 1.45 | 2.05 | 1.54 | 3.82 | 2.51 | 4.28 | 0.58 | 2.44 | 0.77 | 7.02 |
| AudioMAE | 0.87 | 4.49 | 1.53 | 2.02 | 1.06 | 1.52 | 1.35 | 4.15 | 1.90 | 3.67 | 0.69 | 2.31 | 0.63 | 4.79 |
| VGGish | 0.66 | 2.68 | 1.15 | 2.03 | 1.36 | 1.72 | 1.23 | 2.88 | 2.48 | 4.03 | 0.45 | 2.09 | 0.40 | 4.89 |
| ECAPA-TDNN (speaker) | 1.17 | 5.75 | 1.73 | 1.52 | 1.25 | 1.68 | 1.47 | 4.90 | 3.13 | 3.73 | 0.14 | 1.49 | 0.77 | 3.95 |
| ResNet (speaker) | 1.01 | 6.35 | 1.73 | 1.73 | 1.17 | 1.90 | 1.47 | 4.22 | 2.87 | 3.11 | 0.34 | 1.61 | 0.68 | 4.41 |
| x-vector (speaker) | 1.02 | 4.88 | 1.39 | 2.02 | 1.69 | 2.26 | 2.13 | 5.15 | 3.54 | 4.66 | 0.38 | 1.97 | 0.85 | 4.79 |
| wav2vec 2.0 Base | 1.18 | 4.13 | 1.69 | 2.91 | 1.95 | 2.37 | 1.87 | 4.13 | 4.56 | 5.80 | 0.92 | 2.60 | 0.40 | 6.98 |
| wav2vec 2.0 Large (robust) | 1.24 | 3.35 | 1.81 | 3.09 | 1.70 | 2.63 | 1.66 | 5.52 | 5.30 | 6.60 | 0.93 | 2.60 | 0.47 | 7.68 |

| Neural network | Zebra finch, group of four | Zebra finch, one bird per recording | Zebra finch, group of eight | Great tit, across year | Tree pipit, across year | Chiffchaff, across year | Tree pipit, within year | Little owl, across year | Little penguin, across night | Red-tailed black cockatoo, random split | Egyptian fruit bat, across year | Chiffchaff, within year | Rook, across year | Great tit, 50 birds, across year |
|---|---|---|---|---|---|---|---|---|---|---|---|---|---|---|
| XLS-R 300M | 1.16 | 4.68 | 2.25 | 2.89 | 1.87 | 2.70 | 1.72 | 5.48 | 5.32 | 6.89 | 0.88 | 2.57 | 0.61 | 7.29 |
| wav2vec 2.0 Conformer Large | 1.27 | 4.88 | 2.21 | 3.36 | 1.74 | 2.71 | 1.75 | 6.02 | 5.40 | 7.22 | 1.04 | 2.92 | 0.50 | 8.18 |
| MMS 300M | 1.16 | 4.47 | 2.32 | 2.81 | 1.89 | 2.70 | 1.66 | 5.51 | 5.36 | 6.95 | 0.93 | 2.59 | 0.58 | 7.45 |
| HuBERT Base | 0.81 | 4.46 | 1.80 | 2.95 | 1.85 | 2.30 | 2.01 | 5.38 | 5.09 | 6.77 | 0.96 | 2.51 | 0.52 | 7.09 |
| HuBERT Large | 0.99 | 4.22 | 1.06 | 3.29 | 1.74 | 2.66 | 1.87 | 5.59 | 5.09 | 7.04 | 0.98 | 2.72 | 0.56 | 7.76 |
| data2vec Base (100 h) | 1.20 | 2.82 | 0.99 | 3.26 | 2.20 | 2.96 | 1.97 | 5.98 | 5.06 | 7.44 | 0.91 | 2.96 | 0.34 | 8.18 |
| data2vec Base (960 h) | 1.17 | 2.64 | 1.14 | 3.28 | 2.14 | 2.91 | 1.98 | 6.05 | 4.85 | 7.06 | 0.96 | 2.68 | 0.51 | 8.09 |
| WavLM Base+ | 1.20 | 5.06 | 1.96 | 2.89 | 1.81 | 2.39 | 1.97 | 5.67 | 5.18 | 6.60 | 0.96 | 2.69 | 0.30 | 7.28 |
| WavLM Large | 1.05 | 4.20 | 1.98 | 3.15 | 1.74 | 2.64 | 1.85 | 5.59 | 5.26 | 6.68 | 1.00 | 2.78 | 0.56 | 8.10 |
| UniSpeech-SAT Base+ | 1.23 | 3.96 | 1.92 | 3.02 | 1.82 | 2.34 | 1.94 | 5.44 | 4.97 | 6.74 | 0.93 | 2.62 | 0.36 | 7.28 |
| XEUS | 1.39 | 4.85 | 2.10 | 3.07 | 2.13 | 2.70 | 2.16 | 5.76 | 5.54 | 7.68 | 0.82 | 2.96 | 0.63 | 7.82 |

*Notes:* Computed on all call clips of each dataset from principal components (Appendix S1.7). It measures how well clips group by label, whether the grouping comes from the animal or the recording.

**Table S19.** BirdNET, the 16 speech network checkpoints of the earlier run and two controls, run twice: candidate embeddings compared on each dataset.

| Dataset | Candidates in the first run | Compared | Identical |
|---|---|---|---|
| Zebra finch, group of four | 734 | 734 | 734 |
| Zebra finch, one bird per recording | 735 | 735 | 735 |
| Zebra finch, group of eight | 492 | 492 | 492 |
| Great tit, across year | 735 | 735 | 735 |
| Tree pipit, across year | 659 | 659 | 659 |
| Chiffchaff, across year | 710 | 710 | 708 |
| Tree pipit, within year | 659 | 659 | 659 |
| Little owl, across year | 735 | 735 | 735 |
| Little penguin, across night | 735 | 735 | 735 |
| Red-tailed black cockatoo, random split | 735 | 735 | 735 |
| Egyptian fruit bat, across year | 735 | 735 | 735 |
| Chiffchaff, within year | 633 | 633 | 633 |

*Notes:* Identical: the same re-ID accuracy and the same right or wrong answer on every test clip. The two candidates that differed (Appendix S1.3) were not chosen by either layer rule, and every reported embedding was identical.

**Table S20.** Datasets screened and not used, with the reason.

| Dataset | Reported scale | Reason not used |
|---|---|---|
| Roroa (great spotted kiwi, *Apteryx maxima*), Bedoya & Molles (2021) | 849 calls, 30 individuals | Call labels derive from the nest associated with each recorder and the published workflow divides calls at random, so nest, recorder, location and identity are not independently crossed. |
| Wild zebra finches, Chauhan et al. (2025) | 2,915 clips, 173 individuals | Identity labels were assigned from distinctive song exemplars, with a median of eight clips per individual; too few to hold a group of strangers back. |
| House wren, Krieg & Wade (2023) | 35 birds (17 male, 18 female) | No bird has ten calls on each of two recordings or dates, which the admission rule specified in advance requires. |
| Black-capped and Carolina chickadee hybrid zone, Palmer et al. (2025) | 55 genotyped birds; song recordings from 10 | Ten recorded birds are too few for a design across several days. |
| Ovenbird, Lapp et al. (2025) | Public code sample: 100 localised calls from 10 individuals; evaluation set described in the paper: 3,963 clips from 45 individuals | The public sample is too small, and the evaluation set named in the paper was not obtained. |

**Table S21.** Within-recording permutation test for all 38 embeddings and controls on the datasets in which recordings hold several individuals.

| Embedding | Zebra finch, group of four | Zebra finch, group of eight | Rook, across year | Zebra finch, one bird per recording |
|---|---|---|---|---|
| BirdNET v2.4 | 0.807; 0.587; 0.0001 | 0.498; 0.362; 0.0002 | 0.375; 0.224; 0.0001 | 0.967; 0.856; 0.0002 |
| Perch 2.0 | 0.830; 0.651; 0.0001 | 0.559; 0.354; 0.0001 | 0.418; 0.234; 0.0001 | 0.956; 0.789; 0.0001 |
| Perch | 0.803; 0.587; 0.0001 | 0.528; 0.371; 0.0001 | 0.393; 0.212; 0.0001 | 0.878; 0.800; 0.0008 |
| SurfPerch | 0.780; 0.596; 0.0002 | 0.397; 0.297; 0.0014 | 0.306; 0.166; 0.0001 | 0.900; 0.689; 0.0001 |
| BirdNET v3 (preview) | 0.794; 0.633; 0.0013 | 0.515; 0.354; 0.0001 | 0.328; 0.193; 0.0001 | 0.844; 0.700; 0.0001 |
| AVEX sl-BEATs | 0.417; 0.472; 0.1958 | 0.201; 0.179; 0.0115 | 0.284; 0.238; 0.0005 | 0.733; 0.578; 0.0001 |
| AVEX EfficientNet-B0 | 0.798; 0.578; 0.0001 | 0.485; 0.376; 0.0002 | 0.345; 0.192; 0.0001 | 0.878; 0.778; 0.0002 |
| AvesEcho | 0.541; 0.394; 0.0031 | 0.297; 0.240; 0.0066 | 0.343; 0.199; 0.0001 | 0.756; 0.633; 0.0001 |
| AudioProtoPNet | 0.835; 0.638; 0.0001 | 0.576; 0.419; 0.0002 | 0.386; 0.227; 0.0001 | 0.933; 0.822; 0.0005 |
| ConvNeXt (BirdSet) | 0.839; 0.615; 0.0001 | 0.502; 0.354; 0.0002 | 0.409; 0.225; 0.0001 | 0.978; 0.867; 0.0001 |
| Bird-MAE | 0.367; 0.422; 0.5322 | 0.188; 0.175; 0.0330 | 0.125; 0.221; 0.8013 | 0.567; 0.511; 0.0007 |
| ProtoCLR | 0.372; 0.394; 0.0938 | 0.118; 0.188; 0.5933 | 0.101; 0.233; 0.9833 | 0.378; 0.389; 0.0938 |
| RCL_FS_BSED | 0.661; 0.454; 0.0001 | 0.210; 0.170; 0.0201 | 0.105; 0.219; 0.9063 | 0.456; 0.478; 0.3139 |
| BirdAVES | 0.638; 0.509; 0.0011 | 0.245; 0.245; 0.0530 | 0.151; 0.191; 0.3141 | 0.511; 0.433; 0.0001 |
| AVES | 0.445; 0.399; 0.0171 | 0.175; 0.183; 0.0942 | 0.179; 0.183; 0.0665 | 0.556; 0.489; 0.0011 |
| NatureBEATs | 0.734; 0.491; 0.0001 | 0.201; 0.166; 0.0069 | 0.166; 0.242; 0.7836 | 0.600; 0.456; 0.0001 |
| BioLingual | 0.417; 0.376; 0.0054 | 0.192; 0.201; 0.0741 | 0.281; 0.248; 0.0008 | 0.956; 0.811; 0.0001 |
| BEATs | 0.495; 0.500; 0.0881 | 0.197; 0.188; 0.0446 | 0.207; 0.193; 0.0167 | 0.500; 0.444; 0.0007 |
| AudioMAE | 0.509; 0.413; 0.0006 | 0.275; 0.179; 0.0005 | 0.185; 0.237; 0.4203 | 0.733; 0.656; 0.0015 |
| VGGish | 0.702; 0.477; 0.0001 | 0.445; 0.402; 0.0064 | 0.180; 0.152; 0.0009 | 0.533; 0.511; 0.0280 |
| ECAPA-TDNN (speaker) | 0.711; 0.537; 0.0001 | 0.454; 0.245; 0.0001 | 0.271; 0.188; 0.0001 | 0.844; 0.678; 0.0001 |
| ResNet (speaker) | 0.771; 0.532; 0.0001 | 0.467; 0.271; 0.0001 | 0.297; 0.227; 0.0001 | 0.878; 0.789; 0.0005 |
| x-vector (speaker) | 0.789; 0.518; 0.0001 | 0.454; 0.319; 0.0001 | 0.296; 0.186; 0.0001 | 0.767; 0.667; 0.0004 |
| wav2vec 2.0 Base | 0.826; 0.560; 0.0001 | 0.380; 0.240; 0.0010 | 0.153; 0.142; 0.0128 | 0.622; 0.622; 0.0530 |
| wav2vec 2.0 Large (robust) | 0.555; 0.399; 0.0048 | 0.310; 0.245; 0.0124 | 0.125; 0.217; 0.7978 | 0.256; 0.311; 0.3913 |
| XLS-R 300M | 0.789; 0.606; 0.0004 | 0.367; 0.227; 0.0025 | 0.187; 0.185; 0.0408 | 0.756; 0.700; 0.0023 |
| wav2vec 2.0 Conformer Large | 0.780; 0.555; 0.0001 | 0.406; 0.367; 0.0192 | 0.256; 0.179; 0.0001 | 0.700; 0.611; 0.0003 |
| MMS 300M | 0.798; 0.587; 0.0002 | 0.402; 0.271; 0.0015 | 0.163; 0.164; 0.0511 | 0.722; 0.667; 0.0013 |
| HuBERT Base | 0.638; 0.550; 0.0035 | 0.376; 0.314; 0.0083 | 0.235; 0.156; 0.0001 | 0.744; 0.644; 0.0001 |
| HuBERT Large | 0.716; 0.546; 0.0008 | 0.218; 0.197; 0.0175 | 0.197; 0.209; 0.2489 | 0.656; 0.600; 0.0011 |
| data2vec Base (100 h) | 0.743; 0.606; 0.0011 | 0.480; 0.371; 0.0001 | 0.169; 0.162; 0.0250 | 0.478; 0.433; 0.0005 |
| data2vec Base (960 h) | 0.743; 0.583; 0.0007 | 0.371; 0.419; 0.2285 | 0.236; 0.187; 0.0001 | 0.422; 0.378; 0.0020 |
| WavLM Base+ | 0.817; 0.573; 0.0001 | 0.397; 0.310; 0.0022 | 0.105; 0.149; 0.8075 | 0.756; 0.689; 0.0010 |
| WavLM Large | 0.743; 0.500; 0.0001 | 0.314; 0.288; 0.0287 | 0.166; 0.192; 0.5178 | 0.711; 0.644; 0.0003 |
| UniSpeech-SAT Base+ | 0.794; 0.573; 0.0001 | 0.410; 0.328; 0.0017 | 0.181; 0.162; 0.0067 | 0.789; 0.678; 0.0001 |
| XEUS | 0.817; 0.560; 0.0001 | 0.445; 0.354; 0.0019 | 0.247; 0.190; 0.0001 | 0.867; 0.767; 0.0002 |
| Loudness control | 0.381; 0.353; 0.0266 | 0.210; 0.275; 0.3593 | 0.126; 0.125; 0.0480 | 0.233; 0.278; 0.5674 |
| Duration control | 0.468; 0.468; 0.9617 | 0.284; 0.284; 0.1037 | 0.030; 0.180; 0.9615 | 0.189; 0.200; 0.7396 |

*Notes:* Each cell: re-ID accuracy; 95th percentile of the within-recording test; its p (9,999 shuffles of labels among clips of the same recording, or, for the zebra finch recorded singly, the same day). Entries with $p < 0.05$: Zebra finch, group of four 33; Zebra finch, group of eight 31; Rook, across year 25; Zebra finch, one bird per recording 32. Each embedding at its layer chosen by the Fisher rule.

**Table S22.** Exploratory analysis, specified after the fourth round of review: each neural network read at its own final output against BirdNET v2.4.

*Zebra finch, group of four*: BirdNET v2.4 0.807 (0.669 to 0.963).

| **Neural network** | **Output read, re-ID accuracy** | **Difference from BirdNET** | **Holm p** |
|---|---|---|---|
| Perch 2.0 | embedding: 0.830 (0.679 to 0.923) | 0.023 (−0.037 to 0.073) | 1.0000 |
| Perch | embedding: 0.803 (0.717 to 0.838) | −0.005 (−0.125 to 0.079) | 1.0000 |
| SurfPerch | embedding: 0.780 (0.682 to 0.860) | −0.028 (−0.092 to 0.057) | 1.0000 |
| BirdNET v3 (preview) | embedding: 0.794 (0.696 to 0.900) | −0.014 (−0.062 to 0.027) | 1.0000 |
| AVEX sl-BEATs | embedding: 0.716 (0.596 to 0.900) | −0.092 (−0.140 to 0.017) | 0.9594 |
| AVEX EfficientNet-B0 | embedding: 0.798 (0.660 to 0.867) | −0.009 (−0.094 to 0.055) | 1.0000 |
| AvesEcho | embedding: 0.798 (0.702 to 0.942) | −0.009 (−0.036 to 0.075) | 1.0000 |
| AudioProtoPNet | embedding: 0.835 (0.679 to 0.950) | 0.028 (−0.013 to 0.064) | 1.0000 |
| ConvNeXt (BirdSet) | embedding: 0.839 (0.698 to 0.950) | 0.032 (−0.010 to 0.070) | 1.0000 |
| Bird-MAE | embedding: 0.532 (0.302 to 0.606) | −0.275 (−0.437 to −0.175) | < 0.0074 |
| ProtoCLR | embedding: 0.610 (0.506 to 0.672) | −0.197 (−0.376 to −0.063) | < 0.0074 |
| RCL_FS_BSED | embedding: 0.661 (0.491 to 0.835) | −0.147 (−0.273 to −0.027) | 0.6216 |
| BirdAVES | embedding: 0.638 (0.396 to 0.774) | −0.170 (−0.319 to −0.123) | < 0.0074 |
| AVES | embedding: 0.596 (0.385 to 0.742) | −0.211 (−0.287 to −0.152) | < 0.0074 |
| NatureBEATs | embedding: 0.780 (0.660 to 0.908) | −0.028 (−0.054 to −0.009) | 0.1716 |
| BioLingual | embedding: 0.761 (0.623 to 0.938) | −0.046 (−0.072 to −0.012) | 0.1716 |
| BEATs | embedding: 0.725 (0.561 to 0.946) | −0.083 (−0.130 to −0.012) | 0.1716 |
| AudioMAE | embedding: 0.683 (0.482 to 0.958) | −0.124 (−0.251 to 0.023) | 1.0000 |
| VGGish | embedding: 0.702 (0.486 to 0.873) | −0.106 (−0.182 to −0.058) | < 0.0074 |
| ECAPA-TDNN (speaker) | embedding: 0.702 (0.528 to 0.850) | −0.106 (−0.151 to −0.086) | < 0.0074 |
| ResNet (speaker) | embedding: 0.794 (0.669 to 0.896) | −0.014 (−0.059 to 0.038) | 1.0000 |
| x-vector (speaker) | embedding: 0.729 (0.509 to 0.914) | −0.078 (−0.170 to −0.042) | < 0.0074 |
| wav2vec 2.0 Base | layer 12, full speed: 0.390 (0.188 to 0.532) | −0.417 (−0.758 to −0.202) | < 0.0074 |
| wav2vec 2.0 Large (robust) | layer 24, full speed: 0.298 (0.123 to 0.585) | −0.509 (−0.620 to −0.170) | < 0.0074 |
| XLS-R 300M | layer 24, full speed: 0.683 (0.434 to 0.908) | −0.124 (−0.245 to −0.047) | < 0.0074 |
| wav2vec 2.0 Conformer Large | layer 24, full speed: 0.651 (0.517 to 0.824) | −0.156 (−0.224 to −0.046) | 0.1716 |
| MMS 300M | layer 24, full speed: 0.665 (0.392 to 0.937) | −0.142 (−0.254 to −0.019) | 0.1716 |
| HuBERT Base | layer 12, full speed: 0.555 (0.424 to 0.648) | −0.252 (−0.483 to −0.088) | < 0.0074 |
| HuBERT Large | layer 24, full speed: 0.683 (0.561 to 0.824) | −0.124 (−0.140 to −0.075) | < 0.0074 |
| data2vec Base (100 h) | layer 12, full speed: 0.569 (0.447 to 0.647) | −0.239 (−0.459 to −0.098) | < 0.0074 |
| data2vec Base (960 h) | layer 12, full speed: 0.390 (0.342 to 0.491) | −0.417 (−0.617 to −0.189) | 0.1716 |
| WavLM Base+ | layer 12, full speed: 0.528 (0.423 to 0.623) | −0.280 (−0.538 to −0.088) | 0.1716 |
| WavLM Large | layer 24, full speed: 0.752 (0.604 to 0.892) | −0.055 (−0.108 to −0.008) | 0.1716 |
| UniSpeech-SAT Base+ | layer 12, full speed: 0.670 (0.568 to 0.769) | −0.138 (−0.188 to −0.075) | < 0.0074 |
| XEUS | layer 18, full speed: 0.495 (0.000 to 0.917) | −0.312 (−0.912 to 0.174) | 1.0000 |
| Loudness control | summary: 0.381 (0.018 to 0.846) | −0.427 (−0.693 to −0.118) | < 0.0074 |
| Duration control | summary: 0.468 (0.000 to 0.850) | −0.339 (−0.906 to 0.107) | 1.0000 |

*Zebra finch, one bird per recording*: BirdNET v2.4 0.967 (0.933 to 0.989).

| **Neural network** | **Output read, re-ID accuracy** | **Difference from BirdNET** | **Holm p** |
|---|---|---|---|
| Perch 2.0 | embedding: 0.956 (0.911 to 1.000) | −0.011 (−0.044 to 0.022) | 1.0000 |

| Neural network | Output read, re-ID accuracy | Difference from BirdNET | Holm p |
|---|---|---|---|
| Perch | embedding: 0.878 (0.722 to 0.978) | −0.089 (−0.222 to 0.000) | 0.4338 |
| SurfPerch | embedding: 0.900 (0.833 to 0.967) | −0.067 (−0.111 to −0.022) | 0.0154 |
| BirdNET v3 (preview) | embedding: 0.844 (0.744 to 0.933) | −0.122 (−0.200 to −0.044) | 0.0240 |
| AVEX sl-BEATs | embedding: 0.889 (0.811 to 0.967) | −0.078 (−0.156 to −0.022) | 0.4338 |
| AVEX EfficientNet-B0 | embedding: 0.878 (0.767 to 0.978) | −0.089 (−0.189 to 0.000) | 0.4416 |
| AvesEcho | embedding: 0.889 (0.756 to 0.978) | −0.078 (−0.189 to 0.000) | 0.4338 |
| AudioProtoPNet | embedding: 0.933 (0.844 to 1.000) | −0.033 (−0.111 to 0.022) | 1.0000 |
| ConvNeXt (BirdSet) | embedding: 0.978 (0.933 to 1.000) | 0.011 (−0.022 to 0.044) | 1.0000 |
| Bird-MAE | embedding: 0.689 (0.522 to 0.833) | −0.278 (−0.444 to −0.122) | < 0.0074 |
| ProtoCLR | embedding: 0.389 (0.189 to 0.589) | −0.578 (−0.767 to −0.389) | < 0.0074 |
| RCL_FS_BSED | embedding: 0.456 (0.189 to 0.722) | −0.511 (−0.756 to −0.267) | < 0.0074 |
| BirdAVES | embedding: 0.511 (0.333 to 0.678) | −0.456 (−0.611 to −0.300) | < 0.0074 |
| AVES | embedding: 0.567 (0.411 to 0.722) | −0.400 (−0.533 to −0.267) | < 0.0074 |
| NatureBEATs | embedding: 0.856 (0.767 to 0.944) | −0.111 (−0.178 to −0.044) | 0.0052 |
| BioLingual | embedding: 0.956 (0.911 to 0.989) | −0.011 (−0.044 to 0.022) | 1.0000 |
| BEATs | embedding: 0.700 (0.500 to 0.878) | −0.267 (−0.444 to −0.111) | < 0.0074 |
| AudioMAE | embedding: 0.700 (0.522 to 0.845) | −0.267 (−0.467 to −0.100) | 0.0052 |
| VGGish | embedding: 0.533 (0.333 to 0.733) | −0.433 (−0.622 to −0.244) | < 0.0074 |
| ECAPA-TDNN (speaker) | embedding: 0.722 (0.533 to 0.878) | −0.244 (−0.411 to −0.100) | < 0.0074 |
| ResNet (speaker) | embedding: 0.878 (0.756 to 0.967) | −0.089 (−0.200 to 0.000) | 0.4416 |
| x-vector (speaker) | embedding: 0.767 (0.622 to 0.900) | −0.200 (−0.333 to −0.089) | 0.0028 |
| wav2vec 2.0 Base | layer 12, full speed: 0.311 (0.144 to 0.489) | −0.656 (−0.822 to −0.478) | < 0.0074 |
| wav2vec 2.0 Large (robust) | layer 24, full speed: 0.256 (0.056 to 0.478) | −0.711 (−0.889 to −0.511) | < 0.0074 |
| XLS-R 300M | layer 24, full speed: 0.356 (0.178 to 0.556) | −0.611 (−0.778 to −0.422) | < 0.0074 |
| wav2vec 2.0 Conformer Large | layer 24, full speed: 0.478 (0.267 to 0.678) | −0.489 (−0.667 to −0.322) | < 0.0074 |
| MMS 300M | layer 24, full speed: 0.578 (0.367 to 0.778) | −0.389 (−0.589 to −0.189) | < 0.0074 |
| HuBERT Base | layer 12, full speed: 0.433 (0.267 to 0.589) | −0.533 (−0.689 to −0.378) | < 0.0074 |
| HuBERT Large | layer 24, full speed: 0.422 (0.244 to 0.622) | −0.544 (−0.700 to −0.367) | < 0.0074 |
| data2vec Base (100 h) | layer 12, full speed: 0.389 (0.167 to 0.622) | −0.578 (−0.778 to −0.367) | < 0.0074 |
| data2vec Base (960 h) | layer 12, full speed: 0.367 (0.156 to 0.589) | −0.600 (−0.789 to −0.400) | < 0.0074 |
| WavLM Base+ | layer 12, full speed: 0.422 (0.233 to 0.622) | −0.544 (−0.722 to −0.356) | < 0.0074 |
| WavLM Large | layer 24, full speed: 0.611 (0.411 to 0.800) | −0.356 (−0.556 to −0.167) | < 0.0074 |
| UniSpeech-SAT Base+ | layer 12, full speed: 0.433 (0.233 to 0.633) | −0.533 (−0.711 to −0.344) | < 0.0074 |
| XEUS | layer 18, full speed: 0.278 (0.100 to 0.478) | −0.689 (−0.856 to −0.500) | < 0.0074 |
| Loudness control | summary: 0.233 (0.100 to 0.389) | −0.733 (−0.878 to −0.578) | < 0.0074 |
| Duration control | summary: 0.189 (0.000 to 0.456) | −0.778 (−0.967 to −0.533) | < 0.0074 |

*Zebra finch, group of eight*: BirdNET v2.4 0.498 (0.181 to 0.731).

| Neural network | Output read, re-ID accuracy | Difference from BirdNET | Holm p |
|---|---|---|---|
| Perch 2.0 | embedding: 0.559 (0.374 to 0.719) | 0.061 (−0.034 to 0.209) | 1.0000 |
| Perch | embedding: 0.528 (0.365 to 0.725) | 0.031 (−0.093 to 0.199) | 1.0000 |
| SurfPerch | embedding: 0.397 (0.171 to 0.689) | −0.100 (−0.213 to 0.016) | 1.0000 |
| BirdNET v3 (preview) | embedding: 0.515 (0.347 to 0.692) | 0.017 (−0.099 to 0.179) | 1.0000 |
| AVEX sl-BEATs | embedding: 0.437 (0.232 to 0.664) | −0.061 (−0.163 to 0.085) | 1.0000 |
| AVEX EfficientNet-B0 | embedding: 0.485 (0.234 to 0.705) | −0.013 (−0.070 to 0.067) | 1.0000 |
| AvesEcho | embedding: 0.568 (0.434 to 0.759) | 0.070 (−0.151 to 0.341) | 1.0000 |

| Neural network | Output read, re-ID accuracy | Difference from BirdNET | Holm p |
|---|---|---|---|
| AudioProtoPNet | embedding: 0.576 (0.397 to 0.748) | 0.079 (−0.037 to 0.228) | 1.0000 |
| ConvNeXt (BirdSet) | embedding: 0.502 (0.284 to 0.743) | 0.004 (−0.095 to 0.128) | 1.0000 |
| Bird-MAE | embedding: 0.223 (0.030 to 0.611) | −0.275 (−0.530 to −0.026) | < 0.0074 |
| ProtoCLR | embedding: 0.118 (0.000 to 0.450) | −0.380 (−0.640 to −0.047) | 0.4898 |
| RCL_FS_BSED | embedding: 0.210 (0.013 to 0.627) | −0.288 (−0.539 to −0.004) | 1.0000 |
| BirdAVES | embedding: 0.245 (0.091 to 0.481) | −0.253 (−0.409 to −0.059) | < 0.0074 |
| AVES | embedding: 0.175 (0.061 to 0.406) | −0.323 (−0.550 to −0.038) | 0.4288 |
| NatureBEATs | embedding: 0.472 (0.309 to 0.722) | −0.026 (−0.312 to 0.367) | 1.0000 |
| BioLingual | embedding: 0.397 (0.226 to 0.726) | −0.100 (−0.296 to 0.099) | 1.0000 |
| BEATs | embedding: 0.419 (0.131 to 0.670) | −0.079 (−0.179 to 0.011) | 1.0000 |
| AudioMAE | embedding: 0.376 (0.229 to 0.609) | −0.122 (−0.291 to 0.093) | 1.0000 |
| VGGish | embedding: 0.445 (0.114 to 0.681) | −0.052 (−0.182 to 0.010) | 1.0000 |
| ECAPA-TDNN (speaker) | embedding: 0.358 (0.222 to 0.535) | −0.140 (−0.291 to 0.076) | 1.0000 |
| ResNet (speaker) | embedding: 0.467 (0.314 to 0.720) | −0.031 (−0.180 to 0.155) | 1.0000 |
| x-vector (speaker) | embedding: 0.310 (0.148 to 0.607) | −0.188 (−0.408 to 0.100) | 1.0000 |
| wav2vec 2.0 Base | layer 12, full speed: 0.192 (0.115 to 0.310) | −0.306 (−0.462 to −0.054) | 0.3894 |
| wav2vec 2.0 Large (robust) | layer 24, full speed: 0.245 (0.000 to 0.639) | −0.253 (−0.706 to 0.451) | 1.0000 |
| XLS-R 300M | layer 24, full speed: 0.249 (0.064 to 0.573) | −0.249 (−0.532 to 0.100) | 1.0000 |
| wav2vec 2.0 Conformer Large | layer 24, full speed: 0.389 (0.201 to 0.552) | −0.109 (−0.328 to 0.174) | 1.0000 |
| MMS 300M | layer 24, full speed: 0.223 (0.037 to 0.607) | −0.275 (−0.528 to 0.013) | 1.0000 |
| HuBERT Base | layer 12, full speed: 0.310 (0.146 to 0.562) | −0.188 (−0.379 to 0.065) | 1.0000 |
| HuBERT Large | layer 24, full speed: 0.507 (0.260 to 0.684) | 0.009 (−0.204 to 0.199) | 1.0000 |
| data2vec Base (100 h) | layer 12, full speed: 0.210 (0.087 to 0.429) | −0.288 (−0.495 to 0.000) | 1.0000 |
| data2vec Base (960 h) | layer 12, full speed: 0.205 (0.064 to 0.319) | −0.293 (−0.416 to −0.116) | < 0.0074 |
| WavLM Base+ | layer 12, full speed: 0.376 (0.147 to 0.552) | −0.122 (−0.311 to 0.047) | 1.0000 |
| WavLM Large | layer 24, full speed: 0.489 (0.257 to 0.692) | −0.009 (−0.161 to 0.143) | 1.0000 |
| UniSpeech-SAT Base+ | layer 12, full speed: 0.397 (0.208 to 0.581) | −0.100 (−0.331 to 0.220) | 1.0000 |
| XEUS | layer 18, full speed: 0.236 (0.016 to 0.500) | −0.262 (−0.453 to −0.059) | 0.3604 |
| Loudness control | summary: 0.210 (0.014 to 0.608) | −0.288 (−0.565 to 0.079) | 1.0000 |
| Duration control | summary: 0.284 (0.000 to 0.572) | −0.214 (−0.661 to 0.000) | 1.0000 |

*Great tit, across year*: BirdNET v2.4 0.444 (0.319 to 0.569).

| Neural network | Output read, re-ID accuracy | Difference from BirdNET | Holm p |
|---|---|---|---|
| Perch 2.0 | embedding: 0.450 (0.312 to 0.588) | 0.006 (−0.094 to 0.100) | 1.0000 |
| Perch | embedding: 0.294 (0.194 to 0.400) | −0.150 (−0.256 to −0.056) | 0.0088 |
| SurfPerch | embedding: 0.375 (0.275 to 0.475) | −0.069 (−0.131 to −0.012) | 0.1452 |
| BirdNET v3 (preview) | embedding: 0.156 (0.106 to 0.206) | −0.287 (−0.437 to −0.144) | < 0.0074 |
| AVEX sl-BEATs | embedding: 0.381 (0.287 to 0.475) | −0.062 (−0.131 to 0.006) | 0.3304 |
| AVEX EfficientNet-B0 | embedding: 0.338 (0.225 to 0.456) | −0.106 (−0.231 to 0.000) | 0.2970 |
| AvesEcho | embedding: 0.438 (0.294 to 0.581) | −0.006 (−0.069 to 0.056) | 1.0000 |
| AudioProtoPNet | embedding: 0.294 (0.194 to 0.394) | −0.150 (−0.269 to −0.044) | 0.0484 |
| ConvNeXt (BirdSet) | embedding: 0.325 (0.200 to 0.463) | −0.119 (−0.225 to −0.025) | 0.0960 |
| Bird-MAE | embedding: 0.138 (0.044 to 0.244) | −0.306 (−0.469 to −0.131) | 0.0168 |
| ProtoCLR | embedding: 0.113 (0.031 to 0.212) | −0.331 (−0.500 to −0.150) | 0.0128 |
| RCL_FS_BSED | embedding: 0.219 (0.094 to 0.362) | −0.225 (−0.338 to −0.106) | 0.0088 |
| BirdAVES | embedding: 0.362 (0.231 to 0.506) | −0.081 (−0.175 to 0.019) | 0.3462 |

| Neural network | Output read, re-ID accuracy | Difference from BirdNET | Holm p |
|---|---|---|---|
| AVES | embedding: 0.287 (0.169 to 0.425) | −0.156 (−0.281 to −0.037) | 0.0960 |
| NatureBEATs | embedding: 0.269 (0.131 to 0.412) | −0.175 (−0.294 to −0.056) | 0.0522 |
| BioLingual | embedding: 0.319 (0.206 to 0.444) | −0.125 (−0.206 to −0.050) | 0.0150 |
| BEATs | embedding: 0.250 (0.163 to 0.344) | −0.194 (−0.325 to −0.075) | 0.0114 |
| AudioMAE | embedding: 0.219 (0.131 to 0.306) | −0.225 (−0.356 to −0.112) | < 0.0074 |
| VGGish | embedding: 0.169 (0.087 to 0.250) | −0.275 (−0.394 to −0.150) | < 0.0074 |
| ECAPA-TDNN (speaker) | embedding: 0.256 (0.175 to 0.344) | −0.188 (−0.331 to −0.056) | 0.0480 |
| ResNet (speaker) | embedding: 0.237 (0.144 to 0.338) | −0.206 (−0.319 to −0.106) | < 0.0074 |
| x-vector (speaker) | embedding: 0.212 (0.106 to 0.338) | −0.231 (−0.350 to −0.125) | < 0.0074 |
| wav2vec 2.0 Base | layer 12, full speed: 0.219 (0.106 to 0.350) | −0.225 (−0.356 to −0.094) | 0.0114 |
| wav2vec 2.0 Large (robust) | layer 24, full speed: 0.119 (0.031 to 0.231) | −0.325 (−0.506 to −0.131) | 0.0168 |
| XLS-R 300M | layer 24, full speed: 0.119 (0.056 to 0.194) | −0.325 (−0.462 to −0.181) | < 0.0074 |
| wav2vec 2.0 Conformer Large | layer 24, full speed: 0.181 (0.100 to 0.269) | −0.262 (−0.412 to −0.113) | 0.0114 |
| MMS 300M | layer 24, full speed: 0.138 (0.056 to 0.231) | −0.306 (−0.463 to −0.138) | 0.0088 |
| HuBERT Base | layer 12, full speed: 0.156 (0.100 to 0.212) | −0.287 (−0.425 to −0.150) | < 0.0074 |
| HuBERT Large | layer 24, full speed: 0.094 (0.037 to 0.163) | −0.350 (−0.494 to −0.212) | < 0.0074 |
| data2vec Base (100 h) | layer 12, full speed: 0.131 (0.056 to 0.219) | −0.312 (−0.462 to −0.163) | < 0.0074 |
| data2vec Base (960 h) | layer 12, full speed: 0.056 (0.013 to 0.119) | −0.387 (−0.537 to −0.237) | < 0.0074 |
| WavLM Base+ | layer 12, full speed: 0.175 (0.113 to 0.244) | −0.269 (−0.375 to −0.169) | < 0.0074 |
| WavLM Large | layer 24, full speed: 0.131 (0.069 to 0.200) | −0.312 (−0.450 to −0.181) | < 0.0074 |
| UniSpeech-SAT Base+ | layer 12, full speed: 0.194 (0.119 to 0.275) | −0.250 (−0.363 to −0.144) | < 0.0074 |
| XEUS | layer 18, full speed: 0.237 (0.131 to 0.350) | −0.206 (−0.344 to −0.069) | 0.0484 |
| Loudness control | summary: 0.113 (0.031 to 0.206) | −0.331 (−0.494 to −0.169) | < 0.0074 |
| Duration control | summary: 0.069 (0.000 to 0.175) | −0.375 (−0.537 to −0.200) | < 0.0074 |

*Tree pipit, across year*: BirdNET v2.4 0.214 (0.117 to 0.336).

| Neural network | Output read, re-ID accuracy | Difference from BirdNET | Holm p |
|---|---|---|---|
| Perch 2.0 | embedding: 0.447 (0.294 to 0.625) | 0.233 (0.149 to 0.310) | < 0.0074 |
| Perch | embedding: 0.351 (0.219 to 0.517) | 0.137 (0.064 to 0.207) | 0.0348 |
| SurfPerch | embedding: 0.243 (0.153 to 0.358) | 0.029 (−0.015 to 0.083) | 1.0000 |
| BirdNET v3 (preview) | embedding: 0.160 (0.096 to 0.228) | −0.054 (−0.173 to 0.035) | 1.0000 |
| AVEX sl-BEATs | embedding: 0.240 (0.121 to 0.379) | 0.026 (−0.051 to 0.100) | 1.0000 |
| AVEX EfficientNet-B0 | embedding: 0.211 (0.114 to 0.319) | −0.003 (−0.091 to 0.091) | 1.0000 |
| AvesEcho | embedding: 0.089 (0.019 to 0.190) | −0.125 (−0.192 to −0.055) | 0.0448 |
| AudioProtoPNet | embedding: 0.201 (0.102 to 0.317) | −0.013 (−0.070 to 0.036) | 1.0000 |
| ConvNeXt (BirdSet) | embedding: 0.288 (0.138 to 0.464) | 0.073 (−0.023 to 0.156) | 1.0000 |
| Bird-MAE | embedding: 0.051 (0.018 to 0.095) | −0.163 (−0.294 to −0.052) | 0.0300 |
| ProtoCLR | embedding: 0.083 (0.030 to 0.156) | −0.131 (−0.261 to 0.000) | 1.0000 |
| RCL_FS_BSED | embedding: 0.045 (0.017 to 0.071) | −0.169 (−0.283 to −0.080) | < 0.0074 |
| BirdAVES | embedding: 0.214 (0.116 to 0.334) | 0.000 (−0.048 to 0.039) | 1.0000 |
| AVES | embedding: 0.096 (0.033 to 0.196) | −0.118 (−0.202 to −0.057) | < 0.0074 |
| NatureBEATs | embedding: 0.192 (0.087 to 0.301) | −0.022 (−0.106 to 0.064) | 1.0000 |
| BioLingual | embedding: 0.227 (0.101 to 0.375) | 0.013 (−0.044 to 0.058) | 1.0000 |
| BEATs | embedding: 0.058 (0.037 to 0.083) | −0.157 (−0.275 to −0.065) | < 0.0074 |
| AudioMAE | embedding: 0.169 (0.113 to 0.237) | −0.045 (−0.135 to 0.054) | 1.0000 |
| VGGish | embedding: 0.070 (0.034 to 0.109) | −0.144 (−0.249 to −0.062) | < 0.0074 |

| Neural network | Output read, re-ID accuracy | Difference from BirdNET | Holm p |
|---|---|---|---|
| ECAPA-TDNN (speaker) | embedding: 0.131 (0.082 to 0.180) | −0.083 (−0.227 to 0.034) | 1.0000 |
| ResNet (speaker) | embedding: 0.150 (0.077 to 0.247) | −0.064 (−0.143 to 0.011) | 1.0000 |
| x-vector (speaker) | embedding: 0.182 (0.098 to 0.303) | −0.032 (−0.118 to 0.045) | 1.0000 |
| wav2vec 2.0 Base | layer 12, full speed: 0.086 (0.019 to 0.172) | −0.128 (−0.255 to 0.014) | 1.0000 |
| wav2vec 2.0 Large (robust) | layer 24, full speed: 0.064 (0.000 to 0.142) | −0.150 (−0.296 to −0.013) | 0.7550 |
| XLS-R 300M | layer 24, full speed: 0.083 (0.028 to 0.143) | −0.131 (−0.239 to −0.049) | < 0.0074 |
| wav2vec 2.0 Conformer Large | layer 24, full speed: 0.195 (0.130 to 0.270) | −0.019 (−0.085 to 0.041) | 1.0000 |
| MMS 300M | layer 24, full speed: 0.115 (0.040 to 0.196) | −0.099 (−0.257 to 0.041) | 1.0000 |
| HuBERT Base | layer 12, full speed: 0.147 (0.053 to 0.300) | −0.067 (−0.153 to 0.020) | 1.0000 |
| HuBERT Large | layer 24, full speed: 0.147 (0.053 to 0.269) | −0.067 (−0.175 to 0.022) | 1.0000 |
| data2vec Base (100 h) | layer 12, full speed: 0.141 (0.044 to 0.266) | −0.073 (−0.191 to 0.056) | 1.0000 |
| data2vec Base (960 h) | layer 12, full speed: 0.134 (0.048 to 0.223) | −0.080 (−0.221 to 0.052) | 1.0000 |
| WavLM Base+ | layer 12, full speed: 0.144 (0.064 to 0.263) | −0.070 (−0.162 to 0.000) | 1.0000 |
| WavLM Large | layer 24, full speed: 0.137 (0.053 to 0.273) | −0.077 (−0.164 to −0.009) | 0.6396 |
| UniSpeech-SAT Base+ | layer 12, full speed: 0.144 (0.059 to 0.262) | −0.070 (−0.134 to −0.003) | 1.0000 |
| XEUS | layer 18, full speed: 0.070 (0.019 to 0.140) | −0.144 (−0.237 to −0.072) | < 0.0074 |
| Loudness control | summary: 0.073 (0.018 to 0.138) | −0.141 (−0.261 to −0.033) | 0.2484 |
| Duration control | summary: 0.083 (0.000 to 0.234) | −0.131 (−0.307 to 0.079) | 1.0000 |

*Chiffchaff, across year*: BirdNET v2.4 0.160 (0.035 to 0.330).

| Neural network | Output read, re-ID accuracy | Difference from BirdNET | Holm p |
|---|---|---|---|
| Perch 2.0 | embedding: 0.160 (0.034 to 0.289) | 0.000 (−0.091 to 0.087) | 1.0000 |
| Perch | embedding: 0.150 (0.044 to 0.277) | −0.010 (−0.078 to 0.068) | 1.0000 |
| SurfPerch | embedding: 0.255 (0.103 to 0.421) | 0.095 (0.015 to 0.192) | 0.6808 |
| BirdNET v3 (preview) | embedding: 0.085 (0.033 to 0.151) | −0.075 (−0.245 to 0.077) | 1.0000 |
| AVEX sl-BEATs | embedding: 0.150 (0.054 to 0.267) | −0.010 (−0.090 to 0.086) | 1.0000 |
| AVEX EfficientNet-B0 | embedding: 0.120 (0.036 to 0.234) | −0.040 (−0.116 to 0.033) | 1.0000 |
| AvesEcho | embedding: 0.220 (0.018 to 0.432) | 0.060 (−0.043 to 0.207) | 1.0000 |
| AudioProtoPNet | embedding: 0.115 (0.047 to 0.197) | −0.045 (−0.155 to 0.054) | 1.0000 |
| ConvNeXt (BirdSet) | embedding: 0.135 (0.034 to 0.262) | −0.025 (−0.098 to 0.051) | 1.0000 |
| Bird-MAE | embedding: 0.095 (0.025 to 0.175) | −0.065 (−0.227 to 0.083) | 1.0000 |
| ProtoCLR | embedding: 0.195 (0.041 to 0.379) | 0.035 (−0.141 to 0.222) | 1.0000 |
| RCL_FS_BSED | embedding: 0.150 (0.016 to 0.312) | −0.010 (−0.083 to 0.064) | 1.0000 |
| BirdAVES | embedding: 0.175 (0.061 to 0.310) | 0.015 (−0.054 to 0.095) | 1.0000 |
| AVES | embedding: 0.140 (0.015 to 0.305) | −0.020 (−0.081 to 0.053) | 1.0000 |
| NatureBEATs | embedding: 0.170 (0.050 to 0.307) | 0.010 (−0.041 to 0.056) | 1.0000 |
| BioLingual | embedding: 0.160 (0.031 to 0.289) | 0.000 (−0.086 to 0.078) | 1.0000 |
| BEATs | embedding: 0.140 (0.049 to 0.234) | −0.020 (−0.123 to 0.058) | 1.0000 |
| AudioMAE | embedding: 0.150 (0.046 to 0.279) | −0.010 (−0.071 to 0.045) | 1.0000 |
| VGGish | embedding: 0.095 (0.000 to 0.275) | −0.065 (−0.126 to −0.005) | 1.0000 |
| ECAPA-TDNN (speaker) | embedding: 0.155 (0.044 to 0.295) | −0.005 (−0.087 to 0.092) | 1.0000 |
| ResNet (speaker) | embedding: 0.080 (0.028 to 0.140) | −0.080 (−0.203 to 0.027) | 1.0000 |
| x-vector (speaker) | embedding: 0.105 (0.042 to 0.181) | −0.055 (−0.157 to 0.026) | 1.0000 |
| wav2vec 2.0 Base | layer 12, full speed: 0.195 (0.017 to 0.390) | 0.035 (−0.053 to 0.112) | 1.0000 |
| wav2vec 2.0 Large (robust) | layer 24, full speed: 0.140 (0.000 to 0.341) | −0.020 (−0.247 to 0.218) | 1.0000 |
| XLS-R 300M | layer 24, full speed: 0.165 (0.028 to 0.349) | 0.005 (−0.083 to 0.109) | 1.0000 |

| Neural network | Output read, re-ID accuracy | Difference from BirdNET | Holm p |
|---|---|---|---|
| wav2vec 2.0 Conformer Large | layer 24, full speed: 0.195 (0.094 to 0.308) | 0.035 (−0.054 to 0.116) | 1.0000 |
| MMS 300M | layer 24, full speed: 0.170 (0.000 to 0.404) | 0.010 (−0.093 to 0.151) | 1.0000 |
| HuBERT Base | layer 12, full speed: 0.195 (0.016 to 0.438) | 0.035 (−0.073 to 0.190) | 1.0000 |
| HuBERT Large | layer 24, full speed: 0.190 (0.049 to 0.354) | 0.030 (−0.062 to 0.126) | 1.0000 |
| data2vec Base (100 h) | layer 12, full speed: 0.130 (0.028 to 0.262) | −0.030 (−0.107 to 0.049) | 1.0000 |
| data2vec Base (960 h) | layer 12, full speed: 0.170 (0.066 to 0.294) | 0.010 (−0.071 to 0.093) | 1.0000 |
| WavLM Base+ | layer 12, full speed: 0.135 (0.011 to 0.323) | −0.025 (−0.078 to 0.023) | 1.0000 |
| WavLM Large | layer 24, full speed: 0.195 (0.044 to 0.375) | 0.035 (−0.074 to 0.170) | 1.0000 |
| UniSpeech-SAT Base+ | layer 12, full speed: 0.165 (0.029 to 0.332) | 0.005 (−0.057 to 0.065) | 1.0000 |
| XEUS | layer 18, full speed: 0.165 (0.000 to 0.379) | 0.005 (−0.086 to 0.122) | 1.0000 |
| Loudness control | summary: 0.025 (0.000 to 0.058) | −0.135 (−0.308 to 0.000) | 1.0000 |
| Duration control | summary: 0.105 (0.000 to 0.298) | −0.055 (−0.271 to 0.156) | 1.0000 |

*Tree pipit, within year*: BirdNET v2.4 0.376 (0.195 to 0.564).

| Neural network | Output read, re-ID accuracy | Difference from BirdNET | Holm p |
|---|---|---|---|
| Perch 2.0 | embedding: 0.548 (0.361 to 0.738) | 0.172 (0.103 to 0.251) | < 0.0074 |
| Perch | embedding: 0.426 (0.230 to 0.628) | 0.050 (−0.054 to 0.171) | 1.0000 |
| SurfPerch | embedding: 0.455 (0.265 to 0.634) | 0.079 (−0.010 to 0.171) | 1.0000 |
| BirdNET v3 (preview) | embedding: 0.257 (0.128 to 0.417) | −0.119 (−0.304 to 0.054) | 1.0000 |
| AVEX sl-BEATs | embedding: 0.380 (0.220 to 0.543) | 0.003 (−0.125 to 0.125) | 1.0000 |
| AVEX EfficientNet-B0 | embedding: 0.281 (0.111 to 0.475) | −0.096 (−0.244 to 0.051) | 1.0000 |
| AvesEcho | embedding: 0.208 (0.075 to 0.364) | −0.168 (−0.336 to 0.024) | 1.0000 |
| AudioProtoPNet | embedding: 0.442 (0.239 to 0.657) | 0.066 (−0.087 to 0.237) | 1.0000 |
| ConvNeXt (BirdSet) | embedding: 0.502 (0.298 to 0.704) | 0.125 (0.012 to 0.265) | 0.8758 |
| Bird-MAE | embedding: 0.191 (0.055 to 0.384) | −0.185 (−0.414 to 0.042) | 1.0000 |
| ProtoCLR | embedding: 0.152 (0.025 to 0.308) | −0.224 (−0.448 to −0.012) | 1.0000 |
| RCL_FS_BSED | embedding: 0.172 (0.054 to 0.337) | −0.205 (−0.441 to 0.051) | 1.0000 |
| BirdAVES | embedding: 0.419 (0.241 to 0.607) | 0.043 (−0.101 to 0.194) | 1.0000 |
| AVES | embedding: 0.360 (0.190 to 0.554) | −0.017 (−0.217 to 0.212) | 1.0000 |
| NatureBEATs | embedding: 0.297 (0.148 to 0.452) | −0.079 (−0.200 to 0.061) | 1.0000 |
| BioLingual | embedding: 0.327 (0.179 to 0.475) | −0.050 (−0.141 to 0.053) | 1.0000 |
| BEATs | embedding: 0.195 (0.084 to 0.314) | −0.182 (−0.332 to −0.042) | 0.2380 |
| AudioMAE | embedding: 0.211 (0.137 to 0.281) | −0.165 (−0.319 to −0.020) | 0.7320 |
| VGGish | embedding: 0.116 (0.028 to 0.216) | −0.261 (−0.489 to −0.052) | 0.3776 |
| ECAPA-TDNN (speaker) | embedding: 0.244 (0.121 to 0.395) | −0.132 (−0.325 to 0.034) | 1.0000 |
| ResNet (speaker) | embedding: 0.211 (0.105 to 0.345) | −0.165 (−0.342 to 0.000) | 1.0000 |
| x-vector (speaker) | embedding: 0.300 (0.131 to 0.490) | −0.076 (−0.232 to 0.086) | 1.0000 |
| wav2vec 2.0 Base | layer 12, full speed: 0.165 (0.049 to 0.281) | −0.211 (−0.361 to −0.043) | 0.4340 |
| wav2vec 2.0 Large (robust) | layer 24, full speed: 0.162 (0.016 to 0.345) | −0.215 (−0.470 to 0.036) | 1.0000 |
| XLS-R 300M | layer 24, full speed: 0.145 (0.048 to 0.261) | −0.231 (−0.430 to −0.043) | 0.3498 |
| wav2vec 2.0 Conformer Large | layer 24, full speed: 0.277 (0.120 to 0.453) | −0.099 (−0.227 to 0.014) | 1.0000 |
| MMS 300M | layer 24, full speed: 0.178 (0.040 to 0.382) | −0.198 (−0.437 to 0.040) | 1.0000 |
| HuBERT Base | layer 12, full speed: 0.248 (0.049 to 0.474) | −0.129 (−0.315 to 0.055) | 1.0000 |
| HuBERT Large | layer 24, full speed: 0.241 (0.063 to 0.439) | −0.135 (−0.288 to −0.007) | 0.9990 |
| data2vec Base (100 h) | layer 12, full speed: 0.132 (0.047 to 0.234) | −0.244 (−0.405 to −0.101) | 0.0070 |
| data2vec Base (960 h) | layer 12, full speed: 0.112 (0.042 to 0.194) | −0.264 (−0.431 to −0.110) | < 0.0074 |

| Neural network | Output read, re-ID accuracy | Difference from BirdNET | Holm p |
|---|---|---|---|
| WavLM Base+ | layer 12, full speed: 0.274 (0.069 to 0.505) | −0.102 (−0.289 to 0.067) | 1.0000 |
| WavLM Large | layer 24, full speed: 0.234 (0.070 to 0.415) | −0.142 (−0.298 to −0.010) | 0.9128 |
| UniSpeech-SAT Base+ | layer 12, full speed: 0.248 (0.086 to 0.428) | −0.129 (−0.293 to −0.003) | 1.0000 |
| XEUS | layer 18, full speed: 0.241 (0.078 to 0.425) | −0.135 (−0.314 to 0.027) | 1.0000 |
| Loudness control | summary: 0.228 (0.032 to 0.461) | −0.149 (−0.407 to 0.140) | 1.0000 |
| Duration control | summary: 0.145 (0.000 to 0.334) | −0.231 (−0.490 to 0.023) | 1.0000 |

*Little owl, across year*: BirdNET v2.4 0.501 (0.372 to 0.631).

| Neural network | Output read, re-ID accuracy | Difference from BirdNET | Holm p |
|---|---|---|---|
| Perch 2.0 | embedding: 0.651 (0.494 to 0.786) | 0.150 (0.044 to 0.244) | 0.1840 |
| Perch | embedding: 0.646 (0.501 to 0.775) | 0.145 (0.057 to 0.229) | 0.0432 |
| SurfPerch | embedding: 0.442 (0.291 to 0.593) | −0.059 (−0.218 to 0.097) | 1.0000 |
| BirdNET v3 (preview) | embedding: 0.373 (0.237 to 0.523) | −0.128 (−0.258 to 0.003) | 0.7392 |
| AVEX sl-BEATs | embedding: 0.418 (0.238 to 0.611) | −0.084 (−0.238 to 0.067) | 1.0000 |
| AVEX EfficientNet-B0 | embedding: 0.373 (0.237 to 0.522) | −0.128 (−0.253 to −0.003) | 0.6104 |
| AvesEcho | embedding: 0.464 (0.294 to 0.629) | −0.037 (−0.193 to 0.117) | 1.0000 |
| AudioProtoPNet | embedding: 0.631 (0.472 to 0.782) | 0.130 (0.033 to 0.246) | 0.0960 |
| ConvNeXt (BirdSet) | embedding: 0.624 (0.479 to 0.754) | 0.123 (0.006 to 0.229) | 0.6104 |
| Bird-MAE | embedding: 0.251 (0.103 to 0.412) | −0.251 (−0.453 to −0.036) | 0.3424 |
| ProtoCLR | embedding: 0.172 (0.067 to 0.305) | −0.329 (−0.467 to −0.182) | < 0.0074 |
| RCL_FS_BSED | embedding: 0.189 (0.071 to 0.331) | −0.312 (−0.495 to −0.111) | 0.0936 |
| BirdAVES | embedding: 0.285 (0.155 to 0.427) | −0.216 (−0.346 to −0.089) | 0.0120 |
| AVES | embedding: 0.351 (0.209 to 0.484) | −0.150 (−0.323 to 0.019) | 0.7480 |
| NatureBEATs | embedding: 0.364 (0.215 to 0.527) | −0.138 (−0.258 to −0.031) | 0.2232 |
| BioLingual | embedding: 0.408 (0.234 to 0.598) | −0.093 (−0.256 to 0.080) | 1.0000 |
| BEATs | embedding: 0.364 (0.203 to 0.539) | −0.138 (−0.295 to 0.017) | 0.7480 |
| AudioMAE | embedding: 0.437 (0.276 to 0.598) | −0.064 (−0.226 to 0.106) | 1.0000 |
| VGGish | embedding: 0.231 (0.098 to 0.379) | −0.270 (−0.398 to −0.157) | < 0.0074 |
| ECAPA-TDNN (speaker) | embedding: 0.376 (0.256 to 0.492) | −0.125 (−0.236 to −0.027) | 0.1862 |
| ResNet (speaker) | embedding: 0.344 (0.231 to 0.456) | −0.157 (−0.308 to 0.009) | 0.7392 |
| x-vector (speaker) | embedding: 0.364 (0.220 to 0.515) | −0.138 (−0.321 to 0.068) | 1.0000 |
| wav2vec 2.0 Base | layer 12, full speed: 0.201 (0.095 to 0.325) | −0.300 (−0.442 to −0.128) | 0.0174 |
| wav2vec 2.0 Large (robust) | layer 24, full speed: 0.130 (0.028 to 0.262) | −0.371 (−0.526 to −0.215) | < 0.0074 |
| XLS-R 300M | layer 24, full speed: 0.160 (0.074 to 0.250) | −0.342 (−0.483 to −0.206) | < 0.0074 |
| wav2vec 2.0 Conformer Large | layer 24, full speed: 0.295 (0.166 to 0.433) | −0.206 (−0.371 to −0.024) | 0.4590 |
| MMS 300M | layer 24, full speed: 0.197 (0.102 to 0.298) | −0.305 (−0.479 to −0.125) | 0.0224 |
| HuBERT Base | layer 12, full speed: 0.275 (0.145 to 0.426) | −0.226 (−0.379 to −0.069) | 0.1056 |
| HuBERT Large | layer 24, full speed: 0.209 (0.097 to 0.334) | −0.292 (−0.428 to −0.155) | 0.0062 |
| data2vec Base (100 h) | layer 12, full speed: 0.258 (0.130 to 0.393) | −0.243 (−0.410 to −0.077) | 0.0966 |
| data2vec Base (960 h) | layer 12, full speed: 0.265 (0.132 to 0.414) | −0.236 (−0.406 to −0.068) | 0.1176 |
| WavLM Base+ | layer 12, full speed: 0.310 (0.176 to 0.450) | −0.192 (−0.383 to 0.019) | 0.7480 |
| WavLM Large | layer 24, full speed: 0.327 (0.174 to 0.487) | −0.174 (−0.355 to 0.024) | 0.7480 |
| UniSpeech-SAT Base+ | layer 12, full speed: 0.265 (0.127 to 0.424) | −0.236 (−0.381 to −0.080) | 0.0936 |
| XEUS | layer 18, full speed: 0.285 (0.155 to 0.424) | −0.216 (−0.389 to −0.036) | 0.3366 |
| Loudness control | summary: 0.069 (0.014 to 0.134) | −0.432 (−0.570 to −0.305) | < 0.0074 |
| Duration control | summary: 0.115 (0.000 to 0.293) | −0.386 (−0.526 to −0.243) | < 0.0074 |

*Little penguin, across night*: BirdNET v2.4 0.693 (0.607 to 0.773).

| Neural network | Output read, re-ID accuracy | Difference from BirdNET | Holm p |
|---|---|---|---|
| Perch 2.0 | embedding: 0.705 (0.599 to 0.791) | 0.011 (−0.051 to 0.069) | 1.0000 |
| Perch | embedding: 0.635 (0.528 to 0.727) | −0.058 (−0.134 to 0.013) | 1.0000 |
| SurfPerch | embedding: 0.646 (0.551 to 0.728) | −0.047 (−0.107 to 0.011) | 1.0000 |
| BirdNET v3 (preview) | embedding: 0.501 (0.371 to 0.611) | −0.192 (−0.268 to −0.124) | < 0.0074 |
| AVEX sl-BEATs | embedding: 0.731 (0.673 to 0.795) | 0.038 (−0.017 to 0.092) | 1.0000 |
| AVEX EfficientNet-B0 | embedding: 0.569 (0.444 to 0.681) | −0.124 (−0.199 to −0.051) | 0.0036 |
| AvesEcho | embedding: 0.750 (0.650 to 0.834) | 0.057 (−0.011 to 0.132) | 1.0000 |
| AudioProtoPNet | embedding: 0.674 (0.569 to 0.760) | −0.019 (−0.076 to 0.036) | 1.0000 |
| ConvNeXt (BirdSet) | embedding: 0.658 (0.538 to 0.755) | −0.035 (−0.130 to 0.052) | 1.0000 |
| Bird-MAE | embedding: 0.419 (0.271 to 0.541) | −0.274 (−0.388 to −0.171) | < 0.0074 |
| ProtoCLR | embedding: 0.447 (0.310 to 0.560) | −0.246 (−0.366 to −0.132) | < 0.0074 |
| RCL_FS_BSED | embedding: 0.622 (0.532 to 0.711) | −0.071 (−0.117 to −0.022) | 0.0728 |
| BirdAVES | embedding: 0.634 (0.517 to 0.731) | −0.059 (−0.140 to 0.018) | 1.0000 |
| AVES | embedding: 0.670 (0.553 to 0.769) | −0.023 (−0.096 to 0.048) | 1.0000 |
| NatureBEATs | embedding: 0.759 (0.692 to 0.826) | 0.066 (−0.001 to 0.138) | 0.6336 |
| BioLingual | embedding: 0.617 (0.500 to 0.724) | −0.076 (−0.145 to −0.006) | 0.4392 |
| BEATs | embedding: 0.569 (0.444 to 0.681) | −0.124 (−0.202 to −0.051) | 0.0090 |
| AudioMAE | embedding: 0.528 (0.398 to 0.636) | −0.165 (−0.246 to −0.099) | < 0.0074 |
| VGGish | embedding: 0.463 (0.341 to 0.576) | −0.230 (−0.300 to −0.166) | < 0.0074 |
| ECAPA-TDNN (speaker) | embedding: 0.442 (0.324 to 0.552) | −0.251 (−0.327 to −0.182) | < 0.0074 |
| ResNet (speaker) | embedding: 0.423 (0.302 to 0.538) | −0.270 (−0.373 to −0.170) | < 0.0074 |
| x-vector (speaker) | embedding: 0.441 (0.317 to 0.544) | −0.253 (−0.354 to −0.155) | < 0.0074 |
| wav2vec 2.0 Base | layer 12, full speed: 0.475 (0.340 to 0.601) | −0.218 (−0.308 to −0.126) | 0.0036 |
| wav2vec 2.0 Large (robust) | layer 24, full speed: 0.237 (0.118 to 0.351) | −0.456 (−0.559 to −0.362) | < 0.0074 |
| XLS-R 300M | layer 24, full speed: 0.510 (0.356 to 0.638) | −0.183 (−0.284 to −0.093) | < 0.0074 |
| wav2vec 2.0 Conformer Large | layer 24, full speed: 0.432 (0.295 to 0.560) | −0.261 (−0.385 to −0.151) | < 0.0074 |
| MMS 300M | layer 24, full speed: 0.577 (0.413 to 0.720) | −0.116 (−0.219 to −0.022) | 0.1742 |
| HuBERT Base | layer 12, full speed: 0.431 (0.305 to 0.557) | −0.263 (−0.356 to −0.165) | 0.0036 |
| HuBERT Large | layer 24, full speed: 0.434 (0.295 to 0.561) | −0.259 (−0.357 to −0.161) | < 0.0074 |
| data2vec Base (100 h) | layer 12, full speed: 0.356 (0.209 to 0.484) | −0.337 (−0.440 to −0.250) | < 0.0074 |
| data2vec Base (960 h) | layer 12, full speed: 0.289 (0.203 to 0.368) | −0.404 (−0.486 to −0.327) | < 0.0074 |
| WavLM Base+ | layer 12, full speed: 0.399 (0.281 to 0.516) | −0.294 (−0.391 to −0.192) | < 0.0074 |
| WavLM Large | layer 24, full speed: 0.460 (0.322 to 0.585) | −0.234 (−0.327 to −0.142) | < 0.0074 |
| UniSpeech-SAT Base+ | layer 12, full speed: 0.391 (0.239 to 0.543) | −0.302 (−0.417 to −0.182) | < 0.0074 |
| XEUS | layer 18, full speed: 0.601 (0.445 to 0.743) | −0.092 (−0.207 to 0.021) | 1.0000 |
| Loudness control | summary: 0.159 (0.048 to 0.267) | −0.534 (−0.659 to −0.411) | < 0.0074 |
| Duration control | summary: 0.167 (0.000 to 0.394) | −0.527 (−0.738 to −0.287) | < 0.0074 |

*Red-tailed black cockatoo, random split*: BirdNET v2.4 0.961 (0.930 to 0.980).

| Neural network | Output read, re-ID accuracy | Difference from BirdNET | Holm p |
|---|---|---|---|
| Perch 2.0 | embedding: 0.961 (0.933 to 0.979) | 0.000 (−0.015 to 0.023) | 1.0000 |
| Perch | embedding: 0.947 (0.920 to 0.973) | −0.014 (−0.043 to 0.029) | 1.0000 |
| SurfPerch | embedding: 0.958 (0.920 to 0.981) | −0.003 (−0.019 to 0.017) | 1.0000 |

| Neural network | Output read, re-ID accuracy | Difference from BirdNET | Holm p |
|---|---|---|---|
| BirdNET v3 (preview) | embedding: 0.854 (0.791 to 0.892) | −0.106 (−0.158 to −0.070) | < 0.0074 |
| AVEX sl-BEATs | embedding: 0.936 (0.894 to 0.964) | −0.025 (−0.053 to 0.000) | 0.5742 |
| AVEX EfficientNet-B0 | embedding: 0.891 (0.839 to 0.925) | −0.070 (−0.107 to −0.034) | 0.0048 |
| AvesEcho | embedding: 0.969 (0.926 to 0.993) | 0.008 (−0.020 to 0.032) | 1.0000 |
| AudioProtoPNet | embedding: 0.941 (0.910 to 0.967) | −0.020 (−0.050 to 0.023) | 1.0000 |
| ConvNeXt (BirdSet) | embedding: 0.969 (0.936 to 0.991) | 0.008 (−0.019 to 0.045) | 1.0000 |
| Bird-MAE | embedding: 0.527 (0.169 to 0.753) | −0.434 (−0.778 to −0.216) | < 0.0074 |
| ProtoCLR | embedding: 0.515 (0.219 to 0.713) | −0.445 (−0.726 to −0.259) | < 0.0074 |
| RCL_FS_BSED | embedding: 0.709 (0.436 to 0.841) | −0.252 (−0.510 to −0.122) | < 0.0074 |
| BirdAVES | embedding: 0.947 (0.893 to 0.980) | −0.014 (−0.050 to 0.014) | 1.0000 |
| AVES | embedding: 0.924 (0.848 to 0.963) | −0.036 (−0.092 to −0.010) | 0.0154 |
| NatureBEATs | embedding: 0.944 (0.873 to 0.978) | −0.017 (−0.070 to 0.013) | 1.0000 |
| BioLingual | embedding: 0.933 (0.886 to 0.960) | −0.028 (−0.057 to −0.004) | 0.3060 |
| BEATs | embedding: 0.908 (0.845 to 0.945) | −0.053 (−0.091 to −0.029) | < 0.0074 |
| AudioMAE | embedding: 0.782 (0.615 to 0.871) | −0.179 (−0.327 to −0.101) | < 0.0074 |
| VGGish | embedding: 0.812 (0.680 to 0.902) | −0.148 (−0.258 to −0.071) | < 0.0074 |
| ECAPA-TDNN (speaker) | embedding: 0.725 (0.614 to 0.796) | −0.235 (−0.328 to −0.166) | < 0.0074 |
| ResNet (speaker) | embedding: 0.812 (0.752 to 0.849) | −0.148 (−0.187 to −0.116) | < 0.0074 |
| x-vector (speaker) | embedding: 0.796 (0.652 to 0.868) | −0.165 (−0.290 to −0.100) | < 0.0074 |
| wav2vec 2.0 Base | layer 12, full speed: 0.768 (0.676 to 0.824) | −0.193 (−0.266 to −0.144) | < 0.0074 |
| wav2vec 2.0 Large (robust) | layer 24, full speed: 0.499 (0.237 to 0.705) | −0.462 (−0.712 to −0.266) | < 0.0074 |
| XLS-R 300M | layer 24, full speed: 0.737 (0.519 to 0.859) | −0.224 (−0.424 to −0.109) | < 0.0074 |
| wav2vec 2.0 Conformer Large | layer 24, full speed: 0.739 (0.591 to 0.831) | −0.221 (−0.345 to −0.141) | < 0.0074 |
| MMS 300M | layer 24, full speed: 0.742 (0.548 to 0.854) | −0.218 (−0.401 to −0.114) | < 0.0074 |
| HuBERT Base | layer 12, full speed: 0.798 (0.675 to 0.872) | −0.162 (−0.261 to −0.103) | < 0.0074 |
| HuBERT Large | layer 24, full speed: 0.641 (0.422 to 0.818) | −0.319 (−0.524 to −0.154) | < 0.0074 |
| data2vec Base (100 h) | layer 12, full speed: 0.616 (0.384 to 0.770) | −0.345 (−0.560 to −0.205) | < 0.0074 |
| data2vec Base (960 h) | layer 12, full speed: 0.655 (0.443 to 0.791) | −0.305 (−0.497 to −0.184) | < 0.0074 |
| WavLM Base+ | layer 12, full speed: 0.826 (0.733 to 0.887) | −0.134 (−0.204 to −0.087) | < 0.0074 |
| WavLM Large | layer 24, full speed: 0.832 (0.693 to 0.913) | −0.129 (−0.243 to −0.063) | < 0.0074 |
| UniSpeech-SAT Base+ | layer 12, full speed: 0.812 (0.680 to 0.902) | −0.148 (−0.260 to −0.070) | < 0.0074 |
| XEUS | layer 18, full speed: 0.863 (0.753 to 0.927) | −0.098 (−0.190 to −0.046) | < 0.0074 |
| Loudness control | summary: 0.406 (0.153 to 0.587) | −0.555 (−0.793 to −0.375) | < 0.0074 |
| Duration control | summary: 0.336 (0.000 to 0.666) | −0.625 (−0.958 to −0.304) | < 0.0074 |

*Egyptian fruit bat, across year*: BirdNET v2.4 0.381 (0.189 to 0.561).

| Neural network | Output read, re-ID accuracy | Difference from BirdNET | Holm p |
|---|---|---|---|
| Perch 2.0 | embedding: 0.350 (0.161 to 0.532) | −0.031 (−0.115 to 0.104) | 1.0000 |
| Perch | embedding: 0.268 (0.131 to 0.537) | −0.113 (−0.353 to 0.180) | 1.0000 |
| SurfPerch | embedding: 0.353 (0.202 to 0.496) | −0.028 (−0.136 to 0.104) | 1.0000 |
| BirdNET v3 (preview) | embedding: 0.293 (0.194 to 0.475) | −0.088 (−0.272 to 0.103) | 1.0000 |
| AVEX sl-BEATs | embedding: 0.371 (0.233 to 0.490) | −0.011 (−0.110 to 0.083) | 1.0000 |
| AVEX EfficientNet-B0 | embedding: 0.307 (0.258 to 0.359) | −0.074 (−0.237 to 0.088) | 1.0000 |
| AvesEcho | embedding: 0.223 (0.160 to 0.292) | −0.158 (−0.299 to −0.027) | 0.0504 |
| AudioProtoPNet | embedding: 0.305 (0.131 to 0.512) | −0.077 (−0.189 to 0.100) | 1.0000 |
| ConvNeXt (BirdSet) | embedding: 0.333 (0.127 to 0.530) | −0.048 (−0.121 to 0.061) | 1.0000 |

| Neural network | Output read, re-ID accuracy | Difference from BirdNET | Holm p |
|---|---|---|---|
| Bird-MAE | embedding: 0.121 (0.054 to 0.218) | −0.261 (−0.467 to −0.056) | < 0.0074 |
| ProtoCLR | embedding: 0.133 (0.052 to 0.233) | −0.249 (−0.494 to −0.001) | 1.0000 |
| RCL_FS_BSED | embedding: 0.191 (0.077 to 0.327) | −0.191 (−0.467 to 0.056) | 1.0000 |
| BirdAVES | embedding: 0.347 (0.142 to 0.540) | −0.034 (−0.124 to 0.019) | 1.0000 |
| AVES | embedding: 0.287 (0.055 to 0.517) | −0.095 (−0.275 to 0.011) | 1.0000 |
| NatureBEATs | embedding: 0.337 (0.107 to 0.579) | −0.044 (−0.122 to 0.027) | 1.0000 |
| BioLingual | embedding: 0.209 (0.122 to 0.319) | −0.172 (−0.417 to 0.054) | 1.0000 |
| BEATs | embedding: 0.248 (0.173 to 0.361) | −0.134 (−0.358 to 0.052) | 1.0000 |
| AudioMAE | embedding: 0.182 (0.111 to 0.300) | −0.199 (−0.430 to 0.027) | 1.0000 |
| VGGish | embedding: 0.166 (0.081 to 0.216) | −0.216 (−0.370 to −0.033) | 0.5250 |
| ECAPA-TDNN (speaker) | embedding: 0.258 (0.183 to 0.354) | −0.124 (−0.333 to 0.080) | 1.0000 |
| ResNet (speaker) | embedding: 0.319 (0.204 to 0.412) | −0.062 (−0.244 to 0.144) | 1.0000 |
| x-vector (speaker) | embedding: 0.331 (0.201 to 0.437) | −0.051 (−0.151 to 0.061) | 1.0000 |
| wav2vec 2.0 Base | layer 12, full speed: 0.237 (0.015 to 0.376) | −0.144 (−0.367 to 0.109) | 1.0000 |
| wav2vec 2.0 Large (robust) | layer 24, full speed: 0.210 (0.046 to 0.395) | −0.171 (−0.462 to 0.167) | 1.0000 |
| XLS-R 300M | layer 24, full speed: 0.248 (0.037 to 0.406) | −0.134 (−0.328 to 0.161) | 1.0000 |
| wav2vec 2.0 Conformer Large | layer 24, full speed: 0.287 (0.082 to 0.422) | −0.095 (−0.289 to 0.072) | 1.0000 |
| MMS 300M | layer 24, full speed: 0.204 (0.091 to 0.292) | −0.178 (−0.340 to 0.019) | 1.0000 |
| HuBERT Base | layer 12, full speed: 0.309 (0.054 to 0.495) | −0.072 (−0.279 to 0.086) | 1.0000 |
| HuBERT Large | layer 24, full speed: 0.232 (0.091 to 0.393) | −0.150 (−0.405 to 0.176) | 1.0000 |
| data2vec Base (100 h) | layer 12, full speed: 0.219 (0.086 to 0.378) | −0.163 (−0.441 to 0.157) | 1.0000 |
| data2vec Base (960 h) | layer 12, full speed: 0.225 (0.097 to 0.365) | −0.156 (−0.396 to 0.117) | 1.0000 |
| WavLM Base+ | layer 12, full speed: 0.268 (0.100 to 0.405) | −0.113 (−0.319 to 0.156) | 1.0000 |
| WavLM Large | layer 24, full speed: 0.309 (0.138 to 0.405) | −0.072 (−0.223 to 0.131) | 1.0000 |
| UniSpeech-SAT Base+ | layer 12, full speed: 0.323 (0.146 to 0.430) | −0.058 (−0.231 to 0.164) | 1.0000 |
| XEUS | layer 18, full speed: 0.287 (0.052 to 0.458) | −0.095 (−0.251 to 0.044) | 1.0000 |
| Loudness control | summary: 0.248 (0.013 to 0.620) | −0.134 (−0.531 to 0.395) | 1.0000 |
| Duration control | summary: 0.208 (0.000 to 0.462) | −0.173 (−0.537 to 0.249) | 1.0000 |

*Chiffchaff, within year*: BirdNET v2.4 0.765 (0.680 to 0.841).

| Neural network | Output read, re-ID accuracy | Difference from BirdNET | Holm p |
|---|---|---|---|
| Perch 2.0 | embedding: 0.831 (0.756 to 0.884) | 0.066 (−0.008 to 0.122) | 0.5782 |
| Perch | embedding: 0.778 (0.656 to 0.858) | 0.013 (−0.105 to 0.094) | 1.0000 |
| SurfPerch | embedding: 0.729 (0.632 to 0.808) | −0.036 (−0.115 to 0.013) | 0.8892 |
| BirdNET v3 (preview) | embedding: 0.624 (0.370 to 0.770) | −0.141 (−0.381 to −0.003) | 0.3504 |
| AVEX sl-BEATs | embedding: 0.655 (0.572 to 0.748) | −0.110 (−0.161 to −0.058) | < 0.0074 |
| AVEX EfficientNet-B0 | embedding: 0.691 (0.577 to 0.760) | −0.074 (−0.173 to −0.021) | 0.0440 |
| AvesEcho | embedding: 0.691 (0.518 to 0.772) | −0.073 (−0.217 to −0.008) | 0.2400 |
| AudioProtoPNet | embedding: 0.806 (0.646 to 0.890) | 0.042 (−0.096 to 0.114) | 1.0000 |
| ConvNeXt (BirdSet) | embedding: 0.805 (0.687 to 0.866) | 0.041 (−0.066 to 0.089) | 1.0000 |
| Bird-MAE | embedding: 0.341 (0.145 to 0.470) | −0.424 (−0.603 to −0.323) | < 0.0074 |
| ProtoCLR | embedding: 0.325 (0.211 to 0.477) | −0.439 (−0.548 to −0.305) | < 0.0074 |
| RCL_FS_BSED | embedding: 0.256 (0.121 to 0.494) | −0.508 (−0.643 to −0.289) | < 0.0074 |
| BirdAVES | embedding: 0.749 (0.683 to 0.812) | −0.016 (−0.083 to 0.050) | 1.0000 |
| AVES | embedding: 0.515 (0.396 to 0.645) | −0.250 (−0.329 to −0.168) | < 0.0074 |
| NatureBEATs | embedding: 0.739 (0.558 to 0.824) | −0.026 (−0.164 to 0.042) | 1.0000 |

| Neural network | Output read, re-ID accuracy | Difference from BirdNET | Holm p |
|---|---|---|---|
| BioLingual | embedding: 0.659 (0.452 to 0.769) | −0.106 (−0.283 to −0.008) | 0.2610 |
| BEATs | embedding: 0.514 (0.310 to 0.624) | −0.251 (−0.432 to −0.163) | < 0.0074 |
| AudioMAE | embedding: 0.570 (0.494 to 0.694) | −0.195 (−0.271 to −0.083) | 0.0072 |
| VGGish | embedding: 0.458 (0.256 to 0.561) | −0.307 (−0.501 to −0.223) | < 0.0074 |
| ECAPA-TDNN (speaker) | embedding: 0.437 (0.309 to 0.531) | −0.328 (−0.441 to −0.263) | < 0.0074 |
| ResNet (speaker) | embedding: 0.392 (0.273 to 0.534) | −0.373 (−0.470 to −0.271) | < 0.0074 |
| x-vector (speaker) | embedding: 0.385 (0.258 to 0.523) | −0.379 (−0.470 to −0.290) | < 0.0074 |
| wav2vec 2.0 Base | layer 12, full speed: 0.347 (0.251 to 0.480) | −0.418 (−0.502 to −0.314) | < 0.0074 |
| wav2vec 2.0 Large (robust) | layer 24, full speed: 0.274 (0.070 to 0.397) | −0.491 (−0.698 to −0.376) | < 0.0074 |
| XLS-R 300M | layer 24, full speed: 0.384 (0.193 to 0.526) | −0.381 (−0.553 to −0.272) | < 0.0074 |
| wav2vec 2.0 Conformer Large | layer 24, full speed: 0.501 (0.351 to 0.592) | −0.263 (−0.412 to −0.185) | < 0.0074 |
| MMS 300M | layer 24, full speed: 0.349 (0.187 to 0.499) | −0.416 (−0.559 to −0.301) | < 0.0074 |
| HuBERT Base | layer 12, full speed: 0.492 (0.329 to 0.584) | −0.273 (−0.415 to −0.201) | < 0.0074 |
| HuBERT Large | layer 24, full speed: 0.469 (0.313 to 0.575) | −0.296 (−0.455 to −0.203) | < 0.0074 |
| data2vec Base (100 h) | layer 12, full speed: 0.316 (0.176 to 0.451) | −0.449 (−0.573 to −0.348) | < 0.0074 |
| data2vec Base (960 h) | layer 12, full speed: 0.310 (0.208 to 0.424) | −0.454 (−0.568 to −0.349) | < 0.0074 |
| WavLM Base+ | layer 12, full speed: 0.485 (0.328 to 0.575) | −0.280 (−0.425 to −0.211) | < 0.0074 |
| WavLM Large | layer 24, full speed: 0.585 (0.426 to 0.682) | −0.179 (−0.307 to −0.113) | < 0.0074 |
| UniSpeech-SAT Base+ | layer 12, full speed: 0.498 (0.346 to 0.594) | −0.267 (−0.399 to −0.201) | < 0.0074 |
| XEUS | layer 18, full speed: 0.440 (0.290 to 0.576) | −0.324 (−0.444 to −0.235) | < 0.0074 |
| Loudness control | summary: 0.094 (0.010 to 0.267) | −0.671 (−0.747 to −0.522) | < 0.0074 |
| Duration control | summary: 0.040 (0.000 to 0.147) | −0.725 (−0.794 to −0.607) | < 0.0074 |

*Rook, across year*: BirdNET v2.4 0.375 (0.261 to 0.506).

| Neural network | Output read, re-ID accuracy | Difference from BirdNET | Holm p |
|---|---|---|---|
| Perch 2.0 | embedding: 0.418 (0.306 to 0.535) | 0.043 (0.017 to 0.065) | 0.0240 |
| Perch | embedding: 0.393 (0.282 to 0.516) | 0.018 (−0.027 to 0.062) | 0.9816 |
| SurfPerch | embedding: 0.306 (0.200 to 0.425) | −0.069 (−0.106 to −0.038) | 0.0038 |
| BirdNET v3 (preview) | embedding: 0.328 (0.212 to 0.439) | −0.048 (−0.098 to −0.008) | 0.1632 |
| AVEX sl-BEATs | embedding: 0.332 (0.191 to 0.444) | −0.043 (−0.119 to 0.011) | 0.6640 |
| AVEX EfficientNet-B0 | embedding: 0.345 (0.247 to 0.442) | −0.031 (−0.073 to −0.001) | 0.3010 |
| AvesEcho | embedding: 0.343 (0.223 to 0.437) | −0.033 (−0.125 to 0.024) | 0.9816 |
| AudioProtoPNet | embedding: 0.386 (0.269 to 0.506) | 0.011 (−0.008 to 0.024) | 0.9248 |
| ConvNeXt (BirdSet) | embedding: 0.409 (0.260 to 0.536) | 0.034 (−0.044 to 0.083) | 0.9816 |
| Bird-MAE | embedding: 0.194 (0.065 to 0.365) | −0.182 (−0.255 to −0.107) | < 0.0074 |
| ProtoCLR | embedding: 0.084 (0.018 to 0.219) | −0.292 (−0.374 to −0.165) | < 0.0074 |
| RCL_FS_BSED | embedding: 0.105 (0.032 to 0.215) | −0.270 (−0.350 to −0.178) | < 0.0074 |
| BirdAVES | embedding: 0.174 (0.053 to 0.384) | −0.202 (−0.305 to −0.073) | 0.0120 |
| AVES | embedding: 0.176 (0.058 to 0.379) | −0.199 (−0.284 to −0.087) | 0.0096 |
| NatureBEATs | embedding: 0.314 (0.154 to 0.475) | −0.061 (−0.136 to −0.0004) | 0.3010 |
| BioLingual | embedding: 0.303 (0.186 to 0.404) | −0.073 (−0.139 to −0.025) | 0.0120 |
| BEATs | embedding: 0.315 (0.192 to 0.428) | −0.060 (−0.107 to −0.034) | < 0.0074 |
| AudioMAE | embedding: 0.236 (0.119 to 0.411) | −0.140 (−0.224 to −0.047) | 0.0198 |
| VGGish | embedding: 0.180 (0.086 to 0.329) | −0.195 (−0.266 to −0.117) | < 0.0074 |
| ECAPA-TDNN (speaker) | embedding: 0.254 (0.145 to 0.402) | −0.121 (−0.161 to −0.076) | < 0.0074 |
| ResNet (speaker) | embedding: 0.239 (0.125 to 0.435) | −0.137 (−0.221 to −0.035) | 0.0414 |

| Neural network | Output read, re-ID accuracy | Difference from BirdNET | Holm p |
|---|---|---|---|
| x-vector (speaker) | embedding: 0.262 (0.136 to 0.417) | −0.114 (−0.163 to −0.064) | < 0.0074 |
| wav2vec 2.0 Base | layer 12, full speed: 0.157 (0.095 to 0.261) | −0.218 (−0.301 to −0.132) | < 0.0074 |
| wav2vec 2.0 Large (robust) | layer 24, full speed: 0.125 (0.012 to 0.267) | −0.250 (−0.409 to −0.089) | 0.0120 |
| XLS-R 300M | layer 24, full speed: 0.177 (0.060 to 0.373) | −0.198 (−0.297 to −0.089) | < 0.0074 |
| wav2vec 2.0 Conformer Large | layer 24, full speed: 0.196 (0.105 to 0.302) | −0.179 (−0.234 to −0.138) | < 0.0074 |
| MMS 300M | layer 24, full speed: 0.184 (0.039 to 0.375) | −0.191 (−0.312 to −0.076) | < 0.0074 |
| HuBERT Base | layer 12, full speed: 0.194 (0.077 to 0.378) | −0.181 (−0.268 to −0.084) | 0.0038 |
| HuBERT Large | layer 24, full speed: 0.190 (0.086 to 0.335) | −0.185 (−0.269 to −0.114) | < 0.0074 |
| data2vec Base (100 h) | layer 12, full speed: 0.158 (0.061 to 0.300) | −0.218 (−0.293 to −0.147) | < 0.0074 |
| data2vec Base (960 h) | layer 12, full speed: 0.147 (0.045 to 0.318) | −0.229 (−0.330 to −0.126) | < 0.0074 |
| WavLM Base+ | layer 12, full speed: 0.121 (0.038 to 0.298) | −0.254 (−0.351 to −0.139) | < 0.0074 |
| WavLM Large | layer 24, full speed: 0.163 (0.041 to 0.372) | −0.212 (−0.309 to −0.090) | 0.0120 |
| UniSpeech-SAT Base+ | layer 12, full speed: 0.164 (0.063 to 0.356) | −0.211 (−0.309 to −0.097) | < 0.0074 |
| XEUS | layer 18, full speed: 0.218 (0.112 to 0.387) | −0.157 (−0.241 to −0.073) | < 0.0074 |
| Loudness control | summary: 0.126 (0.019 to 0.342) | −0.249 (−0.363 to −0.108) | < 0.0074 |
| Duration control | summary: 0.030 (0.000 to 0.113) | −0.345 (−0.467 to −0.210) | 0.0038 |

*Notes:* Own final output: the output embedding for networks trained on animal or general sound, the last transformer layer at the recording's own speed for speech networks, and the output at the recording's own speed for speaker networks; networks with one embedding are as in Table S13. Difference and Holm p as in Table S13. Against the layer chosen by the Fisher rule, the final output was higher in 90 of 130 network-dataset pairs for the ten animal networks with several layers (median change +0.069) and in 53 of 169 for the 13 speech networks with several layers (median −0.041).

#### Appendix S2. Follow-up sensitivity analyses after review

These follow-up analyses were specified after review, before their execution; they were not preregistered. The fixed protocol was committed as 84c26f1664d95eea36923d7b826bf1a0ebf7f433; its SHA-256 is 22d2276709936df17b48ef208e5211dff7b3c69838afe4d3d9856242f927b0ec. The open-set runner was corrected to retain the original rook enrolment cap before any rook result was written (commit 7d52438). All 194 new outputs were copied to the content-addressed GCS archive with checksum verification. Public aggregate projections, source hashes, split hashes and runtime inventories are included in the v10 evidence deposit (https://doi.org/10.5281/zenodo.23174588). Software 0.3.0 is archived at https://doi.org/10.5281/zenodo.23167850. Recordings, embeddings, real identities and per-clip observations are not redistributed.

##### S2.1 Count-matched clip splits

Randomisation describes how units are assigned; it does not define the unit. Randomly assigning whole recording sessions can be consistent with a valid session-disjoint evaluation. Randomly assigning individual clips can instead mix the context of a session between enrolment and test. Which design is appropriate depends on the target use. Recognition of a familiar animal in a later session requires evaluation on sessions untouched by training and model selection.

For each species and encoder, both arms used exactly the published train/test clip pool: chiffchaff 5,107/1,131 enrolment/test clips, little owl 545/407 and tree pipit 409/303. The tree-pipit more/years entries were excluded in both arms. Each bird retained its own enrolment/test counts. Ten fixed seeds (101–110) shuffled clips within bird. Both arms used the same frozen vectors, enrolment-only preprocessing, three classifier rules and ridge penalty 1.0. The paired bootstrap resampled individuals jointly over the session arm and mean of the ten random arms (10,000 draws, seed 17); its intervals condition on those ten fixed randomisations. They are not intervals over all possible split choices. All 18 mean differences were positive, from 0.196 to 0.508, with every interval excluding zero.

Detailed results: values below give session accuracy; mean random-clip accuracy [minimum–maximum across ten seeds]; mean difference [paired 95% interval]; and balanced accuracy, session → random mean. Balanced accuracy is mean recall over birds.

littleowl, birdnet, kernel ridge: 0.477; 0.963 [0.953–0.973]; 0.486 [0.359–0.606]; balanced 0.489 → 0.961.

littleowl, birdnet, class mean: 0.430; 0.897 [0.880–0.912]; 0.467 [0.343–0.582]; balanced 0.452 → 0.894.

littleowl, birdnet, nearest clip: 0.577; 0.973 [0.961–0.980]; 0.396 [0.249–0.546]; balanced 0.570 → 0.973.

littleowl, google-perch, kernel ridge: 0.631; 0.964 [0.958–0.973]; 0.333 [0.201–0.479]; balanced 0.625 → 0.964.

littleowl, google-perch, class mean: 0.543; 0.879 [0.865–0.899]; 0.336 [0.203–0.466]; balanced 0.560 → 0.876.

littleowl, google-perch, nearest clip: 0.582; 0.963 [0.961–0.971]; 0.381 [0.229–0.537]; balanced 0.591 → 0.964.

chiffchaff, birdnet, kernel ridge: 0.734; 0.969 [0.963–0.975]; 0.235 [0.147–0.315]; balanced 0.742 → 0.947.

chiffchaff, birdnet, class mean: 0.502; 0.810 [0.792–0.827]; 0.308 [0.219–0.425]; balanced 0.515 → 0.796.

chiffchaff, birdnet, nearest clip: 0.595; 0.941 [0.930–0.953]; 0.346 [0.253–0.434]; balanced 0.602 → 0.916.

chiffchaff, google-perch, kernel ridge: 0.751; 0.947 [0.935–0.957]; 0.196 [0.120–0.270]; balanced 0.735 → 0.909.

chiffchaff, google-perch, class mean: 0.404; 0.622 [0.599–0.653]; 0.218 [0.128–0.276]; balanced 0.456 → 0.626.

chiffchaff, google-perch, nearest clip: 0.486; 0.818 [0.809–0.842]; 0.332 [0.225–0.389]; balanced 0.494 → 0.785.

pipit, birdnet, kernel ridge: 0.419; 0.927 [0.904–0.947]; 0.508 [0.328–0.688]; balanced 0.404 → 0.922.

pipit, birdnet, class mean: 0.363; 0.782 [0.746–0.815]; 0.418 [0.291–0.548]; balanced 0.342 → 0.772.

pipit, birdnet, nearest clip: 0.406; 0.865 [0.835–0.908]; 0.459 [0.325–0.592]; balanced 0.395 → 0.863.

pipit, google-perch, kernel ridge: 0.452; 0.910 [0.888–0.944]; 0.458 [0.275–0.635]; balanced 0.445 → 0.906.

pipit, google-perch, class mean: 0.277; 0.717 [0.653–0.772]; 0.440 [0.339–0.543]; balanced 0.270 → 0.709.

pipit, google-perch, nearest clip: 0.356; 0.771 [0.749–0.809]; 0.415 [0.263–0.569]; balanced 0.356 → 0.767.

The median cosine similarity to the nearest enrolment clip from the same bird was higher in every randomisation for each of the six species–encoder combinations. This is a descriptive measure on raw unit-normalised vectors, not proof of near-duplicate calls or a causal estimate of the contribution of background, vocal drift or recording conditions. The retained metadata identify original train/test blocks, not finer recording sessions; we do not infer missing session labels.

##### S2.2 Original published-fold comparisons retained

The following are the original Table 3 comparisons, preserved for transparency. Unlike S2.1, the published little-owl fold used 761 rather than 545 enrolment clips, and the tree-pipit fold used additional clips. They therefore do not isolate split composition from training size or clip pool. For each row: original random-fold accuracy; session accuracy; original paired difference [95% interval]. The original rook comparison used one count-matched random split, not the ten-seed procedure.

Little owl, BirdNET, kernel ridge: 0.984; 0.477; 0.508 (0.378 to 0.626).

Little owl, BirdNET, class mean: 0.927; 0.430; 0.497 (0.365 to 0.609).

Little owl, BirdNET, nearest clip: 0.979; 0.577; 0.402 (0.265 to 0.542).

Little owl, Perch, kernel ridge: 0.990; 0.631; 0.358 (0.222 to 0.507).

Little owl, Perch, class mean: 0.880; 0.543; 0.337 (0.196 to 0.474).

Little owl, Perch, nearest clip: 0.979; 0.582; 0.397 (0.249 to 0.548).

Chiffchaff, BirdNET, kernel ridge: 0.961; 0.734; 0.227 (0.149 to 0.309).

Chiffchaff, BirdNET, class mean: 0.800; 0.502; 0.298 (0.219 to 0.444).

Chiffchaff, BirdNET, nearest clip: 0.935; 0.595; 0.340 (0.253 to 0.429).

Chiffchaff, Perch, kernel ridge: 0.944; 0.751; 0.193 (0.125 to 0.290).

Chiffchaff, Perch, class mean: 0.591; 0.404; 0.187 (0.099 to 0.287).

Chiffchaff, Perch, nearest clip: 0.819; 0.486; 0.333 (0.243 to 0.403).

Tree pipit (not the same clips), BirdNET, kernel ridge: 0.875; 0.419; 0.456 (0.270 to 0.645).

Tree pipit (not the same clips), BirdNET, class mean: 0.670; 0.363; 0.307 (0.145 to 0.463).

Tree pipit (not the same clips), BirdNET, nearest clip: 0.839; 0.406; 0.433 (0.307 to 0.558).

Tree pipit (not the same clips), Perch, kernel ridge: 0.894; 0.452; 0.442 (0.249 to 0.629).

Tree pipit (not the same clips), Perch, class mean: 0.575; 0.277; 0.298 (0.212 to 0.403).

Tree pipit (not the same clips), Perch, nearest clip: 0.700; 0.356; 0.343 (0.198 to 0.484).

Rook, shared aviary, BirdNET (this study), kernel ridge: 0.547; 0.422; 0.126 (0.065 to 0.172).

##### S2.3 Background accuracy and class imbalance

These are the retained BirdNET kernel-ridge background/call results, not new model inference. Balanced accuracy gives equal weight to each bird. The query-majority reference is the fraction of clips belonging to the commonest test identity; it is a descriptive best-constant reference using test labels. The separate fitted constant baseline always predicts the most frequent enrolment identity. Ties are resolved by the fixed anonymous identity ordering. These two quantities answer different questions.

chiffchaff-withinyear: foreground: ordinary 0.765, balanced 0.791, query-majority 0.393, enrolment-majority predictor 0.062 (n=1131); background: ordinary 0.737, balanced 0.714, query-majority 0.395, enrolment-majority predictor 0.063 (n=1100).

chiffchaff-acrossyear: foreground: ordinary 0.160, balanced 0.151, query-majority 0.160, enrolment-majority predictor 0.100 (n=200); background: ordinary 0.132, balanced 0.143, query-majority 0.152, enrolment-majority predictor 0.096 (n=197).

littleowl-acrossyear: foreground: ordinary 0.501, balanced 0.518, query-majority 0.084, enrolment-majority predictor 0.044 (n=407); background: ordinary 0.105, balanced 0.108, query-majority 0.086, enrolment-majority predictor 0.044 (n=409).

pipit-withinyear: foreground: ordinary 0.376, balanced 0.362, query-majority 0.125, enrolment-majority predictor 0.116 (n=303); background: ordinary 0.177, balanced 0.177, query-majority 0.126, enrolment-majority predictor 0.116 (n=293).

pipit-acrossyear: foreground: ordinary 0.214, balanced 0.218, query-majority 0.160, enrolment-majority predictor 0.073 (n=313); background: ordinary 0.065, balanced 0.056, query-majority 0.160, enrolment-majority predictor 0.072 (n=306).

Within-year chiffchaff background identification remains substantial under balanced accuracy (0.714 against 0.791 for calls). The two balanced accuracies should not be described as equivalent. Calls and backgrounds have slightly different test counts and therefore separate majority references.

#### S2.4 Open-set scoring and calibration-size sensitivity

The original mean-profile scorer was compared with a nearest-enrolment-clip scorer, using the same frozen selected embeddings and all 16 original allocations. Each scorer independently calibrated its own threshold using only the calibration strangers. The nearest-clip scorer assigns the identity of the most similar enrolled clip; no test clip enters its gallery. Model selection, preprocessing, role assignments and stranger budgets were unchanged. The original 90-clip-per-bird rook enrolment cap was retained. Recomputed mean-profile closed-set accuracy and 10% budget known-correct acceptance reproduced the archived values in every allocation. Both scorers are reported; the better test result is not a deployment-selected model.

Across the 36 distinct neural networks, the highest median accepted-and-correctly-named proportion at a 10% calibration stranger budget was 0.374 with mean profiles and 0.462 with nearest-clip scoring on the four wild session-disjoint endpoints. Both maxima were Perch 2.0 on little owl. For that nearest-clip comparison, the range over allocations was 0.000–0.596, and the actual test-stranger acceptance rate had median 0.099 and range 0.000–0.615. This improvement is conditional on the tested models, scorers, roles and data; neither number is a ceiling on future re-identification methods. Nearest-clip scoring did not improve every model and dataset: BirdNET on rook fell from 0.080 to 0.037.

Detailed comparisons for BirdNET and Perch 2.0: mean-profile → nearest-clip known-correct acceptance, followed by the actual stranger acceptance rates for the same two scorers. All values are medians [allocation minima–maxima], using the original 16 allocations and 10% calibration budget. Table S15 retains all original mean-profile results; the complete two-scorer aggregates for all 36 distinct networks are in the v10 evidence and public repository. The numerically identical WavLM Base+ speaker-verification duplicate is retained in the archive but excluded from the network count.

Great tit, 16 birds, birdnet-v2.4: known 0.175 [0.050–0.350] → 0.250 [0.025–0.400]; strangers 0.087 [0.000–0.350] → 0.087 [0.000–0.250].

Great tit, 16 birds, perch-v2: known 0.225 [0.025–0.350] → 0.300 [0.175–0.500]; strangers 0.087 [0.000–0.275] → 0.100 [0.000–0.300].

Great tit, 50 birds, birdnet-v2.4: known 0.092 [0.015–0.177] → 0.144 [0.050–0.267]; strangers 0.092 [0.025–0.275] → 0.107 [0.017–0.269].

Great tit, 50 birds, perch-v2: known 0.112 [0.015–0.250] → 0.216 [0.085–0.292]; strangers 0.103 [0.025–0.277] → 0.096 [0.017–0.246].

Little owl, birdnet-v2.4: known 0.200 [0.000–0.520] → 0.229 [0.000–0.650]; strangers 0.084 [0.000–0.697] → 0.074 [0.000–0.679].

Little owl, perch-v2: known 0.374 [0.000–0.500] → 0.462 [0.000–0.596]; strangers 0.100 [0.000–0.587] → 0.099 [0.000–0.615].

Rook, birdnet-v2.4: known 0.080 [0.000–0.234] → 0.037 [0.002–0.172]; strangers 0.100 [0.000–0.503] → 0.099 [0.009–0.277].

Rook, perch-v2: known 0.101 [0.000–0.291] → 0.133 [0.046–0.264]; strangers 0.100 [0.000–0.403] → 0.101 [0.026–0.229].

Cockatoo, published random fold, birdnet-v2.4: known 0.741 [0.029–0.921] → 0.863 [0.000–1.000]; strangers 0.082 [0.000–0.584] → 0.086 [0.000–0.636].

Cockatoo, published random fold, perch-v2: known 0.814 [0.029–0.941] → 0.883 [0.088–1.000]; strangers 0.085 [0.000–0.632] → 0.077 [0.000–0.781].

Calibration size was assessed on the 50-bird great-tit endpoint using nested subsets of 4, 8 and 12 calibration strangers selected by fixed sorted identity order within the original calibration role. Galleries, calibration-known birds and both test roles stayed unchanged. The original allocation sometimes had 13 calibration strangers; the 12-stranger variant is therefore not always identical to the original. This is a controlled sensitivity check conditional on these subsets, not a general learning curve over randomly sampled calibration sets.

birdnet-v2.4, mean-profile, 4 calibration strangers: known 0.075 [0.000–0.158]; strangers 0.079 [0.008–0.275]; 5/16 allocations exceeded the 10% test-stranger budget.

birdnet-v2.4, mean-profile, 8 calibration strangers: known 0.071 [0.015–0.175]; strangers 0.075 [0.008–0.333]; 5/16 allocations exceeded the 10% test-stranger budget.

birdnet-v2.4, mean-profile, 12 calibration strangers: known 0.084 [0.015–0.154]; strangers 0.092 [0.025–0.250]; 7/16 allocations exceeded the 10% test-stranger budget.

birdnet-v2.4, nearest-clip, 4 calibration strangers: known 0.136 [0.023–0.308]; strangers 0.096 [0.000–0.269]; 7/16 allocations exceeded the 10% test-stranger budget.

birdnet-v2.4, nearest-clip, 8 calibration strangers: known 0.128 [0.038–0.308]; strangers 0.067 [0.000–0.269]; 5/16 allocations exceeded the 10% test-stranger budget.

birdnet-v2.4, nearest-clip, 12 calibration strangers: known 0.142 [0.050–0.267]; strangers 0.103 [0.017–0.269]; 8/16 allocations exceeded the 10% test-stranger budget.

perch-v2, mean-profile, 4 calibration strangers: known 0.109 [0.015–0.250]; strangers 0.068 [0.000–0.308]; 5/16 allocations exceeded the 10% test-stranger budget.

perch-v2, mean-profile, 8 calibration strangers: known 0.115 [0.015–0.250]; strangers 0.107 [0.000–0.283]; 8/16 allocations exceeded the 10% test-stranger budget.

perch-v2, mean-profile, 12 calibration strangers: known 0.112 [0.023–0.250]; strangers 0.103 [0.025–0.277]; 8/16 allocations exceeded the 10% test-stranger budget.

perch-v2, nearest-clip, 4 calibration strangers: known 0.204 [0.108–0.383]; strangers 0.092 [0.000–0.369]; 7/16 allocations exceeded the 10% test-stranger budget.

perch-v2, nearest-clip, 8 calibration strangers: known 0.212 [0.108–0.308]; strangers 0.088 [0.017–0.215]; 7/16 allocations exceeded the 10% test-stranger budget.

perch-v2, nearest-clip, 12 calibration strangers: known 0.216 [0.085–0.292]; strangers 0.092 [0.017–0.246]; 6/16 allocations exceeded the 10% test-stranger budget.

Increasing calibration size did not yield a monotonic improvement. For BirdNET with mean profiles, median known-correct acceptance was 0.075, 0.071 and 0.084 with 4, 8 and 12 strangers; for Perch 2.0 with nearest-clip scoring it was 0.204, 0.212 and 0.216. Test-stranger rates still exceeded the calibration target in some allocations at every size. Small calibration sets are a plausible source of variability but this check does not show that enlarging them resolves transfer between strangers.

##### S2.5 Timing of paired clips

Using retained BirdNET test embeddings, we enumerated all unordered same-individual/different-recording pairs and different-individual/same-recording pairs. Dates came from the original recording metadata or dated BirdPark recording names. We report cosine means for same-day, 1–7-day and longer gaps. Pairs share clips and individuals; these are descriptive aggregates, not independent biological replicates or additional significance tests. The original sampled-pair error rates are unchanged.

BirdPark group of four: animal 1 to 7 days: 0 pairs, mean not estimable; animal all: 751 pairs, mean 0.818; animal over 7 days: 751 pairs, mean 0.818; animal same day: 0 pairs, mean not estimable; recording: 3,953 pairs, mean 0.657.

BirdPark group of eight: animal 1 to 7 days: 0 pairs, mean not estimable; animal all: 418 pairs, mean 0.726; animal over 7 days: 418 pairs, mean 0.726; animal same day: 0 pairs, mean not estimable; recording: 6,577 pairs, mean 0.729.

Rook: animal 1 to 7 days: 205,017 pairs, mean 0.765; animal all: 1,168,249 pairs, mean 0.764; animal over 7 days: 719,210 pairs, mean 0.770; animal same day: 244,022 pairs, mean 0.748; recording: 56,625 pairs, mean 0.731.

All 751 and 418 eligible same-bird, different-recording pairs in the two BirdPark groups were more than seven days apart. A within-day or within-week BirdPark control therefore cannot be run with these test clips. The rook’s same-day animal mean (0.748) exceeded its recording-pair mean (0.731), but this does not match elapsed time exactly, remove other context or establish temporal invariance. The paired diagnostic can reflect both identity persistence and the interval between recordings; that qualification applies to biological interpretation of the original result.

##### S2.6 Fixed donors, interpretation and access

The added-background experiment used one fixed sorted-order donor mapping. Its bootstrap varies recipients conditional on that mapping; it does not quantify variation over donors. Donor rotation remains an unperformed robustness experiment. No new claim of donor-independent generality is made.

For users seeking better models, choose a split that matches the intended deployment, retain untouched sessions for final evaluation, and tune architecture or training diversity only on development data. A random-clip score measures performance under that clip-sharing design; raising it does not by itself demonstrate improved recognition in a new session. In open-set work, report identification, correct acceptance and actual stranger acceptance together, and calibrate each scorer separately without selecting on test outcomes.

Full aggregates and reproducible code: https://github.com/OpenXeroth/acoustic-re-id-research/tree/main/docs/review-sensitivity. The manuscript, supporting information, figures and original evidence documentation are licensed CC BY-NC 4.0 (https://creativecommons.org/licenses/by-nc/4.0/); software remains Apache-2.0. Original datasets and model weights retain their publishers' terms.

#### References cited only in the supporting information

Babu, A., Wang, C., Tjandra, A., Lakhotia, K., Xu, Q., Goyal, N., Singh, K., von Platen, P., Saraf, Y., Pino, J., Baevski, A., Conneau, A. & Auli, M. (2022) XLS-R: self-supervised cross-lingual speech representation learning at scale. Proceedings of Interspeech 2022, pp. 2278-2282. https://doi.org/10.21437/Interspeech.2022-143

Baevski, A., Zhou, Y., Mohamed, A. & Auli, M. (2020) wav2vec 2.0: a framework for self-supervised learning of speech representations. Advances in Neural Information Processing Systems, 33, 12449-12460.

Baevski, A., Hsu, W.-N., Xu, Q., Babu, A., Gu, J. & Auli, M. (2022) data2vec: a general framework for self-supervised learning in speech, vision and language. Proceedings of the 39th International Conference on Machine Learning, Proceedings of Machine Learning Research, 162, 1298-1312.

Chen, S., Wang, C., Chen, Z., Wu, Y., Liu, S., Chen, Z., Li, J., Kanda, N., Yoshioka, T., Xiao, X., Wu, J., Zhou, L., Ren, S., Qian, Y., Qian, Y., Wu, J., Zeng, M., Yu, X. & Wei, F. (2022a) WavLM: large-scale self-supervised pre-training for full stack speech processing. IEEE Journal of Selected Topics in Signal Processing, 16, 1505-1518. https://doi.org/10.1109/JSTSP.2022.3188113

Chen, S., Wu, Y., Wang, C., Chen, Z., Chen, Z., Liu, S., Wu, J., Qian, Y., Wei, F., Li, J. & Yu, X. (2022b) UniSpeech-SAT: universal speech representation learning with speaker aware pre-training. Proceedings of ICASSP 2022, pp. 6152-6156. https://doi.org/10.1109/ICASSP43922.2022.9747077

Chen, S., Wu, Y., Wang, C., Liu, S., Tompkins, D., Chen, Z., Che, W., Yu, X. & Wei, F. (2023) BEATs: audio pre-training with acoustic tokenizers. Proceedings of the 40th International Conference on Machine Learning, Proceedings of Machine Learning Research, 202, 5178-5193. https://proceedings.mlr.press/v202/chen23ag.html

Desplanques, B., Thienpondt, J. & Demuynck, K. (2020) ECAPA-TDNN: emphasized channel attention, propagation and aggregation in TDNN based speaker verification. Proceedings of Interspeech 2020, pp. 3830-3834. https://doi.org/10.21437/Interspeech.2020-2650

Ghani, B., Kalkman, V.J., Planqué, B., Vellinga, W.-P., Gill, L. & Stowell, D. (2025) Impact of transfer learning methods and dataset characteristics on generalization in birdsong classification. Scientific Reports, 15, 16273. https://doi.org/10.1038/s41598-025-00996-2

Gulati, A., Qin, J., Chiu, C.-C., Parmar, N., Zhang, Y., Yu, J., Han, W., Wang, S., Zhang, Z., Wu, Y. & Pang, R. (2020) Conformer: Convolution-augmented Transformer for Speech Recognition. Proceedings of Interspeech 2020, pp. 5036-5040. https://doi.org/10.21437/Interspeech.2020-3015

Hatch, A.O., Kajarekar, S. & Stolcke, A. (2006) Within-class covariance normalization for SVM-based speaker recognition. Proceedings of Interspeech 2006, paper 1874. https://doi.org/10.21437/Interspeech.2006-183

Heinrich, R., Rauch, L., Sick, B. & Scholz, C. (2025) AudioProtoPNet: an interpretable deep learning model for bird sound classification. Ecological Informatics, 87, 103081. https://doi.org/10.1016/j.ecoinf.2025.103081

Hershey, S., Chaudhuri, S., Ellis, D.P.W., Gemmeke, J.F., Jansen, A., Moore, R.C., Plakal, M., Platt, D., Saurous, R.A., Seybold, B., Slaney, M., Weiss, R.J. & Wilson, K. (2017) CNN architectures for large-scale audio classification. In 2017 IEEE International Conference on Acoustics, Speech and Signal Processing (ICASSP), pp. 131-135. https://doi.org/10.1109/ICASSP.2017.7952132

Hsu, W.-N., Bolte, B., Tsai, Y.-H.H., Lakhotia, K., Salakhutdinov, R. & Mohamed, A. (2021a) HuBERT: self-supervised speech representation learning by masked prediction of hidden units. IEEE/ACM Transactions on Audio, Speech, and Language Processing, 29, 3451-3460. https://doi.org/10.1109/TASLP.2021.3122291

Hsu, W.-N., Sriram, A., Baevski, A., Likhomanenko, T., Xu, Q., Pratap, V., Kahn, J., Lee, A., Collobert, R., Synnaeve, G. & Auli, M. (2021b) Robust wav2vec 2.0: analyzing domain shift in self-supervised pre-training. Proceedings of Interspeech 2021, pp. 721-725. https://doi.org/10.21437/Interspeech.2021-236

Huang, P.-Y., Xu, H., Li, J., Baevski, A., Auli, M., Galuba, W., Metze, F. & Feichtenhofer, C. (2022) Masked autoencoders that listen. Advances in Neural Information Processing Systems, 35, 28708-28720. https://doi.org/10.52202/068431-2081

Kather, V.S., Haupert, S., Ghani, B. & Stowell, D. (2026) Bacpipe: a Python package to make bioacoustic deep learning models accessible. Methods in Ecology and Evolution. https://doi.org/10.1111/2041-210X.70406

Matějka, P., Novotný, O., Plchot, O., Burget, L., Diez Sánchez, M. & Černocký, J. (2017) Analysis of score normalization in multilingual speaker recognition. In Interspeech 2017, pp. 1567-1571. https://doi.org/10.21437/Interspeech.2017-803

Moummad, I., Farrugia, N. & Serizel, R. (2024) Regularized contrastive pre-training for few-shot bioacoustic sound detection. In ICASSP 2024, IEEE International Conference on Acoustics, Speech and Signal Processing, pp. 1436-1440. https://doi.org/10.1109/ICASSP48485.2024.10446409

Moummad, I., Serizel, R., Benetos, E. & Farrugia, N. (2026) Domain-invariant representation learning of bird sounds. In ICASSP 2026, IEEE International Conference on Acoustics, Speech and Signal Processing, pp. 15237-15241. https://doi.org/10.1109/ICASSP55912.2026.11463533

Pratap, V., Tjandra, A., Shi, B., Tomasello, P., Babu, A., Kundu, S., Elkahky, A., Ni, Z., Vyas, A., Fazel-Zarandi, M., Baevski, A., Adi, Y., Zhang, X., Hsu, W.-N., Conneau, A. & Auli, M. (2024) Scaling speech technology to 1,000+ languages. Journal of Machine Learning Research, 25(97), 1-52.

Rauch, L., Heinrich, R., Moummad, I., Joly, A., Sick, B. & Scholz, C. (2025a) Can masked autoencoders also listen to birds? Transactions on Machine Learning Research. https://openreview.net/forum?id=GIBWR0Xo2J

Rauch, L., Schwinger, R., Wirth, M., Heinrich, R., Huseljic, D., Herde, M., Lange, J., Kahl, S., Sick, B., Tomforde, S. & Scholz, C. (2025b) BirdSet: a large-scale dataset for audio classification in avian bioacoustics. In International Conference on Learning Representations (ICLR 2025), pp. 29482-29520.

Ravanelli, M., Parcollet, T., Plantinga, P., Rouhe, A., Cornell, S., Lugosch, L., Subakan, C., Dawalatabad, N., Heba, A., Zhong, J., Chou, J.-C., Yeh, S.-L., Fu, S.-W., Liao, C.-F., Rastorgueva, E., Grondin, F., Aris, W., Na, H., Gao, Y., De Mori, R. & Bengio, Y. (2021) SpeechBrain: a general-purpose speech toolkit. arXiv:2106.04624. https://doi.org/10.48550/arXiv.2106.04624

Robinson, D., Robinson, A. & Akrapongpisak, L. (2024) Transferable models for bioacoustics with human language supervision. In ICASSP 2024, IEEE International Conference on Acoustics, Speech and Signal Processing, pp. 1316-1320. https://doi.org/10.1109/ICASSP48485.2024.10447250

Robinson, D., Miron, M., Hagiwara, M. & Pietquin, O. (2025) NatureLM-audio: an audio-language foundation model for bioacoustics. In International Conference on Learning Representations (ICLR 2025), pp. 21378-21398.

Snyder, D., Garcia-Romero, D., Sell, G., Povey, D. & Khudanpur, S. (2018) X-vectors: robust DNN embeddings for speaker recognition. Proceedings of ICASSP 2018, pp. 5329-5333. https://doi.org/10.1109/ICASSP.2018.8461375

Villalba, J., Chen, N., Snyder, D., Garcia-Romero, D., McCree, A., Sell, G., Borgstrom, J., García-Perera, L.P., Richardson, F., Dehak, R., Torres-Carrasquillo, P.A. & Dehak, N. (2020) State-of-the-art speaker recognition with neural network embeddings in NIST SRE18 and Speakers in the Wild evaluations. Computer Speech & Language, 60, 101026. https://doi.org/10.1016/j.csl.2019.101026

Wang, C., Tang, Y., Ma, X., Wu, A., Okhonko, D. & Pino, J. (2020) Fairseq S2T: fast speech-to-text modeling with fairseq. Proceedings of the 1st Conference of the Asia-Pacific Chapter of the Association for Computational Linguistics and the 10th International Joint Conference on Natural Language Processing: System Demonstrations, pp. 33-39. https://doi.org/10.18653/v1/2020.aacl-demo.6

Williams, B., van Merriënboer, B., Dumoulin, V., Hamer, J., Fleishman, A.B., McKown, M., Munger, J., Rice, A.N., Lillis, A., White, C., Hobbs, C., Razak, T., Curnick, D., Jones, K.E. & Denton, T. (2025) Using tropical reef, bird and unrelated sounds for superior transfer learning in marine bioacoustics. Philosophical Transactions of the Royal Society B, 380(1928), 20240280. https://doi.org/10.1098/rstb.2024.0280
