## Supplementary material for "Recording context and data splitting can inflate acoustic re-identification accuracy across species and studies": Graphical abstract

### Is acoustic re-identification recognising the animal, or the recording?

Four diagnostics applied to 13 public datasets from nine species and seven studies, with BirdNET and 35 other neural networks

As commonly tested | Under the diagnostic

THE PROBLEM

#### Two common practices can inflate reported re-ID accuracy

##### a. Each animal is recorded at its own site

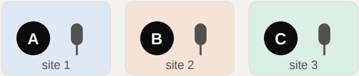


Identity and recording site are linked, so a classifier can score highly by recognising the place.

##### b. Clips are split at random

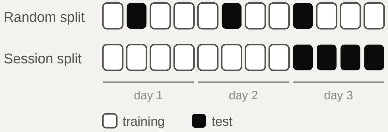


Training and test clips share sessions, so the score no longer measures recognising an animal in a new session.

#### 1 Split by recording session

Does accuracy hold in a new recording session?

Little owl, BirdNET, same clips and per-bird counts

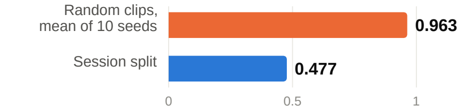


Mixing clips from the same sessions into training and test raised accuracy by 0.20 to 0.51 in 18 comparisons with matched clips, counts and models (three species).

#### 2 Take the animal away

Can ambient sound alone identify individuals?

Chiffchaff within one year, BirdNET, balanced accuracy

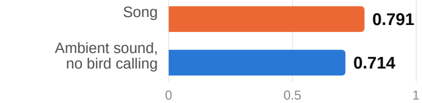


Background alone identified the birds for 35 of 38 embeddings and controls.

#### 3 Hold the recording fixed

Where animals share a microphone, does re-ID survive?

| | | |
|---|---|---|
| ✓ | Zebra finches, group of 4 | **identity signal beyond recording** |
| ? | Zebra finches, group of 8 | recording not ruled out |
| ? | Rooks, one aviary | recording not ruled out |

Only one dataset gave clear evidence beyond recording. Both finch groups lack cross-recording same-bird pairs within seven days; changes over time remain a confound.

#### 4 Let strangers in

Are known animals still named when others call?

Little owl · Perch 2.0 · nearest clip · four enrolled birds

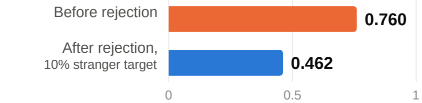


Highest median in wild session-split tests: 46%, across 36 networks and two scorers; 16 role allocations.

Known calls accepted and correctly named; stranger rates can exceed 10%.

**Accuracy reported without these four diagnostics can substantially overstate acoustic re-ID.**

No new models are needed; each diagnostic requires suitable data. Test before planning population monitoring.

Wallington & Lighten (2026), author manuscript v10. CC BY-NC 4.0
